# NPlasmid-seq: Cas9-encoded terminal signatures enable molecule-level nanopore quality control of vector and IVT-template plasmids

**DOI:** 10.64898/2026.11.02.763549

**Authors:** Yan Xu, Wei Wen, Di Sun, Yi-Xue Yin, Xiao-Yu Ma, Jia-Xin Ren, Shu-Man Zhang, Jian-Qiao Chang, Shan-Chen Pang, Shu-Dong Wang, Xiao-Bing Zhang, Jian-Ping Zhang

**Author notes:** Corresponding authors., (Zhang JP); (Zhang XB). Equal contribution (Yan Xu, Wei Wen, Di Sun).

## Abstract

Consensus-based plasmid quality control can overlook structural heterogeneity within a preparation. We developed NPlasmid-seq, a Cas9-guided nanopore workflow that encodes sample identity through paired terminal signatures generated at predefined backbone positions, without sample-specific oligonucleotide barcodes. We cleavage-validated 24 sgRNAs. In five representative assigned read bins, each bin showed its strongest sequence agreement with its cognate reference. Contaminant detection was evaluated with a single class-specific classifier applied both to computational read mixtures and to physical DNA mixtures, placing the two designs on a common scale. In computational read mixtures of 5,000 reads, a contaminant was detected in all 20 independent subsamples at each of 0.2%, 0.5%, 1% and 5% abundance and in 13 of 20 subsamples at 0.1%; precision was 1.000 in all 100 subsamples, with no false-positive calls among 493,200 reads from the declared target plasmid. In physical DNA mixtures carried through cleavage, pooled library preparation and sequencing, a contaminant was detected in all six samples prepared at 0.5 mol% input and all six at 1 mol%. Observed read fractions did not distinguish the 0.5 and 1 mol% input levels. Instead, they differed systematically by 4.0-fold between preparation pools, a larger effect than the difference between the two input levels. We therefore interpret the workflow as a detector, not a quantifier, of sub-percent contamination. Read-level analysis identified a recurrent 93-bp deletion in 61.7% of reads from a preparation that had passed consensus-based quality control. Sanger genotyping of 24 colonies identified deletion-type profiles in 10 colonies and intact-type profiles in three. The remaining 11 colonies showed double peaks at the diagnostic locus, consistent with coexistence of both sequence forms within individual clones. In in vitro transcription template plasmids, longer poly(A) tracts were associated with higher multimeric-read fractions. Across ten matched A90 culture pairs, propagation at 30 °C reduced these fractions relative to 37 °C (paired t-test, P = 0.0098). These measurements describe recovered read populations, not absolute molecular abundances. NPlasmid-seq therefore complements consensus validation by revealing sequence and structural heterogeneity within a preparation and by identifying a culture condition associated with a lower multimeric-read fraction.

## Introduction

Recombinant plasmids are foundational tools in molecular biology and important starting materials for an expanding range of biopharmaceutical applications, including adeno-associated virus (AAV) and lentiviral vector production and the generation of in vitro transcription (IVT) templates for mRNA therapeutics and vaccines(1, 2). In these applications, sequence errors, contaminating plasmids, and structural rearrangements introduced during cloning or bacterial propagation can compromise experimental interpretation, manufacturing consistency, or final-product quality.

The dominant quality-control paradigm remains consensus sequencing. Sanger sequencing is accessible and inexpensive but typically reads only several hundred to approximately one thousand bases per reaction, requires primer walking for complete coverage, and performs poorly in GC-rich, repetitive, or highly structured regions(3). Short-read sequencing can cover an entire plasmid but requires fragmentation and reassembly, may not uniquely resolve shared backbone elements, and is not always practical for routine validation of small construct sets(4). Nanopore sequencing has changed this landscape because individual reads can span complete plasmids, while modern chemistries and polishing workflows can generate highly accurate consensus sequences(5, 6). Workflows such as OnRamp have further enabled pooled, amplification-free plasmid validation at a cost per kilobase below that of Sanger sequencing(4).

Most routine plasmid-sequencing workflows nevertheless emphasize a consensus verdict(7). A preparation is considered correct when the consensus matches the intended design. Consensus sequencing can confirm the intended sequence but does not reveal what fraction of individual molecules retains the expected end-to-end structure. A plasmid preparation is a population of molecules. Repetitive, GC-rich, or homopolymeric inserts can promote deletions through replication slippage or recombination(8), while high-copy-number plasmids can accumulate multimers(9–11). A preparation containing a large deletion-bearing or multimeric subpopulation may therefore yield an apparently correct base-by-base consensus while remaining structurally heterogeneous at the molecule level.

Nanopore reads can, in principle, address this population-level question because each read originates from a physical DNA molecule. A practical multiplexed workflow, however, requires each full-length read to be assigned to its source plasmid without random fragmentation. Oligonucleotide barcodes add cost and can be misassigned, transposase-based rapid kits fragment DNA, and restriction-based linearization requires a suitable site in every construct(4). Cas9 ribonucleoprotein (RNP) cleavage provides a programmable alternative. When a plasmid is linearized at a selected backbone coordinate, the resulting pair of terminal sequence contexts becomes a physical molecular address that can be read directly during nanopore sequencing(12, 13).

Here, we developed NPlasmid-seq, a wet-laboratory and computational workflow for molecule-level plasmid quality control. We evaluated terminal-signature demultiplexing in pooled plasmids spanning 2.40–37.62 kb (Table S1) using 24 validated backbone-targeting sgRNAs (Table S2). We then applied the workflow to three questions that consensus sequences alone cannot answer: whether low-abundance contaminants can be detected, how frequently structural variants appear among recovered reads, and whether multimeric IVT-template plasmids can be detected. For each application, we paired a computational or read-level analysis with a complementary laboratory test. Contaminant detection was evaluated in both computational read mixtures and physical DNA mixtures prepared before Cas9 treatment. The read-level deletion estimate was compared with clone genotyping, and a multimeric species detected in sequencing reads was isolated from a gel and identified as a tandem plasmid.

## Methods

### Plasmid propagation, extraction, and quality control

Plasmids ranging from 2.40 to 37.62 kb (Table S1; Supplementary File 1) were propagated in a recombination-and endonuclease-deficient Escherichia coli strain (TransGen Biotech, Beijing, China). Single colonies were inoculated into 2 mL of LB medium containing the appropriate selective antibiotic and cultured for 4–6 h at 37 °C with shaking at 220 rpm. Seed cultures were diluted 1:1,000 into 50 mL of the same medium and grown for 12–16 h at 37 °C with shaking at 220 rpm. Plasmid DNA was extracted with the ZymoPURE II Midiprep Kit (Zymo Research) according to the manufacturer’s instructions and eluted in 10 mM Tris-HCl (pH 8.5). DNA concentration was measured with the Qubit dsDNA HS Assay, and purity and gross integrity were assessed by NanoDrop spectrophotometry and 1% agarose gel electrophoresis, respectively.

Preparations were accepted for Cas9 treatment if the A260/A280 ratio was 1.80–2.00, the A260/A230 ratio was at least 1.80, and agarose gel electrophoresis showed a predominant supercoiled band without visible genomic DNA contamination or smearing indicative of degradation. Preparations that did not meet these criteria were discarded and remade.

### Temperature-controlled propagation of poly(A)-containing IVT plasmids

To evaluate the effect of propagation temperature on poly(A)-containing IVT plasmids, matched cultures derived from the same seed culture were diluted 1:1,000 into 50 mL of selective LB medium and incubated at either 30 °C or 37 °C with shaking at 220 rpm. Medium composition, inoculation ratio, antibiotic concentration, culture volume, and harvest criterion were held constant. Plasmid DNA was extracted and quality-controlled as described above. Ten matched culture pairs were analysed for the A90 temperature comparison.

### sgRNA design and Cas9 RNP assembly

Candidate sgRNAs were designed against backbone regions shared among plasmids in each sequencing pool(12). The predicted Streptococcus pyogenes Cas9 cleavage coordinate was defined as 3 bp upstream of an NGG protospacer-adjacent motif. Guides were selected so that (i) cleavage sites were located in nonoverlapping backbone neighborhoods and outside annotated functional elements; (ii) cleavage coordinates assigned to different plasmids were separated by at least 200 bp, preventing overlap between the ±50-bp signature windows; and (iii) no additional protospacer with two or fewer mismatches and a compatible PAM was detected elsewhere in any plasmid in the corresponding pool. Each plasmid was assigned one guide that generated a distinct terminal-sequence context after linearization. Spacer lengths of 18 to 20 nt were permitted by design and are listed in Table S2.

For each reaction, 3 pmol S. pyogenes Cas9 protein (V3) was mixed with 6 pmol sgRNA and incubated for 10 min at room temperature. The assembled RNP was added to 200 ng plasmid DNA in a final volume of 10 µL, incubated at 37 °C for 4 h, and heat-inactivated at 65 °C for 10 min. Cleavage was assessed by 1% agarose gel electrophoresis, and reactions in which the expected linearized band predominated over residual circular forms were pooled for library preparation.

### Partial Cas9 cleavage for multimer analysis

To retain covalently linked multimers while generating sequenceable DNA ends, poly(A)-containing IVT plasmids were subjected to a shortened Cas9 reaction. RNPs were assembled as above, added to 200 ng plasmid DNA in a final volume of 10 µL, incubated for 1 h at 37 °C, and heat-inactivated for 10 min at 65 °C.

### Contaminant detection in physical DNA mixtures

To test contaminant detection through the complete laboratory workflow rather than after sequencing alone, purified plasmids were mixed before Cas9 treatment. P7 (7.02 kb) served as the declared target and P6 (5.74 kb) as the contaminant; both had independently verified reference sequences and both were also used in the computational benchmark, allowing the two designs to be compared directly.

Concentrations were measured in triplicate by Qubit dsDNA HS assay and converted to molar concentration using an average double-stranded DNA mass of 650 g/mol per base pair. Mixtures were prepared at 0, 0.5, 1, 5, 50 and 100 mol% P6; because the two plasmids differ in length, mass ratios were calculated from molar ratios rather than assumed to be equal (for example, 1 mol% P6 corresponds to 0.82 mass% and to 1.64 ng of P6 in a 200 ng reaction). Working solutions were prepared in 10 mM Tris-HCl (pH 8.5) in DNA LoBind tubes so that no single pipetting step was below 1.5 µL.

The replicate structure is defined as follows and is reported in full in Table S5. A total of 26 mixture samples were analysed, distributed across three pools: Pool 1 and Pool 2 each contained samples V01-V09 at 0, 0.5 and 1 mol% P6, and Pool 3 contained samples V10-V17 at 5, 50 and 100 mol% P6. Each final P6/P7 mixture was prepared separately in its own tube; no mixture was split into aliquots. Within each pool and input level, the three mixtures were prepared separately from the same upstream P6 working solution. They therefore represent separate mixture preparations, but not independent dilutions from the P6 stock. The two pools were prepared using different upstream working solutions. Each tube received a different cleavage guide, and the same three guides were used at a given input level in both pools, so pool and guide are crossed. The 0.5 mol% and 1 mol% conditions were each represented by six samples (three per pool), the 5 mol% and 50 mol% conditions by three samples each, the 0 mol% condition by six and the 100 mol% condition by two. Because the upstream working solution is shared within a pool, the pool rather than the individual tube is the level at which preparation error is shared, and results are reported accordingly.

Each mixture (200 ng total DNA) was cleaved in a single reaction with one guide whose target site and flanking ±50-bp windows are identical in both plasmids, so that target and contaminant received identical ends and identical exposure to the cleavage reaction. Reactions were incubated at 37 °C for 4 h, heat-inactivated, treated with Proteinase K (20 mg/mL, 37 °C, 15 min), purified with 0.5x AMPure XP beads and requantified. Different mixtures were assigned different backbone cleavage addresses, so that mixtures could be pooled into a single library and separated by terminal-signature demultiplexing without oligonucleotide barcodes; within a mixture, target and contaminant were indistinguishable by their ends. Single-plasmid controls were pooled, prepared and sequenced within the same libraries as the mixtures they control for: the P7-only (0 mol%) controls were pooled with the 0.5 and 1 mol% mixtures, and the P6-only (100 mol%) controls with the 5 and 50 mol% mixtures. Libraries were prepared with SQK-LSK114 and sequenced on R10.4.1 flow cells (FLO-MIN114) for 16 h, longer than the 6–8 h used for routine quality control, to reach the read depth required at 0.5% abundance.

### Single-colony and subcolony genotyping of the P9 deletion

The glycerol stock used to generate the sequenced P9 preparation was streaked onto LB agar containing the appropriate antibiotic and incubated overnight at 37 °C. Twenty-four well-separated colonies were selected for Sanger sequencing across the 93-bp deletion locus within the rabbit SOX9 region. Chromatograms were compared with the intact P9 reference and the expected deletion-junction sequence. Colonies were classified as deletion-type when interpretable sequence spanned the deletion junction, or intact-type when sequence supported retention of the affected region. Chromatograms showing overlapping peaks at the diagnostic locus were recorded separately as double-peak profiles, which indicate that both sequence forms were present in the material derived from that colony. Counts and percentages were calculated using all 24 colonies as the denominator (Fig. 3E, Table S6).

To test whether the deletion phenotype emerged during further propagation of an intact-type parent, one parent classified as intact by Sanger sequencing was streak-plated. Eight isolated subcolonies were selected for colony PCR (Fig. 3F). Each subcolony was lightly touched with a sterile tip and suspended in 20 µl of water; 1 µl of the suspension was used as the PCR template.

The deletion locus was amplified using forward primer 5′-GACCTTCGACGTCAACGAGT-3′ and reverse primer 5′-AGGTGGAGTAGAGGCTGGAG-3′ (Table S3). Expected amplicon sizes were 548 bp for the intact allele and 455 bp for the deletion allele. PCR was performed in 10-µl reactions using KAPA HiFi, 5% DMSO (v/v) and 0.5 µM of each primer. Cycling comprised an initial denaturation at 98 °C for 2 min, followed by 30 cycles of 98 °C for 5 s, 64 °C for 5 s, 68 °C for 5 s and 72 °C for 15 s. Products were resolved on a 1.2% agarose gel at 110 V for approximately 30 min. Subcolonies were scored by the presence of the expected intact or deletion amplicon; band intensity was not used to estimate allele proportions.

### Pooling and nanopore library preparation

After Cas9 linearization, plasmids were pooled using length-adjusted DNA inputs to reduce underrepresentation of larger constructs. For a pool containing a total DNA mass m, the input mass of plasmid i was calculated as mi proportional to Li^1.5, where Li is the length of plasmid i. Larger plasmids were therefore included at proportionally higher mass. This empirical weighting was applied to pools containing 12–24 plasmids to improve read-depth balance across plasmids of different sizes. For the physical-mixture pools, in which all members were of similar length, equal masses were used.

The pooled linearized DNA was prepared with the Ligation Sequencing Kit V14 (SQK-LSK114; Oxford Nanopore Technologies) according to the manufacturer’s protocol(14, 15). No sample-specific oligonucleotide barcodes were introduced. DNA underwent end repair and dA-tailing, adapter ligation, and Long Fragment Buffer washes before final elution and Qubit quantification.

### Sequencing and FAST/HAC/SUP basecalling

The final sequencing library was combined with sequencing buffer and loading beads to a total loading volume of approximately 75 µL, loaded onto a primed R10.4.1 flow cell (FLO-MIN114; Oxford Nanopore Technologies), and sequenced on a MinION Mk1C. Raw nanopore signals were recorded in POD5 format. Routine quality-control runs were 6–8 h; the physical-mixture validation runs were extended to 16 h to reach the pre-specified read depth.

To compare the effects of basecalling accuracy, the same POD5 datasets were independently basecalled using Dorado v0.4.3 with the FAST, high-accuracy (HAC), and super-accuracy (SUP) models for R10.4.1/E8.2 400-bps chemistry: dna_r10.4.1_e8.2_400bps_fast@v4.3.0, dna_r10.4.1_e8.2_400bps_hac@v4.3.0, and dna_r10.4.1_e8.2_400bps_sup@v4.3.0, respectively. Identical input datasets and filtering parameters were used for all three basecalling modes. Sequencing adapters were trimmed using Dorado, and reads with a mean predicted Q-score below 10 were excluded(16).

Unless otherwise specified, SUP-basecalled reads were used for read assignment, alignment, consensus generation, and structural analysis. Reads lacking the expected terminal signatures or carrying conflicting signature combinations were retained for failure-mode auditing but excluded from confident sample assignment and consensus reconstruction.

### End-signature generation and dual-end demultiplexing

For each plasmid reference, 20-nt probes with a 7-nt stride were generated across the ±50-bp region flanking the predicted Cas9 cleavage site. For confident sample assignment, a read was required to contain perfect matches to the expected 5′ and 3′ terminal-signature probe sets within the prespecified terminal windows. Reads with only one matching terminus or conflicting signatures for multiple plasmids were classified as unassigned and excluded from consensus reconstruction. Assigned reads were independently cross-mapped to all plasmid references with BLASTN v2.16.0+(17). A fit index, defined as fractional identity multiplied by fractional coverage, was calculated for each read-bin/reference pair.

Assignment outcomes were tabulated for every pooled run as four mutually exclusive categories summing to the total number of passing reads: concordant dual-end signatures (assigned), a match at only one terminus, conflicting dual-end signatures, and no signature match. Absolute counts and percentages are reported in Table S4. The fit index describes how well an assigned read bin matches its cognate reference and is not a measure of assignment accuracy, because the same terminal-signature rule generates both the assignment and the bin under test.

### Two-layer contaminant classification

Contaminant classification within a mixture proceeds in two independent layers, which are reported separately throughout. The first layer is terminal-signature demultiplexing, which assigns a read to a mixture (that is, to a cleavage address) using the dual-end rule described above with a minimum mean read quality of Q10, a 50-bp scan window at each read terminus, and 20-nt probes at a 7-nt stride. The second layer is class-specific sequence classification, which assigns an already-demultiplexed read to the target or the contaminant plasmid using sequence intervals that are unique to one of the two plasmids. Reads for which the class-specific evidence does not reach the prespecified requirement are labelled ambiguous. Ambiguous reads are reported as a separate category and are not counted as misclassifications. The same two-layer procedure and the same class-specific classifier (v1.0.1) were applied to computational and physical mixtures. As an implementation check, the physical-mixture reads were also classified with an independent two-class implementation of the same decision rule; this returned identical P6, P7 and ambiguous counts for all 26 samples.

### Detection endpoint and performance metrics

Because the intended application is quality control, the primary endpoint is detection rather than per-read classification accuracy. We set the detection threshold at three contaminant-class reads on the basis of the observed false-call rate. No false-positive call occurred among 493,200 reads from the declared target plasmid across the 100 computational subsamples, giving a one-sided 95% upper bound of 6.1 × 10□□ per read. At this upper-bound rate, fewer than 0.03 false calls are expected per 5,000-read sample, and the probability of three or more false calls is below 10□□. Detection rate is reported as the number of samples or subsamples in which the contaminant was detected over the number tested, with the denominator stated.

Precision, defined as the fraction of reads called as contaminant that truly originate from the contaminant, is reported as the primary measure of classification trustworthiness, because a false alarm in quality control triggers unnecessary rejection of a preparation. Recall is reported as a depth-dependent sensitivity parameter rather than as a performance ceiling, because reads with insufficient class-specific evidence are conservatively labelled ambiguous by design. F1 is reported in Table S7 as a secondary summary statistic to permit comparison with reference-first decision rules; F1 values obtained under different decision rules and different ambiguous-read conventions are not interchangeable and are not pooled.

Detection rate and precision are computable for both the computational and the physical mixtures, whereas recall and F1 require per-read ground truth and are therefore available only for the computational mixtures. Reporting detection rate and precision as the primary endpoints places the two designs on a common scale.

### Computational mixture benchmark

Computational benchmark datasets were generated by combining reads from independently sequenced plasmid datasets at prespecified contaminant abundances. No plasmid DNA was mixed in the laboratory for these datasets; they therefore test classification after sequencing but not co-cleavage, co-library preparation or co-sequencing. Each simulated sample contained a fixed total of 5,000 passing reads, and each abundance level was represented by 20 independent random subsamples. Contaminant abundances of 0.1%, 0.2%, 0.5%, 1% and 5% were evaluated. Classification used the class-specific classifier (v1.0.1) and the ambiguous-read convention described above, identical to the classifier applied to the physical mixtures. Per-subsample results, including recall and F1, are reported in Table S7.

The classifier assigns reads using region-alignment criteria over class-specific intervals and does not expose a diagnostic-probe-hit count as a tunable threshold; no operating-point sweep was therefore performed, and the classifier was applied with its released parameters throughout (Fig. 2B). Alignment anomalies caused by partial-length or internally deleted reads were manually audited in IGV.

An earlier version of this benchmark used a reference-first decision rule in which reads were flagged when alignment to the declared reference had identity below 95% or coverage below 90%. That rule differs from the class-specific classifier in both its decision criterion and its treatment of reads with incomplete evidence, and the two are therefore not reported together. All computational-mixture results presented here were generated with the class-specific classifier, and the corresponding analysis code is provided in the repository cited under Code and data availability.

### Near-identical plasmid classification

For near-identical plasmids (differing by approximately 20 bp), reads were scanned across known variant loci using a 20-nt sliding window; a read was classified as a contaminant if at least 9 substitutions occurred within any window, provided the local mean base quality was at least Q18. To exclude chimeric artifacts, reads were required to have concordant dual-end signatures matching the declared backbone.

### poly(A)-associated multimer classification

Multimeric plasmid forms were identified from concordant read-alignment and electrophoretic evidence. Candidate multimeric reads contained at least two tandem, reference-length alignment units or a prespecified plasmid-plasmid junction verified in IGV. Agarose gels were used as an orthogonal qualitative assay for bands with an apparent size near twice that of the linear monomer(22). Reads were classified as monomer, dimer, or at least trimer. Because recovery and sequencing efficiency can vary with DNA length, the reported quantity is termed the multimeric-read fraction rather than the molecular or molar multimer fraction. For each multimeric read, the relative orientation of the two tandem units and the coordinate of the plasmid-plasmid junction relative to the poly(A) tract were recorded, so that the read-level structure could be compared with the gel-based identity test.

### Signal-assisted estimation of poly(A) length

poly(A) tract length was estimated by integrating raw nanopore signal with basecalled sequence. For each read, a scaling factor (signal points per base) was computed as the ratio of total raw signal points to FASTQ read length. Candidate poly(A) start sites were located in the basecalled sequence by fuzzy matching of flanking conserved motifs (up to 6 mismatches) or detection of at least 10 consecutive adenines. Each candidate position was mapped to the raw signal by multiplying the start position by the scaling factor, and a 3,000-point window centred there was extracted. Local variance was computed with a sliding window (size 50 points, step 20 points); the poly(A) region was defined as the longest contiguous low-variance segment, reflecting the low signal variance characteristic of a homopolymer, with boundaries refined at inflection points of the variance profile. poly(A) length in nucleotides was obtained by dividing the refined signal interval by the scaling factor. The resulting estimates were compared with Sanger sequencing as a qualitative orthogonal check.

### Read-integrity analysis, alignment and consensus generation

A read was classified as left-end intact, right-end intact, or dual-end intact when aligned to the reference DNA. Read integrity (%) was calculated as dual-end intact read count divided by total read count, multiplied by 100 (Fig. S8). Demultiplexed reads were aligned to plasmid references with Minimap2(18), processed with Samtools(19), and inspected in IGV v2.19.1(20). Consensus sequences were generated using three rounds of Racon followed by one round of Medaka(21).

### Statistical analysis and reproducibility

Statistical tests are specified in the figure legends and were two-sided unless otherwise stated. For sample-level inference, the independent units were biological preparations or matched culture pairs, not individual sequencing reads. For the physical mixtures, independent mixture preparations rather than individual cleavage guides were treated as the unit of replication. Computational mixtures and subsampling iterations were not treated as biological replicates. Exact P values, effect estimates, 95% confidence intervals, sample sizes, replicate definitions, data summaries, pairing structures, and multiple-comparison procedures are reported for each analysis.

Proportions derived from counts (colony genotypes, flagged-read fractions, background misassignment) are reported with Wilson 95% confidence intervals; for zero-count observations the one-sided 95% upper bound is reported using the rule of three. The matched temperature comparison is reported as the mean paired difference with a 95% confidence interval in addition to the paired t test, and a Wilcoxon signed-rank test is provided as a sensitivity analysis because normality cannot be assessed reliably at n = 10. Thresholds and analysis scripts for the physical-mixture experiment were fixed before unblinding.

### Code and data availability

The sequencing data are available through NCBI BioProject PRJNA1347537. The analysis pipeline that generated the figures presented here is publicly available at https://github.com/XBlab-Bioinfo/NPlasmid-seq.

## Results

### Cas9-generated terminal signatures encode programmable, barcode-free sample identities

NPlasmid-seq encodes sample identity through the programmed position of plasmid linearization (Fig. 1A). Cas9 ribonucleoprotein complexes carrying different backbone-targeting sgRNAs cleave plasmids at distinct coordinates, generating paired terminal signatures defined by the sequence context surrounding each cleavage site (Fig. 1B). Reads retaining both signatures can be assigned to the corresponding plasmid preparation without sample-specific oligonucleotide barcodes, while preserving information about end-to-end molecule architecture. The workflow required less than 4 h of hands-on preparation, 6–8 h of sequencing, and approximately 2 h of computational analysis (Fig. 1A).

**Figure 1.**
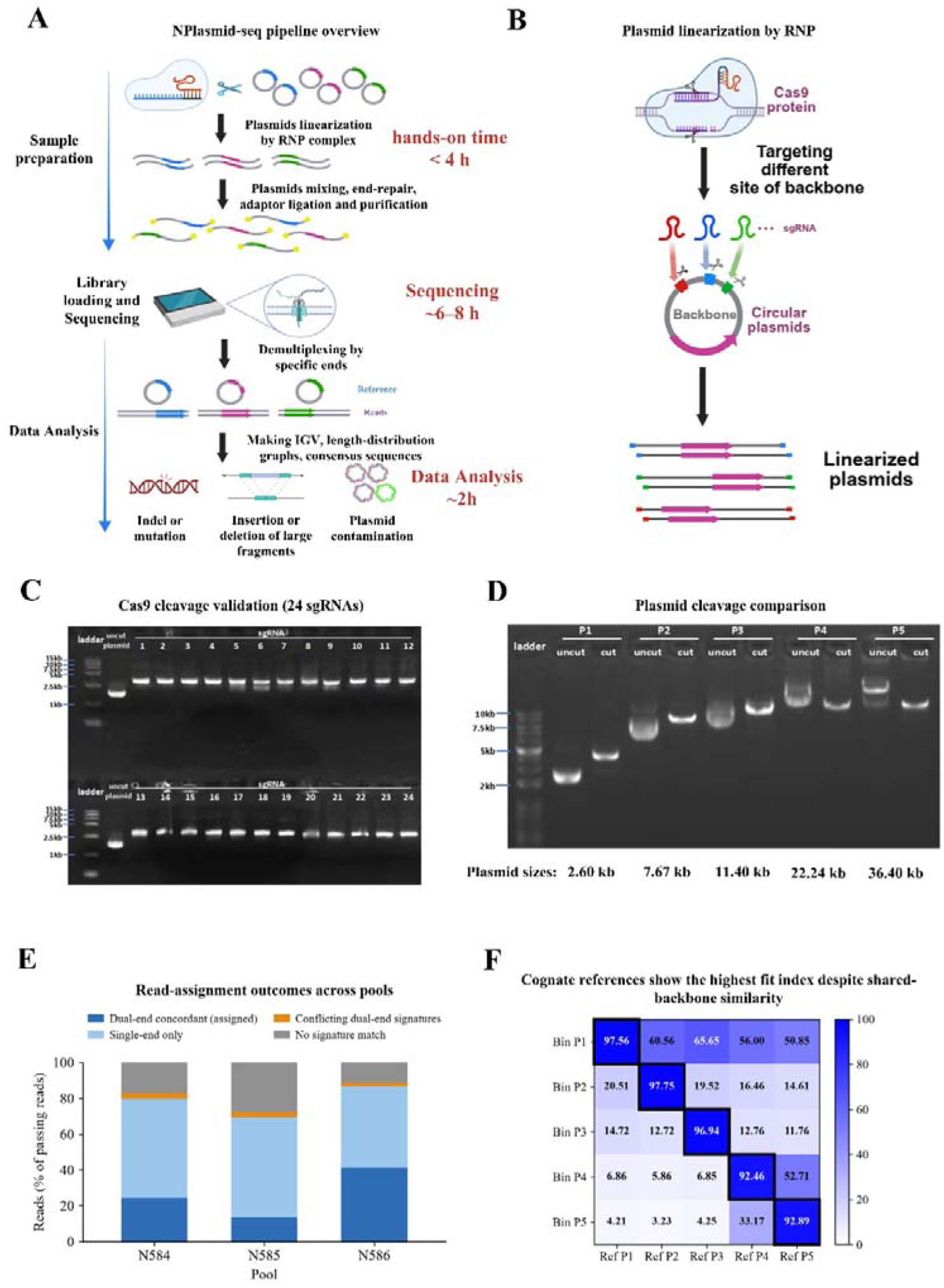
NPlasmid-seq workflow, Cas9-directed addressing and reference agreement of assigned read bins. (A) Overview of the NPlasmid-seq workflow. Plasmids are linearized at predefined backbone sites using Cas9 ribonucleoprotein (RNP) complexes, pooled, end-repaired, ligated to sequencing adapters, purified and subjected to Oxford Nanopore sequencing. Reads are demultiplexed using cleavage-derived terminal signatures and analysed to generate read-alignment profiles, read-length distributions and consensus sequences. Approximate processing times are indicated: less than 4 h of hands-on preparation, 6-8 h of sequencing and 2 h of computational analysis. (B) Principle of Cas9-directed sample addressing. Cas9 RNPs containing different sgRNAs target distinct positions within the plasmid backbone, generating linearized molecules with different pairs of terminal sequence contexts. These cleavage-derived signatures encode sample identity without sample-specific oligonucleotide barcodes. (C) Agarose gel electrophoresis of Cas9 cleavage products generated using 24 sgRNAs. Numbered lanes correspond to individual sgRNAs; ladder and untreated plasmid controls are indicated. Each guide produced a predominant band consistent with plasmid linearization. This assay validates cleavage activity rather than pooled discrimination among all 24 addresses. (D) Cas9-mediated cleavage of five plasmids spanning 2.60-36.40 kb: P1, 2.60 kb; P2, 7.67 kb; P3, 11.40 kb; P4, 22.24 kb; and P5, 36.40 kb. Paired lanes show untreated plasmids (uncut) and Cas9-treated products (cut). DNA size markers are indicated. Migration of uncut plasmids depends on topology and was not used to infer molecular length. (E) Read-assignment outcomes in a 16-plex pooled sequencing run. Mutually exclusive categories are shown as proportions of the 149,082 passing reads: concordant dual-end signatures (assigned), 29,945 reads (20.09%); single-end match only or conflicting dual-end signatures, 48,197 reads (32.33%); and no signature match, 70,940 reads (47.58%). Absolute counts are indicated for each category. Per-run values are reported in Table S4. (F) Fit-index matrix comparing reads assigned to five representative bins (P1-P5) with plasmid references Ref P1-Ref P5. Rows denote assigned bins and columns denote references. Values are percentages, calculated as fractional alignment identity x fractional reference coverage x 100; colour intensity indicates the fit index. Black outlines highlight cognate bin-reference pairs, which show the highest value in each row. Shared backbone sequences contribute to nonzero off-diagonal values. The fit index measures sequence agreement, not read-assignment accuracy or assignment yield.

**Figure 2.**
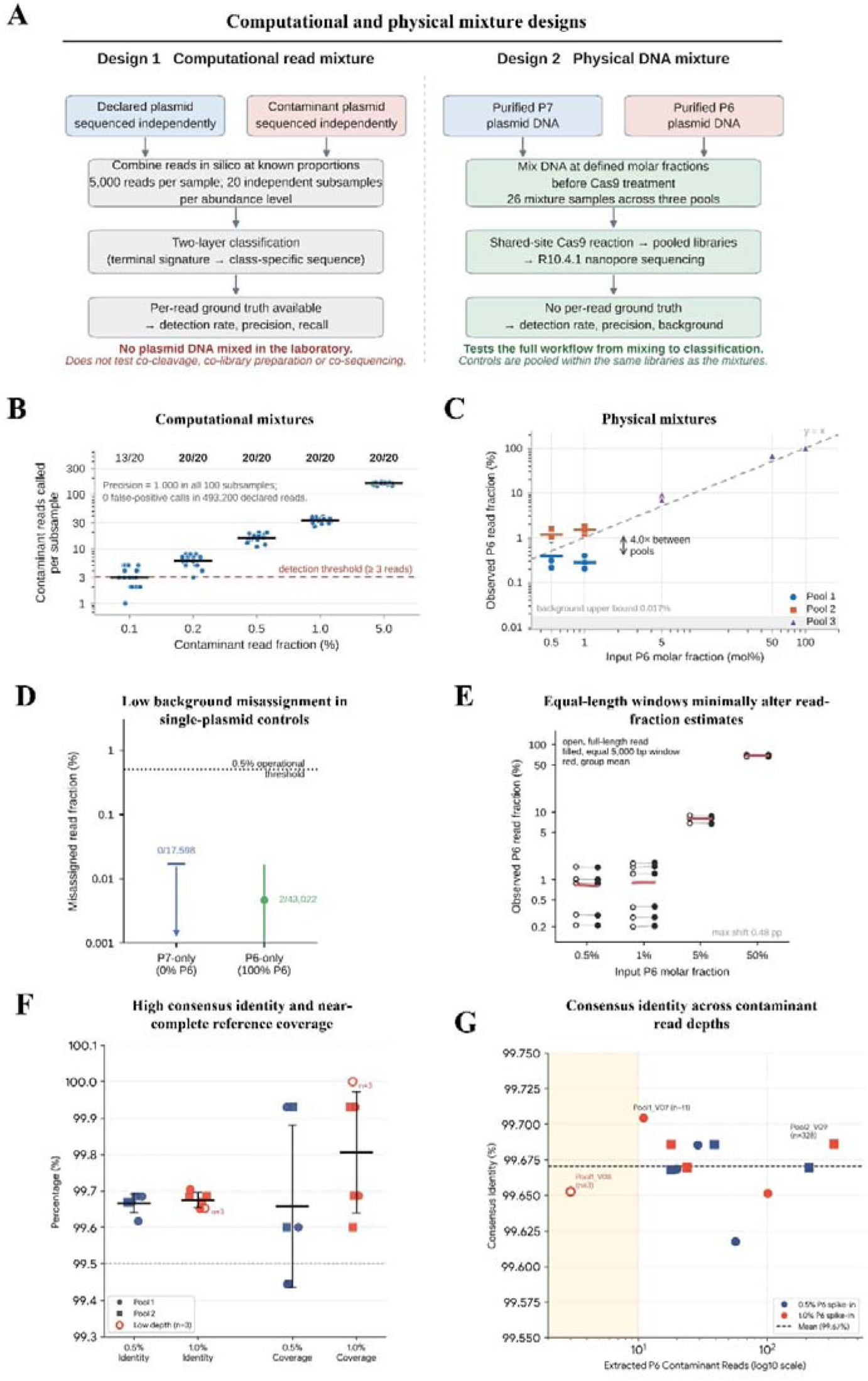
Contaminant detection in computational read mixtures and physical DNA mixtures. (A) Experimental designs. Computational read mixtures combine reads from independently sequenced plasmids at defined read fractions, testing classification after sequencing without physically mixing DNA. Physical DNA mixtures combine purified P7 and P6 DNA at defined molar fractions before Cas9 treatment, testing cleavage, pooled library preparation, sequencing and classification. The same two-layer procedure (terminal-signature demultiplexing followed by class-specific sequence classification) and the same classifier were applied to both designs. (B) Contaminant detection in computational read mixtures. Each point is one of 20 independent subsamples of 5,000 passing reads at the indicated contaminant read fraction; horizontal bars are geometric means and the y axis is logarithmic. The dashed line marks the prespecified detection threshold of at least three contaminant reads, defined in Methods from the classifier’s observed false-positive behaviour. Fractions above each level give the number of subsamples in which the contaminant was detected. Precision was 1.000 in all 100 subsamples, with no false-positive calls among 493,200 declared reads; recall and F1 for the same subsamples are reported in Table S7. The classifier uses region-alignment criteria and does not expose a tunable diagnostic-hit threshold, so no operating-point sweep is shown. (C) Observed P6 read fraction versus input P6 molar fraction in physical mixtures with P7, on logarithmic axes. Each point is one separately prepared mixture; symbol colour and shape denote the sequencing pool, and short horizontal bars are per-pool geometric means at 0.5 and 1 mol%. Tubes within a pool shared an upstream P6 working solution; the two pools used separate working solutions. At 0.5 and 1 mol% the two pools separate completely, with geometric means of 0.329% (Pool 1) and 1.323% (Pool 2), a ratio of 4.0 (Mann-Whitney P = 0.0022); within a pool the scatter is much smaller and is largely consistent with Poisson variation in contaminant read counts. The dashed diagonal indicates equality of input molar and observed read fractions (y = x) and is a reference line, not a calibration curve. The shaded band marks the background upper bound of 0.017% from the P7-only control (panel D). (D) Background read misassignment in single-plasmid controls. No P6 reads were assigned among 17,598 classified reads in the P7-only control; two of 43,022 classified reads were assigned to P7 in the P6-only control. For the zero-count control, the cap marks the one-sided 95% upper bound (0.017%, rule of three) and the downward arrow indicates that the observed value is zero. For the P6-only control the point indicates the observed fraction (0.0046%) and the error bar the Wilson 95% confidence interval (0.0013-0.0168%). The y axis is logarithmic. Each control was pooled and sequenced within the same library as the mixtures it controls for, so the measured background includes any contribution from inter-sample cross-talk under those conditions. (E) Comparison of observed P6 read fractions calculated using full-length reads (open circles) or equal 5,000-bp windows comprising the first and last 2,500 bp of each read (filled circles), across 18 mixture samples. Connecting lines link estimates from the same sample; bars indicate group means. The maximum absolute change between the two calculations was 0.478 percentage points. Both axes are logarithmic. The within-level spread at 0.5 and 1 mol% is evident in both calculations. (F) Consensus sequence identity and reference coverage for P6 contaminants recovered from physical mixtures containing 0.5 or 1 mol% P6. Symbol colour indicates input level; symbol shape distinguishes pools. The open symbol identifies Pool1_V08, reconstructed from three contaminant reads and reported as a low-depth exploratory result. Horizontal bars indicate group means and error bars indicate SEM. The dotted horizontal line marks 99.5%. Consensus identities ranged from 99.6175% to 99.7043% (mean 99.6704%). (G) Consensus sequence identity versus the number of extracted P6 contaminant reads (3-328 reads per sample), with read count shown on a logarithmic axis. Colours and symbol shapes follow panel F. Selected samples are labelled with their identifiers and contaminant read counts (n). The dashed horizontal line indicates the mean identity of 99.67%; the shaded region highlights samples with fewer than ten contaminant reads.

**Figure 3.**
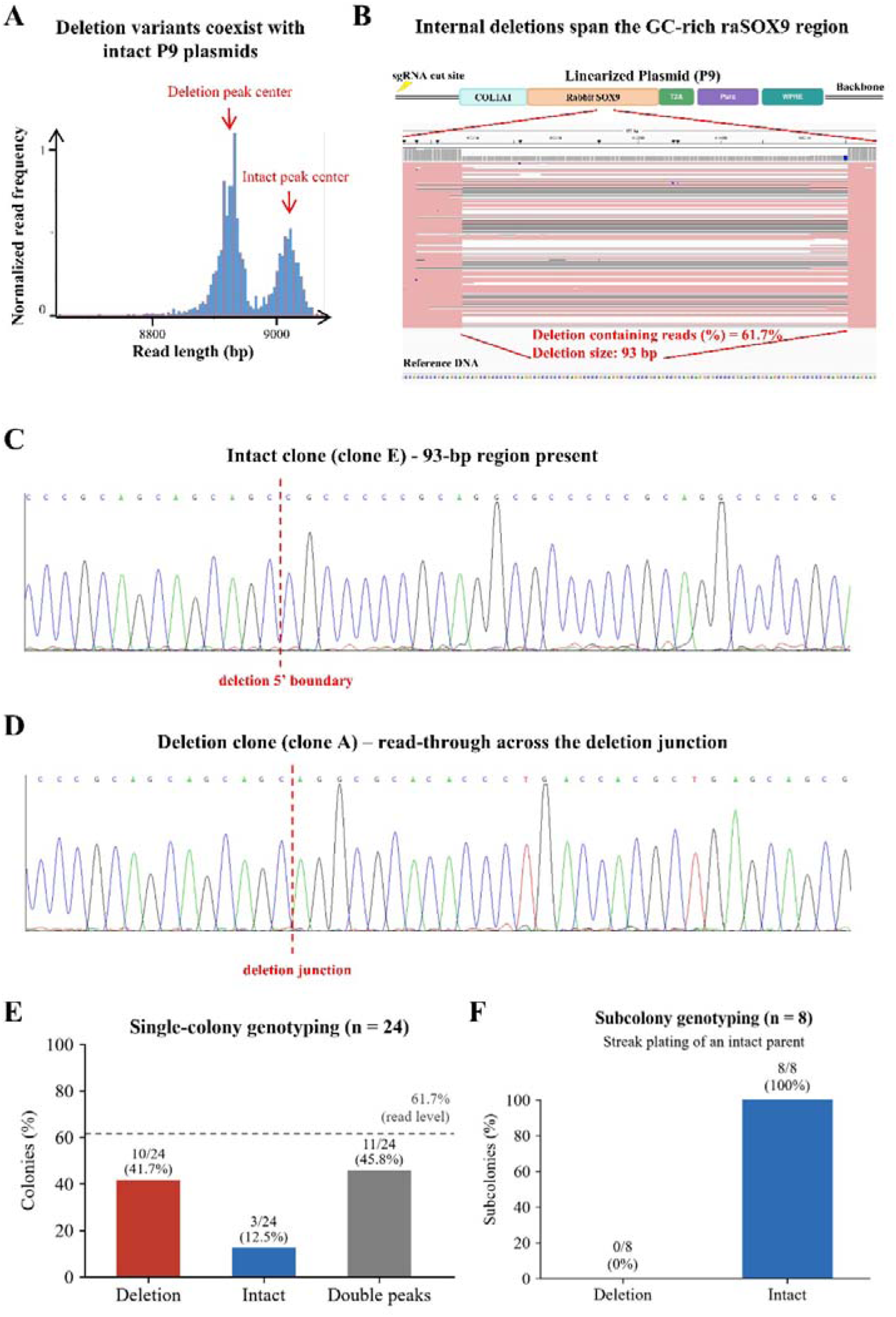
Read-level analysis and clone genotyping reveal a deletion variant in a P9 preparation that passed consensus quality control. (A) Read-length distribution from the P9 preparation, showing two major populations associated with deletion-containing and intact plasmids. Arrows indicate the centres of the two peaks. Read frequencies are normalized. (B) Localization of the deletion within P9. The upper schematic shows the linearized plasmid, including the COL1A1-rabbit SOX9-T2A-Puro-WPRE cassette and the sgRNA-directed cleavage site. Reference-aligned reads reveal a recurrent 93-bp deletion within the GC-rich rabbit SOX9 region. Deletion-containing reads accounted for 61.7% of the analysed P9 reads. (C) Representative Sanger chromatogram from an intact clone (clone E), showing retained sequence at the locus affected by the 93-bp deletion. The vertical dashed line marks the position corresponding to the 5′ boundary of the deletion. (D) Representative Sanger chromatogram from a deletion clone (clone A), showing read-through across the junction formed by joining the sequences flanking the 93-bp deletion. The vertical dashed line marks the deletion junction. (E) Genotyping of 24 single colonies from the glycerol stock that produced the sequenced preparation (Table S6). Bars show the proportions classified as deletion-type (10/24, 41.7%), intact-type (3/24, 12.5%) or double-peak profiles (11/24, 45.8%), with Wilson 95% confidence intervals. A double-peak chromatogram indicates that both sequence forms were present in material from that colony; scored as carrying detectable deletion sequence, 21 of 24 colonies (87.5%) were positive. The horizontal dashed line marks the deletion-containing read fraction of 61.7% for reference; read-level and colony-level proportions describe different sampled populations. (F) Genotyping of eight subcolonies obtained by streak plating an intact parent. All eight yielded only the 548-bp intact amplicon (8/8, 100%); none yielded the 455-bp deletion amplicon (0/8, 0%). Counts and percentages are indicated above the bars. Band intensity was not used to estimate allele proportions.

To encode distinct sample identities, sgRNAs targeted different backbone positions, giving each assigned plasmid a distinct pair of terminal sequence contexts. After the standard 4-h Cas9 reaction, each of the 24 sgRNAs produced a predominant band consistent with plasmid linearization (Fig. 1C and Fig. S1A). Cas9 treatment also yielded discrete bands consistent with linearized products across five plasmids spanning 2.60–36.40 kb (Fig. 1D, Table S1). These results support the selected guide-RNP combinations as 24 cleavage-validated candidate addresses (Table S2). Gel validation established cleavage activity; discrimination between pooled samples was evaluated separately by terminal-signature demultiplexing.

### Terminal-signature-assigned read bins show the highest agreement with their cognate references

We first asked what happens to every read in a pooled run rather than only to the reads that were successfully assigned. Across three sequencing pools comprising 1,224,980 passing reads (Table S4), 303,661 reads (24.79%) carried concordant dual-end signatures and were assigned to a source plasmid. A further 679,114 reads (55.44%) matched only one terminus or carried conflicting dual-end signatures, and 242,205 reads (19.77%) matched no signature (Fig. 1E, Table S4). Assignment yield differed between runs, from 13.76% to 41.42% of passing reads.

Although only about one quarter of passing reads were assigned, the resulting depth was far above the requirement for downstream analysis. The three runs carried 9, 9 and 8 cleavage addresses respectively, giving means of approximately 11,400, 7,300 and 17,000 assigned reads per address, whereas consensus accuracy plateaued at approximately 10 assigned reads (Fig. S3B). Assignment yield, rather than depth per address, therefore sets the practical multiplexing ceiling. Complete assignment statistics are reported here for pools of eight and nine cleavage addresses; the 24 guides are accordingly described as cleavage-validated candidate addresses rather than as a demonstrated 24-plex capacity.

To evaluate the sequence composition of assigned read bins, we cross-mapped reads from five representative bins against the corresponding plasmid references. Agreement was summarized using a fit index calculated as fractional alignment identity multiplied by fractional reference coverage. Each bin showed its highest fit index against its cognate reference, with values ranging from 92.46% to 97.75% (Fig. 1F). Shared backbone sequences produced nonzero fit indices against noncognate references, but cognate-reference values exceeded the strongest competing match by 31.91–82.22 percentage points. This separation was consistent with the assignment of each read bin to its expected plasmid. The fit index measures sequence agreement and does not directly quantify read-assignment accuracy or the fraction of passing reads successfully assigned.

Dual-end read integrity declined with increasing plasmid length across 37 plasmids spanning 2.28–24.30 kb (linear regression, R² = 0.858; Fig. S8A). Approximately 70–80% of reads retained both reference ends for the shortest plasmids, compared with fewer than 20% for plasmids approaching 25 kb. Representative alignments showed greater end-specific truncation in longer plasmids, including a 36.40-kb plasmid that was not included in the regression (Fig. S8B). These results identify read integrity as a constraint on terminal-signature assignment for longer constructs. Accordingly, the 36.40-kb data support structural inspection of recovered reads rather than complete characterization of the plasmid population.

### Sub-percent contaminant detection in computational read mixtures

To select a basecalling model, the same raw nanopore datasets were processed using FAST, HAC, and SUP with identical filtering criteria. SUP produced the highest consensus accuracy and was used for subsequent analyses (Fig. S3A). Random subsampling indicated that consensus accuracy plateaued at approximately 99.95% with about ten assigned reads (Fig. S3B). Relative to individual raw reads, the Racon x3-Medaka workflow reduced the residual error rate by approximately 20-fold (Fig. S3C).

We then evaluated contaminant detection using computational read mixtures, created by combining reads from independently sequenced plasmids at known proportions. For example, a 1% mixture contained 99% reads from the declared plasmid and 1% from a second plasmid designated as the contaminant. No plasmid DNA was mixed in the laboratory for these benchmarks: they test classification after sequencing, and the two designs used in this study are compared side by side in Fig. 2A. Classification used the same class-specific classifier that was applied to the physical mixtures, so that the two designs are reported on a common scale.

Each abundance level was represented by 20 independent subsamples of 5,000 passing reads. Under the prespecified requirement of at least three contaminant reads, the contaminant was detected in all 20 subsamples at 0.2%, 0.5%, 1% and 5% abundance, and in 13 of 20 subsamples at 0.1% (Fig. 2B). Precision was 1.000 in every one of the 100 subsamples: no read was called as contaminant in error anywhere in the benchmark, among 493,200 declared reads. The number of contaminant reads recovered scaled with abundance as expected, with geometric means of 3.0, 6.0, 16.0, 33.7 and 160 reads at 0.1%, 0.2%, 0.5%, 1% and 5% respectively, and minima in any single subsample of 1, 3, 11, 26 and 144 reads. At 0.1% and 0.2%, detection depended on very small numbers of reads and was therefore limited by counting variation. At 0.5%, every subsample yielded at least 11 contaminant reads. Recall averaged 0.62 to 0.68 across levels and was set by the ambiguous-read convention rather than by abundance: a mean of 33% of reads in each subsample carried insufficient class-specific evidence and were labelled ambiguous, bounding recall at approximately 0.67 (Table S7). The corresponding ambiguous fraction in the physical mixtures was 20%; because the same classifier was used for both designs, this difference reflects the different read populations rather than a difference in classification. For near-identical plasmids, classification used known local sequence differences with a base-quality filter and concordant terminal signatures, resolving contaminants that read-length distributions alone could not distinguish (Figs. S3D and S4).

### Physical DNA mixtures detect but do not quantify sub-percent contamination

To test contaminant detection through cleavage, library preparation and sequencing, we mixed purified P6 DNA into P7 DNA at defined molar fractions before Cas9 treatment (Fig. 2A). Within each mixture, both plasmids shared the same cleavage position and terminal signatures, so sequence content rather than read ends distinguished the contaminant. Mixtures were assigned distinct cleavage addresses and pooled, with classification thresholds fixed before analysis.

P6-derived reads were detected in all six samples prepared at 0.5 mol% input and in all six samples prepared at 1 mol% input (Fig. 2C). The observed read fractions did not resolve these two input levels, and their variation was not random. Samples from Pool 1 and Pool 2 separated completely: every Pool 1 sample at 0.5 or 1 mol% gave a lower observed fraction than every Pool 2 sample at either level (geometric means 0.329% and 1.323%, a ratio of 4.0; Mann-Whitney P = 0.0022, the smallest value attainable with six samples per group). Within a pool the scatter was much smaller and, in three of the four pool-by-level groups, was statistically consistent with Poisson variation in the number of contaminant reads recovered. The apparent 7.3-fold and 8.7-fold ranges at 0.5 and 1 mol% are therefore dominated by a single systematic offset between the two pools rather than by tube-to-tube irreproducibility. Because each pool used its own upstream P6 working solution while tubes within a pool shared one, the most parsimonious explanation is a difference between those two working solutions; this could not be resolved retrospectively. At higher inputs the observed fractions tracked the input closely and varied little between samples, with means of 8.082% at 5 mol% and 69.186% at 50 mol%, compared with 0.834% at 0.5 mol% and 0.899% at 1 mol%.

The 50:50 mixture provides the cleanest available estimate of relative recovery, because equimolar mixing requires a single pipetting step from each stock rather than dilution of one component to sub-percent level. Across its three samples the ratio of observed to input odds was 2.02 to 2.56 (mean 2.26), indicating that the shorter plasmid (P6, 5.74 kb) was recovered approximately two-fold more efficiently than P7 (7.02 kb); the three samples at 5 mol% gave a consistent ratio of 1.42 to 1.89 (mean 1.67). The systematic recovery bias is therefore of the order of two-fold and approximately constant across the range in which it can be measured precisely. It is substantially smaller than the 4.0-fold offset between pools at sub-percent input, so recovery during the workflow does not account for that offset; neither does classifier background, which was lower still by a further 50- to 100-fold (Fig. 2D).

Taken together, the workflow detected sub-percent contamination in every sample tested at 0.5 mol% input but did not measure its abundance. Observed read fractions are therefore reported as detected read representation and were not used to estimate input molar fractions; below approximately 5 mol% input the assay is a detector rather than a quantifier.

Single-plasmid controls showed low background misassignment (Fig. 2D). No P6 reads were assigned among 17,598 classified reads in the P7-only control, corresponding to a one-sided 95% upper bound of 0.017%. In the P6-only control, two of 43,022 classified reads were assigned to P7, corresponding to 0.0046% (Wilson 95% CI, 0.0013-0.0168%). Because each control was pooled, prepared and sequenced within the same library as the mixtures it controls for, these values include any contribution from inter-sample cross-talk arising during barcode-free pooled library preparation and sequencing. They describe the observed background under the conditions tested and are not an estimate of a platform-independent false-positive rate.

To test whether classification depended on read length rather than on sequence content, we recalculated P6 fractions using the first 2,500 bp and last 2,500 bp of each qualifying read instead of the full-length read. Across 18 mixture samples, estimates changed by at most 0.478 percentage points (Fig. 2E). Inspection of the 15 cleavage sites in samples V01-V15 confirmed that the equal-length windows retained more than 1 kb of P6- or P7-specific sequence in every case. Classification was therefore not driven by read length alone, although this analysis does not exclude length-dependent recovery bias arising during cleavage, purification, library preparation or sequencing.

We next asked whether the small number of contaminant reads recovered at sub-percent input retained sufficient sequence information to reconstruct the contaminant. P6-called reads were extracted from 12 samples at 0.5 and 1 mol% input (3 to 328 reads per sample) and polished with Racon x3 followed by Medaka. Consensus identities to the 5,754-bp P6 reference ranged from 99.6175% to 99.7043%, with a mean of 99.6704% (Fig. 2F,G). Reference coverage exceeded 99.4% across the displayed samples and varied more than consensus identity (Fig. 2F); in 11 of 12 samples the uncovered intervals were 4-32 bp at the read termini and were located at or immediately adjacent to the Cas9 cleavage site, while one sample reached 100% reference coverage. One sample (Pool1_V08) was reconstructed from only three contaminant reads and is identified separately in Fig. 2F,G; it is reported as a low-depth exploratory result and is not taken as evidence for consensus reliability at comparable depth. These results support both contaminant detection and near-complete sequence reconstruction of the contaminant at 0.5 mol% input under the tested conditions.

### A predominant deletion variant is detected in a P9 preparation that passed consensus QC

To assess whether consensus QC excluded internal structural heterogeneity, we examined individual reads from the P9 preparation that had passed this QC. Read-length profiling revealed two major populations (Fig. 3A). Reference alignment identified a recurrent 93-bp deletion within the GC-rich rabbit SOX9 region, present in 61.7% of the analysed reads (Fig. 3B). Thus, a preparation that passed consensus QC contained a predominant deletion-bearing read population alongside intact plasmids.

Sanger sequencing of individual clones independently confirmed both sequence structures. An intact clone retained the sequence at the deletion boundary, whereas a deletion clone showed direct joining of the flanking sequences across the deletion junction (Fig. 3C,D).

Genotyping 24 single colonies from the same glycerol stock identified 10 deletion-type colonies (41.7%; Wilson 95% CI, 24.5-61.2%), three intact-type colonies (12.5%; 95% CI, 4.3-31.0%) and 11 colonies with double-peak chromatograms at the diagnostic locus (45.8%; 95% CI, 27.9-64.9%; Fig. 3E, Table S6). A double-peak chromatogram from a single colony indicates that both the intact and the deletion sequence were present in the material derived from that colony, that is, that the plasmid population within a single clone was itself heterogeneous. Scored in this way, 21 of 24 colonies (87.5%; 95% CI, 69.0-95.7%) carried detectable deletion sequence and only three yielded exclusively intact sequence. This pattern is consistent with the read-level estimate of 61.7% under a model in which most colonies contain a mixture of both forms, and indicates that the deletion is not clonally fixed but segregates within individual clones. Read-level and colony-level proportions nonetheless describe different sampled populations: the 61.7% read-level estimate describes sequenced molecules from the preparation, whereas colony genotyping describes sequence recovered after colony growth.

To examine genotype retention upon further colony isolation, we streak-plated an intact parent and genotyped eight resulting subcolonies by colony PCR. All eight yielded only the 548-bp intact amplicon, with no 455-bp deletion amplicon detected (Fig. 3F). Band intensity was not used to estimate allele proportions. This result shows retention of the intact genotype in the sampled descendants of that parent under the conditions tested, without establishing a general rate of deletion formation. Together, the read-level analysis and clone sequencing revealed structural heterogeneity that consensus QC had not excluded.

### Reference-spanning coverage across viral vector plasmids

NPlasmid-seq produced continuous reference coverage for representative AAV, lentiviral, and adenoviral vector plasmids ranging from approximately 5.74 to 37.62 kb (Fig. S5A). Representative reads also traversed a 160-bp AAV inverted terminal repeat and a 230-bp lentiviral long terminal repeat (Fig. S5B-C). These data support structural inspection across diverse vector architectures while recognizing the reduced dual-end recovery of the largest plasmids.

### Long poly(A) tracts are associated with increased multimeric-read fractions

Poly(A)-containing IVT-template plasmids generated reads spanning two or more tandem reference-length units. To retain linked plasmid copies for detection, we shortened Cas9 incubation from 4 h to 1 h. This partial-cleavage workflow generated sequenceable ends while limiting cleavage between adjacent plasmid units.

Gel electrophoresis provided supporting evidence for higher-order plasmid species (Fig. 4B). After cleavage, the poly(A)-containing preparation showed a band near the expected dimer size of 4.8 kb alongside the approximately 2.4-kb monomer band. The corresponding higher-molecular-weight band was not apparent in the control lacking poly(A). This pattern was consistent with a dimer-sized species, although migration alone did not establish its sequence or topology.

**Figure 4.**
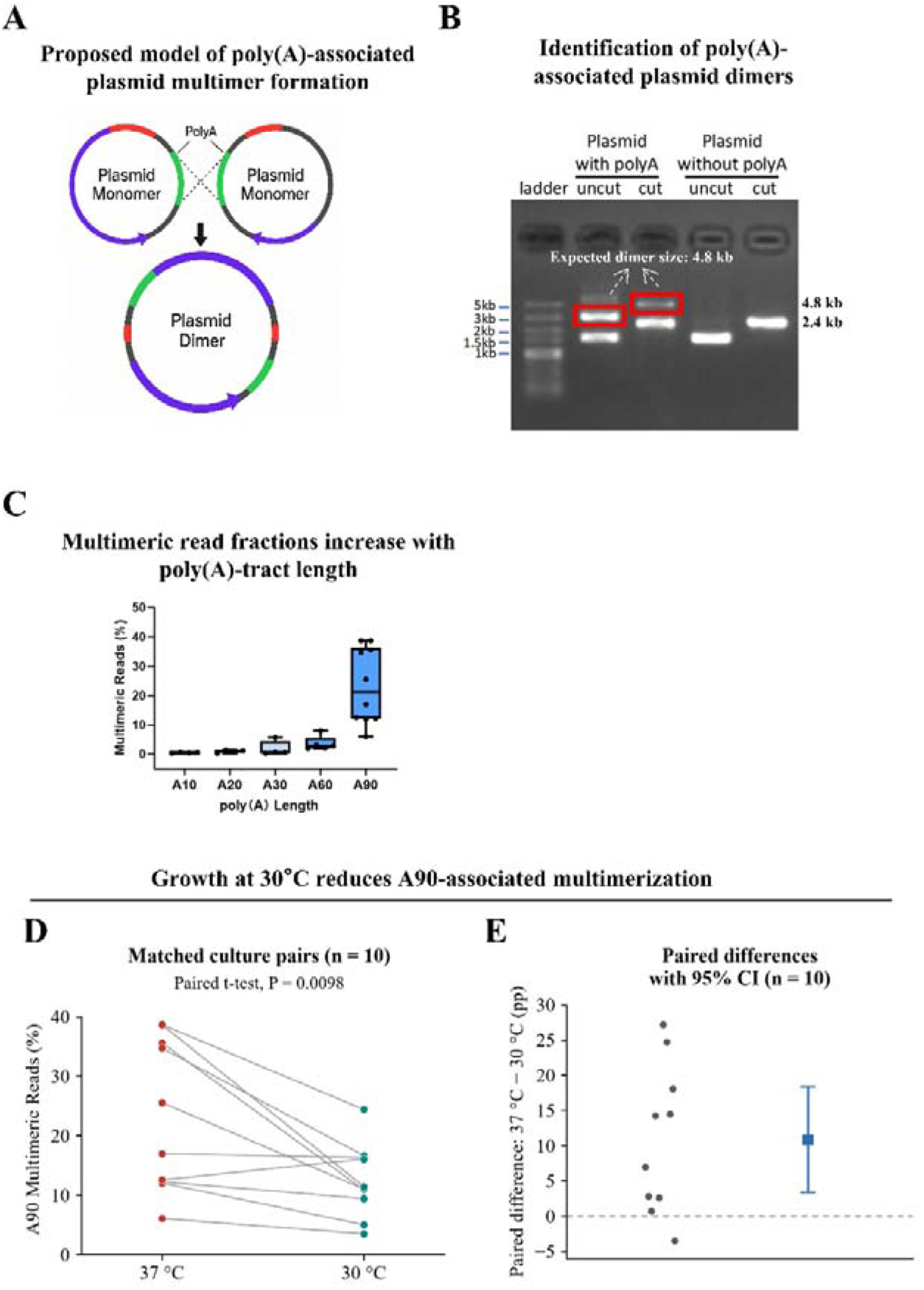
Longer poly(A) tracts are associated with higher multimeric-read fractions, which are reduced by propagation at 30 °C. (A) Hypothetical model of poly(A)-associated plasmid multimer formation. Interactions between homologous sequences may facilitate the formation of linked plasmid copies. Corresponding coloured segments denote homologous regions, and green segments indicate the poly(A) tracts. The proposed mechanism has not been experimentally established. (B) Gel electrophoresis of plasmid preparations with or without a poly(A) tract, before and after Cas9 cleavage. Lanes show the DNA ladder, uncut and cut poly(A)-containing plasmids, and uncut and cut plasmids lacking poly(A). The cut poly(A)-containing preparation shows a band near the expected dimer size of 4.8 kb alongside the approximately 2.4-kb monomer band. Red boxes highlight the candidate higher-order species. Migration of uncut plasmids depends on topology and was not used to infer molecular length; gel migration alone does not establish dimer identity. (C) Multimeric-read fractions in plasmid preparations carrying A10, A20, A30, A60 or A90 tracts after propagation at 37 °C. Each point represents an independent plasmid preparation (A10, A20 and A30, n = 4 each; A60, n = 5; A90, n = 10). Boxes indicate the interquartile range, centre lines indicate medians, and whiskers indicate the data range. (D) Multimeric-read fractions in ten matched pairs of A90 cultures propagated at 37 °C or 30 °C. Each point represents one culture, and connecting lines identify matched cultures derived from the same seed culture. The 37 °C measurements are the same A90 preparations shown in panel C and are not independent additional observations. Paired t-test, P = 0.0098; a Wilcoxon signed-rank test is reported in Methods as a sensitivity analysis. (E) Within-pair differences in multimeric-read fraction for the cultures shown in panel D, calculated as the value at 37 °C minus that at 30 °C. Grey points indicate individual paired differences; the square and error bar indicate the mean difference and its 95% confidence interval. Positive values indicate lower multimeric-read fractions at 30 °C. The dashed horizontal line marks zero difference; pp, percentage points.

We next compared preparations carrying A10, A20, A30, A60 or A90 tracts after propagation at 37 °C using the same partial-cleavage workflow. Multimeric-read fractions were near zero for A10 and A20, remained relatively low for A30 and A60, and were highest for A90 (Fig. 4C). A90 preparations showed substantial variation, with a median of approximately 20% and individual values ranging from approximately 6% to 39%. These results associated longer poly(A) tracts with increased multimeric-read fractions under the tested conditions. The proposed poly(A)-mediated multimer-formation mechanism remains hypothetical (Fig. 4A).

### Propagation at 30 °C reduces the A90 multimeric-read fraction

To test whether propagation temperature modified this phenotype, we compared ten matched pairs of A90 cultures derived from the same seed cultures. Within each pair, cultures were grown at 37 °C or 30 °C under otherwise matched conditions and analysed using the same partial-cleavage and read-classification workflow. The 37 °C measurements are the same A90 data shown in Fig. 4C.

Growth at 30 °C reduced the multimeric-read fraction relative to 37 °C (paired t-test, P = 0.0098; Fig. 4D). The mean paired reduction was approximately 11 percentage points, with a 95% confidence interval of approximately 3-18 percentage points (Fig. 4E). Multimeric reads remained detectable at 30 °C, indicating reduction rather than elimination of the phenotype.

Consistent with this reduction, read-length distributions shifted towards the one-copy population, with fewer reads near twice the plasmid length (Fig. S6B). Raw-signal traces showed one poly(A)-associated signal shift in monomeric reads and two spatially separated shifts in dimeric reads, providing signal-level support for the read-length classification (Fig. S6A). Sanger analysis detected no obvious shortening of the A90 tract under either condition (Fig. S6C). Lower-temperature propagation therefore reduced the A90-associated multimeric-read fraction in the tested strain and construct, as summarized schematically in Fig. S7. These fractions describe sequenced reads rather than molecular abundances, because DNA recovery and sequencing efficiency may depend on molecule length.

## Discussion

### Principal findings

This study distinguishes two questions that are often conflated in plasmid quality control: whether the intended sequence is present and what fraction of individual molecules retains the intended architecture. NPlasmid-seq addresses the second question by combining programmable Cas9 linearization with terminal-signature demultiplexing and read-level structural analysis. In the tested preparations, this approach detected a majority deletion-bearing P9 subpopulation and substantial A90 multimeric-read fractions that were not represented by a simple consensus verdict.

Each of these read-level observations is anchored to an independent laboratory measurement: contaminant classification to physical DNA mixtures prepared before cleavage, the P9 deletion to clone-level genotyping with confirmed chromatograms, and the multimeric species to gel isolation. Where the two kinds of evidence measure different populations, we report them side by side rather than merging them. Reporting detection rate and precision as the primary endpoints, rather than per-read F1, was necessary for this pairing: per-read ground truth exists only in the computational design, whereas detection and precision are measurable in both, and a quality-control assay is judged primarily by whether it detects real contamination without raising false alarms.

The physical mixtures also define the assay’s quantitative limits explicitly. Detection at 0.5 mol% input was reproducible, but the observed read fraction varied several-fold between independently prepared mixtures at that level, while the systematic recovery bias measured at equimolar input was only about two-fold. We therefore present the workflow as a detector of sub-percent contamination rather than as a method for measuring how much contaminant is present. For the intended quality-control application this is the operative question: a preparation either carries detectable foreign sequence or it does not.

### Relationship to existing approaches

Several nanopore-based workflows already support pooled plasmid validation. OnRamp established pooled, amplification-free plasmid sequencing and consensus comparison(4). SAVEMONEY uses reference-aware Bayesian classification and prior construction-error probabilities to resolve barcode-free pools(26). Li et al. developed a high-throughput arrayed in vivo barcoding strategy that requires prior barcode engineering and is primarily optimized for sequence verification(27). nCATS established Cas9-guided adapter ligation for targeted genomic nanopore sequencing(13). Other studies have characterized plasmid multimers across backbones and host strains(11), automated biofoundry-scale validation(28) or used R2C2/Chopper for highly accurate complete-plasmid reconstruction(29).

NPlasmid-seq differs by encoding sample identity in the programmed cleavage coordinate and by treating the read population, rather than only the polished consensus, as a primary analytical output. This preserves native linearized molecules without rolling-circle amplification and enables explicit reporting of dual-end integrity, read-length subpopulations, deletion-bearing reads, and multimeric-read fractions. The 24 guides presented here should be regarded as cleavage-validated candidate addresses; the maximum validated multiplexing capacity depends on complete demultiplexing performance across the selected backbone and guide set, and in the pools tested here the limiting factor was assignment yield rather than depth per plasmid.

### Practical implications

For viral-vector workflows, structural heterogeneity in transfer plasmids may propagate into or affect vector production, particularly for repeat-rich constructs. Molecule-level inspection before large-scale transfection could therefore reduce the risk of committing expensive production resources to a heterogeneous plasmid preparation, although the present study did not directly measure final vector-product consequences.

The P9 result also bears on how clonal isolation is used in practice. Because most colonies derived from the same stock contained both the intact and the deletion sequence, picking and verifying a single colony by Sanger sequencing of one locus may not establish that a preparation is homogeneous. A single double-peak chromatogram, which is often treated as a failed read, is itself informative about within-clone heterogeneity.

For mRNA manufacturing, the data suggest that verifying poly(A)-tract length alone may miss a distinct plasmid-level multimerization phenotype. Multimers can alter the effective molar concentration and copy-number behavior of transcription units(9, 10). Maintaining long encoded poly(A) tracts in plasmid form is a recognized difficulty, and dedicated plasmid designs have been developed to propagate them(23, 24). Under the tested conditions, propagation at 30 °C reduced the A90 multimeric-read fraction without detectable tract shortening, providing a process-level mitigation that requires evaluation across additional strains, origins of replication, copy numbers, and poly(A) designs. A complementary, sequence-level strategy is to interrupt the homopolymer itself: segmented poly(A) tails substantially reduce plasmid recombination without compromising mRNA translation efficiency or half-life(25). The multimeric-read fraction measured here provides a direct read-out with which such design choices can be compared, and the two strategies are not mutually exclusive.

### Limitations

Several limitations define the scope of the conclusions. First, dual-end read recovery decreased with plasmid length; population-level estimates are therefore less complete for constructs above approximately 20 kb. Second, nanopore accuracy is lowest in homopolymeric and GC-rich repetitive regions(30), making orthogonal validation especially important for the P9 and poly(A) findings. Third, the demonstrated multiplexing scale was 12-24 constructs and required a shared backbone containing suitable target sites; complete assignment statistics are reported for a 16-plex pool.

Fourth, the physical contamination mixtures validate the workflow from mixing onward. They were prepared from purified DNA and do not reproduce contamination that arises during co-culture, extraction or library preparation, and they were tested for a single plasmid pair; the near-identical plasmid case remains supported by computational benchmarks only. Background misassignment was measured in single-plasmid controls pooled and sequenced within the same libraries as the mixtures they control for, so the reported background includes any contribution from inter-sample cross-talk under those conditions.

Fifth, the operational detection threshold is a function of read depth, classifier operating point and benchmark design rather than an intrinsic sensitivity; at 0.1% abundance and 5,000 reads per plasmid, roughly five contaminant reads are expected, and performance at that level reflects counting statistics. Recall in the computational benchmark is further limited by the deliberately conservative ambiguous-read convention, which favours precision over recall for quality-control use; a less conservative rule would raise recall at the cost of false alarms.

Sixth, observed read fractions in physical mixtures exceeded nominal molar inputs and, at sub-percent inputs, differed systematically by about four-fold between the two preparation pools. The equimolar mixture indicates a relative recovery bias of approximately two-fold favouring the shorter plasmid, which is smaller than the between-pool offset; because each pool used its own upstream P6 working solution while tubes within a pool shared one, the offset is most plausibly attributable to the preparation of those working solutions rather than to the assay, but this could not be established retrospectively and we cannot exclude that some step of the pooled workflow differed between pools. Observed read fractions are therefore reported as detected read representation rather than molar abundance, and the assay is not presented as quantitative below approximately 5 mol% input. Preparing both pools from a single, independently quantified working solution, and spiking each plasmid into the other at matched molar fractions within one library, would separate preparation error from length-dependent recovery.

Seventh, complete cleavage separates tandem repeat units, so gel evidence alone cannot exclude multimers, and uncut lanes contain topoisomers whose migration is not proportional to length; our conclusion rests on the purified-band re-cleavage experiment rather than on apparent size. Eighth, the signal-assisted poly(A) estimator used an approximate mapping between basecalled position and raw signal and requires quantitative validation against defined standards. Ninth, colony genotyping classified double-peak chromatograms as evidence of within-clone coexistence of both sequence forms but did not quantify the proportion of each form within a colony. Finally, the poly(A)-associated multimerization mechanism was not established, and the temperature effect was tested in one host-strain context and a limited construct set.

## Conclusion

NPlasmid-seq uses a programmed Cas9 cleavage coordinate as a physical sample address to support pooled, molecule-level nanopore plasmid quality control. In computational mixtures, the workflow detected the contaminant in all 20 subsamples at read fractions down to 0.2% and in 13 of 20 subsamples at 0.1%, with a precision of 1.000 throughout; detection at 0.2% rested on as few as three contaminant reads, so 0.5% is adopted as the operational threshold. In physical mixtures, the contaminant was detected in all six samples prepared at 0.5 mol% input and all six prepared at 1 mol% input, and contaminant consensus sequences reached a mean identity of 99.67%, although one exploratory reconstruction used only three reads. Read-level analysis estimated that 61.7% of reads carried a recurrent internal deletion, which was independently supported by clone-level genotyping; double-peak chromatograms were consistent with coexistence of intact and deletion sequences within individual clones. Long poly(A) tracts were associated with higher multimeric-read fractions, and the multimeric species was confirmed as a tandem plasmid. Propagation at 30 °C reduced the A90 multimeric-read fraction under the tested conditions. These findings support the use of read-level structural metrics alongside consensus sequence validation. Further work is needed to assess performance with long plasmids and homopolymeric regions, to validate contamination detection across additional plasmid pairs, to characterize length-dependent recovery, and to establish whether reliable quantification is possible below approximately 5 mol% input.

## Declarations

### Generative-AI declaration

During preparation of this work the authors used ChatGPT (OpenAI) to improve language clarity and readability (grammar, phrasing, conciseness). After using this tool the authors reviewed and edited the content as needed and take full responsibility for the content of the published article.

### Author contributions

X.B.Z. and J.P.Z. conceived and designed the study, obtained funding, and supervised the work. Y.X. was the primary experimental contributor and drafted the manuscript. X.Y.M. and D.S. performed the main data analysis and code development. W.W. contributed to data curation, organization and interpretation of the additional validation results, and writing – review and editing. S.M.Z. and J.Q.C. assisted with figure preparation; Y.X.Y., D.S. and J.X.R. assisted with experiments. S.C.P. and S.D.W. provided administrative support and critical revisions. X.B.Z. integrated feedback and approved the final version. All authors read and approved the manuscript prior to submission.

