## Supplementary material for "NPlasmid-seq: Cas9-encoded terminal signatures enable molecule-level nanopore quality control of vector and IVT-template plasmids": Supplementary File 1.pdf

P1:

>S1488 pU6-sgKLRC1a-gN21(Sa-v2)

AGCTGGCACGACAGGTTTCCCGACTGGGCACGACAGGTTTCCCGACTGGAAAGCGGGCAGTGAG
CGCAACGCAATTAAGGTGAGTTAGCTCACTCATTAGGCACCCCAGGCTTTACACTTTATGCTTCCGG
CTCGTATGTTGTGTGGAATTGTGAGCGGATAACAATTTCACACAGGACCCCAGTGGAAAGACGCGC
AGGCAAAACGCACCACGTGACGGAGCGTGACCGCGCGCCGAGCGCGCGCCAAGGTCGGGCAGG
AAGAGGGCCTATTTCCCATGATTCCTTCATATTTGCATATACGATACAAGGCTGTTAGAGAGATAATTA
GAATTAATTTGACTGTAAACACAAAGATATTAGTACAAAATACGTGACGTAGAAAGTAATAATTTCTTGG
GTAGTTTGCAGTTTTAAAATTATGTTTTAAAATGGACTATCATATGCTTACCGTAACTTGAAAGTATTTC
GATTTCTTGGGTTTATATATCTTGTGGAAAGGACGAAACACCGCACTGCAGAGATGGATAACCAGTTT
AAGTACTCTGTGCTGGAAACAGCACAGAATCTACTTAAACAAGGCAAAATGCCGTGTTTATCTCGTC
AACTTGTTGGCGAGATTTTTTTCTCGAGCAGGTGGCACTTTTCGGGGAAATGTGCGCGGAACCCCT
ATTTGTTTATTTTTCTAAATACATTCAAATATGTATCCGCTCATGAGACAATAACCCTGATAAATGCTTCA
ATAATATTGAAAAAGGAAGAGTATGAGTATTCAACATTTCCGTGTCGCCCTTATTCCCTTTTTTGCGGC
ATTTTGCCTTCCTGTTTTTGCTCACCCAGAAACGCTGGTGAAAGTAAAAGATGCTGAAGATCAGTTG
GGTGCACGAGTGGGTTACATCGAACTGGATCTCAACAGCGGTAAGATCCTTGAGAGTTTTCGCCCC
GAAGAACGTTTTCCAATGATGAGCACTTTTAAAGTTCTGCTATGTGGCGCGGTATTATCCCGTATTGA
CGCCGGGCAAGAGCAACTCGGTCGCCGCATACACTATTCTCAGAATGACTTGGTTGAGTACTCACC
AGTCACAGAAAAGCATCTTACGGATGGCATGACAGTAAGAGAATTATGCAGTGCTGCCATAACCATG
AGTGATAACACTGCGGCCAACTTACTTCTGACAACGATCGGAGGACCGAAGGAGCTAACCGCTTTT
TTGCACAACATGGGGGATCATGTAACTCGCCTTGATCGTTGGGAACCGGAGCTGAATGAAGCCATA
CCAAACGACGAGCGTGACACCACGATGCCTGTAGCAATGGCAACAACGTTGCGCAAACTATTAACT
GGCGAACTACTTACTCTAGCTTCCCGGCAACAATTAATAGACTGGATGGAGGCGGATAAAGTTGCAG
GACCACTTCTGCGCTCGGCCCTTCCGGCTGGCTGGTTTATTGCTGATAAATCTGGAGCCGGTGAGC
GTGGGTCTCGCGGTATCATTGCAGCACTGGGGCCAGATGGTAAGCCCTCCCGTATCGTAGTTATCTA
CACGACGGGGAGTCAGGCAACTATGGATGAACGAAATAGACAGATCGCTGAGATAGGTGCCTCACT
GATTAAGCATTGGTAACTGTCAGACCAAGTTTACTCATATATACTTTAGATTGATTTAAAACTTCATTTTT
AATTTAAAAGGATCTAGGTGAAGATCCTTTTTGATAATCTCATGACCAAAATCCCTTAACGTGAGTTTT
CGTTCCACTGAGCGTCAGACCCCGTAGAAAAGATCAAAGGATCTTCTTGAGATCCTTTTTTTCTGCG
CGTAATCTGCTGCTTGCAAACAAAAAAACCACCGCTACCAGCGGTGGTTTGTTTGCCGGATCAAGA
GCTACCAACTCTTTTTCCGAAGGTAACTGGCTTCAGCAGAGCGCAGATACCAAATACTGTTCTTCTA
GTGTAGCCGTAGTTAGGCCACCACTTCAAGAACTCTGTAGCACCGCCTACATACCTCGCTCTGCTAA
TCCTGTTACCAGTGGCTGCTGCCAGTGGCGATAAGTCGTGTCTTACCGGGTTGGACTCAAGACGAT
AGTTACCGGATAAGGCGCAGCGGTCGGGCTGAACGGGGGGTTCGTGCACACAGCCCAGCTTGGA
GCGAACGACCTACACCGAACTGAGATACCTACAGCGTGAGCTATGAGAAAGCGCCACGCTTCCCGA
AGGGAGAAAGGCGGACAGGTATCCGGTAAGCGGCAGGGTCGGAACAGGAGAGCGCACGAGGGA
GCTTCCAGGGGGAAACGCCTGGTATCTTTATAGTCCTGTCGGGTTTCGCCACCTCTGACTTGAGCG
TCGATTTTTGTGATGCTCGTCAGGGGGGCGGAGCCTATGGAAAAACGCCAGCAACGCGGCCTTTTT
ACGGTTCCTGGCCTTTTGCTGGCCTTTTGCTCACATGTTCTTTCCTGCGTTATCCCCTGATTCTGTG
GATAACCGTATTACCGCCTTTGAGTGAGCTGATACCGCTCGCCGCAGCCGAACGACCGAGCGCAGC
GAGTCAGTGAGCGAGGAAGCGGAAGAGCGCCCAATACGCAAACCGCCTCTCCCCGCGCGTTGGC
CGATTCATTAATGC

P2:

>P2525 pAAV-Alb2(SA85-0)-P2A-BDDF8-N6-mir122T-15(S1002FF)

TTTCTGGGTGAGCAAAAACAGGAAGGCAAAATGCCGCAAAAAAGGGAATAAGGGCGACACGGAAAT
GTTGAATACTCATACTCTTCCTTTTTCAATATTATTGAAGCATTTATCAGGGTTATTGTCTCATGAGCGG
ATACATATTTGAATGTATTTAGAAAAATAAACAAATAGGGGTTCCGCGCACATTTCCCCGAAAAGTGCC
ACCTGACGTCTAAGAAACCATTATTATCATGACATTAACCTATAAAAATAGGCGTATCACGAGGCCCTT
TCGTCTCGCGCGTTTCGGTGATGACGGTGAAAACCTCTGACACATGCAGCTCCCGGAGACGGTCA
CAGCTTGTCTGTAAGCGGATGCCGGGAGCAGACAAGCCCGTCAGGGCGCGTCAGCGGGTGTTGG
CGGGTGTCGGGGCTGGCTTAACTATGCGGCATCAGAGCAGATTGTACTGAGAGTGCACCATAAAAT
TGTAAACGTTAATATTTTGTTAAAATTCGCGTTAAATTTTTGTTAAATCAGCTCATTTTTTAACCAATAGA
CCGAAATCGGCAAAATCCCTTATAAATCAAAAGAATAGCCCGAGATAGAGTTGAGTGTTGTTCCAGTT
TGGAACAAGAGTCCACTATTAAAGAACGTGGACTCCAACGTCAAAGGGCGAAAAACCGTCTATCAG
GGCGATGGCCCACTACGTGAACCATCACCCAAATCAAGTTTTTTGGGGTCGAGGTGCCGTAAAGCA
CTAAATCGGAACCCTAAAGGGAGCCCCCGATTTAGAGCTTGACGGGGAAAGCCGGCGAACGTGGC
GAGAAAGGAAGGGAAGAAAGCGAAAGGAGCGGGCGCTAAGGCGCTGGCAAGTGTAGCGGTCACG
CTGCGCGTAACCACCACACCCGCCGCGCTTAATGCGCCGCTACAGGGCGCGTACTATGGTTGCTTT
GACGTATGCGGTGTGAAATACCGCACAGATGCGTAAGGAGAAAATACCGCATCAGGCGCCCCTGCA
GGCAGCTGCGCGCTCGCTCGCTCACTGAGGCCGCCCGGGCAAAGCCCGGGCGTCGGGCGACCT
TTGGTCGCCCGGCCTCAGTGAGCGAGCGAGCGCGCAGAGAGGGAGTGGCCAACTCCATCACTAG
GGGTTCCTGCGGCCGCACCCCAATGGCCTAACAACAGTTTACCGAGTGAGAGACACAAAAAATTCC
AACACACTATTGCAATGAAAATAAATTTCCTTTATTAGCCAGAAGTCAGATGCTCAAGGGGCTTCATGA
TGTCCCCATAATTTTTGGCAGAGGGAAAAAGATTGGAGTGTGACAATGGTTTAAACTCAGTAGAGGT
CCTGTGCCTCGCAGCCCAGAACCTCCATCCTCAGGGCAATCTGGTGCACCCAACTCTGGGGGTGA
ATTCGAAGGTAGCGAGTCAGTAACGGTGGGTCTAGAGAGTTCACCACAGGTGTGAAGGAGTCTTGA
TTTCCCTGAAAAACCTTTACTTTGCCATTCTGAAAAAAGAGAGTCCACTGATGGCCATCTTGACTGCT
GGAGATGAGGAACTCCTTCACATACATGCTGGTAAGCAGAGATTTTACTCCCTGAGTAGTTACTCCT
GTGACTTTCATTGTCTTCTGGAAGTCCACTTGCAGCCACTCTTTTGGATTATTCACCTGAGGTCTCCA
GGCATTACTCCTCCCTTGGAGGTGAAGTCGAGCTTTTGAAGGAGACCAGGTGGCAAACATATTGGT
AAAGTAGGATGAAGCAGTAATCTGTGCATCTGATATTGCTTTACTCTCCATTCCCAATGGCATGCTGC
AACTATTTAAATCACAGCCCATCAACTCCATGCGAAGAGTGCTGCGAATGCTATAATGAGTTGGGTGC
AAACGGATGTATCGAGCAATAATTGGAGGGTTAAAAATATTGTGTTTTATCCCAGATGAATCCACATTG
CCAAAGAAGACCATTAAGGTTCCAGTGGAATTTCCTCGATAAGTCTGCCACTTCTTCCCATCAAGACT
ATACATGATGATAAACTGAGAGATGTAGAGGCTGGAGAACTTCTGACGGGCACCCTGGGTCTTGATG
CCGTGAATAATCATTGGTGCCAACAGATCCACCTTGATCCAAGAAAAGGGCTCCTTGGTGCTCCAG
GCATTGATTGATCCGGAATAATGAAGTCTGGCCAGCTTTGGGGCCCACTGTCCATATTGTCCTGAAG
CTGTAATCTGAAAATCTCTAATGTGTCCAGAAGCCATTCCCAGGGGAGTCTGACACTTATTGCTGTAC
ACCAGAAAAAGTGTGCTCATCCCAGCATGTAGATGCTCGCCAATAAGGCATTCCACCCGCCAAATTC
CAGCTTTGGATGGTAACATTTCCACTGTCTCAAAAACACCTGGATAGAGATTGTACAGTGCCATTTTA
TACTCCTCTTTTTTTCGTACAGTGAACACATGTCCACTGAAATGAATAGAATGGATGTTTTCATTGCTG
CCCATGCTGAGCAGATACCATCGAATCCTTTGATCCTGAGCCATTACTAAGCCAGGTAGTGTATCCAT
TATGTAGCCATTGATTGCATGGAAGCGATAATTCTCTTTAAAAGTGGGATCTTCCATCTGGATATTGCA
GGGAGCCCTGCAGTTTCTTTCCATATTTTCAGTGAAGTACCAGCTTTTGGTCTCATCAAAGATGGTG
AAAAACAGAGCAAATTCCTGTACTGTCACTTGTCTCCCATGAGCAGGGTTCAGTGTGTTAGTGTGGC
AGACCAGAAGGGGTCCAATCAGGCCTGAGTGCACATCTTTTTCCAGGTCAACATCAGAGAAATAAG

CCCAGGCTTTGCAGTCAAACTCATCTTTAGTGGGTGCCATATGATGTTGCACTTTCCAAAAGTAAGTT
TTGGTTTCATTAGGCTTGACAAAGTTTTTTCTAGGTTCTGCTCCTTGCCTCTGATCTTCCTCATAAGAA
ATAAGGCTAGAATAGAAGGAATAGGGACGAGAGGCCTGATTTCTGAAAGTTACCATGATATTATCTTC
AACTTCTGCTCTTATATATGGCCCCAGGAGTCCCAAATGTTCATTTAGTTCTCCACGGTATAAGGGCT
GAGTAAAGGAGCCATCAGTAAATTCCTGGAAAACAACTTTCTTGAACTGAGGGACACTGCCACTCTG
AGCCCTGTTTCTTAGAACATGTGGGGAGCTACTCATCCCATAATCCCAGAGCCTCTCCACTGCAGCA
ATAAAATAGTGTCGTGTTTTCTTTTGAAAGCTGCGGGGGCTCTGATTTTCATCCTCATCATAAATGTCA
AAATCTTCCTTCTTCATTTCAACTGATATGGTATCATCATAGTCAATTTCCTCTTGATCTGACTGAAGAG
TAGTACGAGTTATTTCCCGTTGATGGCGTTTCAAGACTGGTGGCGAAACGTTCGAATCATTACTAGTG
TTTGAGTTGTTACTCACGTTCGTCGCGTTTTGAGAGAAGCTTCTTGGTTCAATGGCATTGTTTTTACT
CAGCAAGTATGCTGAAATATCTTCATAACTGTCCTCGTAATAATCACCAGTGTTCTTGTCACAACTAGA
AACCTTCAGTAAGGCGGTCATGCCTCTGTTCCGAAAGTCTGAGTTGTGGCACCCCAGAATCCATAG
ACCTGGGTTTTCCATCGACATGAAGACAGTTTCTCCTGAGAATGGGAATAGGGTGAGTGTGTCTTCA
TAGACCATTTTGTGTTTGAAGGTATATCCAGAGAAGAAGACAGAAAGGAAGTCAGTCTGTGCTCCAA
TGCTTAGAATGTACCAGTATGCCACCTCATGCAAACAAACTGACAACTGCAAACTATCAAAAACATAG
CCATTGATGCTGTGCATGATGTTGGAGGCTTGGAACTCTGGATCCTCAAGCTGCACTCCAGCTGGA
TTGGGGAGAAAGCGTTGTATATTCTCTGTGAGGTACCAGCTTCGGTTCTCATCAAATACAGAAAACA
GGATGACATTCCTCTTGTCTGACATTATCTGGTTTCCTCTTTGATCTACAGATTCTTTGTAGCAGATGA
GGAGAGGGCCAATGAGTCCTGAAGCTAGATCTCTCTCCATATTAACGAAACTAGAGTAATAGCGGGT
CAGGCACCGAGGATCTGATTTAGTTGGCCCATCTTCTACAGTCACTGTCCATTTATATTTGAATATTTC
TCCTGGCAGAATTGGAAAATCCTTCAAATGTTTTACACCTTTTGGTAATCTCCTTGAATACAAAGGAC
GGACATCAGTGATTCCGTGAGGGTAGATGTTATATGGTCTGCTTGCTTGATTCTTAAATATAATCAACA
GTGTGTCTCCAACTTCCCCATAAAGTAAAGGTCCCAAGATTCCTGATTCATGCTGAATAGCTTCACGA
GTCTTAAAGGTTTCATCTGTGTATGCCATAAATCGGACTTTTTTGTACTTCCTACCAATCCGCTGAGG
GCCATTGTTCAAATATTGACTTTTATAACTTCTGTCATCGGGGGCGAGGACTAAGGGAGCATAGTCCC
AGTCCTCCTCTTCAGCAGCAATGTAATGTACCCAAGTTTTAGGATGCTTCTTGGCAACTGAGCGAATT
TGGATAAAGGAAGGAGAGTTGTCATCATCAAACCTGACCACATCCATTTCAGAATCAGTAAGATCATC
ATCATAGTCTTCCGCTTCTTCATTATTTTTCATTCGTAGTTGGGGTTCCTCTGGACAGCTGTCTACTTT
GACATAAGCTTCCATGCCATCATGTTGGTGGGAAGAGATATGACAAAACAGTAGAAACTGTCCAAGG
TCCATCAAGAGTGTTTGAGCAGTAAGGAAAGTTATTGGCGAGATTTCCAAGGACGCCTGGCGATGG
TTCCTCACAAGAAATGTGTGACCTTCGAGGAATATTGAGTGCACTTCAGGAGTGGTGCCCATTCCAA
TCACATGCCAATAGACTGATTTCCTGTGGCATCCAATCAGACCTGGCAGAGACCTGTTTACATAACCA
TTGACTGTGTGCATTTTAGGCCAGGCCCGAGCAGATGCAGCATCCCTATCCTGCATCAAGGAGTTCT
TTGTTTCTGAGTGCCAACTTTTCCCTTCATCAAATACAGCAAAAAGTAGTATAAATTTGTGCAAGGTCT
GTGTCTTTTCCTTGGCCAGACTCCCTTCTCTACATACTAGTAGGGCTCCAATGAGGCCTGAATTCAA
GTCTTTTACCAGGTCCACATGAGAAAGATATGAGTAGGTAAGGCACAGTGGGTCAGAGGCCATTGG
ACCATTCTCTTTCAGGACCTGCCAGACATATGTATGGCTTCCACCAGGGAAGACTTTATCATCTTCTT
TCTCCCTTTGACTGGTCTGATCATCATATTCAGCTCCCTCAGAAGCTTTCCAGTAGGATACACCAACA
GCATGAAGACTGACAGGATGGGAAGCCATGTTCTTAAGTGTAATGACCACTGTATCATAAACCTCAG
CCTGGATGGTAGGACCTAGCAGACCCATCCAGGGTGGCCTTGGCTTAGCGATGTTGAAAAGGTGAT
CCGTGAATTCTACAAACAGAGTCTTTTTGTACACGACTGAGGTGTTGAATGGAAAAGATTTTGGCAC
TCTAGGAGGAAATCTTGCGTCCACAGGCAGCTCACCGAGATCACTTTGCATATAGTCCCATGACAGT
TCCACTGCACCCAGGTAGTATCTTCTGGTGGCACTAAAGCAGAATCGCAAAAGGCACAGAAAGAAG
CAGGTGGAGAGCTCTATTTGCATACGCGTGGGGCCAGGGTTTTCTTCGACGTCGCCGGCCTGTTTC

AGCAGAGAGAAGTTTGTAGCGCCGGAGCCGGCTAAGGCGTCTTTGCATCTAGTGACAAGGTTTGGA
CCCTGAAAACAAAGAAATGATGGATGTTAGAAAAGTATTAGCAGGACCAATGGCCTAACAACAGTTTA
CGATATCGGCCGCAGGAACCCCTAGTGATGGAGTTGGCCACTCCCTCTCTGCGCGCTCGCTCGCT
CACTGAGGCCGGGCGACCAAAGGTCGCCCGACGCCCGGGCGGCCTCAGTGAGCGAGCGAGCGC
GCAGCTGCCTGCAGGACATGTGAGCAAAAGGCCAGCAAAAGGCCAGGAACCGTAAAAAGGCCGC
GTTGCTGGCGTTTTTCCATAGGCTCCGCCCCCCTGACGAGCATCACAAAAATCGACGCTCAAGTCA
GAGGTGGCGAAACCCGACAGGACTATAAAGATACCAGGCGTTTCCCCCTGGAAGCTCCCTCGTGC
GCTCTCCTGTTCCGACCCTGCCGCTTACCGGATACCTGTCCGCCTTTCTCCCTTCGGGAAGCGTGG
CGCTTTCTCATAGCTCACGCTGTAGGTATCTCAGTTCGGTGTAGGTCGTTCGCTCCAAGCTGGGCT
GTGTGCACGAACCCCCCGTTCAGCCCGACCGCTGCGCCTTATCCGGTAACTATCGTCTTGAGTCCA
ACCCGGTAAGACACGACTTATCGCCACTGGCAGCAGCCACTGGTAACAGGATTAGCAGAGCGAGGT
ATGTAGGCGGTGCTACAGAGTTCTTGAAGTGGTGGCCTAACTACGGCTACACTAGAAGAACAGTATT
TGGTATCTGCGCTCTGCTGAAGCCAGTTACCTTCGGAAAAAGAGTTGGTAGCTCTTGATCCGGCAAA
CAAACCACCGCTGGTAGCGGTGGTTTTTTTGTTTGCAAGCAGCAGATTACGCGCAGAAAAAAAGGA
TCTCAAGAAGATCCTTTGATCTTTTCTACGGGGTCTGACGCTCAGTGGAACGAAAACTCACGTTAAG
GGATTTTGGTCATGAGATTATCAAAAAGGATCTTCACCTAGATCCTTTTAAATTAAAAATGAAGTTTTAA
ATCAATCTAAAGTATATATGAGTAAACTTGGTCTGACAGTTACCAATGCTTAATCAGTGAGGCACCTAT
CTCAGCGATCTGTCTATTTCGTTCATCCATAGTTGCCTGACTCCCCGTCGTGTAGATAACTACGATAC
GGGAGGGCTTACCATCTGGCCCCAGTGCTGCAATGATACCGCGGCTTCCACGCTCACCGGCTCCA
GATTTATCAGCAATAAACCAGCCAGCCGGAAGGGCCGAGCGCAGAAGTGGTCCTGCAACTTTATCC
GCCTCCATCCAGTCTATTAATTGTTGCCGGGAAGCTAGAGTAAGTAGTTCGCCAGTTAATAGTTTGCG
CAACGTTGTTGCCATTGCTACAGGCATCGTGGTGTCACGCTCGTCGTTTGGTATGGCTTCATTCAGC
TCCGGTTCCCAACGATCAAGGCGAGTTACATGATCCCCCATGTTGTGCAAAAAAGCGGTTAGCTCCT
TCGGTCCTCCGATCGTTGTCAGAAGTAAGTTGGCCGCAGTGTTATCACTCATGGTTATGGCAGCACT
GCATAATTCTCTTACTGTCATGCCATCCGTAAGATGCTTTTCTGTGACTGGTGAGTACTCAACCAAGT
CATTCTGAGAATAGTGTATGCGGCGACCGAGTTGCTCTTGCCCGGCGTCAATACGGGATAATACCGC
GCCACATAGCAGAACTTTAAAAGTGCTCATCATTGGAAAACGTTCTTCGGGGCGAAAACTCTCAAGG
ATCTTACCGCTGTTGAGATCCAGTTCGATGTAACCCACTCGTGCACCCAACTGATCTTCAGCATCTTT
TACTTTCACCAGCG

P3:

>P2417 pRSC-SFFV-Cas9NG-E2A-Puro-wpre(Sp-NG)-(SpG)

GGGCGGAGCCTATGGAAAAACGCCAGCAACGCGGCCTTTTTACGGTTCCTGGCCTTTTGCTGGCC
TTTTGCTCACATGTTCTTTCCTGCGTTATCCCCTGATTCTGTGGATAACCGTATTACCGCCTTTGAGT
GAGCTGATACCGCTCGCCGCAGCCGAACGACCGAGCGCAGCGAGTCAGTGAGCGAGGAAGCGGA
AGAGCGCCCAATACGCAAACCGCCTCTCCCCGCGCGTTGGCCGATTCATTAATGCAGCTGGCACGA
CAGGTTTCCCGACTGGAAAGCGGGCAGTGAGCGCAACGCAATTAAGGTGAGTTAGCTCACTCATTA
GGCACCCCAGGCTTTACACTTTATGCTTCCGGCTCGTATGTTGTGTGGAATTGTGAGCGGATAACAA
TTTCACACAGGAAACAGCTATGACCATGATTACGCCAAGCGCGCAATTAACCCTCACTAAAGGGAAC
AAAAGCTGGAGCTGCAAGCTTAATGTAGTCTTATGCAATACTCTTGTAGTCTTGCAACATGGTAACGA
TGAGTTAGCAACATGCCTTACAAGGAGAGAAAAAGCACCGTGCATGCCGATTGGTGGAAGTAAGGT
GGTACGATCGTGCCTTATTAGGAAGGCAACAGACGGGTCTGACATGGATTGGACGAACCACTGAAT
TGCCGCATTGCAGAGATATTGTATTTAAGTGCCTAGCTCGATACATAAACGGGTCTCTCTGGTTAGAC
CAGATCTGAGCCTGGGAGCTCTCTGGCTAACTAGGGAACCCACTGCTTAAGCCTCAATAAAGCTTG
CCTTGAGTGCTTCAAGTAGTGTGTGCCCGTCTGTTGTGTGACTCTGGTAACTAGAGATCCCTCAGAC
CCTTTTAGTCAGTGTGGAAAATCTCTAGCAGTGGCGCCCGAACAGGGACTTGAAAGCGAAAGGGAA
ACCAGAGGAGCTCTCTCGACGCAGGACTCGGCTTGCTGAAGCGCGCACGGCAAGAGGCGAGGGG
CGGCGACTGGTGAGTACGCCAAAAATTTTGACTAGCGGAGGCTAGAAGGAGAGAGATGGGTGCGA
GAGCGTCAGTATTAAGCGGGGGAGAATTAGATCGCGATGGGAAAAAATTCGGTTAAGGCCAGGGGG
AAAGAAAAAATATAAATTAAAACATATAGTATGGGCAAGCAGGGAGCTAGAACGATTCGCAGTTAATCC
TGGCCTGTTAGAAACATCAGAAGGCTGTAGACAAATACTGGGACAGCTACAACCATCCCTTCAGACA
GGATCAGAAGAACTTAGATCATTATATAATACAGTAGCAACCCTCTATTGTGTGCATCAAAGGATAGAG
ATAAAAGACACCAAGGAAGCTTTAGACAAGATAGAGGAAGAGCAAAACAAAAGTAAGACCACCGCA
CAGCAAGCGGCCGCTGATCTTCAGACCTGGAGGAGGAGATATGAGGGACAATTGGAGAAGTGAATT
ATATAAATATAAAGTAGTAAAAATTGAACCATTAGGAGTAGCACCCACCAAGGCAAAGAGAAGAGTGG
TGCAGAGAGAAAAAAGAGCAGTGGGAATAGGAGCTTTGTTCCTTGGGTTCTTGGGAGCAGCAGGA
AGCACTATGGGCGCAGCGTCAATGACGCTGACGGTACAGGCCAGACAATTATTGTCTGGTATAGTG
CAGCAGCAGAACAATTTGCTGAGGGCTATTGAGGCGCAACAGCATCTGTTGCAACTCACAGTCTGG
GGCATCAAGCAGCTCCAGGCAAGAATCCTGGCTGTGGAAAGATACCTAAAGGATCAACAGCTCCTG
GGGATTTGGGGTTGCTCTGGAAAACTCATTTGCACCACTGCTGTGCCTTGGAATGCTAGTTGGAGTA
ATAAATCTCTGGAACAGATTTGGAATCACACGACCTGGATGGAGTGGGACAGAGAAATTAACAATTA
CACAAGCTTAATACACTCCTTAATTGAAGAATCGCAAAACCAGCAAGAAAAGAATGAACAAGAATTAT
TGGAATTAGATAAATGGGCAAGTTTGTGGAATTGGTTTAACATAACAAATTGGCTGTGGTATATAAAAT
TATTCATAATGATAGTAGGAGGCTTGGTAGGTTTAAGAATAGTTTTTGCTGTACTTTCTATAGTGAATAG
AGTTAGGCAGGGATATTCACCATTATCGTTTCAGACCCACCTCCCAACCCCGAGGGGACCCGACAG
GCCCGAAGGAATAGAAGAAGAAGGTGGAGAGAGAGACAGAGACAGAATCCATTCGATTAGTGAACG
GATCTCGACGGTATCGGTTAACTTTTAAAAGAAAAGGGGGGATTGGGGGGTACAGTGCAGGGGAAA
GAATAGTAGACATAATAGCAACAGACATACAAACTAAAGAATTACAAAAACAAATTACAAAAATTCAAAA
TTTTATCGATCACGAGACTAGCCTCGAGAGCTAGCTGCAGTAACGCCATTTTGCAAGGCATGGAAAA
ATACCAAACCAAGAATAGAGAAGTTCAGATCAAGGGCGGGTACATGAAAATAGCTAACGTTGGGCCA
AACAGGATATCTGCGGTGAGCAGTTTCGGCCCCGGCCCGGGGCCAAGAACAGATGGTCACCGCAG
TTTCGGCCCCGGCCCGAGGCCAAGAACAGATGGTCCCCAGATATGGCCCAACCCTCAGCAGTTTC
TTAAGACCCATCAGATGTTTCCAGGCTCCCCCAAGGACCTGAAATGACCCTGCGCCTTATTTGAATT
AACCAATCAGCCTGCTTCTCGCTTCTGTTCGCGCGCTTCTGCTTCCCGAGCTCTATAAAAGAGCTCA

CAACCCCTCACTCGGCGCGCCAGTCCTCCGACAGACTGAGTCGCCCGGGTACCGCGATCGCCATG
GCCCCAAAGAAGAAGCGGAAGGTCGGTATCCACGGAGTCCCAGCAGCCGACAAGAAGTACAGCAT
CGGCCTGGACATCGGCACCAACTCTGTGGGCTGGGCCGTGATCACCGACGAGTACAAGGTGCCCA
GCAAGAAATTCAAGGTGCTGGGCAACACCGACCGGCACAGCATCAAGAAGAACCTGATCGGAGCC
CTGCTGTTCGACAGCGGCGAAACAGCCGAGGCCACCCGGCTGAAGAGAACCGCCAGAAGAAGATA
CACCAGACGGAAGAACCGGATCTGCTATCTGCAAGAGATCTTCAGCAACGAGATGGCCAAGGTGGA
CGACAGCTTCTTCCACAGACTGGAAGAGTCCTTCCTGGTGGAAGAGGATAAGAAGCACGAGCGGC
ACCCCATCTTCGGCAACATCGTGGACGAGGTGGCCTACCACGAGAAGTACCCCACCATCTACCACC
TGAGAAAGAAACTGGTGGACAGCACCGACAAGGCCGACCTGCGGCTGATCTATCTGGCCCTGGCC
CACATGATCAAGTTCCGGGGCCACTTCCTGATCGAGGGCGACCTGAACCCCGACAACAGCGACGT
GGACAAGCTGTTCATCCAGCTGGTGCAGACCTACAACCAGCTGTTCGAGGAAAACCCCATCAACGC
CAGCGGCGTGGACGCCAAGGCCATCCTGTCTGCCAGACTGAGCAAGAGCAGACGGCTGGAAAATC
TGATCGCCCAGCTGCCCGGCGAGAAGAAGAATGGCCTGTTCGGAAACCTGATTGCCCTGAGCCTG
GGCCTGACCCCCAACTTCAAGAGCAACTTCGACCTGGCCGAGGATGCCAAACTGCAGCTGAGCAA
GGACACCTACGACGACGACCTGGACAACCTGCTGGCCCAGATCGGCGACCAGTACGCCGACCTGT
TTCTGGCCGCCAAGAACCTGTCCGACGCCATCCTGCTGAGCGACATCCTGAGAGTGAACACCGAG
ATCACCAAGGCCCCCCTGAGCGCCTCTATGATCAAGAGATACGACGAGCACCACCAGGACCTGACC
CTGCTGAAAGCTCTCGTGCGGCAGCAGCTGCCTGAGAAGTACAAAGAGATTTTCTTCGACCAGAGC
AAGAACGGCTACGCCGGCTACATTGACGGCGGAGCCAGCCAGGAAGAGTTCTACAAGTTCATCAAG
CCCATCCTGGAAAAGATGGACGGCACCGAGGAACTGCTCGTGAAGCTGAACAGAGAGGACCTGCT
GCGGAAGCAGCGGACCTTCGACAACGGCAGCATCCCCCACCAGATCCACCTGGGAGAGCTGCAC
GCCATTCTGCGGCGGCAGGAAGATTTTTACCCATTCCTGAAGGACAACCGGGAAAAGATCGAGAAG
ATCCTGACCTTCCGCATCCCCTACTACGTGGGCCCTCTGGCCAGGGGAAACAGCAGATTCGCCTGG
ATGACCAGAAAGAGCGAGGAAACCATCACCCCCTGGAACTTCGAGGAAGTGGTGGACAAGGGCGC
TTCCGCCCAGAGCTTCATCGAGCGGATGACCAACTTCGATAAGAACCTGCCCAACGAGAAGGTGCT
GCCCAAGCACAGCCTGCTGTACGAGTACTTCACCGTGTATAACGAGCTGACCAAAGTGAAATACGT
GACCGAGGGAATGAGAAAGCCCGCCTTCCTGAGCGGCGAGCAGAAAAAGGCCATCGTGGACCTG
CTGTTCAAGACCAACCGGAAAGTGACCGTGAAGCAGCTGAAAGAGGACTACTTCAAGAAAATCGAG
TGCTTCGACTCCGTGGAAATCTCCGGCGTGGAAGATCGGTTCAACGCCTCCCTGGGCACATACCAC
GATCTGCTGAAAATTATCAAGGACAAGGACTTCCTGGACAATGAGGAAAACGAGGACATTCTGGAAG
ATATCGTGCTGACCCTGACACTGTTTGAGGACAGAGAGATGATCGAGGAACGGCTGAAAACCTATG
CCCACCTGTTCGACGACAAAGTGATGAAGCAGCTGAAGCGGCGGAGATACACCGGCTGGGGCAG
GCTGAGCCGGAAGCTGATCAACGGCATCCGGGACAAGCAGTCCGGCAAGACAATCCTGGATTTCC
TGAAGTCCGACGGCTTCGCCAACAGAAACTTCATGCAGCTGATCCACGACGACAGCCTGACCTTTA
AAGAGGACATCCAGAAAGCCCAGGTGTCCGGCCAGGGCGATAGCCTGCACGAGCACATTGCCAAT
CTGGCCGGCAGCCCCGCCATTAAGAAGGGCATCCTGCAGACAGTGAAGGTGGTGGACGAGCTCGT
GAAAGTGATGGGCCGGCACAAGCCCGAGAACATCGTGATCGAAATGGCCAGAGAGAACCAGACCA
CCCAGAAGGGACAGAAGAACAGCCGCGAGAGAATGAAGCGGATCGAAGAGGGCATCAAAGAGCT
GGGCAGCCAGATCCTGAAAGAACACCCCGTGGAAAACACCCAGCTGCAGAACGAGAAGCTGTACC
TGTACTACCTGCAGAATGGGCGGGATATGTACGTGGACCAGGAACTGGACATCAACCGGCTGTCCG
ACTACGATGTGGACCATATCGTGCCTCAGAGCTTTCTGAAGGACGACTCCATCGACAACAAGGTGC
TGACCAGAAGCGACAAGAACCGGGGCAAGAGCGACAACGTGCCCTCCGAAGAGGTCGTGAAGAA
GATGAAGAACTACTGGCGGCAGCTGCTGAACGCCAAGCTGATTACCCAGAGAAAGTTCGACAATCT
GACCAAGGCCGAGAGAGGCGGCCTGAGCGAACTGGATAAGGCCGGCTTCATCAAGAGACAGCTG

GTGGAAACCCGGCAGATCACAAAGCACGTGGCACAGATCCTGGACTCCCGGATGAACACTAAGTAC
GACGAGAATGACAAGCTGATCCGGGAAGTGAAAGTGATCACCCTGAAGTCCAAGCTGGTGTCCGAT
TTCCGGAAGGATTTCCAGTTTTACAAAGTGCGCGAGATCAACAACTACCACCACGCCCACGACGCC
TACCTGAACGCCGTCGTGGGAACCGCCCTGATCAAAAAGTACCCTAAGCTGGAAAGCGAGTTCGTG
TACGGCGACTACAAGGTGTACGACGTGCGGAAGATGATCGCCAAGAGCGAGCAGGAAATCGGCAA
GGCTACCGCCAAGTACTTCTTCTACAGCAACATCATGAACTTTTTCAAGACCGAGATTACCCTGGCC
AACGGCGAGATCCGGAAGCGGCCTCTGATCGAGACAAACGGCGAAACCGGGGAGATCGTGTGGG
ATAAGGGCCGGGATTTTGCCACCGTGCGGAAAGTGCTGAGCATGCCCCAAGTGAATATCGTGAAAA
AGACCGAGGTGCAGACAGGCGGCTTCAGCAAAGAGTCTATCCTGCCCAAGAGGAACAGCGATAAG
CTGATCGCCAGAAAGAAGGACTGGGACCCTAAGAAGTACGGCGGCTTCTTGTGGCCCACCGTGGC
CTATTCTGTGCTGGTGGTGGCCAAAGTGGAAAAGGGCAAGTCCAAGAAACTGAAGAGTGTGAAAGA
GCTGCTGGGGATCACCATCATGGAAAGAAGCAGCTTCGAGAAGAATCCCATCGACTTTCTGGAAGC
CAAGGGCTACAAAGAAGTGAAAAAGGACCTGATCATCAAGCTGCCTAAGTACTCCCTGTTCGAGCT
GGAAAACGGCCGGAAGAGAATGCTGGCCTCTGCCAAGCAGCTGCAGAAGGGAAACGAACTGGCC
CTGCCCTCCAAATATGTGAACTTCCTGTACCTGGCCAGCCACTATGAGAAGCTGAAGGGCTCCCCC
GAGGATAATGAGCAGAAACAGCTGTTTGTGGAACAGCACAAGCACTACCTGGACGAGATCATCGAG
CAGATCAGCGAGTTCTCCAAGAGAGTGATCCTGGCCGACGCTAATCTGGACAAAGTGCTGTCCGCC
TACAACAAGCACCGGGATAAGCCCATCAGAGAGCAGGCCGAGAATATCATCCACCTGTTTACCCTGA
CCAATCTGGGAGCCCCTGCCGCCTTCAAGTACTTTGACACCACCATCGACCGGAAGCAATACCGAA
GCACCAAAGAGGTGCTGGACGCCACCCTGATCCACCAGAGCATCACCGGCCTGTACGAGACACGG
ATCGACCTGTCTCAGCTGGGAGGCGACAAAAGGCCGGCGGCCACGAAAAAGGCCGGCCAGGCAA
AAAAGAAAAAGACGCGTCAGTGTACTAATTATGCTCTCTTGAAATTGGCTGGAGATGTTGAGAGCAA
CCCAGGTCCCATGACCGAGTACAAGCCCACGGTGCGCCTCGCCACCCGCGACGACGTCCCCAGG
GCCGTACGCACCCTCGCCGCCGCGTTCGCCGACTACCCCGCCACGCGCCACACCGTCGATCCGG
ACCGCCACATCGAGCGGGTCACCGAGCTGCAAGAACTCTTCCTCACGCGCGTCGGGCTCGACATC
GGCAAGGTGTGGGTCGCGGACGACGGCGCCGCGGTGGCGGTCTGGACCACGCCGGAGAGCGTC
GAAGCGGGGGCGGTGTTCGCCGAGATCGGCCCGCGCATGGCCGAGTTGAGCGGTTCCCGGCTG
GCCGCGCAGCAACAGATGGAAGGCCTCCTGGCGCCGCACCGGCCCAAGGAGCCCGCGTGGTTCC
TGGCCACCGTCGGCGTCTCGCCCGACCACCAGGGCAAGGGTCTGGGCAGCGCCGTCGTGCTCCC
CGGAGTGGAGGCGGCCGAGCGCGCCGGGGTGCCCGCCTTCCTGGAGACCTCCGCGCCCCGCAA
CCTCCCCTTCTACGAGCGGCTCGGCTTCACCGTCACCGCCGACGTCGAGGTGCCCGAAGGACCG
CGCACCTGGTGCATGACCCGCAAGCCCGGTGCCTGAGTTTGACAATCAACCTCTGGATTACAAAAT
TTGTGAAAGATTGACTGGTATTCTTAACTATGTTGCTCCTTTTACGCTATGTGGATACGCTGCTTTAAT
GCCTTTGTATCATGCTATTGCTTCCCGTATGGCTTTCATTTTCTCCTCCTTGTATAAATCCTGGTTGCT
GTCTCTTTATGAGGAGTTGTGGCCCGTTGTCAGGCAACGTGGCGTGGTGTGCACTGTGTTTGCTGA
CGCAACCCCCACTGGTTGGGGCATTGCCACCACCTGTCAGCTCCTTTCCGGGACTTTCGCTTTCCC
CCTCCCTATTGCCACGGCGGAACTCATCGCCGCCTGCCTTGCCCGCTGCTGGACAGGGGCTCGGC
TGTTGGGCACTGACAATTCCGTGGTGTTGTCGGGGAAGCTGACGTCCTTTCCATGGCTGCTCGCCT
GTGTTGCCACCTGGATTCTGCGCGGGACGTCCTTCTGCTACGTCCCTTCGGCCCTCAATCCAGCG
GACCTTCCTTCCCGCGGCCTGCTGCCGGCTCTGCGGCCTCTTCCGCGTCTCGCCTTCGCCCTCAG
ACGAGTCGGATCTCCCTTTGGGCCGCCTCCCCGCCTGGAATTCGAGCTCGGTACCTTTAAGACCAA
TGACTTACAAGGCAGCTGTAGATCTTAGCCACTTTTTAAAAGAAAAGGGGGGACTGGAAGGGCTAAT
TCACTCCCAACGAGACAAGATCTGCTTTTTGCTTGTACTGGGTCTCTCTGGTTAGACCAGAATCTGA
GCCTGGGAGCTCTCTGGCTAACTAGGGAACCCACTGCTTAAGCCTCAATAAAGCTTGCCTTGAGTG

CTTCAAGTAGTGTGTGCCCGTCTGTTGTGTGACTCTGGTAACTAGAGAATCCCTCAGACCCTTTTAG
TCAGTGTGGAAAATCTCTAGCAGTAGTAGTTCATGTCATCTTATTATTCAGTATTTATAACTTGCAAAGA
AATGAATATCAGAGAGTGAGAGGAACTTGTTTATTGCAGCTTATAATGGTTACAAATAAAGCAATAGCA
TCACAAATTTCACAAATAAAGCATTTTTTTCACTGCATTCTAGTTGTGGTTTGTCCAAACTCATCAATG
TATCTTATCATGTCTGGCTCTAGCTATCCCGCCCCTAACTCCGCCCATCCCGCCCCTAACTCCGCCC
AGTTCCGCCCATTCTCCGCCCCATGGCTGACTAATTTTTTTTATTTATGCAGAGGCCGAGGCCGCCT
CGGCCTCTGAGCTATTCCAGAAGTAGTGAGGAGGCTTTTTTGGAGGCCTAGGGACGTACCCAATTC
GCCCTATAGTGAGTCGTATTACGCGCGCTCACTGGCCGTCGTTTTACAACGTCGTGACTGGGAAAA
CCCTGGCGTTACCCAACTTAATCGCCTTGCAGCACATCCCCCTTTCGCCAGCTGGCGTAATAGCGA
AGAGGCCCGCACCGATCGCCCTTCCCAACAGTTGCGCAGCCTGAATGGCGAATGGGACGCGCCCT
GTAGCGGCGCATTAAGCGCGGCGGGTGTGGTGGTTACGCGCAGCGTGACCGCTACACTTGCCAGC
GCCCTAGCGCCCGCTCCTTTCGCTTTCTTCCCTTCCTTTCTCGCCACGTTCGCCGGCTTTCCCCGT
CAAGCTCTAAATCGGGGGCTCCCTTTAGGGTTCCGATTTAGTGCTTTACGGCACCTCGACCCCAAAA
AACTTGATTAGGGTGATGGTTCACGTAGTGGGCCATCGCCCTGATAGACGGTTTTTCGCCCTTTGAC
GTTGGAGTCCACGTTCTTTAATAGTGGACTCTTGTTCCAAACTGGAACAACACTCAACCCTATCTCG
GTCTATTCTTTTGATTTATAAGGGATTTTGCCGATTTCGGCCTATTGGTTAAAAAATGAGCTGATTTAAC
AAAAATTTAACGCGAATTTTAACAAAATATTAACGCTTACAATTTAGGTGGCACTTTTCGGGGAAATGT
GCGCGGAACCCCTATTTGTTTATTTTTCTAAATACATTCAAATATGTATCCGCTCATGAGACAATAACC
CTGATAAATGCTTCAATAATATTGAAAAAGGAAGAGTATGAGTATTCAACATTTCCGTGTCGCCCTTAT
TCCCTTTTTTGCGGCATTTTGCCTTCCTGTTTTTGCTCACCCAGAAACGCTGGTGAAAGTAAAAGAT
GCTGAAGATCAGTTGGGTGCACGAGTGGGTTACATCGAACTGGATCTCAACAGCGGTAAGATCCTT
GAGAGTTTTCGCCCCGAAGAACGTTTTCCAATGATGAGCACTTTTAAAGTTCTGCTATGTGGCGCGG
TATTATCCCGTATTGACGCCGGGCAAGAGCAACTCGGTCGCCGCATACACTATTCTCAGAATGACTT
GGTTGAGTACTCACCAGTCACAGAAAAGCATCTTACGGATGGCATGACAGTAAGAGAATTATGCAGT
GCTGCCATAACCATGAGTGATAACACTGCGGCCAACTTACTTCTGACAACGATCGGAGGACCGAAG
GAGCTAACCGCTTTTTTGCACAACATGGGGGATCATGTAACTCGCCTTGATCGTTGGGAACCGGAG
CTGAATGAAGCCATACCAAACGACGAGCGTGACACCACGATGCCTGTAGCAATGGCAACAACGTTG
CGCAAACTATTAACTGGCGAACTACTTACTCTAGCTTCCCGGCAACAATTAATAGACTGGATGGAGGC
GGATAAAGTTGCAGGACCACTTCTGCGCTCGGCCCTTCCGGCTGGCTGGTTTATTGCTGATAAATCT
GGAGCCGGTGAGCGTGGGTCTCGCGGTATCATTGCAGCACTGGGGCCAGATGGTAAGCCCTCCCG
TATCGTAGTTATCTACACGACGGGGAGTCAGGCAACTATGGATGAACGAAATAGACAGATCGCTGAG
ATAGGTGCCTCACTGATTAAGCATTGGTAACTGTCAGACCAAGTTTACTCATATATACTTTAGATTGATT
TAAAACTTCATTTTTAATTTAAAAGGATCTAGGTGAAGATCCTTTTTGATAATCTCATGACCAAAATCCC
TTAACGTGAGTTTTCGTTCCACTGAGCGTCAGACCCCGTAGAAAAGATCAAAGGATCTTCTTGAGAT
CCTTTTTTTCTGCGCGTAATCTGCTGCTTGCAAACAAAAAAACCACCGCTACCAGCGGTGGTTTGTT
TGCCGGATCAAGAGCTACCAACTCTTTTTCCGAAGGTAACTGGCTTCAGCAGAGCGCAGATACCAAA
TACTGTTCTTCTAGTGTAGCCGTAGTTAGGCCACCACTTCAAGAACTCTGTAGCACCGCCTACATACC
TCGCTCTGCTAATCCTGTTACCAGTGGCTGCTGCCAGTGGCGATAAGTCGTGTCTTACCGGGTTGG
ACTCAAGACGATAGTTACCGGATAAGGCGCAGCGGTCGGGCTGAACGGGGGGTTCGTGCACACAG
CCCAGCTTGGAGCGAACGACCTACACCGAACTGAGATACCTACAGCGTGAGCTATGAGAAAGCGCC
ACGCTTCCCGAAGGGAGAAAGGCGGACAGGTATCCGGTAAGCGGCAGGGTCGGAACAGGAGAGC
GCACGAGGGAGCTTCCAGGGGGAAACGCCTGGTATCTTTATAGTCCTGTCGGGTTTCGCCACCTCT
GACTTGAGCGTCGATTTTTGTGATGCTCGTCAGGG

P4：

>P2141 pAdHM4-SFFV-OCT4Cherry-cutted

GGGGGGTTCGTGCACACAGCCCAGCTTGGAGCGAACGACCTACACCGAACTGAGATACCTACAGC
GTGAGCTATGAGAAAGCGCCACGCTTCCCGAAGGGAGAAAGGCGGACAGGTATCCGGTAAGCGGC
AGGGTCGGAACAGGAGAGCGCACGAGGGAGCTTCCAGGGGGAAACGCCTGGTATCTTTATAGTCC
TGTCGGGTTTCGCCACCTCTGACTTGAGCGTCGATTTTTGTGATGCTCGTCAGGGGGGCGGAGCCT
ATGGAAAAACGCCAGCAACGCGGCCTTTTTACGGTTCCTGGCCTTTTGCTGGCCTTTTGCTCACAT
GTTCTTTCCTGCGTTATCCCCTGATTCTGTGGATAACCGTATTACCGCCTTTGAGTGAGCTGATACCG
CTCGCCGCAGCCGAACGACCGAGCGCAGCGAGTCAGTGAGCGAGGAAGCGGAAGAGCGCCCAAT
ACGCAAACCGCCTCTCCCCGCGCGTTGGCCGATTCATTAATGCAGCTGGCACGACAGGTTTCCCGA
CTGGAAAGCGGGCAGTGAGCGCAACGCAATTAATGTGAGTTAGCTCACTCATTAGGCACCCCAGGC
TTTACACTTTATGCTTCCGGCTCGTATGTTGTGTGGAATTGTGAGCGGATAACAATTTCACACAGGAA
ACAGCTATGACCATGATTACGCCAAGCTATTTAGGTGACACTATAGAATACTCAAGCTAGTTAATTAAC
GTTAATTAACATCATCAATAATATACCTTATTTTGGATTGAAGCCAATATGATAATGAGGGGGTGGAGTT
TGTGACGTGGCGCGGGGCGTGGGAACGGGGCGGGTGACGTAGTAGTGTGGCGGAAGTGTGATGT
TGCAAGTGTGGCGGAACACATGTAAGCGACGGATGTGGCAAAAGTGACGTTTTTGGTGTGCGCCG
GTGTACACAGGAAGTGACAATTTTCGCGCGGTTTTAGGCGGATGTTGTAGTAAATTTGGGCGTAACC
GAGTAAGATTTGGCCATTTTCGCGGGAAAACTGAATAAGAGGAAGTGAAATCTGAATAATTTTGTGTT
ACTCATAGCGCGTAATCTCTAGCATCGTAACTATAACGGTCCTAAGGTAGCGAGAATTATTTAAATATCT
ATGTCGGGTGCGGAGAAAGAGGTAATGAAATGGCATCGACTCGAAGATCTGGGCGTGGTTAAGGGT
GGGAAAGAATATATAAGGTGGGGGTCTTATGTAGTTTTGTATCTGTTTTGCAGCAGCCGCCGCCGCC
ATGAGCACCAACTCGTTTGATGGAAGCATTGTGAGCTCATATTTGACAACGCGCATGCCCCCATGGG
CCGGGGTGCGTCAGAATGTGATGGGCTCCAGCATTGATGGTCGCCCCGTCCTGCCCGCAAACTCT
ACTACCTTGACCTACGAGACCGTGTCTGGAACGCCGTTGGAGACTGCAGCCTCCGCCGCCGCTTC
AGCCGCTGCAGCCACCGCCCGCGGGATTGTGACTGACTTTGCTTTCCTGAGCCCGCTTGCAAGCA
GTGCAGCTTCCCGTTCATCCGCCCGCGATGACAAGTTGACGGCTCTTTTGGCACAATTGGATTCTTT
GACCCGGGAACTTAATGTCGTTTCTCAGCAGCTGTTGGATCTGCGCCAGCAGGTTTCTGCCCTGAA
GGCTTCCTCCCCTCCCAATGCGGTTTAAAACATAAATAAAAAACCAGACTCTGTTTGGATTTGGATCA
AGCAAGTGTCTTGCTGTCTTTATTTAGGGGTTTTGCGCGCGCGGTAGGCCCGGGACCAGCGGTCTC
GGTCGTTGAGGGTCCTGTGTATTTTTTCCAGGACGTGGTAAAGGTGACTCTGGATGTTCAGATACAT
GGGCATAAGCCCGTCTCTGGGGTGGAGGTAGCACCACTGCAGAGCTTCATGCTGCGGGGTGGTGT
TGTAGATGATCCAGTCGTAGCAGGAGCGCTGGGCGTGGTGCCTAAAAATGTCTTTCAGTAGCAAGC
TGATTGCCAGGGGCAGGCCCTTGGTGTAAGTGTTTACAAAGCGGTTAAGCTGGGATGGGTGCATAC
GTGGGGATATGAGATGCATCTTGGACTGTATTTTTAGGTTGGCTATGTTCCCAGCCATATCCCTCCGG
GGATTCATGTTGTGCAGAACCACCAGCACAGTGTATCCGGTGCACTTGGGAAATTTGTCATGTAGCT
TAGAAGGAAATGCGTGGAAGAACTTGGAGACGCCCTTGTGACCTCCAAGATTTTCCATGCATTCGTC
CATAATGATGGCAATGGGCCCACGGGCGGCGGCCTGGGCGAAGATATTTCTGGGATCACTAACGTC
ATAGTTGTGTTCCAGGATGAGATCGTCATAGGCCATTTTTACAAAGCGCGGGCGGAGGGTGCCAGA
CTGCGGTATAATGGTTCCATCCGGCCCAGGGGCGTAGTTACCCTCACAGATTTGCATTTCCCACGCT
TTGAGTTCAGATGGGGGGATCATGTCTACCTGCGGGGCGATGAAGAAAACGGTTTCCGGGGTAGG
GGAGATCAGCTGGGAAGAAAGCAGGTTCCTGAGCAGCTGCGACTTACCGCAGCCGGTGGGCCCG
TAAATCACACCTATTACCGGCTGCAACTGGTAGTTAAGAGAGCTGCAGCTGCCGTCATCCCTGAGCA
GGGGGGCCACTTCGTTAAGCATGTCCCTGACTCGCATGTTTTCCCTGACCAAATCCGCCAGAAGGC
GCTCGCCGCCCAGCGATAGCAGTTCTTGCAAGGAAGCAAAGTTTTTCAACGGTTTGAGACCGTCCG

CCGTAGGCATGCTTTTGAGCGTTTGACCAAGCAGTTCCAGGCGGTCCCACAGCTCGGTCACCTGCT
CTACGGCATCTCGATCCAGCATATCTCCTCGTTTCGCGGGTTGGGGCGGCTTTCGCTGTACGGCAG
TAGTCGGTGCTCGTCCAGACGGGCCAGGGTCATGTCTTTCCACGGGCGCAGGGTCCTCGTCAGCG
TAGTCTGGGTCACGGTGAAGGGGTGCGCTCCGGGCTGCGCGCTGGCCAGGGTGCGCTTGAGGCT
GGTCCTGCTGGTGCTGAAGCGCTGCCGGTCTTCGCCCTGCGCGTCGGCCAGGTAGCATTTGACCA
TGGTGTCATAGTCCAGCCCCTCCGCGGCGTGGCCCTTGGCGCGCAGCTTGCCCTTGGAGGAGGC
GCCGCACGAGGGGCAGTGCAGACTTTTGAGGGCGTAGAGCTTGGGCGCGAGAAATACCGATTCCG
GGGAGTAGGCATCCGCGCCGCAGGCCCCGCAGACGGTCTCGCATTCCACGAGCCAGGTGAGCTC
TGGCCGTTCGGGGTCAAAAACCAGGTTTCCCCCATGCTTTTTGATGCGTTTCTTACCTCTGGTTTCC
ATGAGCCGGTGTCCACGCTCGGTGACGAAAAGGCTGTCCGTGTCCCCGTATACAGACTTGAGAGG
CCTGTCCTCGAGCGGTGTTCCGCGGTCCTCCTCGTATAGAAACTCGGACCACTCTGAGACAAAGGC
TCGCGTCCAGGCCAGCACGAAGGAGGCTAAGTGGGAGGGGTAGCGGTCGTTGTCCACTAGGGGG
TCCACTCGCTCCAGGGTGTGAAGACACATGTCGCCCTCTTCGGCATCAAGGAAGGTGATTGGTTTG
TAGGTGTAGGCCACGTGACCGGGTGTTCCTGAAGGGGGGCTATAAAAGGGGGTGGGGGCGCGTT
CGTCCTCACTCTCTTCCGCATCGCTGTCTGCGAGGGCCAGCTGTTGGGGTGAGTACTCCCTCTGAA
AAGCGGGCATGACTTCTGCGCTAAGATTGTCAGTTTCCAAAAACGAGGAGGATTTGATATTCACCTG
GCCCGCGGTGATGCCTTTGAGGGTGGCCGCATCCATCTGGTCAGAAAAGACAATCTTTTTGTTGTC
AAGCTTGGTGGCAAACGACCCGTAGAGGGCGTTGGACAGCAACTTGGCGATGGAGCGCAGGGTTT
GGTTTTTGTCGCGATCGGCGCGCTCCTTGGCCGCGATGTTTAGCTGCACGTATTCGCGCGCAACGC
ACCGCCATTCGGGAAAGACGGTGGTGCGCTCGTCGGGCACCAGGTGCACGCGCCAACCGCGGTT
GTGCAGGGTGACAAGGTCAACGCTGGTGGCTACCTCTCCGCGTAGGCGCTCGTTGGTCCAGCAGA
GGCGGCCGCCCTTGCGCGAGCAGAATGGCGGTAGGGGGTCTAGCTGCGTCTCGTCCGGGGGGTC
TGCGTCCACGGTAAAGACCCCGGGCAGCAGGCGCGCGTCGAAGTAGTCTATCTTGCATCCTTGCAA
GTCTAGCGCCTGCTGCCATGCGCGGGCGGCAAGCGCGCGCTCGTATGGGTTGAGTGGGGGACCC
CATGGCATGGGGTGGGTGAGCGCGGAGGCGTACATGCCGCAAATGTCGTAAACGTAGAGGGGCTC
TCTGAGTATTCCAAGATATGTAGGGTAGCATCTTCCACCGCGGATGCTGGCGCGCACGTAATCGTAT
AGTTCGTGCGAGGGAGCGAGGAGGTCGGGACCGAGGTTGCTACGGGCGGGCTGCTCTGCTCGGA
AGACTATCTGCCTGAAGATGGCATGTGAGTTGGATGATATGGTTGGACGCTGGAAGACGTTGAAGCT
GGCGTCTGTGAGACCTACCGCGTCACGCACGAAGGAGGCGTAGGAGTCGCGCAGCTTGTTGACCA
GCTCGGCGGTGACCTGCACGTCTAGGGCGCAGTAGTCCAGGGTTTCCTTGATGATGTCATACTTAT
CCTGTCCCTTTTTTTTCCACAGCTCGCGGTTGAGGACAAACTCTTCGCGGTCTTTCCAGTACTCTTG
GATCGGAAACCCGTCGGCCTCCGAACGGTAAGAGCCTAGCATGTAGAACTGGTTGACGGCCTGGT
AGGCGCAGCATCCCTTTTCTACGGGTAGCGCGTATGCCTGCGCGGCCTTCCGGAGCGAGGTGTGG
GTGAGCGCAAAGGTGTCCCTGACCATGACTTTGAGGTACTGGTATTTGAAGTCAGTGTCGTCGCAT
CCGCCCTGCTCCCAGAGCAAAAAGTCCGTGCGCTTTTTGGAACGCGGATTTGGCAGGGCGAAGGT
GACATCGTTGAAGAGTATCTTTCCCGCGCGAGGCATAAAGTTGCGTGTGATGCGGAAGGGTCCCGG
CACCTCGGAACGGTTGTTAATTACCTGGGCGGCGAGCACGATCTCGTCAAAGCCGTTGATGTTGTG
GCCCACAATGTAAAGTTCCAAGAAGCGCGGGATGCCCTTGATGGAAGGCAATTTTTTAAGTTCCTCG
TAGGTGAGCTCTTCAGGGGAGCTGAGCCCGTGCTCTGAAAGGGCCCAGTCTGCAAGATGAGGGTT
GGAAGCGACGAATGAGCTCCACAGGTCACGGGCCATTAGCATTTGCAGGTGGTCGCGAAAGGTCC
TAAACTGGCGACCTATGGCCATTTTTTCTGGGGTGATGCAGTAGAAGGTAAGCGGGTCTTGTTCCCA
GCGGTCCCATCCAAGGTTCGCGGCTAGGTCTCGCGCGGCAGTCACTAGAGGCTCATCTCCGCCGA
ACTTCATGACCAGCATGAAGGGCACGAGCTGCTTCCCAAAGGCCCCCATCCAAGTATAGGTCTCTA
CATCGTAGGTGACAAAGAGACGCTCGGTGCGAGGATGCGAGCCGATCGGGAAGAACTGGATCTCC

CGCCACCAATTGGAGGAGTGGCTATTGATGTGGTGAAAGTAGAAGTCCCTGCGACGGGCCGAACA
CTCGTGCTGGCTTTTGTAAAAACGTGCGCAGTACTGGCAGCGGTGCACGGGCTGTACATCCTGCAC
GAGGTTGACCTGACGACCGCGCACAAGGAAGCAGAGTGGGAATTTGAGCCCCTCGCCTGGCGGG
TTTGGCTGGTGGTCTTCTACTTCGGCTGCTTGTCCTTGACCGTCTGGCTGCTCGAGGGGAGTTACG
GTGGATCGGACCACCACGCCGCGCGAGCCCAAAGTCCAGATGTCCGCGCGCGGCGGTCGGAGCT
TGATGACAACATCGCGCAGATGGGAGCTGTCCATGGTCTGGAGCTCCCGCGGCGTCAGGTCAGGC
GGGAGCTCCTGCAGGTTTACCTCGCATAGACGGGTCAGGGCGCGGGCTAGATCCAGGTGATACCT
AATTTCCAGGGGCTGGTTGGTGGCGGCGTCGATGGCTTGCAAGAGGCCGCATCCCCGCGGCGCG
ACTACGGTACCGCGCGGCGGGCGGTGGGCCGCGGGGGTGTCCTTGGATGATGCATCTAAAAGCG
GTGACGCGGGCGAGCCCCCGGAGGTAGGGGGGGCTCCGGACCCGCCGGGAGAGGGGGCAGGG
GCACGTCGGCGCCGCGCGCGGGCAGGAGCTGGTGCTGCGCGCGTAGGTTGCTGGCGAACGCGA
CGACGCGGCGGTTGATCTCCTGAATCTGGCGCCTCTGCGTGAAGACGACGGGCCCGGTGAGCTT
GAACCTGAAAGAGAGTTCGACAGAATCAATTTCGGTGTCGTTGACGGCGGCCTGGCGCAAAATCTC
CTGCACGTCTCCTGAGTTGTCTTGATAGGCGATCTCGGCCATGAACTGCTCGATCTCTTCCTCCTGG
AGATCTCCGCGTCCGGCTCGCTCCACGGTGGCGGCGAGGTCGTTGGAAATGCGGGCCATGAGCT
GCGAGAAGGCGTTGAGGCCTCCCTCGTTCCAGACGCGGCTGTAGACCACGCCCCCTTCGGCATCG
CGGGCGCGCATGACCACCTGCGCGAGATTGAGCTCCACGTGCCGGGCGAAGACGGCGTAGTTTC
GCAGGCGCTGAAAGAGGTAGTTGAGGGTGGTGGCGGTGTGTTCTGCCACGAAGAAGTACATAACC
CAGCGTCGCAACGTGGATTCGTTGATATCCCCCAAGGCCTCAAGGCGCTCCATGGCCTCGTAGAAG
TCCACGGCGAAGTTGAAAAACTGGGAGTTGCGCGCCGACACGGTTAACTCCTCCTCCAGAAGACG
GATGAGCTCGGCGACAGTGTCGCGCACCTCGCGCTCAAAGGCTACAGGGGCCTCTTCTTCTTCTT
CAATCTCCTCTTCCATAAGGGCCTCCCCTTCTTCTTCTTCTGGCGGCGGTGGGGGAGGGGGGACA
CGGCGGCGACGACGGCGCACCGGGAGGCGGTCGACAAAGCGCTCGATCATCTCCCCGCGGCGA
CGGCGCATGGTCTCGGTGACGGCGCGGCCGTTCTCGCGGGGGCGCAGTTGGAAGACGCCGCCC
GTCATGTCCCGGTTATGGGTTGGCGGGGGGCTGCCATGCGGCAGGGATACGGCGCTAACGATGCA
TCTCAACAATTGTTGTGTAGGTACTCCGCCGCCGAGGGACCTGAGCGAGTCCGCATCGACCGGATC
GGAAAACCTCTCGAGAAAGGCGTCTAACCAGTCACAGTCGCAAGGTAGGCTGAGCACCGTGGCGG
GCGGCAGCGGGCGGCGGTCGGGGTTGTTTCTGGCGGAGGTGCTGCTGATGATGTAATTAAAGTAG
GCGGTCTTGAGACGGCGGATGGTCGACAGAAGCACCATGTCCTTGGGTCCGGCCTGCTGAATGCG
CAGGCGGTCGGCCATGCCCCAGGCTTCGTTTTGACATCGGCGCAGGTCTTTGTAGTAGTCTTGCAT
GAGCCTTTCTACCGGCACTTCTTCTTCTCCTTCCTCTTGTCCTGCATCTCTTGCATCTATCGCTGCGG
CGGCGGCGGAGTTTGGCCGTAGGTGGCGCCCTCTTCCTCCCATGCGTGTGACCCCGAAGCCCCT
CATCGGCTGAAGCAGGGCTAGGTCGGCGACAACGCGCTCGGCTAATATGGCCTGCTGCACCTGCG
TGAGGGTAGACTGGAAGTCATCCATGTCCACAAAGCGGTGGTATGCGCCCGTGTTGATGGTGTAAG
TGCAGTTGGCCATAACGGACCAGTTAACGGTCTGGTGACCCGGCTGCGAGAGCTCGGTGTACCTG
AGACGCGAGTAAGCCCTCGAGTCAAATACGTAGTCGTTGCAAGTCCGCACCAGGTACTGGTATCCC
ACCAAAAAGTGCGGCGGCGGCTGGCGGTAGAGGGGCCAGCGTAGGGTGGCCGGGGCTCCGGGG
GCGAGATCTTCCAACATAAGGCGATGATATCCGTAGATGTACCTGGACATCCAGGTGATGCCGGCGG
CGGTGGTGGAGGCGCGCGGAAAGTCGCGGACGCGGTTCCAGATGTTGCGCAGCGGCAAAAAGTG
CTCCATGGTCGGGACGCTCTGGCCGGTCAGGCGCGCGCAATCGTTGACGCTCTAGCGTGCAAAAG
GAGAGCCTGTAAGCGGGCACTCTTCCGTGGTCTGGTGGATAAATTCGCAAGGGTATCATGGCGGAC
GACCGGGGTTCGAGCCCCGTATCCGGCCGTCCGCCGTGATCCATGCGGTTACCGCCCGCGTGTCG
AACCCAGGTGTGCGACGTCAGACAACGGGGGAGTGCTCCTTTTGGCTTCCTTCCAGGCGCGGCG
GCTGCTGCGCTAGCTTTTTTGGCCACTGGCCGCGCGCAGCGTAAGCGGTTAGGCTGGAAAGCGAA

AGCATTAAGTGGCTCGCTCCCTGTAGCCGGAGGGTTATTTTCCAAGGGTTGAGTCGCGGGACCCCC
GGTTCGAGTCTCGGACCGGCCGGACTGCGGCGAACGGGGGTTTGCCTCCCCGTCATGCAAGACC
CCGCTTGCAAATTCCTCCGGAAACAGGGACGAGCCCCTTTTTTGCTTTTCCCAGATGCATCCGGTG
CTGCGGCAGATGCGCCCCCCTCCTCAGCAGCGGCAAGAGCAAGAGCAGCGGCAGACATGCAGGG
CACCCTCCCCTCCTCCTACCGCGTCAGGAGGGGCGACATCCGCGGTTGACGCGGCAGCAGATGG
TGATTACGAACCCCCGCGGCGCCGGGCCCGGCACTACCTGGACTTGGAGGAGGGCGAGGGCCTG
GCGCGGCTAGGAGCGCCCTCTCCTGAGCGGCACCCAAGGGTGCAGCTGAAGCGTGATACGCGTG
AGGCGTACGTGCCGCGGCAGAACCTGTTTCGCGACCGCGAGGGAGAGGAGCCCGAGGAGATGCG
GGATCGAAAGTTCCACGCAGGGCGCGAGCTGCGGCATGGCCTGAATCGCGAGCGGTTGCTGCGC
GAGGAGGACTTTGAGCCCGACGCGCGAACCGGGATTAGTCCCGCGCGCGCACACGTGGCGGCCG
CCGACCTGGTAACCGCATACGAGCAGACGGTGAACCAGGAGATTAACTTTCAAAAAAGCTTTAACAA
CCACGTGCGTACGCTTGTGGCGCGCGAGGAGGTGGCTATAGGACTGATGCATCTGTGGGACTTTGT
AAGCGCGCTGGAGCAAAACCCAAATAGCAAGCCGCTCATGGCGCAGCTGTTCCTTATAGTGCAGCA
CAGCAGGGACAACGAGGCATTCAGGGATGCGCTGCTAAACATAGTAGAGCCCGAGGGCCGCTGGC
TGCTCGATTTGATAAACATCCTGCAGAGCATAGTGGTGCAGGAGCGCAGCTTGAGCCTGGCTGACA
AGGTGGCCGCCATCAACTATTCCATGCTTAGCCTGGGCAAGTTTTACGCCCGCAAGATATACCATAC
CCCTTACGTTCCCATAGACAAGGAGGTAAAGATCGAGGGGTTCTACATGCGCATGGCGCTGAAGGT
GCTTACCTTGAGCGACGACCTGGGCGTTTATCGCAACGAGCGCATCCACAAGGCCGTGAGCGTGA
GCCGGCGGCGCGAGCTCAGCGACCGCGAGCTGATGCACAGCCTGCAAAGGGCCCTGGCTGGCA
CGGGCAGCGGCGATAGAGAGGCCGAGTCCTACTTTGACGCGGGCGCTGACCTGCGCTGGGCCCC
AAGCCGACGCGCCCTGGAGGCAGCTGGGGCCGGACCTGGGCTGGCGGTGGCACCCGCGCGCGC
TGGCAACGTCGGCGGCGTGGAGGAATATGACGAGGACGATGAGTACGAGCCAGAGGACGGCGAG
TACTAAGCGGTGATGTTTCTGATCAGATGATGCAAGACGCAACGGACCCGGCGGTGCGGGCGGCG
CTGCAGAGCCAGCCGTCCGGCCTTAACTCCACGGACGACTGGCGCCAGGTCATGGACCGCATCAT
GTCGCTGACTGCGCGCAATCCTGACGCGTTCCGGCAGCAGCCGCAGGCCAACCGGCTCTCCGCA
ATTCTGGAAGCGGTGGTCCCGGCGCGCGCAAACCCCACGCACGAGAAGGTGCTGGCGATCGTAAA
CGCGCTGGCCGAAAACAGGGCCATCCGGCCCGACGAGGCCGGCCTGGTCTACGACGCGCTGCTT
CAGCGCGTGGCTCGTTACAACAGCGGCAACGTGCAGACCAACCTGGACCGGCTGGTGGGGGATG
TGCGCGAGGCCGTGGCGCAGCGTGAGCGCGCGCAGCAGCAGGGCAACCTGGGCTCCATGGTTG
CACTAAACGCCTTCCTGAGTACACAGCCCGCCAACGTGCCGCGGGGACAGGAGGACTACACCAAC
TTTGTGAGCGCACTGCGGCTAATGGTGACTGAGACACCGCAAAGTGAGGTGTACCAGTCTGGGCC
AGACTATTTTTTCCAGACCAGTAGACAAGGCCTGCAGACCGTAAACCTGAGCCAGGCTTTCAAAAAC
TTGCAGGGGCTGTGGGGGGTGCGGGCTCCCACAGGCGACCGCGCGACCGTGTCTAGCTTGCTGA
CGCCCAACTCGCGCCTGTTGCTGCTGCTAATAGCGCCCTTCACGGACAGTGGCAGCGTGTCCCGG
GACACATACCTAGGTCACTTGCTGACACTGTACCGCGAGGCCATAGGTCAGGCGCATGTGGACGAG
CATACTTTCCAGGAGATTACAAGTGTCAGCCGCGCGCTGGGGCAGGAGGACACGGGCAGCCTGGA
GGCAACCCTAAACTACCTGCTGACCAACCGGCGGCAGAAGATCCCCTCGTTGCACAGTTTAAACAG
CGAGGAGGAGCGCATTTTGCGCTACGTGCAGCAGAGCGTGAGCCTTAACCTGATGCGCGACGGGG
TAACGCCCAGCGTGGCGCTGGACATGACCGCGCGCAACATGGAACCGGGCATGTATGCCTCAAAC
CGGCCGTTTATCAACCGCCTAATGGACTACTTGCATCGCGCGGCCGCCGTGAACCCCGAGTATTTC
ACCAATGCCATCTTGAACCCGCACTGGCTACCGCCCCCTGGTTTCTACACCGGGGGATTCGAGGTG
CCCGAGGGTAACGATGGATTCCTCTGGGACGACATAGACGACAGCGTGTTTTCCCCGCAACCGCA
GACCCTGCTAGAGTTGCAACAGCGCGAGCAGGCAGAGGCGGCGCTGCGAAAGGAAAGCTTCCGC
AGGCCAAGCAGCTTGTCCGATCTAGGCGCTGCGGCCCCGCGGTCAGATGCTAGTAGCCCATTTCC

AAGCTTGATAGGGTCTCTTACCAGCACTCGCACCACCCGCCCGCGCCTGCTGGGCGAGGAGGAGT
ACCTAAACAACTCGCTGCTGCAGCCGCAGCGCGAAAAAAACCTGCCTCCGGCATTTCCCAACAACG
GGATAGAGAGCCTAGTGGACAAGATGAGTAGATGGAAGACGTACGCGCAGGAGCACAGGGACGTG
CCAGGCCCGCGCCCGCCCACCCGTCGTCAAAGGCACGACCGTCAGCGGGGTCTGGTGTGGGAG
GACGATGACTCGGCAGACGACAGCAGCGTCCTGGATTTGGGAGGGAGTGGCAACCCGTTTGCGCA
CCTTCGCCCCAGGCTGGGGAGAATGTTTTAAAAAAAAAAAAGCATGATGCAAAATAAAAAACTCACC
AAGGCCATGGCACCGAGCGTTGGTTTTCTTGTATTCCCCTTAGTATGCGGCGCGCGGCGATGTATG
AGGAAGGTCCTCCTCCCTCCTACGAGAGTGTGGTGAGCGCGGCGCCAGTGGCGGCGGCGCTGGG
TTCTCCCTTCGATGCTCCCCTGGACCCGCCGTTTGTGCCTCCGCGGTACCTGCGGCCTACCGGGG
GGAGAAACAGCATCCGTTACTCTGAGTTGGCACCCCTATTCGACACCACCCGTGTGTACCTGGTGG
ACAACAAGTCAACGGATGTGGCATCCCTGAACTACCAGAACGACCACAGCAACTTTCTGACCACGG
TCATTCAAAACAATGACTACAGCCCGGGGGAGGCAAGCACACAGACCATCAATCTTGACGACCGGT
CGCACTGGGGCGGCGACCTGAAAACCATCCTGCATACCAACATGCCAAATGTGAACGAGTTCATGT
TTACCAATAAGTTTAAGGCGCGGGTGATGGTGTCGCGCTTGCCTACTAAGGACAATCAGGTGGAGC
TGAAATACGAGTGGGTGGAGTTCACGCTGCCCGAGGGCAACTACTCCGAGACCATGACCATAGACC
TTATGAACAACGCGATCGTGGAGCACTACTTGAAAGTGGGCAGACAGAACGGGGTTCTGGAAAGC
GACATCGGGGTAAAGTTTGACACCCGCAACTTCAGACTGGGGTTTGACCCCGTCACTGGTCTTGTC
ATGCCTGGGGTATATACAAACGAAGCCTTCCATCCAGACATCATTTTGCTGCCAGGATGCGGGGTGG
ACTTCACCCACAGCCGCCTGAGCAACTTGTTGGGCATCCGCAAGCGGCAACCCTTCCAGGAGGGC
TTTAGGATCACCTACGATGATCTGGAGGGTGGTAACATTCCCGCACTGTTGGATGTGGACGCCTACC
AGGCGAGCTTGAAAGATGACACCGAACAGGGCGGGGGTGGCGCAGGCGGCAGCAACAGCAGTG
GCAGCGGCGCGGAAGAGAACTCCAACGCGGCAGCCGCGGCAATGCAGCCGGTGGAGGACATGAA
CGATCATGCCATTCGCGGCGACACCTTTGCCACACGGGCTGAGGAGAAGCGCGCTGAGGCCGAA
GCAGCGGCCGAAGCTGCCGCCCCCGCTGCGCAACCCGAGGTCGAGAAGCCTCAGAAGAAACCGG
TGATCAAACCCCTGACAGAGGACAGCAAGAAACGCAGTTACAACCTAATAAGCAATGACAGCACCTT
CCCCAGTACCGCAGCTGGTACCTTGCATACAACTACGGCGACCCTCAGACCGGAATCCGCTCATGG
ACCCTGCTTTGCACTCCTGACGTAACCTGCGGCTCGGAGCAGGTCTACTGGTCGTTGCCAGACATG
ATGCAAGACCCCGTGACCTTCCGCTCCACGCGCCAGATCAGCAACTTTCCGGTGGTGGGCGCCGA
GCTGTTGCCCGTGCACTCCAAGAGCTTCTACAACGACCAGGCCGTCTACTCCCAACTCATCCGCCA
GTTTACCTCTCTGACCCACGTGTTCAATCGCTTTCCCGAGAACCAGATTTTGGCGCGCCAGTCCTC
CGACAGACTGAGTCGCCCGGGTACCGCGATCGCCATGGCGGGACACCTGGCTTCGGATTTCGCCT
TCTCGCCCCCTCCAGGTGGTGGAGGTGATGGGCCAGGGGGGCCGGAGCCGGGCTGGGTTGATCC
TCGGACCTGGCTAAGCTTCCAAGGCCCTCCTGGAGGGCCAGGAATCGGGCCGGGGGTTGGGCCA
GGCTCTGAGGTGTGGGGGATTCCCCCATGCCCCCCGCCGTATGAGTTCTGTGGGGGGATGGCGTA
CTGTGGGCCCCAGGTTGGAGTGGGGCTAGTGCCCCAAGGCGGCTTGGAGACCTCTCAGCCTGAG
GGTGAAGCAGGAGTCGGGGTGGAGAGCAACTCCGATGGGGCCTCCCCGGAGCCCTGCACCGTCA
CCCCTGGTGCCGTGAAGCTGGAGAAGGAGAAGCTGGAGCAAAACCCGGAGGAGTCCCAGGACAT
CAAAGCTCTGCAGAAAGAACTCGAGCAATTTGCCAAGCTCCTGAAGCAGAAGAGGATCACCCTGGG
ATATACACAGGCCGATGTGGGGCTCACCCTGGGGGTTCTATTTGGGAAGGTATTCAGCCAAACGAC
CATCTGCCGCTTTGAGGCTCTGCAGCTTAGCTTCAAGAACATGTGTAAGCTGCGGCCCTTGCTGCA
GAAGTGGGTGGAGGAAGCTGACAACAATGAAAATCTTCAGGAGATATGCAAAGCAGAAACCCTCGT
GCAGGCCCGAAAGAGAAAGCGAACCAGTATCGAGAACCGAGTGAGAGGCAACCTGGAGAATTTGT
TCCTGCAGTGCCCGAAACCCACACTGCAGCAGATCAGCCACATCGCCCAGCAGCTTGGGCTCGAG
AAGGATGTGGTCCGAGTGTGGTTCTGTAACCGGCGCCAGAAGGGCAAGCAATCAAGCAGCGACTA

TGCACAACGAGAGGATTTTGAGGCTGCTGGGTCTCCTTTCTCAGGGGGACCAGTGTCCTTTCCTCT
GGCCCCAGGGCCCCATTTTGGTACCCCAGGCTATGGGAGCCCTCACTTCACTGCACTGTACTCCTC
GGTCCCTTTCCCTGAGGGGGAAGCCTTTCCCCCTGTCTCCGTCACCACTCTGGGCTCTCCCATGCA
TTCAAACGGGGGAGGAGGCTCCATGGTGAGCAAGGGCGAGGAGGATAACATGGCCATCATCAAGG
AGTTCATGCGCTTCAAGGTGCACATGGAGGGCTCCGTGAACGGCCACGAGTTCGAGAATCGAGGG
CGAGGGCGAGGGCCGCCCCTACGAGGGCACCCAGACCGCCAAGCTGAAGGTGACCAAGGGTGG
CCCCCTGCCCTTCGCCTGGGACATCCTGTCCCCTCAGTTCATGTACGGCTCCAAGGCCTACGTGAA
GCACCCCGCCGACATCCCCGACTACTTGAAGCTGTCCTTCCCCGAGGGCTTCAAGTGGGAGCGCG
TGATGAACTTCGAGGACGGCGGCGTGGTGACCGTGACCCAGGACTCCTCCCTGCAGGACGGCGA
GTTCATCTACAAGGTGAAGCTGCGCGGCACCAACTTCCCCTCCGACGGCCCCGTAATGCAGAAGAA
GACCATGGGCTGGGAGGCCTCCTCCGAGCGGATGTACCCCGAGGACGGCGCCCTGAAGGGCGAG
ATCAAGCAGAGGCTGAAGCTGAAGGACGGCGGCCACTACGACGCTGAGGTCAAGACCACCTACAA
GGCCAAGAAGCCCGTGCAGCTGCCCGGCGCCTACAACGTCAACATCAAGTTGGACATCACCTCCC
ACAACGAGGACTACACCATCGTGGAACAGTACGAACGCGCCGAGGGCCGCCACTCCACCGGCGG
CATGGACGAGCTGTACAAGTAACTCGAGTCTAGAGGGCCCGTTTAAACCCGCTGATCAGCCTCGAC
TGTGCCTTCTAGTTGCCAGCCATCTGTTGTTTGCCCCTCCCCCGTGCCTTCCTTGACCCTGGAAGG
TGCCACTCCCACTGTCCTTTCCTAATAAAATGAGGAAATTGCATCGCATTGTCTGAGTAGGTGTCATT
CTATTCTGGGGGGTGGGGTGGGGCAGGACAGCAAGGGGGAGGATTGGGAAGACAATAGCAGGCA
TGCTGGGGATGCGGTGGGCTCTATGGCTTCTGAGGCGGAAAGAACCAGCTGGGGCTCTAGGGGGT
ATCCCCACGCGCCCTGTAGCGGCGCATTAAGCGCGGCGGGTGTGGTGGTTACGCGCAGCGTGAC
CGCTACACTTGCCAGCGCCCTAGCGCCCGCTCCTTTCGCTTTCTTCCCTTCCTTTCTCGCCACGTT
CGCCGGCTTTCCCCGTCAAGCTCTAAATCGGGGGCTCCCTTTAGGGTTCCGATTTAGTGCTTTACG
GCACCTCGACCCCAAAAAACTTGATTAGGGTGATGGTTCACATCGAATTCCTGCAGCCCGGGGGAT
CCATTAACTAATAAAAAAAAATAATAAAGCATCACTTACTTAAAATCAGTTAGCAAATTTCTGTCCAGTT
TATTCAGCAGCACCTCCTTGCCCTCCTCCCAGCTCTGGTATTGCAGCTTCCTCCTGGCTGCAAACTT
TCTCCACAATCTAAATGGAATGTCAGTTTCCTCCTGTTCCTGTCCATCCGCACCCACTATCTTCATGT
TGTTGCAGATGAAGCGCGCAAGACCGTCTGAAGATACCTTCAACCCCGTGTATCCATATGACACGGA
AACCGGTCCTCCAACTGTGCCTTTTCTTACTCCTCCCTTTGTATCCCCCAATGGGTTTCAAGAGAGT
CCCCCTGGAGTTCTTACTTTAAAATGTTTAACCCCACTAACAACCACAGGCGGATCTCTACAGCTAAA
AGTGGGAGGGGGACTTACAGTGGATGACACTGATGGTACCTTACAAGAAAACATACGTGCTACAGC
ACCCATTACTAAAAATAATCACTCTGTAGAACTATCCATTGGAAATGGATTAGAAACTCAAAACAATAAA
CTATGTGCCAAATTGGGAAATGGGTTAAAATTTAACAACGGTGACATTTGTATAAAGGATAGTATTAAC
ACCTTATGGACTGGAATAAACCCTCCACCTAACTGTCAAATTGTGGAAAACACTAATACAAATGATGG
CAAACTTACTTTAGTATTAGTAAAAAACGGAGGGCTTGTTAATGGCTACGTGTCTCTAGTTGGTGTATC
AGACACTGTGAACCAAATGTTCACACAAAAGACAGCAAACATCCAATTAAGATTATATTTTGACTCTTC
TGGAAATCTATTAACTGATGAATCAGACTTAAAAATTCCACTTAAAAATAAATCTTCTACAGCGACCAG
TGAAACTGTAGCCAGCAGCAAAGCCTTTATGCCAAGTACTACAGCTTATCCCTTCAACACCACTACTA
GGGATAGTGAAAACTACATTCATGGAATATGTTACTACATGACTAGTTATGATAGAAGTCTATTTCCCTT
GAACATTTCTATAATGCTAAACAGCCGTATGATTTCTTCCAATGTTGCCTATGCCATACAATTTGAATG
GAATCTAAATGCAAGTGAATCTCCAGAAAGCAACATAGCTACGCTGACCACATCCCCCTTTTTCTTTT
CTTACATTACAGAAGACGACAACTAAAATGCCCAAGAATAAAGAATCGTTTGTGTTATGTTTCAACGT
GTTTATTTTTCAATTGCAGAAAATTTCAAGTCATTTTTCATTCAGTAGTATAGCCCCACCACCACATAG
CTTATACAGATCACCGTACCTTAATCAAACTCACAGAACCCTAGTATTCAACCTGCCACCTCCCTCCC
AACACACAGAGTACACAGTCCTTTCTCCCCGGCTGGCCTTAAAAAGCATCATATCATGGGTAACAGA

CATATTCTTAGGTGTTATATTCCACACGGTTTCCTGTCGAGCCAAACGCTCATCAGTGATATTAATAAA
CTCCCCGGGCAGCTCACTTAAGTTCATGTCGCTGTCCAGCTGCTGAGCCACAGGCTGCTGTCCAAC
TTGCGGTTGCTTAACGGGCGGCGAAGGAGAAGTCCACGCCTACATGGGGGTAGAGTCATAATCGTG
CATCAGGATAGGGCGGTGGTGCTGCAGCAGCGCGCGAATAAACTGCTGCCGCCGCCGCTCCGTCC
TGCAGGAATACAACATGGCAGTGGTCTCCTCAGCGATGATTCGCACCGCCCGCAGCATAAGGCGCC
TTGTCCTCCGGGCACAGCAGCGCACCCTGATCTCACTTAAATCAGCACAGTAACTGCAGCACAGCA
CCACAATATTGTTCAAAATCCCACAGTGCAAGGCGCTGTATCCAAAGCTCATGGCGGGGACCACAG
AACCCACGTGGCCATCATACCACAAGCGCAGGTAGATTAAGTGGCGACCCCTCATAAACACGCTGG
ACATAAACATTACCTCTTTTGGCATGTTGTAATTCACCACCTCCCGGTACCATATAAACCTCTGATTAA
ACATGGCGCCATCCACCACCATCCTAAACCAGCTGGCCAAAACCTGCCCGCCGGCTATACACTGCA
GGGAACCGGGACTGGAACAATGACAGTGGAGAGCCCAGGACTCGTAACCATGGATCATCATGCTC
GTCATGATATCAATGTTGGCACAACACAGGCACACGTGCATACACTTCCTCAGGATTACAAGCTCCT
CCCGCGTTAGAACCATATCCCAGGGAACAACCCATTCCTGAATCAGCGTAAATCCCACACTGCAGG
GAAGACCTCGCACGTAACTCACGTTGTGCATTGTCAAAGTGTTACATTCGGGCAGCAGCGGATGAT
CCTCCAGTATGGTAGCGCGGGTTTCTGTCTCAAAAGGAGGTAGACGATCCCTACTGTACGGAGTGC
GCCGAGACAACCGAGATCGTGTTGGTCGTAGTGTCATGCCAAATGGAACGCCGGACGTAGTCATAT
TTCCTGAAGCAAAACCAGGTGCGGGCGTGACAAACAGATCTGCGTCTCCGGTCTCGCCGCTTAGAT
CGCTCTGTGTAGTAGTTGTAGTATATCCACTCTCTCAAAGCATCCAGGCGCCCCCTGGCTTCGGGTT
CTATGTAAACTCCTTCATGCGCCGCTGCCCTGATAACATCCACCACCGCAGAATAAGCCACACCCAG
CCAACCTACACATTCGTTCTGCGAGTCACACACGGGAGGAGCGGGAAGAGCTGGAAGAACCATGT
TTTTTTTTTATTCCAAAAGATTATCCAAAACCTCAAAATGAAGATCTATTAAGTGAACGCGCTCCCCTC
CGGTGGCGTGGTCAAACTCTACAGCCAAAGAACAGATAATGGCATTTGTAAGATGTTGCACAATGGC
TTCCAAAAGGCAAACGGCCCTCACGTCCAAGTGGACGTAAAGGCTAAACCCTTCAGGGTGAATCTC
CTCTATAAACATTCCAGCACCTTCAACCATGCCCAAATAATTCTCATCTCGCCACCTTCTCAATATATC
TCTAAGCAAATCCCGAATATTAAGTCCGGCCATTGTAAAAATCTGCTCCAGAGCGCCCTCCACCTTCA
GCCTCAAGCAGCGAATCATGATTGCAAAAATTCAGGTTCCTCACAGACCTGTATAAGATTCAAAAGC
GGAACATTAACAAAAATACCGCGATCCCGTAGGTCCCTTCGCAGGGCCAGCTGAACATAATCGTGCA
GGTCTGCACGGACCAGCGCGGCCACTTCCCCGCCAGGAACCATGACAAAAGAACCCACACTGATT
ATGACACGCATACTCGGAGCTATGCTAACCAGCGTAGCCCCGATGTAAGCTTGTTGCATGGGCGGC
GATATAAAATGCAAGGTGCTGCTCAAAAAATCAGGCAAAGCCTCGCGCAAAAAAGAAAGCACATCGT
AGTCATGCTCATGCAGATAAAGGCAGGTAAGCTCCGGAACCACCACAGAAAAAGACACCATTTTTCT
CTCAAACATGTCTGCGGGTTTCTGCATAAACACAAAATAAAATAACAAAAAAACATTTAAACATTAGAA
GCCTGTCTTACAACAGGAAAAACAACCCTTATAAGCATAAGACGGACTACGGCCATGCCGGCGTGA
CCGTAAAAAAACTGGTCACCGTGATTAAAAAGCACCACCGACAGCTCCTCGGTCATGTCCGGAGTC
ATAATGTAAGACTCGGTAAACACATCAGGTTGATTCACATCGGTCAGTGCTAAAAAGCGACCGAAATA
GCCCGGGGGAATACATACCCGCAGGCGTAGAGACAACATTACAGCCCCCATAGGAGGTATAACAAA
ATTAATAGGAGAGAAAAACACATAAACACCTGAAAAACCCTCCTGCCTAGGCAAAATAGCACCCTCC
CGCTCCAGAACAACATACAGCGCTTCCACAGCGGCAGCCATAACAGTCAGCCTTACCAGTAAAAAA
GAAAACCTATTAAAAAAACACCACTCGACACGGCACCAGCTCAATCAGTCACAGTGTAAAAAAGGGC
CAAGTGCAGAGCGAGTATATATAGGACTAAAAAATGACGTAACGGTTAAAGTCCACAAAAAACACCCA
GAAAACCGCACGCGAACCTACGCCCAGAAACGAAAGCCAAAAAACCCACAACTTCCTCAAATCGTC
ACTTCCGTTTTCCCACGTTACGTCACTTCCCATTTTAAGAAAACTACAATTCCCAACACATACAAGTTA
CTCCGCCCTAAAACCTACGTCACCCGCCCCGTTCCCACGCCCCGCGCCACGTCACAAACTCCACC
CCCTCATTATCATATTGGCTTCAATCCAAAATAAGGTATATTATTGATGATGTTACATCGTTAATTAACGA

TTTCGAACCCGGGGTACCGAATTCCTCGAGTCTAGAGGAGCATGCGACGTCGCAATTCGCCCTATA
GTGAGTCGTATTACAATTCACTGGCCGTCGTTTTACAACGTCGTGACTGGGAAAACCCTGGCGTTAC
CCAACTTAATCGCCTTGCAGCACATCCCCCTTTCGCCAGCTGGCGTAATAGCGAAGAGGCCCGCAC
CGATCGCCCTTCCCAACAGTTGCGCAGCCTGAATGGCGAATGGAAATTGTAAGCGTTAATATTTTGTT
AAAATTCGCGTTAAATTTTTGTTAAATCAGCTCATTTTTTAACCAATAGGCCGAAATCGGCAAAATCCC
TTATAAATCAAAAGAATAGACCGAGATAGGGTTGAGTGTTGTTCCAGTTTGGAACAAGAGTCCACTAT
TAAAGAACGTGGACTCCAACGTCAAAGGGCGAAAAACCGTCTATCAGGGCGATGGCCCACTACGTG
AACCATCACCCTAATCAAGTTTTTTGGGGTCGAGGTGCCGTAAAGCACTAAATCGGAACCCTAAAGG
GAGCCCCCGATTTAGAGCTTGACGGGGAAAGCCGGCGAACGTGGCGAGAAAGGAAGGGAAGAAA
GCGAAAGGAGCGGGCGCTAGGGCGCTGGCAAGTGTAGCGGTCACGCTGCGCGTAACCACCACAC
CCGCCGCGCTTAATGCGCCGCTACAGGGCGCGTCCTGATGCGGTATTTTCTCCTTACGCATCTGTG
CGGTATTTCACACCGCATACAGGTGGCACTTTTCGGGGAAATGTGCGCGGAACCCCTATTTGTTTAT
TTTTCTAAATACATTCAAATATGTATCCGCTCATGAGACAATAACCCTGATAAATGCTTCAATAATATTGA
AAAAGGAAGAGTATGAGTATTCAACATTTCCGTGTCGCCCTTATTCCCTTTTTTGCGGCATTTTGCCT
TCCTGTTTTTGCTCACCCAGAAACGCTGGTGAAAGTAAAAGATGCTGAAGATCAGTTGGGTGCACG
AGTGGGTTACATCGAACTGGATCTCAACAGCGGTAAGATCCTTGAGAGTTTTCGCCCCGAAGAACG
TTTTCCAATGATGAGCACTTTTAAAGTTCTGCTATGTGGCGCGGTATTATCCCGTATTGACGCCGGGC
AAGAGCAACTCGGTCGCCGCATACACTATTCTCAGAATGACTTGGTTGAGTACTCACCAGTCACAGA
AAAGCATCTTACGGATGGCATGACAGTAAGAGAATTATGCAGTGCTGCCATAACCATGAGTGATAACA
CTGCGGCCAACTTACTTCTGACAACGATCGGAGGACCGAAGGAGCTAACCGCTTTTTTGCACAACA
TGGGGGATCATGTAACTCGCCTTGATCGTTGGGAACCGGAGCTGAATGAAGCCATACCAAACGACG
AGCGTGACACCACGATGCCTGTAGCAATGGCAACAACGTTGCGCAAACTATTAACTGGCGAACTACT
TACTCTAGCTTCCCGGCAACAATTAATAGACTGGATGGAGGCGGATAAAGTTGCAGGACCACTTCTG
CGCTCGGCCCTTCCGGCTGGCTGGTTTATTGCTGATAAATCTGGAGCCGGTGAGCGTGGGTCTCG
CGGTATCATTGCAGCACTGGGGCCAGATGGTAAGCCCTCCCGTATCGTAGTTATCTACACGACGGG
GAGTCAGGCAACTATGGATGAACGAAATAGACAGATCGCTGAGATAGGTGCCTCACTGATTAAGCAT
TGGTAACTGTCAGACCAAGTTTACTCATATATACTTTAGATTGATTTAAAACTTCATTTTTAATTTAAAAG
GATCTAGGTGAAGATCCTTTTTGATAATCTCATGACCAAAATCCCTTAACGTGAGTTTTCGTTCCACT
GAGCGTCAGACCCCGTAGAAAAGATCAAAGGATCTTCTTGAGATCCTTTTTTTCTGCGCGTAATCTG
CTGCTTGCAAACAAAAAAACCACCGCTACCAGCGGTGGTTTGTTTGCCGGATCAAGAGCTACCAAC
TCTTTTTCCGAAGGTAACTGGCTTCAGCAGAGCGCAGATACCAAATACTGTTCTTCTAGTGTAGCCGT
AGTTAGGCCACCACTTCAAGAACTCTGTAGCACCGCCTACATACCTCGCTCTGCTAATCCTGTTACC
AGTGGCTGCTGCCAGTGGCGATAAGTCGTGTCTTACCGGGTTGGACTCAAGACGATAGTTACCGGA
TAAGGCGCAGCGGTCGGGCTGAAC

P5：

>P141 pAdHM4-SFFV-M2K

TAGAATACTCAAGCTAGTTAATTAACGTTAATTAACATCATCAATAATATACCTTATTTTGGATTGAAGCC
AATATGATAATGAGGGGGTGGAGTTTGTGACGTGGCGCGGGGCGTGGGAACGGGGCGGGTGACG
TAGTAGTGTGGCGGAAGTGTGATGTTGCAAGTGTGGCGGAACACATGTAAGCGACGGATGTGGCAA
AAGTGACGTTTTTGGTGTGCGCCGGTGTACACAGGAAGTGACAATTTTCGCGCGGTTTTAGGCGGA
TGTTGTAGTAAATTTGGGCGTAACCGAGTAAGATTTGGCCATTTTCGCGGGAAAACTGAATAAGAGG
AAGTGAAATCTGAATAATTTTGTGTTACTCATAGCGCGTAATCTCTAGCATCGTAACTATAACGGTCCT
AAGGTAGCGAGAATTATTTAAATATCTATGTCGGGTGCGGAGAAAGAGGTAATGAAATGGCATCGACT
CGAAGATCTGGGCGTGGTTAAGGGTGGGAAAGAATATATAAGGTGGGGGTCTTATGTAGTTTTGTAT
CTGTTTTGCAGCAGCCGCCGCCGCCATGAGCACCAACTCGTTTGATGGAAGCATTGTGAGCTCATA
TTTGACAACGCGCATGCCCCCATGGGCCGGGGTGCGTCAGAATGTGATGGGCTCCAGCATTGATG
GTCGCCCCGTCCTGCCCGCAAACTCTACTACCTTGACCTACGAGACCGTGTCTGGAACGCCGTTG
GAGACTGCAGCCTCCGCCGCCGCTTCAGCCGCTGCAGCCACCGCCCGCGGGATTGTGACTGACT
TTGCTTTCCTGAGCCCGCTTGCAAGCAGTGCAGCTTCCCGTTCATCCGCCCGCGATGACAAGTTGA
CGGCTCTTTTGGCACAATTGGATTCTTTGACCCGGGAACTTAATGTCGTTTCTCAGCAGCTGTTGGA
TCTGCGCCAGCAGGTTTCTGCCCTGAAGGCTTCCTCCCCTCCCAATGCGGTTTAAAACATAAATAAA
AAACCAGACTCTGTTTGGATTTGGATCAAGCAAGTGTCTTGCTGTCTTTATTTAGGGGTTTTGCGCG
CGCGGTAGGCCCGGGACCAGCGGTCTCGGTCGTTGAGGGTCCTGTGTATTTTTTCCAGGACGTGG
TAAAGGTGACTCTGGATGTTCAGATACATGGGCATAAGCCCGTCTCTGGGGTGGAGGTAGCACCAC
TGCAGAGCTTCATGCTGCGGGGTGGTGTTGTAGATGATCCAGTCGTAGCAGGAGCGCTGGGCGTG
GTGCCTAAAAATGTCTTTCAGTAGCAAGCTGATTGCCAGGGGCAGGCCCTTGGTGTAAGTGTTTACA
AAGCGGTTAAGCTGGGATGGGTGCATACGTGGGGATATGAGATGCATCTTGGACTGTATTTTTAGGT
TGGCTATGTTCCCAGCCATATCCCTCCGGGGATTCATGTTGTGCAGAACCACCAGCACAGTGTATCC
GGTGCACTTGGGAAATTTGTCATGTAGCTTAGAAGGAAATGCGTGGAAGAACTTGGAGACGCCCTT
GTGACCTCCAAGATTTTCCATGCATTCGTCCATAATGATGGCAATGGGCCCACGGGCGGCGGCCTG
GGCGAAGATATTTCTGGGATCACTAACGTCATAGTTGTGTTCCAGGATGAGATCGTCATAGGCCATTT
TTACAAAGCGCGGGCGGAGGGTGCCAGACTGCGGTATAATGGTTCCATCCGGCCCAGGGGCGTAG
TTACCCTCACAGATTTGCATTTCCCACGCTTTGAGTTCAGATGGGGGGATCATGTCTACCTGCGGGG
CGATGAAGAAAACGGTTTCCGGGGTAGGGGAGATCAGCTGGGAAGAAAGCAGGTTCCTGAGCAGC
TGCGACTTACCGCAGCCGGTGGGCCCGTAAATCACACCTATTACCGGCTGCAACTGGTAGTTAAGA
GAGCTGCAGCTGCCGTCATCCCTGAGCAGGGGGGCCACTTCGTTAAGCATGTCCCTGACTCGCAT
GTTTTCCCTGACCAAATCCGCCAGAAGGCGCTCGCCGCCCAGCGATAGCAGTTCTTGCAAGGAAG
CAAAGTTTTTCAACGGTTTGAGACCGTCCGCCGTAGGCATGCTTTTGAGCGTTTGACCAAGCAGTT
CCAGGCGGTCCCACAGCTCGGTCACCTGCTCTACGGCATCTCGATCCAGCATATCTCCTCGTTTCG
CGGGTTGGGGCGGCTTTCGCTGTACGGCAGTAGTCGGTGCTCGTCCAGACGGGCCAGGGTCATG
TCTTTCCACGGGCGCAGGGTCCTCGTCAGCGTAGTCTGGGTCACGGTGAAGGGGTGCGCTCCGG
GCTGCGCGCTGGCCAGGGTGCGCTTGAGGCTGGTCCTGCTGGTGCTGAAGCGCTGCCGGTCTTC
GCCCTGCGCGTCGGCCAGGTAGCATTTGACCATGGTGTCATAGTCCAGCCCCTCCGCGGCGTGGC
CCTTGGCGCGCAGCTTGCCCTTGGAGGAGGCGCCGCACGAGGGGCAGTGCAGACTTTTGAGGGC
GTAGAGCTTGGGCGCGAGAAATACCGATTCCGGGGAGTAGGCATCCGCGCCGCAGGCCCCGCAG
ACGGTCTCGCATTCCACGAGCCAGGTGAGCTCTGGCCGTTCGGGGTCAAAAACCAGGTTTCCCCC
ATGCTTTTTGATGCGTTTCTTACCTCTGGTTTCCATGAGCCGGTGTCCACGCTCGGTGACGAAAAGG
CTGTCCGTGTCCCCGTATACAGACTTGAGAGGCCTGTCCTCGAGCGGTGTTCCGCGGTCCTCCTC

GTATAGAAACTCGGACCACTCTGAGACAAAGGCTCGCGTCCAGGCCAGCACGAAGGAGGCTAAGT
GGGAGGGGTAGCGGTCGTTGTCCACTAGGGGGTCCACTCGCTCCAGGGTGTGAAGACACATGTCG
CCCTCTTCGGCATCAAGGAAGGTGATTGGTTTGTAGGTGTAGGCCACGTGACCGGGTGTTCCTGAA
GGGGGGCTATAAAAGGGGGTGGGGGCGCGTTCGTCCTCACTCTCTTCCGCATCGCTGTCTGCGAG
GGCCAGCTGTTGGGGTGAGTACTCCCTCTGAAAAGCGGGCATGACTTCTGCGCTAAGATTGTCAGT
TTCCAAAAACGAGGAGGATTTGATATTCACCTGGCCCGCGGTGATGCCTTTGAGGGTGGCCGCATC
CATCTGGTCAGAAAAGACAATCTTTTTGTTGTCAAGCTTGGTGGCAAACGACCCGTAGAGGGCGTT
GGACAGCAACTTGGCGATGGAGCGCAGGGTTTGGTTTTTGTCGCGATCGGCGCGCTCCTTGGCCG
CGATGTTTAGCTGCACGTATTCGCGCGCAACGCACCGCCATTCGGGAAAGACGGTGGTGCGCTCG
TCGGGCACCAGGTGCACGCGCCAACCGCGGTTGTGCAGGGTGACAAGGTCAACGCTGGTGGCTA
CCTCTCCGCGTAGGCGCTCGTTGGTCCAGCAGAGGCGGCCGCCCTTGCGCGAGCAGAATGGCGG
TAGGGGGTCTAGCTGCGTCTCGTCCGGGGGGTCTGCGTCCACGGTAAAGACCCCGGGCAGCAGG
CGCGCGTCGAAGTAGTCTATCTTGCATCCTTGCAAGTCTAGCGCCTGCTGCCATGCGCGGGCGGCA
AGCGCGCGCTCGTATGGGTTGAGTGGGGGACCCCATGGCATGGGGTGGGTGAGCGCGGAGGCGT
ACATGCCGCAAATGTCGTAAACGTAGAGGGGCTCTCTGAGTATTCCAAGATATGTAGGGTAGCATCT
TCCACCGCGGATGCTGGCGCGCACGTAATCGTATAGTTCGTGCGAGGGAGCGAGGAGGTCGGGAC
CGAGGTTGCTACGGGCGGGCTGCTCTGCTCGGAAGACTATCTGCCTGAAGATGGCATGTGAGTTG
GATGATATGGTTGGACGCTGGAAGACGTTGAAGCTGGCGTCTGTGAGACCTACCGCGTCACGCAC
GAAGGAGGCGTAGGAGTCGCGCAGCTTGTTGACCAGCTCGGCGGTGACCTGCACGTCTAGGGCG
CAGTAGTCCAGGGTTTCCTTGATGATGTCATACTTATCCTGTCCCTTTTTTTTCCACAGCTCGCGGTT
GAGGACAAACTCTTCGCGGTCTTTCCAGTACTCTTGGATCGGAAACCCGTCGGCCTCCGAACGGTA
AGAGCCTAGCATGTAGAACTGGTTGACGGCCTGGTAGGCGCAGCATCCCTTTTCTACGGGTAGCGC
GTATGCCTGCGCGGCCTTCCGGAGCGAGGTGTGGGTGAGCGCAAAGGTGTCCCTGACCATGACTT
TGAGGTACTGGTATTTGAAGTCAGTGTCGTCGCATCCGCCCTGCTCCCAGAGCAAAAAGTCCGTGC
GCTTTTTGGAACGCGGATTTGGCAGGGCGAAGGTGACATCGTTGAAGAGTATCTTTCCCGCGCGAG
GCATAAAGTTGCGTGTGATGCGGAAGGGTCCCGGCACCTCGGAACGGTTGTTAATTACCTGGGCG
GCGAGCACGATCTCGTCAAAGCCGTTGATGTTGTGGCCCACAATGTAAAGTTCCAAGAAGCGCGGG
ATGCCCTTGATGGAAGGCAATTTTTTAAGTTCCTCGTAGGTGAGCTCTTCAGGGGAGCTGAGCCCG
TGCTCTGAAAGGGCCCAGTCTGCAAGATGAGGGTTGGAAGCGACGAATGAGCTCCACAGGTCACG
GGCCATTAGCATTTGCAGGTGGTCGCGAAAGGTCCTAAACTGGCGACCTATGGCCATTTTTTCTGG
GGTGATGCAGTAGAAGGTAAGCGGGTCTTGTTCCCAGCGGTCCCATCCAAGGTTCGCGGCTAGGT
CTCGCGCGGCAGTCACTAGAGGCTCATCTCCGCCGAACTTCATGACCAGCATGAAGGGCACGAGC
TGCTTCCCAAAGGCCCCCATCCAAGTATAGGTCTCTACATCGTAGGTGACAAAGAGACGCTCGGTG
CGAGGATGCGAGCCGATCGGGAAGAACTGGATCTCCCGCCACCAATTGGAGGAGTGGCTATTGAT
GTGGTGAAAGTAGAAGTCCCTGCGACGGGCCGAACACTCGTGCTGGCTTTTGTAAAAACGTGCGC
AGTACTGGCAGCGGTGCACGGGCTGTACATCCTGCACGAGGTTGACCTGACGACCGCGCACAAGG
AAGCAGAGTGGGAATTTGAGCCCCTCGCCTGGCGGGTTTGGCTGGTGGTCTTCTACTTCGGCTGC
TTGTCCTTGACCGTCTGGCTGCTCGAGGGGAGTTACGGTGGATCGGACCACCACGCCGCGCGAGC
CCAAAGTCCAGATGTCCGCGCGCGGCGGTCGGAGCTTGATGACAACATCGCGCAGATGGGAGCTG
TCCATGGTCTGGAGCTCCCGCGGCGTCAGGTCAGGCGGGAGCTCCTGCAGGTTTACCTCGCATAG
ACGGGTCAGGGCGCGGGCTAGATCCAGGTGATACCTAATTTCCAGGGGCTGGTTGGTGGCGGCGT
CGATGGCTTGCAAGAGGCCGCATCCCCGCGGCGCGACTACGGTACCGCGCGGCGGGCGGTGGG
CCGCGGGGGTGTCCTTGGATGATGCATCTAAAAGCGGTGACGCGGGCGAGCCCCCGGAGGTAGG
GGGGGCTCCGGACCCGCCGGGAGAGGGGGCAGGGGCACGTCGGCGCCGCGCGCGGGCAGGAG

CTGGTGCTGCGCGCGTAGGTTGCTGGCGAACGCGACGACGCGGCGGTTGATCTCCTGAATCTGGC
GCCTCTGCGTGAAGACGACGGGCCCGGTGAGCTTGAACCTGAAAGAGAGTTCGACAGAATCAATT
TCGGTGTCGTTGACGGCGGCCTGGCGCAAAATCTCCTGCACGTCTCCTGAGTTGTCTTGATAGGCG
ATCTCGGCCATGAACTGCTCGATCTCTTCCTCCTGGAGATCTCCGCGTCCGGCTCGCTCCACGGTG
GCGGCGAGGTCGTTGGAAATGCGGGCCATGAGCTGCGAGAAGGCGTTGAGGCCTCCCTCGTTCC
AGACGCGGCTGTAGACCACGCCCCCTTCGGCATCGCGGGCGCGCATGACCACCTGCGCGAGATT
GAGCTCCACGTGCCGGGCGAAGACGGCGTAGTTTCGCAGGCGCTGAAAGAGGTAGTTGAGGGTG
GTGGCGGTGTGTTCTGCCACGAAGAAGTACATAACCCAGCGTCGCAACGTGGATTCGTTGATATCC
CCCAAGGCCTCAAGGCGCTCCATGGCCTCGTAGAAGTCCACGGCGAAGTTGAAAAACTGGGAGTT
GCGCGCCGACACGGTTAACTCCTCCTCCAGAAGACGGATGAGCTCGGCGACAGTGTCGCGCACCT
CGCGCTCAAAGGCTACAGGGGCCTCTTCTTCTTCTTCAATCTCCTCTTCCATAAGGGCCTCCCCTTC
TTCTTCTTCTGGCGGCGGTGGGGGAGGGGGGACACGGCGGCGACGACGGCGCACCGGGAGGCG
GTCGACAAAGCGCTCGATCATCTCCCCGCGGCGACGGCGCATGGTCTCGGTGACGGCGCGGCCG
TTCTCGCGGGGGCGCAGTTGGAAGACGCCGCCCGTCATGTCCCGGTTATGGGTTGGCGGGGGGC
TGCCATGCGGCAGGGATACGGCGCTAACGATGCATCTCAACAATTGTTGTGTAGGTACTCCGCCGC
CGAGGGACCTGAGCGAGTCCGCATCGACCGGATCGGAAAACCTCTCGAGAAAGGCGTCTAACCAG
TCACAGTCGCAAGGTAGGCTGAGCACCGTGGCGGGCGGCAGCGGGCGGCGGTCGGGGTTGTTTC
TGGCGGAGGTGCTGCTGATGATGTAATTAAAGTAGGCGGTCTTGAGACGGCGGATGGTCGACAGAA
GCACCATGTCCTTGGGTCCGGCCTGCTGAATGCGCAGGCGGTCGGCCATGCCCCAGGCTTCGTTT
TGACATCGGCGCAGGTCTTTGTAGTAGTCTTGCATGAGCCTTTCTACCGGCACTTCTTCTTCTCCTT
CCTCTTGTCCTGCATCTCTTGCATCTATCGCTGCGGCGGCGGCGGAGTTTGGCCGTAGGTGGCGC
CCTCTTCCTCCCATGCGTGTGACCCCGAAGCCCCTCATCGGCTGAAGCAGGGCTAGGTCGGCGAC
AACGCGCTCGGCTAATATGGCCTGCTGCACCTGCGTGAGGGTAGACTGGAAGTCATCCATGTCCAC
AAAGCGGTGGTATGCGCCCGTGTTGATGGTGTAAGTGCAGTTGGCCATAACGGACCAGTTAACGGT
CTGGTGACCCGGCTGCGAGAGCTCGGTGTACCTGAGACGCGAGTAAGCCCTCGAGTCAAATACGT
AGTCGTTGCAAGTCCGCACCAGGTACTGGTATCCCACCAAAAAGTGCGGCGGCGGCTGGCGGTAG
AGGGGCCAGCGTAGGGTGGCCGGGGCTCCGGGGGCGAGATCTTCCAACATAAGGCGATGATATCC
GTAGATGTACCTGGACATCCAGGTGATGCCGGCGGCGGTGGTGGAGGCGCGCGGAAAGTCGCGG
ACGCGGTTCCAGATGTTGCGCAGCGGCAAAAAGTGCTCCATGGTCGGGACGCTCTGGCCGGTCAG
GCGCGCGCAATCGTTGACGCTCTAGCGTGCAAAAGGAGAGCCTGTAAGCGGGCACTCTTCCGTGG
TCTGGTGGATAAATTCGCAAGGGTATCATGGCGGACGACCGGGGTTCGAGCCCCGTATCCGGCCGT
CCGCCGTGATCCATGCGGTTACCGCCCGCGTGTCGAACCCAGGTGTGCGACGTCAGACAACGGG
GGAGTGCTCCTTTTGGCTTCCTTCCAGGCGCGGCGGCTGCTGCGCTAGCTTTTTTGGCCACTGGC
CGCGCGCAGCGTAAGCGGTTAGGCTGGAAAGCGAAAGCATTAAGTGGCTCGCTCCCTGTAGCCGG
AGGGTTATTTTCCAAGGGTTGAGTCGCGGGACCCCCGGTTCGAGTCTCGGACCGGCCGGACTGCG
GCGAACGGGGGTTTGCCTCCCCGTCATGCAAGACCCCGCTTGCAAATTCCTCCGGAAACAGGGAC
GAGCCCCTTTTTTGCTTTTCCCAGATGCATCCGGTGCTGCGGCAGATGCGCCCCCCTCCTCAGCAG
CGGCAAGAGCAAGAGCAGCGGCAGACATGCAGGGCACCCTCCCCTCCTCCTACCGCGTCAGGAG
GGGCGACATCCGCGGTTGACGCGGCAGCAGATGGTGATTACGAACCCCCGCGGCGCCGGGCCCG
GCACTACCTGGACTTGGAGGAGGGCGAGGGCCTGGCGCGGCTAGGAGCGCCCTCTCCTGAGCGG
CACCCAAGGGTGCAGCTGAAGCGTGATACGCGTGAGGCGTACGTGCCGCGGCAGAACCTGTTTCG
CGACCGCGAGGGAGAGGAGCCCGAGGAGATGCGGGATCGAAAGTTCCACGCAGGGCGCGAGCT
GCGGCATGGCCTGAATCGCGAGCGGTTGCTGCGCGAGGAGGACTTTGAGCCCGACGCGCGAACC
GGGATTAGTCCCGCGCGCGCACACGTGGCGGCCGCCGACCTGGTAACCGCATACGAGCAGACGG

TGAACCAGGAGATTAACTTTCAAAAAAGCTTTAACAACCACGTGCGTACGCTTGTGGCGCGCGAGG
AGGTGGCTATAGGACTGATGCATCTGTGGGACTTTGTAAGCGCGCTGGAGCAAAACCCAAATAGCA
AGCCGCTCATGGCGCAGCTGTTCCTTATAGTGCAGCACAGCAGGGACAACGAGGCATTCAGGGAT
GCGCTGCTAAACATAGTAGAGCCCGAGGGCCGCTGGCTGCTCGATTTGATAAACATCCTGCAGAGC
ATAGTGGTGCAGGAGCGCAGCTTGAGCCTGGCTGACAAGGTGGCCGCCATCAACTATTCCATGCTT
AGCCTGGGCAAGTTTTACGCCCGCAAGATATACCATACCCCTTACGTTCCCATAGACAAGGAGGTAA
AGATCGAGGGGTTCTACATGCGCATGGCGCTGAAGGTGCTTACCTTGAGCGACGACCTGGGCGTTT
ATCGCAACGAGCGCATCCACAAGGCCGTGAGCGTGAGCCGGCGGCGCGAGCTCAGCGACCGCGA
GCTGATGCACAGCCTGCAAAGGGCCCTGGCTGGCACGGGCAGCGGCGATAGAGAGGCCGAGTCC
TACTTTGACGCGGGCGCTGACCTGCGCTGGGCCCCAAGCCGACGCGCCCTGGAGGCAGCTGGGG
CCGGACCTGGGCTGGCGGTGGCACCCGCGCGCGCTGGCAACGTCGGCGGCGTGGAGGAATATG
ACGAGGACGATGAGTACGAGCCAGAGGACGGCGAGTACTAAGCGGTGATGTTTCTGATCAGATGAT
GCAAGACGCAACGGACCCGGCGGTGCGGGCGGCGCTGCAGAGCCAGCCGTCCGGCCTTAACTC
CACGGACGACTGGCGCCAGGTCATGGACCGCATCATGTCGCTGACTGCGCGCAATCCTGACGCGT
TCCGGCAGCAGCCGCAGGCCAACCGGCTCTCCGCAATTCTGGAAGCGGTGGTCCCGGCGCGCGC
AAACCCCACGCACGAGAAGGTGCTGGCGATCGTAAACGCGCTGGCCGAAAACAGGGCCATCCGG
CCCGACGAGGCCGGCCTGGTCTACGACGCGCTGCTTCAGCGCGTGGCTCGTTACAACAGCGGCA
ACGTGCAGACCAACCTGGACCGGCTGGTGGGGGATGTGCGCGAGGCCGTGGCGCAGCGTGAGC
GCGCGCAGCAGCAGGGCAACCTGGGCTCCATGGTTGCACTAAACGCCTTCCTGAGTACACAGCCC
GCCAACGTGCCGCGGGGACAGGAGGACTACACCAACTTTGTGAGCGCACTGCGGCTAATGGTGAC
TGAGACACCGCAAAGTGAGGTGTACCAGTCTGGGCCAGACTATTTTTTCCAGACCAGTAGACAAGG
CCTGCAGACCGTAAACCTGAGCCAGGCTTTCAAAAACTTGCAGGGGCTGTGGGGGGTGCGGGCTC
CCACAGGCGACCGCGCGACCGTGTCTAGCTTGCTGACGCCCAACTCGCGCCTGTTGCTGCTGCTA
ATAGCGCCCTTCACGGACAGTGGCAGCGTGTCCCGGGACACATACCTAGGTCACTTGCTGACACTG
TACCGCGAGGCCATAGGTCAGGCGCATGTGGACGAGCATACTTTCCAGGAGATTACAAGTGTCAGC
CGCGCGCTGGGGCAGGAGGACACGGGCAGCCTGGAGGCAACCCTAAACTACCTGCTGACCAACC
GGCGGCAGAAGATCCCCTCGTTGCACAGTTTAAACAGCGAGGAGGAGCGCATTTTGCGCTACGTG
CAGCAGAGCGTGAGCCTTAACCTGATGCGCGACGGGGTAACGCCCAGCGTGGCGCTGGACATGA
CCGCGCGCAACATGGAACCGGGCATGTATGCCTCAAACCGGCCGTTTATCAACCGCCTAATGGACT
ACTTGCATCGCGCGGCCGCCGTGAACCCCGAGTATTTCACCAATGCCATCTTGAACCCGCACTGGC
TACCGCCCCCTGGTTTCTACACCGGGGGATTCGAGGTGCCCGAGGGTAACGATGGATTCCTCTGG
GACGACATAGACGACAGCGTGTTTTCCCCGCAACCGCAGACCCTGCTAGAGTTGCAACAGCGCGA
GCAGGCAGAGGCGGCGCTGCGAAAGGAAAGCTTCCGCAGGCCAAGCAGCTTGTCCGATCTAGGC
GCTGCGGCCCCGCGGTCAGATGCTAGTAGCCCATTTCCAAGCTTGATAGGGTCTCTTACCAGCACT
CGCACCACCCGCCCGCGCCTGCTGGGCGAGGAGGAGTACCTAAACAACTCGCTGCTGCAGCCGC
AGCGCGAAAAAAACCTGCCTCCGGCATTTCCCAACAACGGGATAGAGAGCCTAGTGGACAAGATGA
GTAGATGGAAGACGTACGCGCAGGAGCACAGGGACGTGCCAGGCCCGCGCCCGCCCACCCGTCG
TCAAAGGCACGACCGTCAGCGGGGTCTGGTGTGGGAGGACGATGACTCGGCAGACGACAGCAGC
GTCCTGGATTTGGGAGGGAGTGGCAACCCGTTTGCGCACCTTCGCCCCAGGCTGGGGAGAATGTT
TTAAAAAAAAAAAAGCATGATGCAAAATAAAAAACTCACCAAGGCCATGGCACCGAGCGTTGGTTTT
CTTGTATTCCCCTTAGTATGCGGCGCGCGGCGATGTATGAGGAAGGTCCTCCTCCCTCCTACGAGA
GTGTGGTGAGCGCGGCGCCAGTGGCGGCGGCGCTGGGTTCTCCCTTCGATGCTCCCCTGGACCC
GCCGTTTGTGCCTCCGCGGTACCTGCGGCCTACCGGGGGGAGAAACAGCATCCGTTACTCTGAGT
TGGCACCCCTATTCGACACCACCCGTGTGTACCTGGTGGACAACAAGTCAACGGATGTGGCATCCC

TGAACTACCAGAACGACCACAGCAACTTTCTGACCACGGTCATTCAAAACAATGACTACAGCCCGG
GGGAGGCAAGCACACAGACCATCAATCTTGACGACCGGTCGCACTGGGGCGGCGACCTGAAAACC
ATCCTGCATACCAACATGCCAAATGTGAACGAGTTCATGTTTACCAATAAGTTTAAGGCGCGGGTGAT
GGTGTCGCGCTTGCCTACTAAGGACAATCAGGTGGAGCTGAAATACGAGTGGGTGGAGTTCACGCT
GCCCGAGGGCAACTACTCCGAGACCATGACCATAGACCTTATGAACAACGCGATCGTGGAGCACTA
CTTGAAAGTGGGCAGACAGAACGGGGTTCTGGAAAGCGACATCGGGGTAAAGTTTGACACCCGCA
ACTTCAGACTGGGGTTTGACCCCGTCACTGGTCTTGTCATGCCTGGGGTATATACAAACGAAGCCTT
CCATCCAGACATCATTTTGCTGCCAGGATGCGGGGTGGACTTCACCCACAGCCGCCTGAGCAACTT
GTTGGGCATCCGCAAGCGGCAACCCTTCCAGGAGGGCTTTAGGATCACCTACGATGATCTGGAGG
GTGGTAACATTCCCGCACTGTTGGATGTGGACGCCTACCAGGCGAGCTTGAAAGATGACACCGAAC
AGGGCGGGGGTGGCGCAGGCGGCAGCAACAGCAGTGGCAGCGGCGCGGAAGAGAACTCCAACG
CGGCAGCCGCGGCAATGCAGCCGGTGGAGGACATGAACGATCATGCCATTCGCGGCGACACCTTT
GCCACACGGGCTGAGGAGAAGCGCGCTGAGGCCGAAGCAGCGGCCGAAGCTGCCGCCCCCGCT
GCGCAACCCGAGGTCGAGAAGCCTCAGAAGAAACCGGTGATCAAACCCCTGACAGAGGACAGCAA
GAAACGCAGTTACAACCTAATAAGCAATGACAGCACCTTCACCCAGTACCGCAGCTGGTACCTTGCA
TACAACTACGGCGACCCTCAGACCGGAATCCGCTCATGGACCCTGCTTTGCACTCCTGACGTAACC
TGCGGCTCGGAGCAGGTCTACTGGTCGTTGCCAGACATGATGCAAGACCCCGTGACCTTCCGCTC
CACGCGCCAGATCAGCAACTTTCCGGTGGTGGGCGCCGAGCTGTTGCCCGTGCACTCCAAGAGCT
TCTACAACGACCAGGCCGTCTACTCCCAACTCATCCGCCAGTTTACCTCTCTGACCCACGTGTTCAA
TCGCTTTCCCGAGAACCAGATTTTGGCGCGCCCGCCAGCCCCCACCATCACCACCGTCAGTGAAA
ACGTTCCTGCTCTCACAGATCACGGGACGCTACCGCTGCGCAACAGCATCGGAGGAGTCCAGCGA
GTGACCATTACTGACGCCAGACGCCGCACCTGCCCCTACGTTTACAAGGCCCTGGGCATAGTCTCG
CCGCGCGTCCTATCGAGCCGCACTTTTTGAGCAAGCATGTCCATCCTTATATCGCCCAGCAATAACA
CAGGCTGGGGCCTGCGCTTCCCAAGCAAGATGTTTGGCGGGGCCAAGAAGCGCTCCGACCAACA
CCCAGTGCGCGTGCGCGGGCACTACCGCGCGCCCTGGGGCGCGCACAAACGCGGCCGCACTGG
GCGCACCACCGTCGATGACGCCATCGACGCGGTGGTGGAGGAGGCGCGCAACTACACGCCCACG
CCGCCACCAGTGTCCACAGTGGACGCGGCCATTCAGACCGTGGTGCGCGGAGCCCGGCGCTATG
CTAAAATGAAGAGACGGCGGAGGCGCGTAGCACGTCGCCACCGCCGCCGACCCGGCACTGCCGC
CCAACGCGCGGCGGCGGCCCTGCTTAACCGCGCACGTCGCACCGGCCGACGGGCGGCCATGCG
GGCCGCTCGAAGGCTGGCCGCGGGTATTGTCACTGTGCCCCCCAGGTCCAGGCGACGAGCGGCC
GCCGCAGCAGCCGCGGCCATTAGTGCTATGACTCAGGGTCGCAGGGGCAACGTGTATTGGGTGCG
CGACTCGGTTAGCGGCCTGCGCGTGCCCGTGCGCACCCGCCCCCCGCGCAACTAGATTGCAAGA
AAAAACTACTTAGACTCGTACTGTTGTATGTATCCAGCGGCGGCGGCGCGCAACGAAGCTATGTCCA
AGCGCAAAATCAAAGAAGAGATGCTCCAGGTCATCGCGCCGGAGATCTATGGCCCCCCGAAGAAG
GAAGAGCAGGATTACAAGCCCCGAAAGCTAAAGCGGGTCAAAAAGAAAAAGAAAGATGATGATGAT
GAACTTGACGACGAGGTGGAACTGCTGCACGCTACCGCGCCCAGGCGACGGGTACAGTGGAAAG
GTCGACGCGTAAAACGTGTTTTGCGACCCGGCACCACCGTAGTCTTTACGCCCGGTGAGCGCTCC
ACCCGCACCTACAAGCGCGTGTATGATGAGGTGTACGGCGACGAGGACCTGCTTGAGCAGGCCAA
CGAGCGCCTCGGGGAGTTTGCCTACGGAAAGCGGCATAAGGACATGCTGGCGTTGCCGCTGGAC
GAGGGCAACCCAACACCTAGCCTAAAGCCCGTAACACTGCAGCAGGTGCTGCCCGCGCTTGCACC
GTCCGAAGAAAAGCGCGGCCTAAAGCGCGAGTCTGGTGACTTGGCACCCACCGTGCAGCTGATGG
TACCCAAGCGCCAGCGACTGGAAGATGTCTTGGAAAAAATGACCGTGGAACCTGGGCTGGAGCCC
GAGGTCCGCGTGCGGCCAATCAAGCAGGTGGCGCCGGGACTGGGCGTGCAGACCGTGGACGTTC
AGATACCCACTACCAGTAGCACCAGTATTGCCACCGCCACAGAGGGCATGGAGACACAAACGTCCC

CGGTTGCCTCAGCGGTGGCGGATGCCGCGGTGCAGGCGGTCGCTGCGGCCGCGTCCAAGACCT
CTACGGAGGTGCAAACGGACCCGTGGATGTTTCGCGTTTCAGCCCCCCGGCGCCCGCGCCGTTC
GAGGAAGTACGGCGCCGCCAGCGCGCTACTGCCCGAATATGCCCTACATCCTTCCATTGCGCCTAC
CCCCGGCTATCGTGGCTACACCTACCGCCCCAGAAGACGAGCAACTACCCGACGCCGAACCACCA
CTGGAACCCGCCGCCGCCGTCGCCGTCGCCAGCCCGTGCTGGCCCCGATTTCCGTGCGCAGGGT
GGCTCGCGAAGGAGGCAGGACCCTGGTGCTGCCAACAGCGCGCTACCACCCCAGCATCGTTTAAA
AGCCGGTCTTTGTGGTTCTTGCAGATATGGCCCTCACCTGCCGCCTCCGTTTCCCGGTGCCGGGAT
TCCGAGGAAGAATGCACCGTAGGAGGGGCATGGCCGGCCACGGCCTGACGGGCGGCATGCGTCG
TGCGCACCACCGGCGGCGGCGCGCGTCGCACCGTCGCATGCGCGGCGGTATCCTGCCCCTCCTT
ATTCCACTGATCGCCGCGGCGATTGGCGCCGTGCCCGGAATTGCATCCGTGGCCTTGCAGGCGCA
GAGACACTGATTAAAAACAAGTTGCATGTGGAAAAATCAAAATAAAAAGTCTGGACTCTCACGCTCG
CTTGGTCCTGTAACTATTTTGTAGAATGGAAGACATCAACTTTGCGTCTCTGGCCCCGCGACACGGC
TCGCGCCCGTTCATGGGAAACTGGCAAGATATCGGCACCAGCAATATGAGCGGTGGCGCCTTCAGC
TGGGGCTCGCTGTGGAGCGGCATTAAAAATTTCGGTTCCACCGTTAAGAACTATGGCAGCAAGGCC
TGGAACAGCAGCACAGGCCAGATGCTGAGGGATAAGTTGAAAGAGCAAAATTTCCAACAAAAGGTG
GTAGATGGCCTGGCCTCTGGCATTAGCGGGGTGGTGGACCTGGCCAACCAGGCAGTGCAAAATAA
GATTAACAGTAAGCTTGATCCCCGCCCTCCCGTAGAGGAGCCTCCACCGGCCGTGGAGACAGTGT
CTCCAGAGGGGCGTGGCGAAAAGCGTCCGCGCCCCGACAGGGAAGAAACTCTGGTGACGCAAAT
AGACGAGCCTCCCTCGTACGAGGAGGCACTAAAGCAAGGCCTGCCCACCACCCGTCCCATCGCGC
CCATGGCTACCGGAGTGCTGGGCCAGCACACACCCGTAACGCTGGACCTGCCTCCCCCCGCCGA
CACCCAGCAGAAACCTGTGCTGCCAGGCCCGACCGCCGTTGTTGTAACCCGTCCTAGCCGCGCGT
CCCTGCGCCGCGCCGCCAGCGGTCCGCGATCGTTGCGGCCCGTAGCCAGTGGCAACTGGCAAAG
CACACTGAACAGCATCGTGGGTCTGGGGGTGCAATCCCTGAAGCGCCGACGATGCTTCTGATAGCT
AACGTGTCGTATGTGTGTCATGTATGCGTCCATGTCGCCGCCAGAGGAGCTGCTGAGCCGCCGCG
CGCCCGCTTTCCAAGATGGCTACCCCTTCGATGATGCCGCAGTGGTCTTACATGCACATCTCGGGC
CAGGACGCCTCGGAGTACCTGAGCCCCGGGCTGGTGCAGTTTGCCCGCGCCACCGAGACGTACT
TCAGCCTGAATAACAAGTTTAGAAACCCCACGGTGGCGCCTACGCACGACGTGACCACAGACCGGT
CCCAGCGTTTGACGCTGCGGTTCATCCCTGTGGACCGTGAGGATACTGCGTACTCGTACAAGGCGC
GGTTCACCCTAGCTGTGGGTGATAACCGTGTGCTGGACATGGCTTCCACGTACTTTGACATCCGCG
GCGTGCTGGACAGGGGCCCTACTTTTAAGCCCTACTCTGGCACTGCCTACAACGCCCTGGCTCCCA
AGGGTGCCCCAAATCCTTGCGAATGGGATGAAGCTGCTACTGCTCTTGAAATAAACCTAGAAGAAGA
GGACGATGACAACGAAGACGAAGTAGACGAGCAAGCTGAGCAGCAAAAAACTCACGTATTTGGGCA
GGCGCCTTATTCTGGTATAAATATTACAAAGGAGGGTATTCAAATAGGTGTCGAAGGTCAAACACCTA
AATATGCCGATAAAACATTTCAACCTGAACCTCAAATAGGAGAATCTCAGTGGTACGAAACAGAAATT
AATCATGCAGCTGGGAGAGTCCTAAAAAAGACTACCCCAATGAAACCATGTTACGGTTCATATGCAAA
ACCCACAAATGAAAATGGAGGGCAAGGCATTCTTGTAAAGCAACAAAATGGAAAGCTAGAAAGTCAA
GTGGAAATGCAATTTTTCTCAACTACTGAGGCAGCCGCAGGCAATGGTGATAACTTGACTCCTAAAG
TGGTATTGTACAGTGAAGATGTAGATATAGAAACCCCAGACACTCATATTTCTTACATGCCCACTATTA
AGGAAGGTAACTCACGAGAACTAATGGGCCAACAATCTATGCCCAACAGGCCTAATTACATTGCTTTT
AGGGACAATTTTATTGGTCTAATGTATTACAACAGCACGGGTAATATGGGTGTTCTGGCGGGCCAAG
CATCGCAGTTGAATGCTGTTGTAGATTTGCAAGACAGAAACACAGAGCTTTCATACCAGCTTTTGCTT
GATTCCATTGGTGATAGAACCAGGTACTTTTCTATGTGGAATCAGGCTGTTGACAGCTATGATCCAGA
TGTTAGAATTATTGAAAATCATGGAACTGAAGATGAACTTCCAAATTACTGCTTTCCACTGGGAGGTG
TGATTAATACAGAGACTCTTACCAAGGTAAAACCTAAAACAGGTCAGGAAAATGGATGGGAAAAAGAT

GCTACAGAATTTTCAGATAAAAATGAAATAAGAGTTGGAAATAATTTTGCCATGGAAATCAATCTAAAT
GCCAACCTGTGGAGAAATTTCCTGTACTCCAACATAGCGCTGTATTTGCCCGACAAGCTAAAGTACA
GTCCTTCCAACGTAAAAATTTCTGATAACCCAAACACCTACGACTACATGAACAAGCGAGTGGTGGC
TCCCGGGCTAGTGGACTGCTACATTAACCTTGGAGCACGCTGGTCCCTTGACTATATGGACAACGTC
AACCCATTTAACCACCACCGCAATGCTGGCCTGCGCTACCGCTCAATGTTGCTGGGCAATGGTCGC
TATGTGCCCTTCCACATCCAGGTGCCTCAGAAGTTCTTTGCCATTAAAAACCTCCTTCTCCTGCCGG
GCTCATACACCTACGAGTGGAACTTCAGGAAGGATGTTAACATGGTTCTGCAGAGCTCCCTAGGAAA
TGACCTAAGGGTTGACGGAGCCAGCATTAAGTTTGATAGCATTTGCCTTTACGCCACCTTCTTCCCC
ATGGCCCACAACACCGCCTCCACGCTTGAGGCCATGCTTAGAAACGACACCAACGACCAGTCCTTT
AACGACTATCTCTCCGCCGCCAACATGCTCTACCCTATACCCGCCAACGCTACCAACGTGCCCATAT
CCATCCCCTCCCGCAACTGGGCGGCTTTCCGCGGCTGGGCCTTCACGCGCCTTAAGACTAAGGAA
ACCCCATCACTGGGCTCGGGCTACGACCCTTATTACACCTACTCTGGCTCTATACCCTACCTAGATG
GAACCTTTTACCTCAACCACACCTTTAAGAAGGTGGCCATTACCTTTGACTCTTCTGTCAGCTGGCC
TGGCAATGACCGCCTGCTTACCCCCAACGAGTTTGAAATTAAGCGCTCAGTTGACGGGGAGGGTTA
CAACGTTGCCCAGTGTAACATGACCAAAGACTGGTTCCTGGTACAAATGCTAGCTAACTATAACATTG
GCTACCAGGGCTTCTATATCCCAGAGAGCTACAAGGACCGCATGTACTCCTTCTTTAGAAACTTCCA
GCCCATGAGCCGTCAGGTGGTGGATGATACTAAATACAAGGACTACCAACAGGTGGGCATCCTACA
CCAACACAACAACTCTGGATTTGTTGGCTACCTTGCCCCCACCATGCGCGAAGGACAGGCCTACCC
TGCTAACTTCCCCTATCCGCTTATAGGCAAGACCGCAGTTGACAGCATTACCCAGAAAAAGTTTCTTT
GCGATCGCACCCTTTGGCGCATCCCATTCTCCAGTAACTTTATGTCCATGGGCGCACTCACAGACCT
GGGCCAAAACCTTCTCTACGCCAACTCCGCCCACGCGCTAGACATGACTTTTGAGGTGGATCCCAT
GGACGAGCCCACCCTTCTTTATGTTTTGTTTGAAGTCTTTGACGTGGTCCGTGTGCACCAGCCGCA
CCGCGGCGTCATCGAAACCGTGTACCTGCGCACGCCCTTCTCGGCCGGCAACGCCACAACATAAA
GAAGCAAGCAACATCAACAACAGCTGCCGCCATGGGCTCCAGTGAGCAGGAACTGAAAGCCATTG
TCAAAGATCTTGGTTGTGGGCCATATTTTTTGGGCACCTATGACAAGCGCTTTCCAGGCTTTGTTTCT
CCACACAAGCTCGCCTGCGCCATAGTCAATACGGCCGGTCGCGAGACTGGGGGCGTACACTGGAT
GGCCTTTGCCTGGAACCCGCACTCAAAAACATGCTACCTCTTTGAGCCCTTTGGCTTTTCTGACCAG
CGACTCAAGCAGGTTTACCAGTTTGAGTACGAGTCACTCCTGCGCCGTAGCGCCATTGCTTCTTCC
CCCGACCGCTGTATAACGCTGGAAAAGTCCACCCAAAGCGTACAGGGGCCCAACTCGGCCGCCTG
TGGACTATTCTGCTGCATGTTTCTCCACGCCTTTGCCAACTGGCCCCAAACTCCCATGGATCACAAC
CCCACCATGAACCTTATTACCGGGGTACCCAACTCCATGCTCAACAGTCCCCAGGTACAGCCCACC
CTGCGTCGCAACCAGGAACAGCTCTACAGCTTCCTGGAGCGCCACTCGCCCTACTTCCGCAGCCA
CAGTGCGCAGATTAGGAGCGCCACTTCTTTTTGTCACTTGAAAAACATGTAAAAATAATGTACTAGAG
ACACTTTCAATAAAGGCAAATGCTTTTATTTGTACACTCTCGGGTGATTATTTACCCCCACCCTTGCC
GTCTGCGCCGTTTAAAAATCAAAGGGGTTCTGCCGCGCATCGCTATGCGCCACTGGCAGGGACAC
GTTGCGATACTGGTGTTTAGTGCTCCACTTAAACTCAGGCACAACCATCCGCGGCAGCTCGGTGAA
GTTTTCACTCCACAGGCTGCGCACCATCACCAACGCGTTTAGCAGGTCGGGCGCCGATATCTTGAA
GTCGCAGTTGGGGCCTCCGCCCTGCGCGCGCGAGTTGCGATACACAGGGTTGCAGCACTGGAAC
ACTATCAGCGCCGGGTGGTGCACGCTGGCCAGCACGCTCTTGTCGGAGATCAGATCCGCGTCCAG
GTCCTCCGCGTTGCTCAGGGCGAACGGAGTCAACTTTGGTAGCTGCCTTCCCAAAAAGGGCGCGT
GCCCAGGCTTTGAGTTGCACTCGCACCGTAGTGGCATCAAAAGGTGACCGTGCCCGGTCTGGGCG
TTAGGATACAGCGCCTGCATAAAAGCCTTGATCTGCTTAAAAGCCACCTGAGCCTTTGCGCCTTCAG
AGAAGAACATGCCGCAAGACTTGCCGGAAAACTGATTGGCCGGACAGGCCGCGTCGTGCACGCAG
CACCTTGCGTCGGTGTTGGAGATCTGCACCACATTTCGGCCCCACCGGTTCTTCACGATCTTGGCC

TTGCTAGACTGCTCCTTCAGCGCGCGCTGCCCGTTTTCGCTCGTCACATCCATTTCAATCACGTGCT
CCTTATTTATCATAATGCTTCCGTGTAGACACTTAAGCTCGCCTTCGATCTCAGCGCAGCGGTGCAG
CCACAACGCGCAGCCCGTGGGCTCGTGATGCTTGTAGGTCACCTCTGCAAACGACTGCAGGTACG
CCTGCAGGAATCGCCCCATCATCGTCACAAAGGTCTTGTTGCTGGTGAAGGTCAGCTGCAACCCGC
GGTGCTCCTCGTTCAGCCAGGTCTTGCATACGGCCGCCAGAGCTTCCACTTGGTCAGGCAGTAGTT
TGAAGTTCGCCTTTAGATCGTTATCCACGTGGTACTTGTCCATCAGCGCGCGCGCAGCCTCCATGC
CCTTCTCCCACGCAGACACGATCGGCACACTCAGCGGGTTCATCACCGTAATTTCACTTTCCGCTTC
GCTGGGCTCTTCCTCTTCCTCTTGCGTCCGCATACCACGCGCCACTGGGTCGTCTTCATTCAGCCG
CCGCACTGTGCGCTTACCTCCTTTGCCATGCTTGATTAGCACCGGTGGGTTGCTGAAACCCACCATT
TGTAGCGCCACATCTTCTCTTTCTTCCTCGCTGTCCACGATTACCTCTGGTGATGGCGGGCGCTCG
GGCTTGGGAGAAGGGCGCTTCTTTTTCTTCTTGGGCGCAATGGCCAAATCCGCCGCCGAGGTCGA
TGGCCGCGGGCTGGGTGTGCGCGGCACCAGCGCGTCTTGTGATGAGTCTTCCTCGTCCTCGGAC
TCGATACGCCGCCTCATCCGCTTTTTTGGGGGCGCCCGGGGAGGCGGCGGCGACGGGGACGGG
GACGACACGTCCTCCATGGTTGGGGGACGTCGCGCCGCACCGCGTCCGCGCTCGGGGGTGGTTT
CGCGCTGCTCCTCTTCCCGACTGGCCATTTCCTTCTCCTATAGGCAGAAAAAAATCATGGAGTCAGT
CGAGAAGAAGGACAGCCTAACCGCCCCCTCTGAGTTCGCCACCACCGCCTCCACCGATGCCGCCA
ACGCGCCTACCACCTTCCCCGTCGAGGCACCCCCGCTTGAGGAGGAGGAAGTGATTATCGAGCAG
GACCCAGGTTTTGTAAGCGAAGACGACGAGGACCGCTCAGTACCAACAGAGGATAAAAAGCAAGAC
CAGGACAACGCAGAGGCAAACGAGGAACAAGTCGGGCGGGGGGACGAAAGGCATGGCGACTACC
TAGATGTGGGAGACGACGTGCTGTTGAAGCATCTGCAGCGCCAGTGCGCCATTATCTGCGACGCGT
TGCAAGAGCGCAGCGATGTGCCCCTCGCCATAGCGGATGTCAGCCTTGCCTACGAACGCCACCTAT
TCTCACCGCGCGTACCCCCCAAACGCCAAGAAAACGGCACATGCGAGCCCAACCCGCGCCTCAAC
TTCTACCCCGTATTTGCCGTGCCAGAGGTGCTTGCCACCTATCACATCTTTTTCCAAAACTGCAAGAT
ACCCCTATCCTGCCGTGCCAACCGCAGCCGAGCGGACAAGCAGCTGGCCTTGCGGCAGGGCGCT
GTCATACCTGATATCGCCTCGCTCAACGAAGTGCCAAAAATCTTTGAGGGTCTTGGACGCGACGAG
AAGCGCGCGGCAAACGCTCTGCAACAGGAAAACAGCGAAAATGAAAGTCACTCTGGAGTGTTGGT
GGAACTCGAGGGTGACAACGCGCGCCTAGCCGTACTAAAACGCAGCATCGAGGTCACCCACTTTG
CCTACCCGGCACTTAACCTACCCCCCAAGGTCATGAGCACAGTCATGAGTGAGCTGATCGTGCGCC
GTGCGCAGCCCCTGGAGAGGGATGCAAATTTGCAAGAACAAACAGAGGAGGGCCTACCCGCAGTT
GGCGACGAGCAGCTAGCGCGCTGGCTTCAAACGCGCGAGCCTGCCGACTTGGAGGAGCGACGCA
AACTAATGATGGCCGCAGTGCTCGTTACCGTGGAGCTTGAGTGCATGCAGCGGTTCTTTGCTGACC
CGGAGATGCAGCGCAAGCTAGAGGAAACATTGCACTACACCTTTCGACAGGGCTACGTACGCCAGG
CCTGCAAGATCTCCAACGTGGAGCTCTGCAACCTGGTCTCCTACCTTGGAATTTTGCACGAAAACC
GCCTTGGGCAAAACGTGCTTCATTCCACGCTCAAGGGCGAGGCGCGCCGCGACTACGTCCGCGAC
TGCGTTTACTTATTTCTATGCTACACCTGGCAGACGGCCATGGGCGTTTGGCAGCAGTGCTTGGAG
GAGTGCAACCTCAAGGAGCTGCAGAAACTGCTAAAGCAAAACTTGAAGGACCTATGGACGGCCTTC
AACGAGCGCTCCGTGGCCGCGCACCTGGCGGACATCATTTTCCCCGAACGCCTGCTTAAAACCCT
GCAACAGGGTCTGCCAGACTTCACCAGTCAAAGCATGTTGCAGAACTTTAGGAACTTTATCCTAGAG
CGCTCAGGAATCTTGCCCGCCACCTGCTGTGCACTTCCTAGCGACTTTGTGCCCATTAAGTACCGC
GAATGCCCTCCGCCGCTTTGGGGCCACTGCTACCTTCTGCAGCTAGCCAACTACCTTGCCTACCAC
TCTGACATAATGGAAGACGTGAGCGGTGACGGTCTACTGGAGTGTCACTGTCGCTGCAACCTATGC
ACCCCGCACCGCTCCCTGGTTTGCAATTCGCAGCTGCTTAACGAAAGTCAAATTATCGGTACCTTTG
AGCTGCAGGGTCCCTCGCCTGACGAAAAGTCCGCGGCTCCGGGGTTGAAACTCACTCCGGGGCT
GTGGACGTCGGCTTACCTTCGCAAATTTGTACCTGAGGACTACCACGCCCACGAGATTAGGTTCTAC

GAAGACCAATCCCGCCCGCCTAATGCGGAGCTTACCGCCTGCGTCATTACCCAGGGCCACATTCTT
GGCCAATTGCAAGCCATCAACAAAGCCCGCCAAGAGTTTCTGCTACGAAAGGGACGGGGGGTTTA
CTTGGACCCCCAGTCCGGCGAGGAGCTCAACCCAATCCCCCCGCCGCCGCAGCCCTATCAGCAGC
AGCCGCGGGCCCTTGCTTCCCAGGATGGCACCCAAAAAGAAGCTGCAGCTGCCGCCGCGCCACC
CACGGACGAGGAGGAATACTGGGACAGTCAGGCAGAGGAGGTTTTGGACGAGGAGGAGGAGGAC
ATGATGGAAGACTGGGAGAGCCTAGACGAGGAAGCTTCCGAGGTCGAAGAGGTGTCAGACGAAAC
ACCGTCACCCTCGGTCGCATTCCCCTCGCCGGCGCCCCAGAAATCGGCAACCGGTTCCAGCATGG
CTACAACCTCCGCTCCTCAGGCGCCGCCGGCACTGCCCGTTCGCCGACCCAACCGTAGATGGGAC
ACCACTGGAACCAGGGCCGGTAAGTCCAAGCAGCCGCCGCCGTTAGCCCAAGAGCAACAACAGC
GCCAAGGCTACCGCTCATGGCGCGGGCACAAGAACGCCATAGTTGCTTGCTTGCAAGACTGTGGG
GGCAACATCTCCTTCGCCCGCCGCTTTCTTCTCTACCATCACGGCGTGGCCTTCCCCCGTAACATC
CTGCATTACTACCGTCATCTCTACAGCCCATACTGCACCGGCGGCAGCGGCAGCAACAGCAGCGGC
CACACAGAAGCAAAGGCGACCGGATAGCAAGACTCTGACAAAGCCCAAGAAATCCACAGCGGCGG
CAGCAGCAGGAGGAGGAGCGCTGCGTCTGGCGCCCAACGAACCCGTATCGACCCGCGAGCTTAG
AAACAGGATTTTTCCCACTCTGTATGCTATATTTCAACAGAGCAGGGGCCAAGAACAAGAGCTGAAA
ATAAAAAACAGGTCTCTGCGATCCCTCACCCGCAGCTGCCTGTATCACAAAAGCGAAGATCAGCTTC
GGCGCACGCTGGAAGACGCGGAGGCTCTCTTCAGTAAATACTGCGCGCTGACTCTTAAGGACTAGT
TTCGCGCCCTTTCTCAAATTTAAGCGCGAAAACTACGTCATCTCCAGCGGCCACACCCGGCGCCAG
CACCTGTTGTCAGCGCCATTATGAGCAAGGAAATTCCCACGCCCTACATGTGGAGTTACCAGCCACA
AATGGGACTTGCGGCTGGAGCTGCCCAAGACTACTCAACCCGAATAAACTACATGAGCGCGGACC
CCACATGATATCCCGGGTCAACGGAATACGCGCCCACCGAAACCGAATTCTCCTGGAACAGGCGGC
TATTACCACCACACCTCGTAATAACCTTAATCCCCGTAGTTGGCCCGCTGCCCTGGTGTACCAGGAA
AGTCCCGCTCCCACCACTGTGGTACTTCCCAGAGACGCCCAGGCCGAAGTTCAGATGACTAACTCA
GGGGCGCAGCTTGCGGGCGGCTTTCGTCACAGGGTGCGGTCGCCCGGGCAGGGTATAACTCACC
TGACAATCAGAGGGCGAGGTATTCAGCTCAACGACGAGTCGGTGAGCTCCTCGCTTGGTCTCCGT
CCGGACGGGACATTTCAGATCGGCGGCGCCGGCCGCTCTTCATTCACGCCTCGTCAGGCAATCCT
AACTCTGCAGACCTCGTCCTCTGAGCCGCGCTCTGGAGGCATTGGAACTCTGCAATTTATTGAGGA
GTTTGTGCCATCGGTCTACTTTAACCCCTTCTCGGGACCTCCCGGCCACTATCCGGATCAATTTATTC
CTAACTTTGACGCGGTAAAGGACTCGGCGGACGGCTACGACTGAATGTTAAGTGGAGAGGCAGAG
CAACTGCGCCTGAAACACCTGGTCCACTGTCGCCGCCACAAGTGCTTTGCCCGCGACTCCGGTGA
GTTTTGCTACTTTGAATTGCCCGAGGATCATATCGAGGGCCCGGCGCACGGCGTCCGGCTTACCGC
CCAGGGAGAGCTTGCCCGTAGCCTGATTCGGGAGTTTACCCAGCGCCCCCTGCTAGTTGAGCGGG
ACAGGGGACCCTGTGTTCTCACTGTGATTTGCAACTGTCCTAACCCTGGATTACATCAAGATCCTCT
AGTTAATGGATCCAATCGATATCGCGAAGTCGATCGACGGATCGGGAGATCTCCCGATCCCCTATGG
TGCACTCTCAGTACAATCTGCTCTGATGCCGCATAGTTAAGCCAGTATCTGCTCCCTGCTTGTGTGTT
GGAGGTCGCTGAGTAGTGCGCGAGCAAAATTTAAGCTACAACAAGGCAAGGCTTGACCGACAATTG
CATGAAGAATCTGCTTAGGGTTAGGCGTTTTGCGCTGCTTCGGCTAGCTGCAGTAACGCCATTTTGC
AAGGCATGGAAAAATACCAAACCAAGAATAGAGAAGTTCAGATCAAGGGCGGGTACATGAAAATAGC
TAACGTTGGGCCAAACAGGATATCTGCGGTGAGCAGTTTCGGCCCCGGCCCGGGGCCAAGAACAG
ATGGTCACCGCAGTTTCGGCCCCGGCCCGAGGCCAAGAACAGATGGTCCCCAGATATGGCCCAAC
CCTCAGCAGTTTCTTAAGACCCATCAGATGTTTCCAGGCTCCCCCAAGGACCTGAAATGACCCTGC
GCCTTATTTGAATTAACCAATCAGCCTGCTTCTCGCTTCTGTTCGCGCGCTTCTGCTTCCCGAGCTC
TATAAAAGAGCTCACAACCCCTCACTCGGCGCGCCAGTCCTCCGACAGACTGAGTCGCCCGGGTA
CCGCGATCGCCATGGATTTTTTTCGGGTAGTGGAAAACCAGCAGCCTCCCGCGACGATGCCCCTCA

ACGTTAGCTTCACCAACAGGAACTATGACCTCGACTACGACTCGGTGCAGCCGTATTTCTACTGCGA
CGAGGAGGAGAACTTCTACCAGCAGCAGCAGCAGAGCGAGCTGCAGCCCCCGGCGCCCAGCGAG
GATATCTGGAAGAAATTCGAGCTGCTGCCCACCCCGCCCCTGTCCCCTAGCCGCCGCTCCGGGCT
CTGCTCGCCCTCCTACGTTGCGGTCACACCCTTCTCCCTTCGGGGAGACAACGACGGCGGTGGCG
GGAGCTTCTCCACGGCCGACCAGCTGGAGATGGTGACCGAGCTGCTGGGAGGAGACATGGTGAA
CCAGAGTTTCATCTGCGACCCGGACGACGAGACCTTCATCAAAAACATCATCATCCAGGACTGTATG
TGGAGCGGCTTCTCGGCCGCCGCCAAGCTCGTCTCAGAGAAGCTGGCCTCCTACCAGGCTGCGC
GCAAAGACAGCGGCAGCCCGAACCCCGCCCGCGGCCACAGCGTCTGCTCCACCTCCAGCTTGTA
CCTGCAGGATCTGAGCGCCGCCGCCTCAGAGTGCATCGACCCCTCGGTGGTCTTCCCCTACCCTC
TCAACGACAGCAGCTCGCCCAAGTCCTGCGCCTCGCAAGACTCCAGCGCCTTCTCTCCGTCCTCG
GATTCTCTGCTCTCCTCGACGGAGTCCTCCCCGCAGGGCAGCCCCGAGCCCCTGGTGCTCCATGA
GGAGACACCGCCCACCACCAGCAGCGACTCTGAGGAGGAACAAGAAGATGAGGAAGAAATCGATG
TTGTTTCTGTGGAAAAGAGGCAGGCTCCTGGCAAAAGGTCAGAGTCTGGATCACCTTCTGCTGGAG
GCCACAGCAAACCTCCTCACAGCCCACTGGTCCTCAAGAGGTGCCACGTCTCCACACATCAGCAC
AACTACGCAGCGCCTCCCTCCACTCGGAAGGACTATCCTGCTGCCAAGAGGGTCAAGTTGGACAG
TGTCAGAGTCCTGAGACAGAATCAGCAACAACCGAAAATGCACCAGCCCCAGGTCCTCGGACACC
GAGGAGAATGTCAAGAGGCGAACACACAACGTCTTGGAGCGCCAGAGGAGGAACGAGCTAAAACG
GAGCTTTTTTGCCCTGCGTGACCAGATCCCGGAGTTGGAAAACAATGAAAAGGCCCCCAAGGTAGT
TATCCTTAAAAAAGCCACAGCATACATCCTGTCCGTCCAAGCAGAGGAGCAAAAGCTCATTTCTGAA
GAGGACTTGTTGCGGAAACGACGAGAACAGTTGAAACACAAACTTGAACAGCTACGGAACTCTTGT
GCGCAATGTACTAACTACGCTTTGTTGAAACTCGCTGGCGATGTTGAAAGTAACCCCGGTCCTATGA
GGCAGCCACCTGGCGAGTCTGACATGGCTGTCAGCGACGCGCTGCTCCCATCTTTCTCCACGTTC
GCGTCTGGCCCGGCGGGAAGGGAGAAGACACTGCGTCAAGCAGGTGCCCCGAATAACCGCTGGC
GGGAGGAGCTCTCCCACATGAAGCGACTTCCCCCAGTGCTTCCCGGCCGCCCCTATGACCTGGCG
GCGGCGACCGTGGCCACAGACCTGGAGAGCGGCGGAGCCGGTGCGGCTTGCGGCGGTAGCAAC
CTGGCGCCCCTACCTCGGAGAGAGACCGAGGAGTTCAACGATCTCCTGGACCTGGACTTTATTCTC
TCCAATTCGCTGACCCATCCTCCGGAGTCAGTGGCCGCCACCGTGTCCTCGTCAGCGTCAGCCTC
CTCTTCGTCGTCGCCGTCGAGCAGCGGCCCTGCCAGCGCGCCCTCCACCTGCAGCTTCACCTATC
CGATCCGGGCCGGGAACGACCCGGGCGTGGCGCCGGGCGGCACGGGCGGAGGCCTCCTCTATG
GCAGGGAGTCCGCTCCCCCTCCGACGGCTCCCTTCAACCTGGCGGACATCAACGACGTGAGCCC
CTCGGGCGGCTTCGTGGCCGAGCTCCTGCGGCCAGAATTGGACCCGGTGTACATTCCGCCGCAG
CAGCCGCAGCCGCCAGGTGGCGGGCTGATGGGCAAGTTCGTGCTGAAGGCGTCGCTGAGCGCCC
CTGGCAGCGAGTACGGCAGCCCGTCGGTCATCAGCGTCAGCAAAGGCAGCCCTGACGGCAGCCA
CCCGGTGGTGGTGGCGCCCTACAACGGCGGGCCGCCGCGCACGTGCCCCAAGATCAAGCAGGAG
GCGGTCTCTTCGTGCACCCACTTGGGCGCTGGACCCCCTCTCAGCAATGGCCACCGGCCGGCTG
CACACGACTTCCCCCTGGGGCGGCAGCTCCCCAGCAGGACTACCCCGACCCTGGGTCTTGAGGA
AGTGCTGAGCAGCAGGGACTGTCACCCTGCCCTGCCGCTTCCTCCCGGCTTCCATCCCCACCCGG
GGCCCAATTACCCATCCTTCCTGCCCGATCAGATGCAGCCGCAAGTCCCGCCGCTCCATTACCAAG
AGCTCATGCCACCCGGTTCCTGCATGCCAGAGGAGCCCAAGCCAAAGAGGGGAAGACGATCGTGG
CCCCGGAAAAGGACCGCCACCCACACTTGTGATTACGCGGGCTGCGGCAAAACCTACACAAAGAG
TTCCCATCTCAAGGCACACCTGCGAACCCACACAGGTGAGAAACCTTACCACTGTGACTGGGACGG
CTGTGGATGGAAATTCGCCCGCTCAGATGAACTGACCAGGCACTACCGTAAACACACGGGGCACC
GCCCGTTCCAGTGCCAAAAATGCGACCGAGCATTTTCCAGGTCGGACCACCTCGCCTTACACATGA
AGAGGCATTTTTAACTCGAGTCTAGAGGGCCCGTTTAAACCCGCTGATCAGCCTCGACTGTGCCTTC

TAGTTGCCAGCCATCTGTTGTTGTTTGCCCCTCCCCCGTGCCTTCCTTGACCCTGGAAGGTGCCAC
TCCCACTGTCCTTTCCTAATAAAATGAGGAAATTGCATCGCATTGTCTGAGTAGGTGTCATTCTATTCT
GGGGGGTGGGGTGGGGCAGGACAGCAAGGGGGAGGATTGGGAAGACAATAGCAGGCATGCTGG
GGATGCGGTGGGCTCTATGGCTTCTGAGGCGGAAAGAACCAGCTGGGGCTCTAGGGGGTATCCCC
ACGCGCCCTGTAGCGGCGCATTAAGCGCGGCGGGTGTGGTGGTTACGCGCAGCGTGACCGCTAC
ACTTGCCAGCGCCCTAGCGCCCGCTCCTTTCGCTTTCTTCCCTTCCTTTCTCGCCACGTTCGCCGG
CTTTCCCCGTCAAGCTCTAAATCGGGGGCTCCCTTTAGGGTTCCGATTTAGTGCTTTACGGCACCTC
GACCCCAAAAAACTTGATTAGGGTGATGGTTCACATCGAATTCCTGCAGCCCGGGGGATCCATTAAC
TAATAAAAAAAAATAATAAAGCATCACTTACTTAAAATCAGTTAGCAAATTTCTGTCCAGTTTATTCAGC
AGCACCTCCTTGCCCTCCTCCCAGCTCTGGTATTGCAGCTTCCTCCTGGCTGCAAACTTTCTCCACA
ATCTAAATGGAATGTCAGTTTCCTCCTGTTCCTGTCCATCCGCACCCACTATCTTCATGTTGTTGCAG
ATGAAGCGCGCAAGACCGTCTGAAGATACCTTCAACCCCGTGTATCCATATGACACGGAAACCGGT
CCTCCAACTGTGCCTTTTCTTACTCCTCCCTTTGTATCCCCCAATGGGTTTCAAGAGAGTCCCCCTG
GAGTTCTTACTTTAAAATGTTTAACCCCACTAACAACCACAGGCGGATCTCTACAGCTAAAAGTGGGA
GGGGGACTTACAGTGGATGACACTGATGGTACCTTACAAGAAAACATACGTGCTACAGCACCCATTA
CTAAAAATAATCACTCTGTAGAACTATCCATTGGAAATGGATTAGAAACTCAAAACAATAAACTATGTG
CCAAATTGGGAAATGGGTTAAAATTTAACAACGGTGACATTTGTATAAAGGATAGTATTAACACCTTAT
GGACTGGAATAAACCCTCCACCTAACTGTCAAATTGTGGAAAACACTAATACAAATGATGGCAAACTT
ACTTTAGTATTAGTAAAAAACGGAGGGCTTGTTAATGGCTACGTGTCTCTAGTTGGTGTATCAGACAC
TGTGAACCAAATGTTCACACAAAAGACAGCAAACATCCAATTAAGATTATATTTTGACTCTTCTGGAAA
TCTATTAACTGATGAATCAGACTTAAAAATTCCACTTAAAAATAAATCTTCTACAGCGACCAGTGAAAC
TGTAGCCAGCAGCAAAGCCTTTATGCCAAGTACTACAGCTTATCCCTTCAACACCACTACTAGGGATA
GTGAAAACTACATTCATGGAATATGTTACTACATGACTAGTTATGATAGAAGTCTATTTCCCTTGAACAT
TTCTATAATGCTAAACAGCCGTATGATTTCTTCCAATGTTGCCTATGCCATACAATTTGAATGGAATCTA
AATGCAAGTGAATCTCCAGAAAGCAACATAGCTACGCTGACCACATCCCCCTTTTTCTTTTCTTACAT
TACAGAAGACGACAACTAAAATGCCCAAGAATAAAGAATCGTTTGTGTTATGTTTCAACGTGTTTATTT
TTCAATTGCAGAAAATTTCAAGTCATTTTTCATTCAGTAGTATAGCCCCACCACCACATAGCTTATACA
GATCACCGTACCTTAATCAAACTCACAGAACCCTAGTATTCAACCTGCCACCTCCCTCCCAACACAC
AGAGTACACAGTCCTTTCTCCCCGGCTGGCCTTAAAAAGCATCATATCATGGGTAACAGACATATTCT
TAGGTGTTATATTCCACACGGTTTCCTGTCGAGCCAAACGCTCATCAGTGATATTAATAAACTCCCCG
GGCAGCTCACTTAAGTTCATGTCGCTGTCCAGCTGCTGAGCCACAGGCTGCTGTCCAACTTGCGGT
TGCTTAACGGGCGGCGAAGGAGAAGTCCACGCCTACATGGGGGTAGAGTCATAATCGTGCATCAGG
ATAGGGCGGTGGTGCTGCAGCAGCGCGCGAATAAACTGCTGCCGCCGCCGCTCCGTCCTGCAGG
AATACAACATGGCAGTGGTCTCCTCAGCGATGATTCGCACCGCCCGCAGCATAAGGCGCCTTGTCC
TCCGGGCACAGCAGCGCACCCTGATCTCACTTAAATCAGCACAGTAACTGCAGCACAGCACCACAA
TATTGTTCAAAATCCCACAGTGCAAGGCGCTGTATCCAAAGCTCATGGCGGGGACCACAGAACCCA
CGTGGCCATCATACCACAAGCGCAGGTAGATTAAGTGGCGACCCCTCATAAACACGCTGGACATAAA
CATTACCTCTTTTGGCATGTTGTAATTCACCACCTCCCGGTACCATATAAACCTCTGATTAAACATGGC
GCCATCCACCACCATCCTAAACCAGCTGGCCAAAACCTGCCCGCCGGCTATACACTGCAGGGAACC
GGGACTGGAACAATGACAGTGGAGAGCCCAGGACTCGTAACCATGGATCATCATGCTCGTCATGAT
ATCAATGTTGGCACAACACAGGCACACGTGCATACACTTCCTCAGGATTACAAGCTCCTCCCGCGTT
AGAACCATATCCCAGGGAACAACCCATTCCTGAATCAGCGTAAATCCCACACTGCAGGGAAGACCT
CGCACGTAACTCACGTTGTGCATTGTCAAAGTGTTACATTCGGGCAGCAGCGGATGATCCTCCAGTA
TGGTAGCGCGGGTTTCTGTCTCAAAAGGAGGTAGACGATCCCTACTGTACGGAGTGCGCCGAGACA

ACCGAGATCGTGTTGGTCGTAGTGTCATGCCAAATGGAACGCCGGACGTAGTCATATTTCCTGAAGC
AAAACCAGGTGCGGGCGTGACAAACAGATCTGCGTCTCCGGTCTCGCCGCTTAGATCGCTCTGTGT
AGTAGTTGTAGTATATCCACTCTCTCAAAGCATCCAGGCGCCCCCTGGCTTCGGGTTCTATGTAAACT
CCTTCATGCGCCGCTGCCCTGATAACATCCACCACCGCAGAATAAGCCACACCCAGCCAACCTACA
CATTCGTTCTGCGAGTCACACACGGGAGGAGCGGGAAGAGCTGGAAGAACCATGTTTTTTTTTATT
CCAAAAGATTATCCAAAACCTCAAAATGAAGATCTATTAAGTGAACGCGCTCCCCTCCGGTGGCGTG
GTCAAACTCTACAGCCAAAGAACAGATAATGGCATTTGTAAGATGTTGCACAATGGCTTCCAAAAGG
CAAACGGCCCTCACGTCCAAGTGGACGTAAAGGCTAAACCCTTCAGGGTGAATCTCCTCTATAAACA
TTCCAGCACCTTCAACCATGCCCAAATAATTCTCATCTCGCCACCTTCTCAATATATCTCTAAGCAAAT
CCCGAATATTAAGTCCGGCCATTGTAAAAATCTGCTCCAGAGCGCCCTCCACCTTCAGCCTCAAGCA
GCGAATCATGATTGCAAAAATTCAGGTTCCTCACAGACCTGTATAAGATTCAAAAGCGGAACATTAAC
AAAAATACCGCGATCCCGTAGGTCCCTTCGCAGGGCCAGCTGAACATAATCGTGCAGGTCTGCACG
GACCAGCGCGGCCACTTCCCCGCCAGGAACCATGACAAAAGAACCCACACTGATTATGACACGCAT
ACTCGGAGCTATGCTAACCAGCGTAGCCCCGATGTAAGCTTGTTGCATGGGCGGCGATATAAAATGC
AAGGTGCTGCTCAAAAAATCAGGCAAAGCCTCGCGCAAAAAAGAAAGCACATCGTAGTCATGCTCAT
GCAGATAAAGGCAGGTAAGCTCCGGAACCACCACAGAAAAAGACACCATTTTTCTCTCAAACATGTC
TGCGGGTTTCTGCATAAACACAAAATAAAATAACAAAAAAACATTTAAACATTAGAAGCCTGTCTTACA
ACAGGAAAAACAACCCTTATAAGCATAAGACGGACTACGGCCATGCCGGCGTGACCGTAAAAAAACT
GGTCACCGTGATTAAAAAGCACCACCGACAGCTCCTCGGTCATGTCCGGAGTCATAATGTAAGACTC
GGTAAACACATCAGGTTGATTCACATCGGTCAGTGCTAAAAAGCGACCGAAATAGCCCGGGGGAATA
CATACCCGCAGGCGTAGAGACAACATTACAGCCCCCATAGGAGGTATAACAAAATTAATAGGAGAGA
AAAACACATAAACACCTGAAAAACCCTCCTGCCTAGGCAAAATAGCACCCTCCCGCTCCAGAACAAC
ATACAGCGCTTCCACAGCGGCAGCCATAACAGTCAGCCTTACCAGTAAAAAAGAAAACCTATTAAAA
AAACACCACTCGACACGGCACCAGCTCAATCAGTCACAGTGTAAAAAAGGGCCAAGTGCAGAGCG
AGTATATATAGGACTAAAAAATGACGTAACGGTTAAAGTCCACAAAAAACACCCAGAAAACCGCACGC
GAACCTACGCCCAGAAACGAAAGCCAAAAAACCCACAACTTCCTCAAATCGTCACTTCCGTTTTCCC
ACGTTACGTCACTTCCCATTTTAAGAAAACTACAATTCCCAACACATACAAGTTACTCCGCCCTAAAA
CCTACGTCACCCGCCCCGTTCCCACGCCCCGCGCCACGTCACAAACTCCACCCCCTCATTATCATA
TTGGCTTCAATCCAAAATAAGGTATATTATTGATGATGTTACATCGTTAATTAACGATTTCGAACCCGGG
GTACCGAATTCCTCGAGTCTAGAGGAGCATGCGACGTCGCAATTCGCCCTATAGTGAGTCGTATTAC
AATTCACTGGCCGTCGTTTTACAACGTCGTGACTGGGAAAACCCTGGCGTTACCCAACTTAATCGCC
TTGCAGCACATCCCCCTTTCGCCAGCTGGCGTAATAGCGAAGAGGCCCGCACCGATCGCCCTTCCC
AACAGTTGCGCAGCCTGAATGGCGAATGGAAATTGTAAGCGTTAATATTTTGTTAAAATTCGCGTTAA
ATTTTTGTTAAATCAGCTCATTTTTTAACCAATAGGCCGAAATCGGCAAAATCCCTTATAAATCAAAAGA
ATAGACCGAGATAGGGTTGAGTGTTGTTCCAGTTTGGAACAAGAGTCCACTATTAAAGAACGTGGAC
TCCAACGTCAAAGGGCGAAAAACCGTCTATCAGGGCGATGGCCCACTACGTGAACCATCACCCTAA
TCAAGTTTTTTGGGGTCGAGGTGCCGTAAAGCACTAAATCGGAACCCTAAAGGGAGCCCCCGATTT
AGAGCTTGACGGGGAAAGCCGGCGAACGTGGCGAGAAAGGAAGGGAAGAAAGCGAAAGGAGCG
GGCGCTAGGGCGCTGGCAAGTGTAGCGGTCACGCTGCGCGTAACCACCACACCCGCCGCGCTTAA
TGCGCCGCTACAGGGCGCGTCCTGATGCGGTATTTTCTCCTTACGCATCTGTGCGGTATTTCACACC
GCATACAGGTGGCACTTTTCGGGGAAATGTGCGCGGAACCCCTATTTGTTTATTTTTCTAAATACATT
CAAATATGTATCCGCTCATGAGACAATAACCCTGATAAATGCTTCAATAATATTGAAAAAGGAAGAGTA
TGAGTATTCAACATTTCCGTGTCGCCCTTATTCCCTTTTTTGCGGCATTTTGCCTTCCTGTTTTTGCTC
ACCCAGAAACGCTGGTGAAAGTAAAAGATGCTGAAGATCAGTTGGGTGCACGAGTGGGTTACATCG

AACTGGATCTCAACAGCGGTAAGATCCTTGAGAGTTTTCGCCCCGAAGAACGTTTTCCAATGATGAG
CACTTTTAAAGTTCTGCTATGTGGCGCGGTATTATCCCGTATTGACGCCGGGCAAGAGCAACTCGGT
CGCCGCATACACTATTCTCAGAATGACTTGGTTGAGTACTCACCAGTCACAGAAAAGCATCTTACGG
ATGGCATGACAGTAAGAGAATTATGCAGTGCTGCCATAACCATGAGTGATAACACTGCGGCCAACTT
ACTTCTGACAACGATCGGAGGACCGAAGGAGCTAACCGCTTTTTTGCACAACATGGGGGATCATGT
AACTCGCCTTGATCGTTGGGAACCGGAGCTGAATGAAGCCATACCAAACGACGAGCGTGACACCAC
GATGCCTGTAGCAATGGCAACAACGTTGCGCAAACTATTAACTGGCGAACTACTTACTCTAGCTTCC
CGGCAACAATTAATAGACTGGATGGAGGCGGATAAAGTTGCAGGACCACTTCTGCGCTCGGCCCTT
CCGGCTGGCTGGTTTATTGCTGATAAATCTGGAGCCGGTGAGCGTGGGTCTCGCGGTATCATTGCA
GCACTGGGGCCAGATGGTAAGCCCTCCCGTATCGTAGTTATCTACACGACGGGGAGTCAGGCAACT
ATGGATGAACGAAATAGACAGATCGCTGAGATAGGTGCCTCACTGATTAAGCATTGGTAACTGTCAG
ACCAAGTTTACTCATATATACTTTAGATTGATTTAAAACTTCATTTTTAATTTAAAAGGATCTAGGTGAAG
ATCCTTTTTGATAATCTCATGACCAAAATCCCTTAACGTGAGTTTTCGTTCCACTGAGCGTCAGACCC
CGTAGAAAAGATCAAAGGATCTTCTTGAGATCCTTTTTTTCTGCGCGTAATCTGCTGCTTGCAAACAA
AAAAACCACCGCTACCAGCGGTGGTTTGTTTGCCGGATCAAGAGCTACCAACTCTTTTTCCGAAGGT
AACTGGCTTCAGCAGAGCGCAGATACCAAATACTGTTCTTCTAGTGTAGCCGTAGTTAggccaccacttca
agaactctgtagcacCGCCTACATACCTCGCTCTGCTAATCCTGTTACCAGTGGCTGCTGCCAGTGGCGAT
AAGTCGTGTCTTACCGGGTTGGACTCAAGACGATAGTTACCGGATAAGGCGCAGCGGTCGGGCTGA
ACGGGGGGTTCGTGCACACAGCCCAGCTTGGAGCGAACGACCTACACCGAACTGAGATACCTACA
GCGTGAGCTATGAGAAAGCGCCACGCTTCCCGAAGGGAGAAAGGCGGACAGGTATCCGGTAAGCG
GCAGGGTCGGAACAGGAGAGCGCACGAGGGAGCTTCCAGGGGGAAACGCCTGGTATCTTTATAGT
CCTGTCGGGTTTCGCCACCTCTGACTTGAGCGTCGATTTTTGTGATGCTCGTCAGGGGGGCGGAG
CCTATGGAAAAACGCCAGCAACGCGGCCTTTTTACGGTTCCTGGCCTTTTGCTGGCCTTTTGCTCA
CATGTTCTTTCCTGCGTTATCCCCTGATTCTGTGGATAACCGTATTACCGCCTTTGAGTGAGCTGATA
CCGCTCGCCGCAGCCGAACGACCGAGCGCAGCGAGTCAGTGAGCGAGGAAGCGGAAGAGCGCC
CAATACGCAAACCGCCTCTCCCCGCGCGTTGGCCGATTCATTAATGCAGCTGGCACGACAGGTTTC
CCGACTGGAAAGCGGGCAGTGAGCGCAACGCAATTAATGTGAGTTAGCTCACTCATTAGGCACCCC
AGGCTTTACACTTTATGCTTCCGGCTCGTATGTTGTGTGGAATTGTGAGCGGATAACAATTTCACACA
GGAAACAGCTATGACCATGATTACGCCAAGCTATTTAGGTGACACTA

P6:

>P2198 pAAV-CAG-mNeonGreen-Wpre-poly

ACAATGGCAACAACGTTGCGCAAACTATTAACTGGCGAACTACTTACTCTAGCTTCCCGGCAACAATT
AATAGACTGGATGGAGGCGGATAAAGTTGCAGGACCACTTCTGCGCTCGGCCCTTCCGGCTGGCT
GGTTTATTGCTGATAAATCTGGAGCCGGTGAGCGTGGGTCTCGCGGTATCATTGCAGCACTGGGGC
CAGATGGTAAGCCCTCCCGTATCGTAGTTATCTACACGACGGGGAGTCAGGCAACTATGGATGAACG
AAATAGACAGATCGCTGAGATAGGTGCCTCACTGATTAAGCATTGGTAACTGTCAGACCAAGTTTACT
CATATATACTTTAGATTGATTTAAAACTTCATTTTTAATTTAAAAGGATCTAGGTGAAGATCCTTTTTGAT
AATCTCATGACCAAAATCCCTTAACGTGAGTTTTCGTTCCACTGAGCGTCAGACCCCGTAGAAAAGA
TCAAAGGATCTTCTTGAGATCCTTTTTTTCTGCGCGTAATCTGCTGCTTGCAAACAAAAAAACCACCG
CTACCAGCGGTGGTTTGTTTGCCGGATCAAGAGCTACCAACTCTTTTTCCGAAGGTAACTGGCTTCA
GCAGAGCGCAGATACCAAATACTGTCCTTCTAGTGTAGCCGTAGTTAGGCCACCACTTCAAGAACTC
TGTAGCACCGCCTACATACCTCGCTCTGCTAATCCTGTTACCAGTGGCTGCTGCCAGTGGCGATAAG
TCGTGTCTTACCGGGTTGGACTCAAGACGATAGTTACCGGATAAGGCGCAGCGGTCGGGCTGAAC
GGGGGGTTCGTGCACACAGCCCAGCTTGGAGCGAACGACCTACACCGAACTGAGATACCTACAGC
GTGAGCTATGAGAAAGCGCCACGCTTCCCGAAGGGAGAAAGGCGGACAGGTATCCGGTAAGCGGC
AGGGTCGGAACAGGAGAGCGCACGAGGGAGCTTCCAGGGGGAAACGCCTGGTATCTTTATAGTCC
TGTCGGGTTTCGCCACCTCTGACTTGAGCGTCGATTTTTGTGATGCTCGTCAGGGGGGCGGAGCCT
ATGGAAAAACGCCAGCAACGCGGCCTTTTTACGGTTCCTGGCCTTTTGCTGGCCTTTTGCTCACAT
GTCCTGCAGGCAGCTGCGCGCTCGCTCGCTCACTGAGGCCGCCCGGGCGTCGGGCGACCTTTGG
TCGCCCGGCCTCAGTGAGCGAGCGAGCGCGCAGAGAGGGAGTGGCCAACTCCATCACTAGGGGT
TCCTGCGGCCGCACGCGTATCACGAGACTAGCCTCGAGAGCTAGCCCATTGACGTCAATAATGACG
TATGTTCCCATAGTAACGCCAATAGGGACTTTCCATTGACGTCAATGGGTGGAGTATTTACGGTAAAC
TGCCCACTTGGCAGTACATCAAGTGTATCATATGCCAAGTACGCCCCCTATTGACGTCAATGACGGTA
AATGGCCCGCCTGGCATTATGCCCAGTACATGACCTTATGGGACTTTCCTACTTGGCAGTACATCTAC
GTATTAGTCATCGCTATTACCATGGTCGAGGTGAGCCCCACGTTCTGCTTCACTCTCCCCATCTCCC
CCCCCTCCCCACCCCCAATTTTGTATTTATTTATTTTTTAATTATTTTGTGCAGCGATGGGGGCGGGG
GGGGGGGGGGGGGGCGCGCGCCAGGCGGGGCGGGGCGGGGCGAGGGGCGGGGCGGGGCGA
GGCGGAGAGGTGCGGCGGCAGCCAATCAGAGCGGCGCGCTCCGAAAGTTTCCTTTTATGGCGAG
GCGGCGGCGGCGGCGGCCCTATAAAAAGCGAAGCGCGCGGCGGGCGGGAGTCGCTGCGCGCTG
CCTTCGCCCCGTGCCCCGCTCCGCCGCCGCCTCGCGCCGCCCGCCCCGGCTCTGACTGACCGC
GTTACTCCCACAGGTGAGCGGGCGGGACGGCCCTTCTCCTCCGGGCTGTAATTAGCGCTTGGTTTA
ATGACGGCTTGTTTCTTTTCTGTGGCTGCGTGAAAGCCTTGAGGGGCTCCGGGAGGGCCCTTTGT
GCGGGGGGAGCGGCTCGGGGCTGTCCGCGGGGGGACGGCTGCCTTCGGGGGGGACGGGGCAG
GGCGGGGTTCGGCTTCTGGCGTGTGACCGGCGGCTCTAGAGCCTCTGCTAACCATGTTCATGCCT
TCTTCTTTTTCCTACAGCTCCTGGGCAACGTGCTGGTTATTGTGCTGTCTCATCATTTTGGCAAAGAA
TTGGATCCGCGATCGCCATGGTGAGCAAGGGCGAGGAGGATAACATGGCCTCTCTCCCAGCGACA
CATGAGTTACACATCTTTGGCTCCATCAACGGTGTGGACTTTGACATGGTGGGTCAGGGCACCGGC
AATCCAAATGATGGTTATGAGGAGTTAAACCTGAAGTCCACCAAGGGTGACCTCCAGTTCTCCCCCT
GGATTCTGGTCCCTCATATCGGGTATGGCTTCCATCAGTACCTGCCCTACCCTGACGGGATGTCGCC
TTTCCAGGCCGCCATGGTAGATGGCTCCGGATACCAAGTCCATCGCACAATGCAGTTTGAAGATGGT
GCCTCCCTTACTGTTAACTACCGCTACACCTACGAGGGAAGCCACATCAAAGGAGAGGCCCAGGTG
AAGGGGACTGGTTTCCCTGCTGACGGTCCTGTGATGACCAACTCGCTGACCGCTGCGGACTGGTG
CAGGTCGAAGAAGACTTACCCCAACGACAAAACCATCATCAGTACCTTTAAGTGGAGTTACACCACT

GGAAATGGCAAGCGCTACCGGAGCACTGCGCGGACCACCTACACCTTTGCCAAGCCAATGGCGGC
TAACTATCTGAAGAACCAGCCGATGTACGTGTTCCGTAAGACGGAGCTCAAGCACTCCAAGACCGA
GCTCAACTTCAAGGAGTGGCAAAAGGCCTTTACCGATGTGATGGGCATGGACGAGCTGTACAAGTA
AGTTTAAACGTCGACAATCAACCTCTGGATTACAAAATTTGTGAAAGATTGACTGGTATTCTTAACTAT
GTTGCTCCTTTTACGCTATGTGGATACGCTGCTTTAATGCCTTTGTATCATGCTATTGCTTCCCGTATG
GCTTTCATTTTCTCCTCCTTGTATAAATCCTGGTTGCTGTCTCTTTATGAGGAGTTGTGGCCCGTTGT
CAGGCAACGTGGCGTGGTGTGCACTGTGTTTGCTGACGCAACCCCCACTGGTTGGGGCATTGCCA
CCACCTGTCAGCTCCTTTCCGGGACTTTCGCTTTCCCCCTCCCTATTGCCACGGCGGAACTCATCG
CCGCCTGCCTTGCCCGCTGCTGGACAGGGGCTCGGCTGTTGGGCACTGACAATTCCGTGGTGTTG
TCGGGGAAGCTGACGTCCTTTCCATGGCTGCTCGCCTGTGTTGCCACCTGGATTCTGCGCGGGAC
GTCCTTCTGCTACGTCCCTTCGGCCCTCAATCCAGCGGACCTTCCTTCCCGCGGCCTGCTGCCGG
CTCTGCGGCCTCTTCCGCGTCTTCGCCTTCGCCCTCAGACGAGTCGGATCTCCCTTTGGGCGGAT
CCTCTAGAGTCGACCTGCAGAAGCTTGCCTCGAGCAGCGCTGCTCGAGAGATCTACGGGTGGCAT
CCCTGTGACCCCTCCCCAGTGCCTCTCCTGGCCCTGGAAGTTGCCACTCCAGTGCCCACCAGCCT
TGTCCTAATAAAATTAAGTTGCATCATTTTGTCTGACTAGGTGTCCTTCTATAATATTATGGGGTGGAG
GGGGGTGGTATGGAGCAAGGGGCAAGTTGGGAAGACAACCTGTAGGGCCTGCGGGGTCTATTGG
GAACCAAGCTGGAGTGCAGTGGCACAATCTTGGCTCACTGCAATCTCCGCCTCCTGGGTTCAAGC
GATTCTCCTGCCTCAGCCTCCCGAGTTGTTGGGATTCCAGGCATGCATGACCAGGCTCAGCTAATTT
TTGTTTTTTTGGTAGAGACGGGGTTTCACCATATTGGCCAGGCTGGTCTCCAACTCCTAATCTCAGG
TGATCTACCCACCTTGGCCTCCCAAATTGCTGGGATTACAGGCGTGAACCACTGCTCCCTTCCCTGT
CCTTCTGATTTTGTAGGTAACCACGTGCGGACCGAGCGGCCGCAGGAACCCCTAGTGATGGAGTTG
GCCACTCCCTCTCTGCGCGCTCGCTCGCTCACTGAGGCCGGGCGACCAAAGGTCGCCCGACGCC
CGGGCTTTGCCCGGGCGGCCTCAGTGAGCGAGCGAGCGCGCAGCTGCCTGCAGGGGCGCCTGA
TGCGGTATTTTCTCCTTACGCATCTGTGCGGTATTTCACACCGCATACGTCAAAGCAACCATAGTACG
CGCCCTGTAGCGGCGCATTAAGCGCGGCGGGTGTGGTGGTTACGCGCAGCGTGACCGCTACACTT
GCCAGCGCCCTAGCGCCCGCTCCTTTCGCTTTCTTCCCTTCCTTTCTCGCCACGTTCGCCGGCTTT
CCCCGTCAAGCTCTAAATCGGGGGCTCCCTTTAGGGTTCCGATTTAGTGCTTTACGGCACCTCGAC
CCCAAAAAACTTGATTTGGGTGATGGTTCACGTAGTGGGCCATCGCCCTGATAGACGGTTTTTCGCC
CTTTGACGTTGGAGTCCACGTTCTTTAATAGTGGACTCTTGTTCCAAACTGGAACAACACTCAACCC
TATCTCGGGCTATTCTTTTGATTTATAAGGGATTTTGCCGATTTCGGCCTATTGGTTAAAAAATGAGCT
GATTTAACAAAAATTTAACGCGAATTTTAACAAAATATTAACGTTTACAATTTTATGGTGCACTCTCAGT
ACAATCTGCTCTGATGCCGCATAGTTAAGCCAGCCCCGACACCCGCCAACACCCGCTGACGCGCC
CTGACGGGCTTGTCTGCTCCCGGCATCCGCTTACAGACAAGCTGTGACCGTCTCCGGGAGCTGCA
TGTGTCAGAGGTTTTCACCGTCATCACCGAAACGCGCGAGACGAAAGGGCCTCGTGATACGCCTAT
TTTTATAGGTTAATGTCATGATAATAATGGTTTCTTAGACGTCAGGTGGCACTTTTCGGGGAAATGTGC
GCGGAACCCCTATTTGTTTATTTTTCTAAATACATTCAAATATGTATCCGCTCATGAGACAATAACCCTG
ATAAATGCTTCAATAATATTGAAAAAGGAAGAGTATGAGTATTCAACATTTCCGTGTCGCCCTTATTCC
CTTTTTTGCGGCATTTTGCCTTCCTGTTTTTGCTCACCCAGAAACGCTGGTGAAAGTAAAAGATGCT
GAAGATCAGTTGGGTGCACGAGTGGGTTACATCGAACTGGATCTCAACAGCGGTAAGATCCTTGAG
AGTTTTCGCCCCGAAGAACGTTTTCCAATGATGAGCACTTTTAAAGTTCTGCTATGTGGCGCGGTATT
ATCCCGTATTGACGCCGGGCAAGAGCAACTCGGTCGCCGCATACACTATTCTCAGAATGACTTGGTT
GAGTACTCACCAGTCACAGAAAAGCATCTTACGGATGGCATGACAGTAAGAGAATTATGCAGTGCTG
CCATAACCATGAGTGATAACACTGCGGCCAACTTACTTCTGACAACGATCGGAGGACCGAAGGAGC
TAACCGCTTTTTTGCACAACATGGGGGATCATGTAACTCGCCTTGATCGTTGGGAACCGGAGCTGAA

TGAAGCCATACCAAACGACGAGCGTGACACCACGATGCCTGTAG

P7:

>P2202 pAAV-CAG-mNeonGreen-Wpre-polyA

TTTCTGGGTGAGCAAAAACAGGAAGGCAAAATGCCGCAAAAAAGGGAATAAGGGCGACACGGAAAT
GTTGAATACTCATACTCTTCCTTTTTCAATATTATTGAAGCATTTATCAGGGTTATTGTCTCATGAGCGG
ATACATATTTGAATGTATTTAGAAAAATAAACAAATAGGGGTTCCGCGCACATTTCCCCGAAAAGTGCC
ACCTGACGTCTAAGAAACCATTATTATCATGACATTAACCTATAAAAATAGGCGTATCACGAGGCCCTT
TCGTCTCGCGCGTTTCGGTGATGACGGTGAAAACCTCTGACACATGCAGCTCCCGGAGACGGTCA
CAGCTTGTCTGTAAGCGGATGCCGGGAGCAGACAAGCCCGTCAGGGCGCGTCAGCGGGTGTTGG
CGGGTGTCGGGGCTGGCTTAACTATGCGGCATCAGAGCAGATTGTACTGAGAGTGCACCATAAAAT
TGTAAACGTTAATATTTTGTTAAAATTCGCGTTAAATTTTTGTTAAATCAGCTCATTTTTTAACCAATAGA
CCGAAATCGGCAAAATCCCTTATAAATCAAAAGAATAGCCCGAGATAGAGTTGAGTGTTGTTCCAGTT
TGGAACAAGAGTCCACTATTAAAGAACGTGGACTCCAACGTCAAAGGGCGAAAAACCGTCTATCAG
GGCGATGGCCCACTACGTGAACCATCACCCAAATCAAGTTTTTTGGGGTCGAGGTGCCGTAAAGCA
CTAAATCGGAACCCTAAAGGGAGCCCCCGATTTAGAGCTTGACGGGGAAAGCCGGCGAACGTGGC
GAGAAAGGAAGGGAAGAAAGCGAAAGGAGCGGGCGCTAAGGCGCTGGCAAGTGTAGCGGTCACG
CTGCGCGTAACCACCACACCCGCCGCGCTTAATGCGCCGCTACAGGGCGCGTACTATGGTTGCTTT
GACGTATGCGGTGTGAAATACCGCACAGATGCGTAAGGAGAAAATACCGCATCAGGCGCCCCTGCA
GGCAGCTGCGCGCTCGCTCGCTCACTGAGGCCGCCCGGGCAAAGCCCGGGCGTCGGGCGACCT
TTGGTCGCCCGGCCTCAGTGAGCGAGCGAGCGCGCAGAGAGGGAGTGGCCAACTCCATCACTAG
GGGTTCCTGCGGCCGCTCGGTCCGCACGTGCCCCGAAAAGTGCCACCTGGTGGTGCAGATGAAC
TCCATGGGGTTCCCGCATAGCGGCTCACTCTTCCCTCTCTCCCCTCCCCTGCAGTTCGAGTATGGC
GGCCAGGGCTCCGACCCTGCCGATGTGGCCATCCAGCTGACCTTCCTGCGCCTGATGTCCACCGA
GGCCTCCCAGAACATCACCTACCACTGCAAGAACAGCGTGGCCTACATGGACCAGCAGACTGGCA
ACCTCAAGAAGGCCCTGCTCCTCCAGGGCTCCAACGAGATCGAGATCCGCGCCGAGGGCAACAGC
CGCTTCACCTACAGCGTCACTGTCGATGGCTGCACGGTGAGTGCCCAGAATCCCCAGGCAGGGCC
CCACCTCTCCGGCCTTGGGCTTTTTGGCCAGGCCATAGTGCCCTCTCTCCATCACTCCCACGTGGT
AATGCCCCCTCCCGTTGTCTCCACCCCACCCCAGAGTCACACCGGAGCCTGGGGCAAGACAGTGA
TTGAATACAAAACCACCAAGACCTCCCGCCTGCCCATCATCGATGTGGCCCCCTTGGACGTTGGTG
CCCCAGACCAGGAATTCGGCTTCGACGTTGGCCCTGTGGATCCGCCCAAAGGGAGATCCGACTCG
TCTGAGGGCGAAGGCGAGACGCGGAAGAGGCCGCAGAGCCGGCAGCAGGCCGCGGGAAGGAAG
GTCCGCTGGATTGAGGGCCGAAGGGACGTAGCAGAAGGACGTCCCGCGCAGAATCCAGGTGGCA
ACACGGGCGAGCAGCCATGGAAAGGACGTCAGCTTCCCCGACAACACCACGGAATTGTCAGTGCC
CAACAGCCGAGCCCCTGTCCAGCAGCGGGCAAGGCAGGCGGCGATGAGTTCCGCCGTGGCAATA
GGGAGGGGGAAAGCGAAAGTCCCGGAAAGGAGCTGACAGGTGGTGGCAATGCCCCAACCAGTGG
GGGTTGCGTCAGCAAACACAGTGCACACCACGCCACGTTGCCTGACAACGGGCCACAACTCCTCA
TAAAGAGACAGCAACCAGGATTTATACAAGGAGGAGAAAATGAAAGCCATACGGGAAGCAATAGCAT
GATACAAAGGCATTAAAGCAGCGTATCCACATAGCGTAAAAGGAGCAACATAGTTAAGAATACCAGTC
AATCTTTCACAAATTTTGTAATCCAGAGGTTGATTGTCGACGTTTAAACTTACTTGTACAGCTCGTCCA
TGCCCATCACATCGGTAAAGGCCTTTTGCCACTCCTTGAAGTTGAGCTCGGTCTTGGAGTGCTTGA
GCTCCGTCTTACGGAACACGTACATCGGCTGGTTCTTCAGATAGTTAGCCGCCATTGGCTTGGCAAA
GGTGTAGGTGGTCCGCGCAGTGCTCCGGTAGCGCTTGCCATTTCCAGTGGTGTAACTCCACTTAAA
GGTACTGATGATGGTTTTGTCGTTGGGGTAAGTCTTCTTCGACCTGCACCAGTCCGCAGCGGTCAG
CGAGTTGGTCATCACAGGACCGTCAGCAGGGAAACCAGTCCCCTTCACCTGGGCCTCTCCTTTGAT
GTGGCTTCCCTCGTAGGTGTAGCGGTAGTTAACAGTAAGGGAGGCACCATCTTCAAACTGCATTGT

GCGATGGACTTGGTATCCGGAGCCATCTACCATGGCGGCCTGGAAAGGCGACATCCCGTCAGGGT
AGGGCAGGTACTGATGGAAGCCATACCCGATATGAGGGACCAGAATCCAGGGGGAGAACTGGAGG
TCACCCTTGGTGGACTTCAGGTTTAACTCCTCATAACCATCATTTGGATTGCCGGTGCCCTGACCCA
CCATGTCAAAGTCCACACCGTTGATGGAGCCAAAGATGTGTAACTCATGTGTCGCTGGGAGAGAGG
CCATGTTATCCTCCTCGCCCTTGCTCACCATCGGGCCAGGATTCTCCTCCACGTCGCCACAAGTCA
GCAGAGATCCTCGGCCCTCGCCGCTGCCACGCGTAGGTCGAGTGAGCTGTGTGTAGACGGGTTGT
TCCCAGTGCTGGGGGCTGTGGGTCTGCGGGATGGAAGGGACCCCAGAGGTGTCGGCGATGGGG
GTGTACATGGGGCGCTGAGCGGGGTTCATGTAGGTGAAGGTGGAGTAGAGGCCGGTGCCCTGGC
CTGCCGCGTGGCTGTAGTAGGAGCTGGAGTTCTGGTGGTCGGTGTAGTCGTACTGTGAGCGGGTG
ATGGGCGGGTAGGAGGGGCTGTAGTGTGGGAGGTTGAAGGGGCTGTAGGCGATCTGTTGGGGCG
AGTGCTGCTGCTGCTCGCTGTAGTGGCTGGGGCTCAGCTGCTCCGTCTTGATGTGCGTTCGCTGG
GACTGGCCCGGCTCGCTGCTCAGCGTGGTCAGCGTGTGCGCCTGTGGCTGCTGCGGGGGTGCC
GCCGGCTGCTGTGGGGGCTGCTGCGGGGGTGGCGGCGGCGCCTGCTGCTTGGACATCCACACG
TGGCCCGCGCTCGCCGGGGTGGCCGCGGTGCTGCTGATGCCGTAGCTGCCCGTGTAGGTGACCT
GGCCGTGCGTGGCCGGCACCCCCGGGTGGCCGTTGGGCGGCAGGTACTGGTCAAACTCGTTGAC
ATCGAAGGTCTCGATGTTGGAGATGACGTCGCTGCTCAGCTCGCCGATGTCCACGTCGCGGAAGT
CGATAGGGGGCTGTCTGCCCCCCTCTGGCAAGGGGCGCCCCTCTCGCTTCAGGTCAGCCTTGCCC
GGCTGCACGTCGGTTTTGGGGGTGGTGGGTGGGGTCGGTGGGCCCTGGGATTGCCCCGAGTGCT
CGCCGGGGGAGTGCACCTCGCTCATGCCGGAGGAGGAGTGTGGCGAGTCGGCCTGCAGCGCCTT
GAAGATGGCGTTGGGGGAGATGTGCGTCTGCTCCGTGGCCTCCTCTGCCTCCGCCTGCCCGTTCT
TCACCGACTTCCTCCGCCGCGGCTGGTACTTGTAATCCGGGTGGTCCTTCTTGTGCTGCACGCGCA
GCCGCTCCGCCTCCTCCACGAAGGGCCGCTTCTCGCTCTCGTTCAGAAGTCTCCAGAGCTTGCCC
AGCGTCTTGCTGAGCTCGGCGTTGTGCAAGTGCGGGTACTGGTCCGCGAGCTTCCTGCGCGCCG
CCTGCGCCCACACCATGAAGGCGTTCATGGGCCGCTTGACGTGCGGCTTGTTCTTGCTGGAGCCG
TTGACGCGCACCGGCATGGGCACCAGCGTCCAGTCGTAGCCTTTGAGCACCTGGCTGACCGCCTC
GCGGATGCACACGGGGAACTTGTCCTCCTCGCTCTCCTTCTTCAGATCGGGCTCGCCCTTGGGGA
ACGTGTTCTCCTGGGGCCGCGTGTTCTCGGTGTCCGAGCCGGAGCCCGACGGGCAGGGCGAGCC
CGCGGAGTCCTCGGACATGGTGGGGCTGGGGGCGCCGGACAGGCCCTTCTCCTGCTCGTCGGTC
ATCTTCATGAAGGGGTCCAGGAGATTCATGCTAGCGGGGCCAGGGTTTTCTTCGACGTCGCCGGCC
TGTTTCAGCAGAGAGAAGTTTGTAGCGCCGGAGCCACTCCCTCCATCCCAACCTGGCTCCCTCCCA
CCCAACCAACTTTCCCCCCAACCCGGAAACAGACAAGCAACCCAAACTGAACCCCCTCAAAAGCCA
AAAAATGGGAGACAATTTCACATGGACTTTGGAAAATATTTTTTTCCTTTGCATTCATCTCTCAAACTT
AGTTTTTATCTTTGACCAACCGAACATGACCAAAAACCAAAAGTGCATTCAACCTTACCAAAAAAAAA
AAAAAAAAAAAAAAAAAGAATAAATAAATAACTTTTTAAAAAAGGAAGCTTGGTCCACTTGCTTGAAGA
CCCATGCGGGGGTAAGTCCCTTTCTGCCCGTTGGGCTTATGAAACCCCAATGCTGCCCTTTCTGCT
CCTTTCTCCACACCCCCCTTGGGGCCTCCCCTCCACTCCTTCCCAAATCTGTCTCCCCAGAAGACA
CAGGAAACAATGTATTGTCTGCCCAGCAATCAAAGGCAATGCTCAAACACCCAAGTGGCCCCCACC
CTCAGCCCGCTCCTGCCCGCCCAGCACCCCCAGGCCCTGGGGGACCCCATGGAGTTCATCTGCA
CCACCTGTGTGAAATTGTTATCCGCTCACGCGTGCGGCCGCAGGAACCCCTAGTGATGGAGTTGGC
CACTCCCTCTCTGCGCGCTCGCTCGCTCACTGAGGCCGGGCGACCAAAGGTCGCCCGACGCCCG
GGCGGCCTCAGTGAGCGAGCGAGCGCGCAGCTGCCTGCAGGACATGTGAGCAAAAGGCCAGCAA
AAGGCCAGGAACCGTAAAAAGGCCGCGTTGCTGGCGTTTTTCCATAGGCTCCGCCCCCCTGACGA
GCATCACAAAAATCGACGCTCAAGTCAGAGGTGGCGAAACCCGACAGGACTATAAAGATACCAGGC
GTTTCCCCCTGGAAGCTCCCTCGTGCGCTCTCCTGTTCCGACCCTGCCGCTTACCGGATACCTGTC

CGCCTTTCTCCCTTCGGGAAGCGTGGCGCTTTCTCATAGCTCACGCTGTAGGTATCTCAGTTCGGT
GTAGGTCGTTCGCTCCAAGCTGGGCTGTGTGCACGAACCCCCCGTTCAGCCCGACCGCTGCGCCT
TATCCGGTAACTATCGTCTTGAGTCCAACCCGGTAAGACACGACTTATCGCCACTGGCAGCAGCCAC
TGGTAACAGGATTAGCAGAGCGAGGTATGTAGGCGGTGCTACAGAGTTCTTGAAGTGGTGGCCTAA
CTACGGCTACACTAGAAGAACAGTATTTGGTATCTGCGCTCTGCTGAAGCCAGTTACCTTCGGAAAA
AGAGTTGGTAGCTCTTGATCCGGCAAACAAACCACCGCTGGTAGCGGTGGTTTTTTTGTTTGCAAG
CAGCAGATTACGCGCAGAAAAAAAGGATCTCAAGAAGATCCTTTGATCTTTTCTACGGGGTCTGACG
CTCAGTGGAACGAAAACTCACGTTAAGGGATTTTGGTCATGAGATTATCAAAAAGGATCTTCACCTAG
ATCCTTTTAAATTAAAAATGAAGTTTTAAATCAATCTAAAGTATATATGAGTAAACTTGGTCTGACAGTTA
CCAATGCTTAATCAGTGAGGCACCTATCTCAGCGATCTGTCTATTTCGTTCATCCATAGTTGCCTGAC
TCCCCGTCGTGTAGATAACTACGATACGGGAGGGCTTACCATCTGGCCCCAGTGCTGCAATGATACC
GCGAGACCCACGCTCACCGGCTCCAGATTTATCAGCAATAAACCAGCCAGCCGGAAGGGCCGAGC
GCAGAAGTGGTCCTGCAACTTTATCCGCCTCCATCCAGTCTATTAATTGTTGCCGGGAAGCTAGAGT
AAGTAGTTCGCCAGTTAATAGTTTGCGCAACGTTGTTGCCATTGCTACAGGCATCGTGGTGTCACGC
TCGTCGTTTGGTATGGCTTCATTCAGCTCCGGTTCCCAACGATCAAGGCGAGTTACATGATCCCCCA
TGTTGTGCAAAAAAGCGGTTAGCTCCTTCGGTCCTCCGATCGTTGTCAGAAGTAAGTTGGCCGCAG
TGTTATCACTCATGGTTATGGCAGCACTGCATAATTCTCTTACTGTCATGCCATCCGTAAGATGCTTTT
CTGTGACTGGTGAGTACTCAACCAAGTCATTCTGAGAATAGTGTATGCGGCGACCGAGTTGCTCTTG
CCCGGCGTCAATACGGGATAATACCGCGCCACATAGCAGAACTTTAAAAGTGCTCATCATTGGAAAA
CGTTCTTCGGGGCGAAAACTCTCAAGGATCTTACCGCTGTTGAGATCCAGTTCGATGTAACCCACTC
GTGCACCCAACTGATCTTCAGCATCTTTTACTTTCACCAGCG

P8：

>P2207 pRSC-COL1A1-raSOX9-T2A-Puro-Wpre

CTTCAAGAACTCTGTAGCACCGCCTACATACCTCGCTCTGCTAATCCTGTTACCAGTGGCTGCTGCC
AGTGGCGATAAGTCGTGTCTTACCGGGTTGGACTCAAGACGATAGTTACCGGATAAGGCGCAGCGG
TCGGGCTGAACGGGGGGTTCGTGCACACAGCCCAGCTTGGAGCGAACGACCTACACCGAACTGA
GATACCTACAGCGTGAGCTATGAGAAAGCGCCACGCTTCCCGAAGGGAGAAAGGCGGACAGGTAT
CCGGTAAGCGGCAGGGTCGGAACAGGAGAGCGCACGAGGGAGCTTCCAGGGGGAAACGCCTGGT
ATCTTTATAGTCCTGTCGGGTTTCGCCACCTCTGACTTGAGCGTCGATTTTTGTGATGCTCGTCAGG
GGGGCGGAGCCTATGGAAAAACGCCAGCAACGCGGCCTTTTTACGGTTCCTGGCCTTTTGCTGGC
CTTTTGCTCACATGTTCTTTCCTGCGTTATCCCCTGATTCTGTGGATAACCGTATTACCGCCTTTGAG
TGAGCTGATACCGCTCGCCGCAGCCGAACGACCGAGCGCAGCGAGTCAGTGAGCGAGGAAGCGG
AAGAGCGCCCAATACGCAAACCGCCTCTCCCCGCGCGTTGGCCGATTCATTAATGCAGCTGGCACG
ACAGGTTTCCCGACTGGAAAGCGGGCAGTGAGCGCAACGCAATTAAGGTGAGTTAGCTCACTCATT
AGGCACCCCAGGCTTTACACTTTATGCTTCCGGCTCGTATGTTGTGTGGAATTGTGAGCGGATAACA
ATTTCACACAGGAAACAGCTATGACCATGATTACGCCAAGCGCGCAATTAACCCTCACTAAAGGGAA
CAAAAGCTGGAGCTGCAAGCTTAATGTAGTCTTATGCAATACTCTTGTAGTCTTGCAACATGGTAACG
ATGAGTTAGCAACATGCCTTACAAGGAGAGAAAAAGCACCGTGCATGCCGATTGGTGGAAGTAAGG
TGGTACGATCGTGCCTTATTAGGAAGGCAACAGACGGGTCTGACATGGATTGGACGAACCACTGAA
TTGCCGCATTGCAGAGATATTGTATTTAAGTGCCTAGCTCGATACATAAACGGGTCTCTCTGGTTAGA
CCAGATCTGAGCCTGGGAGCTCTCTGGCTAACTAGGGAACCCACTGCTTAAGCCTCAATAAAGCTT
GCCTTGAGTGCTTCAAGTAGTGTGTGCCCGTCTGTTGTGTGACTCTGGTAACTAGAGATCCCTCAG
ACCCTTTTAGTCAGTGTGGAAAATCTCTAGCAGTGGCGCCCGAACAGGGACTTGAAAGCGAAAGGG
AAACCAGAGGAGCTCTCTCGACGCAGGACTCGGCTTGCTGAAGCGCGCACGGCAAGAGGCGAGG
GGCGGCGACTGGTGAGTACGCCAAAAATTTTGACTAGCGGAGGCTAGAAGGAGAGAGATGGGTGC
GAGAGCGTCAGTATTAAGCGGGGGAGAATTAGATCGCGATGGGAAAAAATTCGGTTAAGGCCAGGG
GGAAAGAAAAAATATAAATTAAAACATATAGTATGGGCAAGCAGGGAGCTAGAACGATTCGCAGTTAA
TCCTGGCCTGTTAGAAACATCAGAAGGCTGTAGACAAATACTGGGACAGCTACAACCATCCCTTCAG
ACAGGATCAGAAGAACTTAGATCATTATATAATACAGTAGCAACCCTCTATTGTGTGCATCAAAGGATA
GAGATAAAAGACACCAAGGAAGCTTTAGACAAGATAGAGGAAGAGCAAAACAAAAGTAAGACCACC
GCACAGCAAGCGGCCGCTGATCTTCAGACCTGGAGGAGGAGATATGAGGGACAATTGGAGAAGTG
AATTATATAAATATAAAGTAGTAAAAATTGAACCATTAGGAGTAGCACCCACCAAGGCAAAGAGAAGAG
TGGTGCAGAGAGAAAAAAGAGCAGTGGGAATAGGAGCTTTGTTCCTTGGGTTCTTGGGAGCAGCA
GGAAGCACTATGGGCGCAGCGTCAATGACGCTGACGGTACAGGCCAGACAATTATTGTCTGGTATA
GTGCAGCAGCAGAACAATTTGCTGAGGGCTATTGAGGCGCAACAGCATCTGTTGCAACTCACAGTC
TGGGGCATCAAGCAGCTCCAGGCAAGAATCCTGGCTGTGGAAAGATACCTAAAGGATCAACAGCTC
CTGGGGATTTGGGGTTGCTCTGGAAAACTCATTTGCACCACTGCTGTGCCTTGGAATGCTAGTTGG
AGTAATAAATCTCTGGAACAGATTTGGAATCACACGACCTGGATGGAGTGGGACAGAGAAATTAACA
ATTACACAAGCTTAATACACTCCTTAATTGAAGAATCGCAAAACCAGCAAGAAAAGAATGAACAAGAA
TTATTGGAATTAGATAAATGGGCAAGTTTGTGGAATTGGTTTAACATAACAAATTGGCTGTGGTATATAA
AATTATTCATAATGATAGTAGGAGGCTTGGTAGGTTTAAGAATAGTTTTTGCTGTACTTTCTATAGTGAA
TAGAGTTAGGCAGGGATATTCACCATTATCGTTTCAGACCCACCTCCCAACCCCGAGGGGACCCGA
CAGGCCCGAAGGAATAGAAGAAGAAGGTGGAGAGAGAGACAGAGACAGATCCATTCGATTAGTGAA
CGGATCTCGACGGTATCGGTTAACTTTTAAAAGAAAAGGGGGGATTGGGGGGTACAGTGCAGGGGA
AAGAATAGTAGACATAATAGCAACAGACATACAAACTAAAGAATTACAAAAACAAATTACAAAAATTCAA

AATTTTATCGATCACGAGACTAGCCATATGTATTCCTAAGCTCCATCCCCTAGCCACACCAGAAGACA
CCCCCAAACAGGCACATCTTTTTAATTCCCAGCTTCCTCTGTTTTGGAGAGGTCCTCAGCATGCCTC
TTTATGCCCCTCCCTTAGCTCTTGCCAGGATATCAGAGGGTGACTGGGGCACAGCCAGGAGGACCC
CCTCCCCAACACCCCCAACCCTTCCACCTTTGGAAGTCTCCCCACCCAGCTCCCCAGTTCCCCAGT
TCCACTTCTTCTAGATTGGAGGTCCCAGGAAGAGAGCAGAGGGGCACCCCTACCCACTGGTTAGCC
CACGCCATTCTGAGGACCCAGCTGCACCCCTACCACAGCACCTCTGGCCCAGGCTGGGCTGGGG
GGCTGGGGAGGCAGAGCTGCGAAGAGGGGAGATGTGGGGTGGACTCCCTTCCCTCCTCCTCCCC
CTCTCCATTCCAACTCCCAAATTGGGGGCCGGGCCAGGCAGCTCTGATTGGCTGGGGCACGGGCG
GCCGGCTCCCCCTCTCCGAGGGGCAGGGTTCCTCCCTGCTCTCCATCAGGACAGTATAAAAGGGG
CCCGGGCCAGTCGTCGGAGCAGACGGGAGTTTCTCCTCGGGGTCGGAGCAGGAGGCACGCGGA
GTGTGAGGCCACGCATGAGCGGACGCTAACCCCCTCCCCAGCCACAAAGAGTCTACATGTCTAGG
GTCTAGACGCGATCGCCTGAATCTCCTGGACCCCTTCATGAAGATGACCGACGAGCAGGAGAAGG
GCCTGTCCGGCGCCCCCAGCCCCACCATGTCCGAGGACTCGGCGGGCTCGCCCTGCCCGTCGG
GCTCCGGCTCCGACACCGAGAATACACGGCCCCAGGAGAACACGTTCCCCAAGGGCGAGCCGGA
CCTCAAGAAGGAGAGCGAAGAGGACAAGTTCCCCGTGTGCATCCGCGAGGCGGTCAGCCAGGTG
CTCAAGGGCTACGACTGGACGCTGGTGCCCATGCCCGTGCGCGTCAACGGCTCCAGCAAGAACAA
GCCCCACGTCAAGCGGCCCATGAACGCCTTCATGGTGTGGGCGCAGGCGGCGCGCAGGAAGCTG
GCCGACCAGTACCCGCACCTGCACAACGCCGAGCTCAGCAAGACCCTCGGGAAGCTCTGGAGAC
TGCTGAACGAGAGCGAGAAGCGGCCCTTCGTGGAGGAGGCGGAGCGGCTGCGTGTGCAGCACAA
GAAAGACCACCCGGACTACAAGTACCAGCCGCGGCGGAGGAAGTCGGTGAAGAATGGGCAGGCG
GAGGCCGAGGAGGCCACGGAGCAGACGCACATCTCGCCCAACGCCATCTTCAAGGCGCTGCAGG
CCGACTCGCCGCACTCCTCGTCCGGCATGAGCGAGGTCCATTCCCCCGGCGAGCACTCGGGACAA
TCCCAGGGTCCGCCGACCCCACCCACCACTCCCAAGACCGACGTGCAGCCGGGCAAGGCTGACC
TGAAGCGAGAGGGGCGCCCCCTGCCCGAGGGGGGCCGGCAGCCCCCCATCGACTTCCGCGACG
TGGACATCGGCGAGCTGAGCAGCGACGTCATCTCCAACATCGAGACCTTCGACGTCAACGAGTTC
GACCAGTACCTGCCGCCCAATGGCCACCCGGGGGTGCCGGCCACGCACGGCCAGGTCACCTACA
CCGGCAGCTACGGCATCAGCAGCGCCGCCGCGACCCCCGCGGGCGCGGGCCACGTGTGGATGT
CCAAGCAGCAGGCGCCCCCGCCGCCGCCGCCCCCGCAGCAGCAGCCGCCCCCGCAGGCGCCCC
CGCAGGCCCCGCAGGCGCCCCCGCAGCCGCAGCAGGCGCCCCCGCAGCCGCAGCCGGCGCCC
CCGCAGCCGCAGCAGGCGCACACCCTGACCACGCTGAGCAGCGAGCCGGGCCAGGCGCAGCGA
ACGCACATCAAGACGGAGCAGCTGAGCCCCAGCCACTACAGCGAGCCGCAGCACTCGCCGCAGC
AGATCGCCTACAGCCCCTTCAACCTCCCGCACTACAGCCCCTCGTACCCGCCCATCACGCGCTCGC
AGTTCGACTACGCGGAGCACCAGAACTCGGGGTCCTACTACAGCCACGCGGCCAGCCAGGGCTCC
AGCCTCTACTCCACCTTCACCTACATGAGCCCCGCCCAGCGCCCCATGTACACCCCCATCGCCGAC
ACCTCCGGGGTGCCTTCCATCCCGCAGACCCACAGCCCGCAGCACTGGGAGCAGCCGGTGTACA
CACAGCTCACCAGACCTACGCGTGGCAGCGGCGAGGGCCGAGGATCTCTGCTGACTTGTGGCGA
CGTGGAGGAGAATCCTGGCCCGATGACCGAGTACAAGCCCACGGTGCGCCTCGCCACCCGCGAC
GACGTCCCCAGGGCCGTACGCACCCTCGCCGCCGCGTTCGCCGACTACCCCGCCACGCGCCACA
CCGTCGATCCGGACCGCCACATCGAGCGGGTCACCGAGCTGCAAGAACTCTTCCTCACGCGCGTC
GGGCTCGACATCGGCAAGGTGTGGGTCGCGGACGACGGCGCCGCGGTGGCGGTCTGGACCACG
CCGGAGAGCGTCGAAGCGGGGGCGGTGTTCGCCGAGATCGGCCCGCGCATGGCCGAGTTGAGC
GGTTCCCGGCTGGCCGCGCAGCAACAGATGGAAGGCCTCCTGGCGCCGCACCGGCCCAAGGAG
CCCGCGTGGTTCCTGGCCACCGTCGGCGTCTCGCCCGACCACCAGGGCAAGGGTCTGGGCAGCG
CCGTCGTGCTCCCCGGAGTGGAGGCGGCCGAGCGCGCCGGGGTGCCCGCCTTCCTGGAGACCT

CCGCGCCCCGCAACCTCCCCTTCTACGAGCGGCTCGGCTTCACCGTCACCGCCGACGTCGAGGT
GCCCGAAGGACCGCGCACCTGGTGCATGACCCGCAAGCCCGGTGCCTAGGTTTAAACGTCGACAA
TCAACCTCTGGATTACAAAATTTGTGAAAGATTGACTGGTATTCTTAACTATGTTGCTCCTTTTACGCT
ATGTGGATACGCTGCTTTAATGCCTTTGTATCATGCTATTGCTTCCCGTATGGCTTTCATTTTCTCCTC
CTTGTATAAATCCTGGTTGCTGTCTCTTTATGAGGAGTTGTGGCCCGTTGTCAGGCAACGTGGCGTG
GTGTGCACTGTGTTTGCTGACGCAACCCCCACTGGTTGGGGCATTGCCACCACCTGTCAGCTCCTT
TCCGGGACTTTCGCTTTCCCCCTCCCTATTGCCACGGCGGAACTCATCGCCGCCTGCCTTGCCCGC
TGCTGGACAGGGGCTCGGCTGTTGGGCACTGACAATTCCGTGGTGTTGTCGGGGAAGCTGACGTC
CTTTCCATGGCTGCTCGCCTGTGTTGCCACCTGGATTCTGCGCGGGACGTCCTTCTGCTACGTCCC
TTCGGCCCTCAATCCAGCGGACCTTCCTTCCCGCGGCCTGCTGCCGGCTCTGCGGCCTCTTCCGC
GTCTTCGCCTTCGCCCTCAGACGAGTCGGATCTCCCTTTGGGCCGCCTCCCCGCCTGGAATTCGA
GCTCGGTACCTTTAAGACCAATGACTTACAAGGCAGCTGTAGATCTTAGCCACTTTTTAAAAGAAAAG
GGGGGACTGGAAGGGCTAATTCACTCCCAACGAAGACAAGATCTGCTTTTTGCTTGTACTGGGTCT
CTCTGGTTAGACCAGATCTGAGCCTGGGAGCTCTCTGGCTAACTAGGGAACCCACTGCTTAAGCCT
CAATAAAGCTTGCCTTGAGTGCTTCAAGTAGTGTGTGCCCGTCTGTTGTGTGACTCTGGTAACTAGA
GATCCCTCAGACCCTTTTAGTCAGTGTGGAAAATCTCTAGCAGTAGTAGTTCATGTCATCTTATTATTC
AGTATTTATAACTTGCAAAGAAATGAATATCAGAGAGTGAGAGGAACTTGTTTATTGCAGCTTATAATG
GTTACAAATAAAGCAATAGCATCACAAATTTCACAAATAAAGCATTTTTTTCACTGCATTCTAGTTGTG
GTTTGTCCAAACTCATCAATGTATCTTATCATGTCTGGCTCTAGCTATCCCGCCCCTAACTCCGCCCA
TCCCGCCCCTAACTCCGCCCAGTTCCGCCCATTCTCCGCCCCATGGCTGACTAATTTTTTTTATTTAT
GCAGAGGCCGAGGCCGCCTCGGCCTCTGAGCTATTCCAGAAGTAGTGAGGAGGCTTTTTTGGAGG
CCTAGGGACGTACCCAATTCGCCCTATAGTGAGTCGTATTACGCGCGCTCACTGGCCGTCGTTTTAC
AACGTCGTGACTGGGAAAACCCTGGCGTTACCCAACTTAATCGCCTTGCAGCACATCCCCCTTTCG
CCAGCTGGCGTAATAGCGAAGAGGCCCGCACCGATCGCCCTTCCCAACAGTTGCGCAGCCTGAAT
GGCGAATGGGACGCGCCCTGTAGCGGCGCATTAAGCGCGGCGGGTGTGGTGGTTACGCGCAGCG
TGACCGCTACACTTGCCAGCGCCCTAGCGCCCGCTCCTTTCGCTTTCTTCCCTTCCTTTCTCGCCA
CGTTCGCCGGCTTTCCCCGTCAAGCTCTAAATCGGGGGCTCCCTTTAGGGTTCCGATTTAGTGCTTT
ACGGCACCTCGACCCCAAAAAACTTGATTAGGGTGATGGTTCACGTAGTGGGCCATCGCCCTGATA
GACGGTTTTTCGCCCTTTGACGTTGGAGTCCACGTTCTTTAATAGTGGACTCTTGTTCCAAACTGGA
ACAACACTCAACCCTATCTCGGTCTATTCTTTTGATTTATAAGGGATTTTGCCGATTTCGGCCTATTGG
TTAAAAAATGAGCTGATTTAACAAAAATTTAACGCGAATTTTAACAAAATATTAACGCTTACAATTTAGG
TGGCACTTTTCGGGGAAATGTGCGCGGAACCCCTATTTGTTTATTTTTCTAAATACATTCAAATATGTA
TCCGCTCATGAGACAATAACCCTGATAAATGCTTCAATAATATTGAAAAAGGAAGAGTATGAGTATTCA
ACATTTCCGTGTCGCCCTTATTCCCTTTTTTGCGGCATTTTGCCTTCCTGTTTTTGCTCACCCAGAAA
CGCTGGTGAAAGTAAAAGATGCTGAAGATCAGTTGGGTGCACGAGTGGGTTACATCGAACTGGATC
TCAACAGCGGTAAGATCCTTGAGAGTTTTCGCCCCGAAGAACGTTTTCCAATGATGAGCACTTTTAA
AGTTCTGCTATGTGGCGCGGTATTATCCCGTATTGACGCCGGGCAAGAGCAACTCGGTCGCCGCAT
ACACTATTCTCAGAATGACTTGGTTGAGTACTCACCAGTCACAGAAAAGCATCTTACGGATGGCATG
ACAGTAAGAGAATTATGCAGTGCTGCCATAACCATGAGTGATAACACTGCGGCCAACTTACTTCTGA
CAACGATCGGAGGACCGAAGGAGCTAACCGCTTTTTTGCACAACATGGGGGATCATGTAACTCGCC
TTGATCGTTGGGAACCGGAGCTGAATGAAGCCATACCAAACGACGAGCGTGACACCACGATGCCTG
TAGCAATGGCAACAACGTTGCGCAAACTATTAACTGGCGAACTACTTACTCTAGCTTCCCGGCAACA
ATTAATAGACTGGATGGAGGCGGATAAAGTTGCAGGACCACTTCTGCGCTCGGCCCTTCCGGCTGG
CTGGTTTATTGCTGATAAATCTGGAGCCGGTGAGCGTGGGTCTCGCGGTATCATTGCAGCACTGGG

GCCAGATGGTAAGCCCTCCCGTATCGTAGTTATCTACACGACGGGGAGTCAGGCAACTATGGATGAA
CGAAATAGACAGATCGCTGAGATAGGTGCCTCACTGATTAAGCATTGGTAACTGTCAGACCAAGTTT
ACTCATATATACTTTAGATTGATTTAAAACTTCATTTTTAATTTAAAAGGATCTAGGTGAAGATCCTTTTT
GATAATCTCATGACCAAAATCCCTTAACGTGAGTTTTCGTTCCACTGAGCGTCAGACCCCGTAGAAA
AGATCAAAGGATCTTCTTGAGATCCTTTTTTTCTGCGCGTAATCTGCTGCTTGCAAACAAAAAAACCA
CCGCTACCAGCGGTGGTTTGTTTGCCGGATCAAGAGCTACCAACTCTTTTTCCGAAGGTAACTGGC
TTCAGCAGAGCGCAGATACCAAATACTGTCCTTCTAGTGTAGCCGTAGTTAGGCCACCA

P9:

>P2280 pRSC-EF1-raSOX9-T2A-Puro-wpre

GGGCGGAGCCTATGGAAAAACGCCAGCAACGCGGCCTTTTTACGGTTCCTGGCCTTTTGCTGGCC
TTTTGCTCACATGTTCTTTCCTGCGTTATCCCCTGATTCTGTGGATAACCGTATTACCGCCTTTGAGT
GAGCTGATACCGCTCGCCGCAGCCGAACGACCGAGCGCAGCGAGTCAGTGAGCGAGGAAGCGGA
AGAGCGCCCAATACGCAAACCGCCTCTCCCCGCGCGTTGGCCGATTCATTAATGCAGCTGGCACGA
CAGGTTTCCCGACTGGAAAGCGGGCAGTGAGCGCAACGCAATTAAGGTGAGTTAGCTCACTCATTA
GGCACCCCAGGCTTTACACTTTATGCTTCCGGCTCGTATGTTGTGTGGAATTGTGAGCGGATAACAA
TTTCACACAGGAAACAGCTATGACCATGATTACGCCAAGCGCGCAATTAACCCTCACTAAAGGGAAC
AAAAGCTGGAGCTGCAAGCTTAATGTAGTCTTATGCAATACTCTTGTAGTCTTGCAACATGGTAACGA
TGAGTTAGCAACATGCCTTACAAGGAGAGAAAAAGCACCGTGCATGCCGATTGGTGGAAGTAAGGT
GGTACGATCGTGCCTTATTAGGAAGGCAACAGACGGGTCTGACATGGATTGGACGAACCACTGAAT
TGCCGCATTGCAGAGATATTGTATTTAAGTGCCTAGCTCGATACATAAACGGGTCTCTCTGGTTAGAC
CAGATCTGAGCCTGGGAGCTCTCTGGCTAACTAGGGAACCCACTGCTTAAGCCTCAATAAAGCTTG
CCTTGAGTGCTTCAAGTAGTGTGTGCCCGTCTGTTGTGTGACTCTGGTAACTAGAGATCCCTCAGAC
CCTTTTAGTCAGTGTGGAAAATCTCTAGCAGTGGCGCCCGAACAGGGACTTGAAAGCGAAAGGGAA
ACCAGAGGAGCTCTCTCGACGCAGGACTCGGCTTGCTGAAGCGCGCACGGCAAGAGGCGAGGGG
CGGCGACTGGTGAGTACGCCAAAAATTTTGACTAGCGGAGGCTAGAAGGAGAGAGATGGGTGCGA
GAGCGTCAGTATTAAGCGGGGGAGAATTAGATCGCGATGGGAAAAAATTCGGTTAAGGCCAGGGGG
AAAGAAAAAATATAAATTAAAACATATAGTATGGGCAAGCAGGGAGCTAGAACGATTCGCAGTTAATCC
TGGCCTGTTAGAAACATCAGAAGGCTGTAGACAAATACTGGGACAGCTACAACCATCCCTTCAGACA
GGATCAGAAGAACTTAGATCATTATATAATACAGTAGCAACCCTCTATTGTGTGCATCAAAGGATAGAG
ATAAAAGACACCAAGGAAGCTTTAGACAAGATAGAGGAAGAGCAAAACAAAAGTAAGACCACCGCA
CAGCAAGCGGCCGCTGATCTTCAGACCTGGAGGAGGAGATATGAGGGACAATTGGAGAAGTGAATT
ATATAAATATAAAGTAGTAAAAATTGAACCATTAGGAGTAGCACCCACCAAGGCAAAGAGAAGAGTGG
TGCAGAGAGAAAAAAGAGCAGTGGGAATAGGAGCTTTGTTCCTTGGGTTCTTGGGAGCAGCAGGA
AGCACTATGGGCGCAGCGTCAATGACGCTGACGGTACAGGCCAGACAATTATTGTCTGGTATAGTG
CAGCAGCAGAACAATTTGCTGAGGGCTATTGAGGCGCAACAGCATCTGTTGCAACTCACAGTCTGG
GGCATCAAGCAGCTCCAGGCAAGAATCCTGGCTGTGGAAAGATACCTAAAGGATCAACAGCTCCTG
GGGATTTGGGGTTGCTCTGGAAAACTCATTTGCACCACTGCTGTGCCTTGGAATGCTAGTTGGAGTA
ATAAATCTCTGGAACAGATTTGGAATCACACGACCTGGATGGAGTGGGACAGAGAAATTAACAATTA
CACAAGCTTAATACACTCCTTAATTGAAGAATCGCAAAACCAGCAAGAAAAGAATGAACAAGAATTAT
TGGAATTAGATAAATGGGCAAGTTTGTGGAATTGGTTTAACATAACAAATTGGCTGTGGTATATAAAAT
TATTCATAATGATAGTAGGAGGCTTGGTAGGTTTAAGAATAGTTTTTGCTGTACTTTCTATAGTGAATAG
AGTTAGGCAGGGATATTCACCATTATCGTTTCAGACCCACCTCCCAACCCCGAGGGGACCCGACAG
GCCCGAAGGAATAGAAGAAGAAGGTGGAGAGAGAGACAGAGACAGATCCATTCGATTAGTGAACG
GATCTCGACGGTATCGGTTAACTTTTAAAAGAAAAGGGGGGATTGGGGGGTACAGTGCAGGGGAAA
GAATAGTAGACATAATAGCAACAGACATACAAACTAAAGAATTACAAAAACAAATTACAAAAATTCAAAA
TTTTATCGATCACGAGACTAGCAGCTAGCGGCTCCGGTGCCCGTCAGTGGGCAGAGCGCACATCG
CCCACAGTCCCCGAGAAGTTGGGGGGAGGGGTCGGCAATTGAACCGGTGCCTAGAGAAGGTGGC
GCGGGGTAAACTGGGAAAGTGATGTCGTGTACTGGCTCCGCCTTTTTCCCGAGGGTGGGGGAGAA
CCGTATATAAGTGCAGTAGTCGCCGTGAACGTTCTTTTTCGCAACGGGTTTGCCGCCAGAACACAG
GTAAGTGCCGTGTGTGGTTCCCGCGGGCCTGGCCTCTTTACGGGTTATGGCCCTTGCGTGCCTTGA
ATTACTTCCACCTGGCTGCAGTACGTGATTCTTGATCCCGAGCTTCGGGTTGGAAGTGGGTGGGAG

AGTTCGAGGCCTTGCGCTTAAGGAGCCCCTTCGCCTCGTGCTTGAGTTGAGGCCTGGCCTGGGCG
CTGGGGCCGCCGCGTGCGAATCTGGTGGCACCTTCGCGCCTGTCTCGCTGCTTTCGATAAGTCTC
TAGCCATTTAAAATTTTTGATGACCTGCTGCGACGCTTTTTTTCTGGCAAGATAGTCTTGTAAATGCG
GGCCAAGATCTGCACACTGGTATTTCGGTTTTTGGGGCCGCGGGCGGCGACGGGGCCCGTGCGT
CCCAGCGCACATGTTCGGCGAGGCGGGGCCTGCGAGCGCGGCCACCGAGAATCGGACGGGGGT
AGTCTCAAGCTGGCCGGCCTGCTCTGGTGCCTGGCCTCGCGCCGCCGTGTATCGCCCCGCCCTG
GGCGGCAAGGCTGGCCCGGTCGGCACCAGTTGCGTGAGCGGAAAGATGGCCGCTTCCCGGCCCT
GCTGCAGGGAGCTCAAAATGGAGGACGCGGCGCTCGGGAGAGCGGGCGGGTGAGTCACCCACA
CAAAGGAAAAGGGCCTTTCCGTCCTCAGCCGTCGCTTCATGTGACTCCACGGAGTACCGGGCGCC
GTCCAGGCACCTCGATTAGTTCTCGAGCTTTTGGAGTACGTCGTCTTTAGGTTGGGGGGAGGGGTT
TTATGCGATGGAGTTTCCCCACACTGAGTGGGTGGAGACTGAAGTTAGGCCAGCTTGGCACTTGAT
GTAATTCTCCTTGGAATTTGCCCTTTTTGAGTTTGGATCTTGGTTCATTCTCAAGCCTCAGACAGTGG
TTCAAAGTTTTTTTCTTCCATTTCAGGTGTCGTGAGCGATCGCCATGAATCTCCTGGACCCCTTCATG
AAGATGACCGACGAGCAGGAGAAGGGCCTGTCCGGCGCCCCCAGCCCCACCATGTCCGAGGACT
CGGCGGGCTCGCCCTGCCCGTCGGGCTCCGGCTCCGACACCGAGAATACACGGCCCCAGGAGAA
CACGTTCCCCAAGGGCGAGCCGGACCTCAAGAAGGAGAGCGAAGAGGACAAGTTCCCCGTGTGC
ATCCGCGAGGCGGTCAGCCAGGTGCTCAAGGGCTACGACTGGACGCTGGTGCCCATGCCCGTGC
GCGTCAACGGCTCCAGCAAGAACAAGCCCCACGTCAAGCGGCCCATGAACGCCTTCATGGTGTGG
GCGCAGGCGGCGCGCAGGAAGCTGGCCGACCAGTACCCGCACCTGCACAACGCCGAGCTCAGCA
AGACCCTCGGGAAGCTCTGGAGACTGCTGAACGAGAGCGAGAAGCGGCCCTTCGTGGAGGAGGC
GGAGCGGCTGCGTGTGCAGCACAAGAAAGACCACCCGGACTACAAGTACCAGCCGCGGCGGAGG
AAGTCGGTGAAGAATGGGCAGGCGGAGGCCGAGGAGGCCACGGAGCAGACGCACATCTCGCCCA
ACGCCATCTTCAAGGCGCTGCAGGCCGACTCGCCGCACTCCTCGTCCGGCATGAGCGAGGTCCAT
TCCCCCGGCGAGCACTCGGGACAATCCCAGGGTCCGCCGACCCCACCCACCACTCCCAAGACCG
ACGTGCAGCCGGGCAAGGCTGACCTGAAGCGAGAGGGGCGCCCCCTGCCCGAGGGGGGCCGGC
AGCCCCCCATCGACTTCCGCGACGTGGACATCGGCGAGCTGAGCAGCGACGTCATCTCCAACATC
GAGACCTTCGACGTCAACGAGTTCGACCAGTACCTGCCGCCCAATGGCCACCCGGGGGTGCCGG
CCACGCACGGCCAGGTCACCTACACCGGCAGCTACGGCATCAGCAGCGCCGCCGCGACCCCCGC
GGGCGCGGGCCACGTGTGGATGTCCAAGCAGCAGGCGCCCCCGCCGCCGCCGCCCCCGCAGCA
GCAGCCGCCCCCGCAGGCGCCCCCGCAGGCCCCGCAGGCGCCCCCGCAGCCGCAGCAGGCGC
CCCCGCAGCCGCAGCCGGCGCCCCCGCAGCCGCAGCAGGCGCACACCCTGACCACGCTGAGCA
GCGAGCCGGGCCAGGCGCAGCGAACGCACATCAAGACGGAGCAGCTGAGCCCCAGCCACTACAG
CGAGCCGCAGCACTCGCCGCAGCAGATCGCCTACAGCCCCTTCAACCTCCCGCACTACAGCCCCT
CGTACCCGCCCATCACGCGCTCGCAGTTCGACTACGCGGAGCACCAGAACTCGGGGTCCTACTAC
AGCCACGCGGCCAGCCAGGGCTCCAGCCTCTACTCCACCTTCACCTACATGAGCCCCGCCCAGCG
CCCCATGTACACCCCCATCGCCGACACCTCCGGGGTGCCTTCCATCCCGCAGACCCACAGCCCGC
AGCACTGGGAGCAGCCGGTGTACACACAGCTCACCAGACCTACGCGTGGCAGCGGCGAGGGCCG
AGGATCTCTGCTGACTTGTGGCGACGTGGAGGAGAATCCTGGCCCGATGACCGAGTACAAGCCCA
CGGTGCGCCTCGCCACCCGCGACGACGTCCCCAGGGCCGTACGCACCCTCGCCGCCGCGTTCGC
CGACTACCCCGCCACGCGCCACACCGTCGATCCGGACCGCCACATCGAGCGGGTCACCGAGCTG
CAAGAACTCTTCCTCACGCGCGTCGGGCTCGACATCGGCAAGGTGTGGGTCGCGGACGACGGCG
CCGCGGTGGCGGTCTGGACCACGCCGGAGAGCGTCGAAGCGGGGGCGGTGTTCGCCGAGATCG
GCCCGCGCATGGCCGAGTTGAGCGGTTCCCGGCTGGCCGCGCAGCAACAGATGGAAGGCCTCCT
GGCGCCGCACCGGCCCAAGGAGCCCGCGTGGTTCCTGGCCACCGTCGGCGTCTCGCCCGACCA

CCAGGGCAAGGGTCTGGGCAGCGCCGTCGTGCTCCCCGGAGTGGAGGCGGCCGAGCGCGCCGG
GGTGCCCGCCTTCCTGGAGACCTCCGCGCCCCGCAACCTCCCCTTCTACGAGCGGCTCGGCTTCA
CCGTCACCGCCGACGTCGAGGTGCCCGAAGGACCGCGCACCTGGTGCATGACCCGCAAGCCCGG
TGCCTAGGTTTAAACGTCGACAATCAACCTCTGGATTACAAAATTTGTGAAAGATTGACTGGTATTCTT
AACTATGTTGCTCCTTTTACGCTATGTGGATACGCTGCTTTAATGCCTTTGTATCATGCTATTGCTTCC
CGTATGGCTTTCATTTTCTCCTCCTTGTATAAATCCTGGTTGCTGTCTCTTTATGAGGAGTTGTGGCC
CGTTGTCAGGCAACGTGGCGTGGTGTGCACTGTGTTTGCTGACGCAACCCCCACTGGTTGGGGCA
TTGCCACCACCTGTCAGCTCCTTTCCGGGACTTTCGCTTTCCCCCTCCCTATTGCCACGGCGGAAC
TCATCGCCGCCTGCCTTGCCCGCTGCTGGACAGGGGCTCGGCTGTTGGGCACTGACAATTCCGTG
GTGTTGTCGGGGAAGCTGACGTCCTTTCCATGGCTGCTCGCCTGTGTTGCCACCTGGATTCTGCGC
GGGACGTCCTTCTGCTACGTCCCTTCGGCCCTCAATCCAGCGGACCTTCCTTCCCGCGGCCTGCT
GCCGGCTCTGCGGCCTCTTCCGCGTCTTCGCCTTCGCCCTCAGACGAGTCGGATCTCCCTTTGGG
CCGCCTCCCCGCCTGGAATTCGAGCTCGGTACCTTTAAGACCAATGACTTACAAGGCAGCTGTAGA
TCTTAGCCACTTTTTAAAAGAAAAGGGGGGACTGGAAGGGCTAATTCACTCCCAACGAAGACAAGAT
CTGCTTTTTGCTTGTACTGGGTCTCTCTGGTTAGACCAGATCTGAGCCTGGGAGCTCTCTGGCTAAC
TAGGGAACCCACTGCTTAAGCCTCAATAAAGCTTGCCTTGAGTGCTTCAAGTAGTGTGTGCCCGTCT
GTTGTGTGACTCTGGTAACTAGAGATCCCTCAGACCCTTTTAGTCAGTGTGGAAAATCTCTAGCAGT
AGTAGTTCATGTCATCTTATTATTCAGTATTTATAACTTGCAAAGAAATGAATATCAGAGAGTGAGAGG
AACTTGTTTATTGCAGCTTATAATGGTTACAAATAAAGCAATAGCATCACAAATTTCACAAATAAAGCAT
TTTTTTCACTGCATTCTAGTTGTGGTTTGTCCAAACTCATCAATGTATCTTATCATGTCTGGCTCTAGC
TATCCCGCCCCTAACTCCGCCCATCCCGCCCCTAACTCCGCCCAGTTCCGCCCATTCTCCGCCCCA
TGGCTGACTAATTTTTTTTATTTATGCAGAGGCCGAGGCCGCCTCGGCCTCTGAGCTATTCCAGAAG
TAGTGAGGAGGCTTTTTTGGAGGCCTAGGGACGTACCCAATTCGCCCTATAGTGAGTCGTATTACGC
GCGCTCACTGGCCGTCGTTTTACAACGTCGTGACTGGGAAAACCCTGGCGTTACCCAACTTAATCG
CCTTGCAGCACATCCCCCTTTCGCCAGCTGGCGTAATAGCGAAGAGGCCCGCACCGATCGCCCTTC
CCAACAGTTGCGCAGCCTGAATGGCGAATGGGACGCGCCCTGTAGCGGCGCATTAAGCGCGGCG
GGTGTGGTGGTTACGCGCAGCGTGACCGCTACACTTGCCAGCGCCCTAGCGCCCGCTCCTTTCGC
TTTCTTCCCTTCCTTTCTCGCCACGTTCGCCGGCTTTCCCCGTCAAGCTCTAAATCGGGGGCTCCCT
TTAGGGTTCCGATTTAGTGCTTTACGGCACCTCGACCCCAAAAAACTTGATTAGGGTGATGGTTCAC
GTAGTGGGCCATCGCCCTGATAGACGGTTTTTCGCCCTTTGACGTTGGAGTCCACGTTCTTTAATAG
TGGACTCTTGTTCCAAACTGGAACAACACTCAACCCTATCTCGGTCTATTCTTTTGATTTATAAGGGAT
TTTGCCGATTTCGGCCTATTGGTTAAAAAATGAGCTGATTTAACAAAAATTTAACGCGAATTTTAACAA
AATATTAACGCTTACAATTTAGGTGGCACTTTTCGGGGAAATGTGCGCGGAACCCCTATTTGTTTATTT
TTCTAAATACATTCAAATATGTATCCGCTCATGAGACAATAACCCTGATAAATGCTTCAATAATATTGAAA
AAGGAAGAGTATGAGTATTCAACATTTCCGTGTCGCCCTTATTCCCTTTTTTGCGGCATTTTGCCTTC
CTGTTTTTGCTCACCCAGAAACGCTGGTGAAAGTAAAAGATGCTGAAGATCAGTTGGGTGCACGAG
TGGGTTACATCGAACTGGATCTCAACAGCGGTAAGATCCTTGAGAGTTTTCGCCCCGAAGAACGTTT
TCCAATGATGAGCACTTTTAAAGTTCTGCTATGTGGCGCGGTATTATCCCGTATTGACGCCGGGCAA
GAGCAACTCGGTCGCCGCATACACTATTCTCAGAATGACTTGGTTGAGTACTCACCAGTCACAGAAA
AGCATCTTACGGATGGCATGACAGTAAGAGAATTATGCAGTGCTGCCATAACCATGAGTGATAACACT
GCGGCCAACTTACTTCTGACAACGATCGGAGGACCGAAGGAGCTAACCGCTTTTTTGCACAACATG
GGGGATCATGTAACTCGCCTTGATCGTTGGGAACCGGAGCTGAATGAAGCCATACCAAACGACGAG
CGTGACACCACGATGCCTGTAGCAATGGCAACAACGTTGCGCAAACTATTAACTGGCGAACTACTTA
CTCTAGCTTCCCGGCAACAATTAATAGACTGGATGGAGGCGGATAAAGTTGCAGGACCACTTCTGCG

CTCGGCCCTTCCGGCTGGCTGGTTTATTGCTGATAAATCTGGAGCCGGTGAGCGTGGGTCTCGCG
GTATCATTGCAGCACTGGGGCCAGATGGTAAGCCCTCCCGTATCGTAGTTATCTACACGACGGGGA
GTCAGGCAACTATGGATGAACGAAATAGACAGATCGCTGAGATAGGTGCCTCACTGATTAAGCATTG
GTAACTGTCAGACCAAGTTTACTCATATATACTTTAGATTGATTTAAAACTTCATTTTTAATTTAAAAGGA
TCTAGGTGAAGATCCTTTTTGATAATCTCATGACCAAAATCCCTTAACGTGAGTTTTCGTTCCACTGA
GCGTCAGACCCCGTAGAAAAGATCAAAGGATCTTCTTGAGATCCTTTTTTTCTGCGCGTAATCTGCT
GCTTGCAAACAAAAAAACCACCGCTACCAGCGGTGGTTTGTTTGCCGGATCAAGAGCTACCAACTC
TTTTTCCGAAGGTAACTGGCTTCAGCAGAGCGCAGATACCAAATACTGTTCTTCTAGTGTAGCCGTA
GTTAGGCCACCACTTCAAGAACTCTGTAGCACCGCCTACATACCTCGCTCTGCTAATCCTGTTACCA
GTGGCTGCTGCCAGTGGCGATAAGTCGTGTCTTACCGGGTTGGACTCAAGACGATAGTTACCGGAT
AAGGCGCAGCGGTCGGGCTGAACGGGGGGTTCGTGCACACAGCCCAGCTTGGAGCGAACGACCT
ACACCGAACTGAGATACCTACAGCGTGAGCTATGAGAAAGCGCCACGCTTCCCGAAGGGAGAAAG
GCGGACAGGTATCCGGTAAGCGGCAGGGTCGGAACAGGAGAGCGCACGAGGGAGCTTCCAGGGG
GAAACGCCTGGTATCTTTATAGTCCTGTCGGGTTTCGCCACCTCTGACTTGAGCGTCGATTTTTGTG
ATGCTCGTCAGGG

P10:

>P2389 pAAVD-Grin1(800)-CAG-Crimson-PolyA(rbG)-Grin1(950)

CGAAGGAGCTAACCGCTTTTTTGCACAACATGGGGGATCATGTAACTCGCCTTGATCGTTGGGAAC
CGGAGCTGAATGAAGCCATACCAAACGACGAGCGTGACACCACGATGCCTGTAGCAATGGCAACAA
CGTTGCGCAAACTATTAACTGGCGAACTACTTACTCTAGCTTCCCGGCAACAATTAATAGACTGGATG
GAGGCGGATAAAGTTGCAGGACCACTTCTGCGCTCGGCCCTTCCGGCTGGCTGGTTTATTGCTGAT
AAATCTGGAGCCGGTGAGCGTGGGTCTCGCGGTATCATTGCAGCACTGGGGCCAGATGGTAAGCC
CTCCCGTATCGTAGTTATCTACACGACGGGGAGTCAGGCAACTATGGATGAACGAAATAGACAGATC
GCTGAGATAGGTGCCTCACTGATTAAGCATTGGTAACTGTCAGACCAAGTTTACTCATATATACTTTAG
ATTGATTTAAAACTTCATTTTTAATTTAAAAGGATCTAGGTGAAGATCCTTTTTGATAATCTCATGACCA
AAATCCCTTAACGTGAGTTTTCGTTCCACTGAGCGTCAGACCCCGTAGAAAAGATCAAAGGATCTTC
TTGAGATCCTTTTTTTCTGCGCGTAATCTGCTGCTTGCAAACAAAAAAACCACCGCTACCAGCGGTG
GTTTGTTTGCCGGATCAAGAGCTACCAACTCTTTTTCCGAAGGTAACTGGCTTCAGCAGAGCGCAG
ATACCAAATACTGTTCTTCTAGTGTAGCCGTAGTTAGGCCACCACTTCAAGAACTCTGTAGCACCGCC
TACATACCTCGCTCTGCTAATCCTGTTACCAGTGGCTGCTGCCAGTGGCGATAAGTCGTGTCTTACC
GGGTTGGACTCAAGACGATAGTTACCGGATAAGGCGCAGCGGTCGGGCTGAACGGGGGGTTCGTG
CACACAGCCCAGCTTGGAGCGAACGACCTACACCGAACTGAGATACCTACAGCGTGAGCTATGAGA
AAGCGCCACGCTTCCCGAAGGGAGAAAGGCGGACAGGTATCCGGTAAGCGGCAGGGTCGGAACA
GGAGAGCGCACGAGGGAGCTTCCAGGGGGAAACGCCTGGTATCTTTATAGTCCTGTCGGGTTTCG
CCACCTCTGACTTGAGCGTCGATTTTTGTGATGCTCGTCAGGGGGGCGGAGCCTATGGAAAAACGC
CAGCAACGCGGCCTTTTTACGGTTCCTGGCCTTTTGCTGGCCTTTTGCTCACATGTCCTGCAGGCA
GCTGCGCGCTCGCTCGCTCACTGAGGCCGCCCGGGCGTCGGGCGACCTTTGGTCGCCCGGCCTC
AGTGAGCGAGCGAGCGCGCAGAGAGGGAGTGGCCAACTCCATCACTAGGGGTTCCTGCGGCCGC
ACGCGTTCTAGGGAGAGAGGGGGTCTGAAGGCTGTAGTCAAACATCGATAACCAGAAGCATCCATC
ACGATGCCAATGGCTGTTAGGAATGTTTCTGGCCTCACTGGTTTGTTCAATTTGATTTTATATGTATGA
GTGTTTTGCCTACCTGTATATGTAGATACCATGTGCTCTCCTGGTGCCCACAGAATTCAGGATATTGG
ATGGCCTGTAACTGGAATGGATGGTTGTAAGCCACCGTGTGGGCGGTAAGAACAGAAACCGGGTCT
TCTGTAAGACCTTCGGGTGCTCATAACTGCTCACCCACTTGTCTAACCCCATCACTAATTTTTTTTTTT
TTGACAGCGCTAGCAAAAACACATAATAGAAAAATATGATCCTGCTTCCTTCTCTCTTCCTTCTCAAGA
CTCCCAGTGAGGGCACCTCCTCCAGTGCCTGTCACCATGCCTTGCCTCACTGTCCACTCCTGCCTG
GGCTACCTACCCCTCTGTTTTTTGCATCCGATATAGACCCTCAGAAACCAACTGTGTTCCACTCTGG
CATCCCCCAGACTTCTACCTGTCCCTCAACTATGAAACTGTAGACCCTAGAGAATAGGGAGGGAGAA
TGTCAAAGCTGAAGAGGCAGCCTTGAAAGAGAAGCATTAACTAGGCATGGTGGAGCATGCCTTTAA
ACCCAGCACTTTGGAAGTAGATGAACAAGAATCAGAAGTTCAAGGTCCTCCTCAGCTAAAAGCCAG
CAGGGACAGCCTTGACTCTATGAAGGCTCTTTCTTTTTTGTGGTGGGACATCAAGAAGAGTTCATTT
GGCTTAGACTTTCATATTGTGTTCCATCATTGAAGGAAGTCAGGACAGGAACTTAAACCGGGCAGAA
GCCTGGAAGCAGGAGCTGATGCAGAGGCCTGGAGGGGTGCTGTTGATGCAGAGGCCCTGGAGGG
GTGCTGCTGATGCAAAGGCCCTGGAGGGGTGCTGCTGATGCAGAGGCCCTGGAGGGGTGCTGTT
GATGCAGAGGCCTGGAGGGGTGCTGCTGATGCAGAGGCCTGGATGGGTGCTGCTGATGCAGAGG
CCCTGGAGGGGTGCTGTTGATGCAGAGGCCCTGGAGGGGTGCTGTTGATGCAGAGGCCTGGAGG
GGTGCTGCTGATGCAGAGGCCTGGAGGGGTGCTGCTGATGCAGAGGCCTGGAGGGGTGCTGCTC
ACTGGCTTGCTCCTTATGGCTTGCTTGGCTTGCTTTCTTATAAAACCCGGAACCACCCGCCCAGGGT
GGCTTCACCTACCATGGGTTTGCCCCTCTCTCCTTCAATGGCTATTTAAGAAAATGCTTTACAGGCTG
GCCTAGCTAGCCCATTGACGTCAATAATGACGTATGTTCCCATAGTAACGCCAATAGGGACTTTCCAT

TGACGTCAATGGGTGGAGTATTTACGGTAAACTGCCCACTTGGCAGTACATCAAGTGTATCATATGCC
AAGTACGCCCCCTATTGACGTCAATGACGGTAAATGGCCCGCCTGGCATTATGCCCAGTACATGACC
TTATGGGACTTTCCTACTTGGCAGTACATCTACGTATTAGTCATCGCTATTACCATGGTCGAGGTGAG
CCCCACGTTCTGCTTCACTCTCCCCATCTCCCCCCCCTCCCCACCCCCAATTTTGTATTTATTTATTTT
TTAATTATTTTGTGCAGCGATGGGGGCGGGGGGGGGGGGGGGGGGCGCGCGCCAGGCGGGGCG
GGGCGGGGCGAGGGGCGGGGCGGGGCGAGGCGGAGAGGTGCGGCGGCAGCCAATCAGAGCGG
CGCGCTCCGAAAGTTTCCTTTTATGGCGAGGCGGCGGCGGCGGCGGCCCTATAAAAAGCGAAGCG
CGCGGCGGGCGGGAGTCGCTGCGCGCTGCCTTCGCCCCGTGCCCCGCTCCGCCGCCGCCTCGC
GCCGCCCGCCCCGGCTCTGACTGACCGCGTTACTCCCACAGGTGAGCGGGCGGGACGGCCCTTC
TCCTCCGGGCTGTAATTAGCGCTTGGTTTAATGACGGCTTGTTTCTTTTCTGTGGCTGCGTGAAAGC
CTTGAGGGGCTCCGGGAGGGCCCTTTGTGCGGGGGGAGCGGCTCGGGGCTGTCCGCGGGGGGA
CGGCTGCCTTCGGGGGGGACGGGGCAGGGCGGGGTTCGGCTTCTGGCGTGTGACCGGCGGCTC
TAGAGCCTCTGCTAACCATGTTCATGCCTTCTTCTTTTTCCTACAGCTCCTGGGCAACGTGCTGGTTA
TTGTGCTGTCTCATCATTTTGGCAAAGAATTGGATCCGCGATCGCCATGGATAGCACTGAGAACGTC
ATCAAGCCCTTCATGCGCTTCAAGGTGCACATGGAGGGCTCCGTGAACGGCCACGAGTTCGAGAT
CGAGGGCGTGGGCGAGGGCAAGCCCTACGAGGGCACCCAGACCGCCAAGCTGCAAGTGACCAA
GGGCGGCCCCCTGCCCTTCGCCTGGGACATCCTGTCCCCCCAGTTCTTCTACGGCTCCAAGGCGT
ACATCAAGCACCCCGCCGACATCCCCGACTACCTCAAGCAGTCCTTCCCCGAGGGCTTCAAGTGG
GAGCGCGTGATGAACTTCGAGGACGGCGGCGTGGTGACCGTGACCCAGGACTCCTCCCTGCAGG
ACGGCACCCTCATCTACCACGTGAAGTTCATCGGCGTGAACTTCCCCTCCGACGGCCCCGTAATGC
AGAAGAAGACTCTGGGCTGGGAGCCCTCCACTGAGCGCAACTACCCCCGCGACGGCGTGCTGAA
GGGCGAGAACCACATGGCGCTGAAGCTGAAGGGCGGCGGCCACTACCTGTGTGAGTTCAAGTCCA
TCTACATGGCCAAGAAGCCCGTGAAGCTGCCCGGCTACCACTACGTGGACTACAAGCTCGACATCA
CCTCCCACAACGAGGACTACACCGTGGTGGAGCAGTACGAGCGCGCCGAGGCCCGCCACCACCT
GTTCCAGTAGGTTTAAACGATCTTTTTCCCTCTGCCAAAAATTATGGGGACATCATGAAGCCCCTTGA
GCATCTGACTTCTGGCTAATAAAGGAAATTTATTTTCATTGCAATAGTGTGTTGGAATTTTTTGTGTCT
CTCACTCGGTTGGTCACAAAGCTGGCTGACTAACCAATGCAGCACAGTTACTGTGCTCCAGGCTTA
GAAGAAAACTTGTCAAAAATAATATTAGTGATAATGATAACAATAATAAAAAACAGTGGGACTGGAGAG
ATGGCTCAGTGGTTTAGAGCCCTGGCTATTCTCCCATAGGACCCGGGTTCAAGTCCCAGTACCCAC
ACAGCAGCTCACAACCATCTGTATCTATAGTTCCAGGGGATCTAATGTTCTTTCTGACCTCTGTGGGT
TCTGTATGCACCTGGAGTACAAACGTGTATGCTGGCAAAACACCCATACACTTAAAATAGAGAAGATA
TTCTCTCAGAGGTTAAGAGAAATTGTTGCTCTTCCAAAAAAAAAAAAAAAAAAAAAAAAAAACTGGGCT
CAGTTCCCAGCACCCACATGGTAGTTACAACTATGTGTAGCTTCAGTTCCAGGGGATTTGACGCCTC
CTTCTGGTTTCTGTGGACCCCTGCATGTACGTGATACACATACATACACTGAGGCACACACACACAC
ATAAGATAAAAACAAAAGATAAATATATAAGTGATAAAAGTGATAGAGAAAGACATGGGGCATTATCCTC
CATCCTCTGTGTACTTGCTCATATACATGAATACACAGGAACACATACATACATATAAACATATACATACA
TGTAAACATGTATACATATATACATACATATAAAACTACCTAAAAGGTAGTTTTTTAAAAGATGAGGATCA
AAGACTGAAGGCTACATGCCCATGGAGGCCAGGGTAGCTTTGGAAAGGACACAGCCTTTCTGCTCC
CTGATTTGTCCGAGACAGTGCCTGGTGCACACTTCCTGCACAGTAGCACTATGATAGAATAATGTATA
AAGGGGCTGGAGAGATGGCTCAGTGGTTGAGAGCACTGACTGCTCTTCAGAAGGTCCTGAGTTCA
ATTCCCGGCAACCACATGATGGCTCACAACCATCCGTAATGAGATCTGGCACCCTCTTCTGGAGTGT
TTGAAGACAGCTACAGTGTACTTACATATAATAAATAAATAAATCTTTTTTTAAAAAATCTGAAAAATAAA
AGAAGAATGTATAAGCTTAAGCTGCAGCCTGAGAGAAAGGGGCCACAGGAGAGGTCGCAGTACTGC
AGGATGCATTTTTTGATGAGTCAGGGCAGGTGGCTATACACGTGCGGACCGAGCGGCCGCAGGAA

CCCCTAGTGATGGAGTTGGCCACTCCCTCTCTGCGCGCTCGCTCGCTCACTGAGGCCGGGCGACC
AAAGGTCGCCCGACGCCCGGGCTTTGCCCGGGCGGCCTCAGTGAGCGAGCGAGCGCGCAGCTG
CCTGCAGGGGCGCCTGATGCGGTATTTTCTCCTTACGCATCTGTGCGGTATTTCACACCGCATACGT
CAAAGCAACCATAGTACGCGCCCTGTAGCGGCGCATTAAGCGCGGCGGGTGTGGTGGTTACGCGC
AGCGTGACCGCTACACTTGCCAGCGCCTTAGCGCCCGCTCCTTTCGCTTTCTTCCCTTCCTTTCTC
GCCACGTTCGCCGGCTTTCCCCGTCAAGCTCTAAATCGGGGGCTCCCTTTAGGGTTCCGATTTAGT
GCTTTACGGCACCTCGACCCCAAAAAACTTGATTTGGGTGATGGTTCACGTAGTGGGCCATCGCCC
TGATAGACGGTTTTTCGCCCTTTGACGTTGGAGTCCACGTTCTTTAATAGTGGACTCTTGTTCCAAAC
TGGAACAACACTCAACTCTATCTCGGGCTATTCTTTTGATTTATAAGGGATTTTGCCGATTTCGGTCTA
TTGGTTAAAAAATGAGCTGATTTAACAAAAATTTAACGCGAATTTTAACAAAATATTAACGTTTACAATT
TTATGGTGCACTCTCAGTACAATCTGCTCTGATGCCGCATAGTTAAGCCAGCCCCGACACCCGCCAA
CACCCGCTGACGCGCCCTGACGGGCTTGTCTGCTCCCGGCATCCGCTTACAGACAAGCTGTGACC
GTCTCCGGGAGCTGCATGTGTCAGAGGTTTTCACCGTCATCACCGAAACGCGCGAGACGAAAGGG
CCTCGTGATACGCCTATTTTTATAGGTTAATGTCATGATAATAATGGTTTCTTAGACGTCAGGTGGCAC
TTTTCGGGGAAATGTGCGCGGAACCCCTATTTGTTTATTTTTCTAAATACATTCAAATATGTATCCGCT
CATGAGACAATAACCCTGATAAATGCTTCAATAATATTGAAAAAGGAAGAGTATGAGTATTCAACATTT
CCGTGTCGCCCTTATTCCCTTTTTTGCGGCATTTTGCCTTCCTGTTTTTGCTCACCCAGAAACGCTG
GTGAAAGTAAAAGATGCTGAAGATCAGTTGGGTGCACGAGTGGGTTACATCGAACTGGATCTCAAC
AGCGGTAAGATCCTTGAGAGTTTTCGCCCCGAAGAACGTTTTCCAATGATGAGCACTTTTAAAGTTC
TGCTATGTGGCGCGGTATTATCCCGTATTGACGCCGGGCAAGAGCAACTCGGTCGCCGCATACACTA
TTCTCAGAATGACTTGGTTGAGTACTCACCAGTCACAGAAAAGCATCTTACGGATGGCATGACAGTA
AGAGAATTATGCAGTGCTGCCATAACCATGAGTGATAACACTGCGGCCAACTTACTTCTGACAACGA
TCGGAGGAC

P11:

>S1487 pU6-sgKLRC1a-gN21(Sa-v2)

AGCTGGCACGACAGGTTTCCCGACTGGAAAGCGGGCAGTGAGCGCAACGCAATTAAGGTGAGTTA
GCTCACTCATTAGGCACCCCAGGCTTTACACTTTATGCTTCCGGCTCGTATGTTGTGTGGAATTGTG
AGCGGATAACAATTTCACACAGGACCCCAGTGGAAAGACGCGCAGGCAAAACGCACCACGTGACG
GAGCGTGACCGCGCGCCGAGCGCGCGCCAAGGTCGGGCAGGAAGAGGGCCTATTTCCCATGATT
CCTTCATATTTGCATATACGATACAAGGCTGTTAGAGAGATAATTAGAATTAATTTGACTGTAAACACAA
AGATATTAGTACAAAATACGTGACGTAGAAAGTAATAATTTCTTGGGTAGTTTGCAGTTTTAAAATTATG
TTTTAAAATGGACTATCATATGCTTACCGTAACTTGAAAGTATTTCGATTTCTTGGGTTTATATATCTTGT
GGAAAGGACGAAACACCGCACTGCAGAGATGGATAACCAGTTTAAGTACTCTGTGCTGGAAACAGC
ACAGAATCTACTTAAACAAGGCAAAATGCCGTGTTTATCTCGTCAACTTGTTGGCGAGATTTTTTTCT
CGAGCAGGTGGCACTTTTCGGGGAAATGTGCGCGGAACCCCTATTTGTTTATTTTTCTAAATACATTC
AAATATGTATCCGCTCATGAGACAATAACCCTGATAAATGCTTCAATAATATTGAAAAAGGAAGAGTAT
GAGTATTCAACATTTCCGTGTCGCCCTTATTCCCTTTTTTGCGGCATTTTGCCTTCCTGTTTTTGCTCA
CCCAGAAACGCTGGTGAAAGTAAAAGATGCTGAAGATCAGTTGGGTGCACGAGTGGGTTACATCGA
ACTGGATCTCAACAGCGGTAAGATCCTTGAGAGTTTTCGCCCCGAAGAACGTTTTCCAATGATGAGC
ACTTTTAAAGTTCTGCTATGTGGCGCGGTATTATCCCGTATTGACGCCGGGCAAGAGCAACTCGGTC
GCCGCATACACTATTCTCAGAATGACTTGGTTGAGTACTCACCAGTCACAGAAAAGCATCTTACGGAT
GGCATGACAGTAAGAGAATTATGCAGTGCTGCCATAACCATGAGTGATAACACTGCGGCCAACTTAC
TTCTGACAACGATCGGAGGACCGAAGGAGCTAACCGCTTTTTTGCACAACATGGGGGATCATGTAA
CTCGCCTTGATCGTTGGGAACCGGAGCTGAATGAAGCCATACCAAACGACGAGCGTGACACCACGA
TGCCTGTAGCAATGGCAACAACGTTGCGCAAACTATTAACTGGCGAACTACTTACTCTAGCTTCCCG
GCAACAATTAATAGACTGGATGGAGGCGGATAAAGTTGCAGGACCACTTCTGCGCTCGGCCCTTCC
GGCTGGCTGGTTTATTGCTGATAAATCTGGAGCCGGTGAGCGTGGGTCTCGCGGTATCATTGCAGC
ACTGGGGCCAGATGGTAAGCCCTCCCGTATCGTAGTTATCTACACGACGGGGAGTCAGGCAACTAT
GGATGAACGAAATAGACAGATCGCTGAGATAGGTGCCTCACTGATTAAGCATTGGTAACTGTCAGAC
CAAGTTTACTCATATATACTTTAGATTGATTTAAAACTTCATTTTTAATTTAAAAGGATCTAGGTGAAGAT
CCTTTTTGATAATCTCATGACCAAAATCCCTTAACGTGAGTTTTCGTTCCACTGAGCGTCAGACCCCG
TAGAAAAGATCAAAGGATCTTCTTGAGATCCTTTTTTTCTGCGCGTAATCTGCTGCTTGCAAACAAAA
AAACCACCGCTACCAGCGGTGGTTTGTTTGCCGGATCAAGAGCTACCAACTCTTTTTCCGAAGGTAA
CTGGCTTCAGCAGAGCGCAGATACCAAATACTGTTCTTCTAGTGTAGCCGTAGTTAGGCCACCACTT
CAAGAACTCTGTAGCACCGCCTACATACCTCGCTCTGCTAATCCTGTTACCAGTGGCTGCTGCCAGT
GGCGATAAGTCGTGTCTTACCGGGTTGGACTCAAGACGATAGTTACCGGATAAGGCGCAGCGGTCG
GGCTGAACGGGGGGTTCGTGCACACAGCCCAGCTTGGAGCGAACGACCTACACCGAACTGAGATA
CCTACAGCGTGAGCTATGAGAAAGCGCCACGCTTCCCGAAGGGAGAAAGGCGGACAGGTATCCGG
TAAGCGGCAGGGTCGGAACAGGAGAGCGCACGAGGGAGCTTCCAGGGGGAAACGCCTGGTATCT
TTATAGTCCTGTCGGGTTTCGCCACCTCTGACTTGAGCGTCGATTTTTGTGATGCTCGTCAGGGGG
GCGGAGCCTATGGAAAAACGCCAGCAACGCGGCCTTTTTACGGTTCCTGGCCTTTTGCTGGCCTTT
TGCTCACATGTTCTTTCCTGCGTTATCCCCTGATTCTGTGGATAACCGTATTACCGCCTTTGAGTGAG
CTGATACCGCTCGCCGCAGCCGAACGACCGAGCGCAGCGAGTCAGTGAGCGAGGAAGCGGAAGA
GCGCCCAATACGCAAACCGCCTCTCCCCGCGCGTTGGCCGATTCATTAATGC

P12:

>S1489 pU6-sgKLRC1c-gN21(Sa-v2)

AGCTGGCACGACAGGTTTCCCGACTGGAAAGCGGGCAGTGAGCGCAACGCAATTAAGGTGAGTTA
GCTCACTCATTAGGCACCCCAGGCTTTACACTTTATGCTTCCGGCTCGTATGTTGTGTGGAATTGTG
AGCGGATAACAATTTCACACAGGACCCCAGTGGAAAGACGCGCAGGCAAAACGCACCACGTGACG
GAGCGTGACCGCGCGCCGAGCGCGCGCCAAGGTCGGGCAGGAAGAGGGCCTATTTCCCATGATT
CCTTCATATTTGCATATACGATACAAGGCTGTTAGAGAGATAATTAGAATTAATTTGACTGTAAACACAA
AGATATTAGTACAAAATACGTGACGTAGAAAGTAATAATTTCTTGGGTAGTTTGCAGTTTTAAAATTATG
TTTTAAAATGGACTATCATATGCTTACCGTAACTTGAAAGTATTTCGATTTCTTGGGTTTATATATCTTGT
GGAAAGGACGAAACACCGGTTTGGGGGCAGATTCAGGTGTTTAAGTACTCTGTGCTGGAAACAGCA
CAGAATCTACTTAAACAAGGCAAAATGCCGTGTTTATCTCGTCAACTTGTTGGCGAGATTTTTTTCTC
GAGCAGGTGGCACTTTTCGGGGAAATGTGCGCGGAACCCCTATTTGTTTATTTTTCTAAATACATTCA
AATATGTATCCGCTCATGAGACAATAACCCTGATAAATGCTTCAATAATATTGAAAAAGGAAGAGTATG
AGTATTCAACATTTCCGTGTCGCCCTTATTCCCTTTTTTGCGGCATTTTGCCTTCCTGTTTTTGCTCAC
CCAGAAACGCTGGTGAAAGTAAAAGATGCTGAAGATCAGTTGGGTGCACGAGTGGGTTACATCGAA
CTGGATCTCAACAGCGGTAAGATCCTTGAGAGTTTTCGCCCCGAAGAACGTTTTCCAATGATGAGCA
CTTTTAAAGTTCTGCTATGTGGCGCGGTATTATCCCGTATTGACGCCGGGCAAGAGCAACTCGGTCG
CCGCATACACTATTCTCAGAATGACTTGGTTGAGTACTCACCAGTCACAGAAAAGCATCTTACGGATG
GCATGACAGTAAGAGAATTATGCAGTGCTGCCATAACCATGAGTGATAACACTGCGGCCAACTTACTT
CTGACAACGATCGGAGGACCGAAGGAGCTAACCGCTTTTTTGCACAACATGGGGGATCATGTAACT
CGCCTTGATCGTTGGGAACCGGAGCTGAATGAAGCCATACCAAACGACGAGCGTGACACCACGAT
GCCTGTAGCAATGGCAACAACGTTGCGCAAACTATTAACTGGCGAACTACTTACTCTAGCTTCCCGG
CAACAATTAATAGACTGGATGGAGGCGGATAAAGTTGCAGGACCACTTCTGCGCTCGGCCCTTCCG
GCTGGCTGGTTTATTGCTGATAAATCTGGAGCCGGTGAGCGTGGGTCTCGCGGTATCATTGCAGCA
CTGGGGCCAGATGGTAAGCCCTCCCGTATCGTAGTTATCTACACGACGGGGAGTCAGGCAACTATG
GATGAACGAAATAGACAGATCGCTGAGATAGGTGCCTCACTGATTAAGCATTGGTAACTGTCAGACC
AAGTTTACTCATATATACTTTAGATTGATTTAAAACTTCATTTTTAATTTAAAAGGATCTAGGTGAAGATC
CTTTTTGATAATCTCATGACCAAAATCCCTTAACGTGAGTTTTCGTTCCACTGAGCGTCAGACCCCGT
AGAAAAGATCAAAGGATCTTCTTGAGATCCTTTTTTTCTGCGCGTAATCTGCTGCTTGCAAACAAAAA
AACCACCGCTACCAGCGGTGGTTTGTTTGCCGGATCAAGAGCTACCAACTCTTTTTCCGAAGGTAA
CTGGCTTCAGCAGAGCGCAGATACCAAATACTGTTCTTCTAGTGTAGCCGTAGTTAGGCCACCACTT
CAAGAACTCTGTAGCACCGCCTACATACCTCGCTCTGCTAATCCTGTTACCAGTGGCTGCTGCCAGT
GGCGATAAGTCGTGTCTTACCGGGTTGGACTCAAGACGATAGTTACCGGATAAGGCGCAGCGGTCG
GGCTGAACGGGGGGTTCGTGCACACAGCCCAGCTTGGAGCGAACGACCTACACCGAACTGAGATA
CCTACAGCGTGAGCTATGAGAAAGCGCCACGCTTCCCGAAGGGAGAAAGGCGGACAGGTATCCGG
TAAGCGGCAGGGTCGGAACAGGAGAGCGCACGAGGGAGCTTCCAGGGGGAAACGCCTGGTATCT
TTATAGTCCTGTCGGGTTTCGCCACCTCTGACTTGAGCGTCGATTTTTGTGATGCTCGTCAGGGGG
GCGGAGCCTATGGAAAAACGCCAGCAACGCGGCCTTTTTACGGTTCCTGGCCTTTTGCTGGCCTTT
TGCTCACATGTTCTTTCCTGCGTTATCCCCTGATTCTGTGGATAACCGTATTACCGCCTTTGAGTGAG
CTGATACCGCTCGCCGCAGCCGAACGACCGAGCGCAGCGAGTCAGTGAGCGAGGAAGCGGAAGA
GCGCCCAATACGCAAACCGCCTCTCCCCGCGCGTTGGCCGATTCATTAATGC

P13:

>P2209 pIVT-T7-Nco-U3.Hba-A10(Gaaaac)

CGCTGGTGAAAGTAAAAGATGCTGAAGATCAGTTGGGTGCACGAGTGGGTTACATCGAACTGGATC
TCAACAGCGGTAAGATCCTTGAGAGTTTTCGCCCCGAAGAACGTTTTCCAATGATGAGCACTTTTAA
AGTTCTGCTATGTGGCGCGGTATTATCCCGTATTGACGCCGGGCAAGAGCAACTCGGTCGCCGCAT
ACACTATTCTCAGAATGACTTGGTTGAGTACTCACCAGTCACAGAAAAGCATCTTACGGATGGCATG
ACAGTAAGAGAATTATGCAGTGCTGCCATAACCATGAGTGATAACACTGCGGCCAACTTACTTCTGA
CAACGATCGGAGGACCGAAGGAGCTAACCGCTTTTTTGCACAACATGGGGGATCATGTAACTCGCC
TTGATCGTTGGGAACCGGAGCTGAATGAAGCCATACCAAACGACGAGCGTGACACCACGATGCCTG
TAGCAATGGCAACAACGTTGCGCAAACTATTAACTGGCGAACTACTTACTCTAGCTTCCCGGCAACA
ATTAATAGACTGGATGGAGGCGGATAAAGTTGCAGGACCACTTCTGCGCTCGGCCCTTCCGGCTGG
CTGGTTTATTGCTGATAAATCTGGAGCCGGTGAGCGTGGGTCTCGCGGTATCATTGCAGCACTGGG
GCCAGATGGTAAGCCCTCCCGTATCGTAGTTATCTACACGACGGGGAGTCAGGCAACTATGGATGAA
CGAAATAGACAGATCGCTGAGATAGGTGCCTCACTGATTAAGCATTGGTAACTGTCAGACCAAGTTT
ACTCATATATACTTTAGATTGATTTAAAACTTCATTTTTAATTTAAAAGGATCTAGGTGAAGATCCTTTTT
GATAATCTCATGACCAAAATCCCTTAACGTGAGTTTTCGTTCCACTGAGCGTCAGACCCCGTAGAAA
AGATCAAAGGATCTTCTTGAGATCCTTTTTTTCTGCGCGTAATCTGCTGCTTGCAAACAAAAAAACCA
CCGCTACCAGCGGTGGTTTGTTTGCCGGATCAAGAGCTACCAACTCTTTTTCCGAAGGTAACTGGC
TTCAGCAGAGCGCAGATACCAAATACTGTTCTTCTAGTGTAGCCGTAGTTAGGCCACCACTTCAAGA
ACTCTGTAGCACCGCCTACATACCTCGCTCTGCTAATCCTGTTACCAGTGGCTGCTGCCAGTGGCGA
TAAGTCGTGTCTTACCGGGTTGGACTCAAGACGATAGTTACCGGATAAGGCGCAGCGGTCGGGCTG
AACGGGGGGTTCGTGCACACAGCCCAGCTTGGAGCGAACGACCTACACCGAACTGAGATACCTAC
AGCGTGAGCTATGAGAAAGCGCCACGCTTCCCGAAGGGAGAAAGGCGGACAGGTATCCGGTAAGC
GGCAGGGTCGGAACAGGAGAGCGCACGAGGGAGCTTCCAGGGGGAAACGCCTGGTATCTTTATAG
TCCTGTCGGGTTTCGCCACCTCTGACTTGAGCGTCGATTTTTGTGATGCTCGTCAGGGGGGCGGA
GCCTATGGAAAAACGCCAGCAACGCGGCCTTTTTACGGTTCCTGGCCTTTTGCTGGCCTTTTGCTC
ACATGTTCTTTCCTGCGTTATCCCCTGATTCTGTGGATAACCGTATTACCGCCTTTGAGTGAGCTGAT
ACCGCTCGCCGCAGCCGAACGACCGAGCGCAGCGAGTCAGTGAGCGAGGAAGCGGAAGGCGCC
CAATACGCAAACCGCCTCTCCCCGCGCGTTGGCCGATTCATTAATGCAGCTGGCACGACAGGTTTC
CCGACTGGAAAGCGGGCAGTGAGCGCAACGCAATTAAGGTGAGTTAGCTCACTCATTAGGCACCCC
AGGCTTTACACTTTATGCTTCCGGCTCGTATGTTGTGTGGAATTGTGAGCGGATAACAATTTCACACA
GATATCTAATACGACTCACTATAAGAGGAAATAAGAGAGAAAAGAAGAGTAAGAAGAAATATAAGAGC
CACCATGGTAAGTTTAAACGCTGCCTTCTGCGGGGCTTGCCTTCTGGCCATGCCCTTCTTCTCTCCC
TTGCACCTGTACCTCTTGGTCTTTGAATAAAGCCTGAGTAGGAAAAAAAAAAGAAAACGAAGAGCCA
GGTGGCACTTTTCGGGGAAATGTGCGCGGAACCCCTATTTGTTTATTTTTCTAAATACATTCAAATATG
TATCCGCTCATGAGACAATAACCCTGATAAATGCTTCAATAATATTGAAAAAGGAAGAGTATGAGTATT
CAACATTTCCGTGTCGCCCTTATTCCCTTTTTTGCGGCATTTTGCCTTCCTGTTTTTGCTCACCCAGA
AA

P14:

>P2210 pIVT-T7-Nco-U3.Hba-A20(Gaaaac)

CGCTGGTGAAAGTAAAAGATGCTGAAGATCAGTTGGGTGCACGAGTGGGTTACATCGAACTGGATC
TCAACAGCGGTAAGATCCTTGAGAGTTTTCGCCCCGAAGAACGTTTTCCAATGATGAGCACTTTTAA
AGTTCTGCTATGTGGCGCGGTATTATCCCGTATTGACGCCGGGCAAGAGCAACTCGGTCGCCGCAT
ACACTATTCTCAGAATGACTTGGTTGAGTACTCACCAGTCACAGAAAAGCATCTTACGGATGGCATG
ACAGTAAGAGAATTATGCAGTGCTGCCATAACCATGAGTGATAACACTGCGGCCAACTTACTTCTGA
CAACGATCGGAGGACCGAAGGAGCTAACCGCTTTTTTGCACAACATGGGGGATCATGTAACTCGCC
TTGATCGTTGGGAACCGGAGCTGAATGAAGCCATACCAAACGACGAGCGTGACACCACGATGCCTG
TAGCAATGGCAACAACGTTGCGCAAACTATTAACTGGCGAACTACTTACTCTAGCTTCCCGGCAACA
ATTAATAGACTGGATGGAGGCGGATAAAGTTGCAGGACCACTTCTGCGCTCGGCCCTTCCGGCTGG
CTGGTTTATTGCTGATAAATCTGGAGCCGGTGAGCGTGGGTCTCGCGGTATCATTGCAGCACTGGG
GCCAGATGGTAAGCCCTCCCGTATCGTAGTTATCTACACGACGGGGAGTCAGGCAACTATGGATGAA
CGAAATAGACAGATCGCTGAGATAGGTGCCTCACTGATTAAGCATTGGTAACTGTCAGACCAAGTTT
ACTCATATATACTTTAGATTGATTTAAAACTTCATTTTTAATTTAAAAGGATCTAGGTGAAGATCCTTTTT
GATAATCTCATGACCAAAATCCCTTAACGTGAGTTTTCGTTCCACTGAGCGTCAGACCCCGTAGAAA
AGATCAAAGGATCTTCTTGAGATCCTTTTTTTCTGCGCGTAATCTGCTGCTTGCAAACAAAAAAACCA
CCGCTACCAGCGGTGGTTTGTTTGCCGGATCAAGAGCTACCAACTCTTTTTCCGAAGGTAACTGGC
TTCAGCAGAGCGCAGATACCAAATACTGTTCTTCTAGTGTAGCCGTAGTTAGGCCACCACTTCAAGA
ACTCTGTAGCACCGCCTACATACCTCGCTCTGCTAATCCTGTTACCAGTGGCTGCTGCCAGTGGCGA
TAAGTCGTGTCTTACCGGGTTGGACTCAAGACGATAGTTACCGGATAAGGCGCAGCGGTCGGGCTG
AACGGGGGGTTCGTGCACACAGCCCAGCTTGGAGCGAACGACCTACACCGAACTGAGATACCTAC
AGCGTGAGCTATGAGAAAGCGCCACGCTTCCCGAAGGGAGAAAGGCGGACAGGTATCCGGTAAGC
GGCAGGGTCGGAACAGGAGAGCGCACGAGGGAGCTTCCAGGGGGAAACGCCTGGTATCTTTATAG
TCCTGTCGGGTTTCGCCACCTCTGACTTGAGCGTCGATTTTTGTGATGCTCGTCAGGGGGGCGGA
GCCTATGGAAAAACGCCAGCAACGCGGCCTTTTTACGGTTCCTGGCCTTTTGCTGGCCTTTTGCTC
ACATGTTCTTTCCTGCGTTATCCCCTGATTCTGTGGATAACCGTATTACCGCCTTTGAGTGAGCTGAT
ACCGCTCGCCGCAGCCGAACGACCGAGCGCAGCGAGTCAGTGAGCGAGGAAGCGGAAGGCGCC
CAATACGCAAACCGCCTCTCCCCGCGCGTTGGCCGATTCATTAATGCAGCTGGCACGACAGGTTTC
CCGACTGGAAAGCGGGCAGTGAGCGCAACGCAATTAAGGTGAGTTAGCTCACTCATTAGGCACCCC
AGGCTTTACACTTTATGCTTCCGGCTCGTATGTTGTGTGGAATTGTGAGCGGATAACAATTTCACACA
GATATCTAATACGACTCACTATAAGAGGAAATAAGAGAGAAAAGAAGAGTAAGAAGAAATATAAGAGC
CACCATGGTAAGTTTAAACGCTGCCTTCTGCGGGGCTTGCCTTCTGGCCATGCCCTTCTTCTCTCCC
TTGCACCTGTACCTCTTGGTCTTTGAATAAAGCCTGAGTAGGAAAAAAAAAAAAAAAAAAAAGAAAAC
GAAGAGCCAGGTGGCACTTTTCGGGGAAATGTGCGCGGAACCCCTATTTGTTTATTTTTCTAAATACA
TTCAAATATGTATCCGCTCATGAGACAATAACCCTGATAAATGCTTCAATAATATTGAAAAAGGAAGAG
TATGAGTATTCAACATTTCCGTGTCGCCCTTATTCCCTTTTTTGCGGCATTTTGCCTTCCTGTTTTTGC
TCACCCAGAAA

P15:

>P2210 pIVT-T7-Nco-U3.Hba-A30(Gaaaac)

CGCTGGTGAAAGTAAAAGATGCTGAAGATCAGTTGGGTGCACGAGTGGGTTACATCGAACTGGATC
TCAACAGCGGTAAGATCCTTGAGAGTTTTCGCCCCGAAGAACGTTTTCCAATGATGAGCACTTTTAA
AGTTCTGCTATGTGGCGCGGTATTATCCCGTATTGACGCCGGGCAAGAGCAACTCGGTCGCCGCAT
ACACTATTCTCAGAATGACTTGGTTGAGTACTCACCAGTCACAGAAAAGCATCTTACGGATGGCATG
ACAGTAAGAGAATTATGCAGTGCTGCCATAACCATGAGTGATAACACTGCGGCCAACTTACTTCTGA
CAACGATCGGAGGACCGAAGGAGCTAACCGCTTTTTTGCACAACATGGGGGATCATGTAACTCGCC
TTGATCGTTGGGAACCGGAGCTGAATGAAGCCATACCAAACGACGAGCGTGACACCACGATGCCTG
TAGCAATGGCAACAACGTTGCGCAAACTATTAACTGGCGAACTACTTACTCTAGCTTCCCGGCAACA
ATTAATAGACTGGATGGAGGCGGATAAAGTTGCAGGACCACTTCTGCGCTCGGCCCTTCCGGCTGG
CTGGTTTATTGCTGATAAATCTGGAGCCGGTGAGCGTGGGTCTCGCGGTATCATTGCAGCACTGGG
GCCAGATGGTAAGCCCTCCCGTATCGTAGTTATCTACACGACGGGGAGTCAGGCAACTATGGATGAA
CGAAATAGACAGATCGCTGAGATAGGTGCCTCACTGATTAAGCATTGGTAACTGTCAGACCAAGTTT
ACTCATATATACTTTAGATTGATTTAAAACTTCATTTTTAATTTAAAAGGATCTAGGTGAAGATCCTTTTT
GATAATCTCATGACCAAAATCCCTTAACGTGAGTTTTCGTTCCACTGAGCGTCAGACCCCGTAGAAA
AGATCAAAGGATCTTCTTGAGATCCTTTTTTTCTGCGCGTAATCTGCTGCTTGCAAACAAAAAAACCA
CCGCTACCAGCGGTGGTTTGTTTGCCGGATCAAGAGCTACCAACTCTTTTTCCGAAGGTAACTGGC
TTCAGCAGAGCGCAGATACCAAATACTGTTCTTCTAGTGTAGCCGTAGTTAGGCCACCACTTCAAGA
ACTCTGTAGCACCGCCTACATACCTCGCTCTGCTAATCCTGTTACCAGTGGCTGCTGCCAGTGGCGA
TAAGTCGTGTCTTACCGGGTTGGACTCAAGACGATAGTTACCGGATAAGGCGCAGCGGTCGGGCTG
AACGGGGGGTTCGTGCACACAGCCCAGCTTGGAGCGAACGACCTACACCGAACTGAGATACCTAC
AGCGTGAGCTATGAGAAAGCGCCACGCTTCCCGAAGGGAGAAAGGCGGACAGGTATCCGGTAAGC
GGCAGGGTCGGAACAGGAGAGCGCACGAGGGAGCTTCCAGGGGGAAACGCCTGGTATCTTTATAG
TCCTGTCGGGTTTCGCCACCTCTGACTTGAGCGTCGATTTTTGTGATGCTCGTCAGGGGGGCGGA
GCCTATGGAAAAACGCCAGCAACGCGGCCTTTTTACGGTTCCTGGCCTTTTGCTGGCCTTTTGCTC
ACATGTTCTTTCCTGCGTTATCCCCTGATTCTGTGGATAACCGTATTACCGCCTTTGAGTGAGCTGAT
ACCGCTCGCCGCAGCCGAACGACCGAGCGCAGCGAGTCAGTGAGCGAGGAAGCGGAAGGCGCC
CAATACGCAAACCGCCTCTCCCCGCGCGTTGGCCGATTCATTAATGCAGCTGGCACGACAGGTTTC
CCGACTGGAAAGCGGGCAGTGAGCGCAACGCAATTAAGGTGAGTTAGCTCACTCATTAGGCACCCC
AGGCTTTACACTTTATGCTTCCGGCTCGTATGTTGTGTGGAATTGTGAGCGGATAACAATTTCACACA
GATATCTAATACGACTCACTATAAGAGGAAATAAGAGAGAAAAGAAGAGTAAGAAGAAATATAAGAGC
CACCATGGTAAGTTTAAACGCTGCCTTCTGCGGGGCTTGCCTTCTGGCCATGCCCTTCTTCTCTCCC
TTGCACCTGTACCTCTTGGTCTTTGAATAAAGCCTGAGTAGGAAAAAAAAAAAAAAAAAAAAAAAAAA
AAAAGAAAACGAAGAGCCAGGTGGCACTTTTCGGGGAAATGTGCGCGGAACCCCTATTTGTTTATTT
TTCTAAATACATTCAAATATGTATCCGCTCATGAGACAATAACCCTGATAAATGCTTCAATAATATTGAAA
AAGGAAGAGTATGAGTATTCAACATTTCCGTGTCGCCCTTATTCCCTTTTTTGCGGCATTTTGCCTTC
CTGTTTTTGCTCACCCAGAAA

P16:

>P2210 pIVT-T7-Nco-U3.Hba-A60(Gaaaac)

CGCTGGTGAAAGTAAAAGATGCTGAAGATCAGTTGGGTGCACGAGTGGGTTACATCGAACTGGATC
TCAACAGCGGTAAGATCCTTGAGAGTTTTCGCCCCGAAGAACGTTTTCCAATGATGAGCACTTTTAA
AGTTCTGCTATGTGGCGCGGTATTATCCCGTATTGACGCCGGGCAAGAGCAACTCGGTCGCCGCAT
ACACTATTCTCAGAATGACTTGGTTGAGTACTCACCAGTCACAGAAAAGCATCTTACGGATGGCATG
ACAGTAAGAGAATTATGCAGTGCTGCCATAACCATGAGTGATAACACTGCGGCCAACTTACTTCTGA
CAACGATCGGAGGACCGAAGGAGCTAACCGCTTTTTTGCACAACATGGGGGATCATGTAACTCGCC
TTGATCGTTGGGAACCGGAGCTGAATGAAGCCATACCAAACGACGAGCGTGACACCACGATGCCTG
TAGCAATGGCAACAACGTTGCGCAAACTATTAACTGGCGAACTACTTACTCTAGCTTCCCGGCAACA
ATTAATAGACTGGATGGAGGCGGATAAAGTTGCAGGACCACTTCTGCGCTCGGCCCTTCCGGCTGG
CTGGTTTATTGCTGATAAATCTGGAGCCGGTGAGCGTGGGTCTCGCGGTATCATTGCAGCACTGGG
GCCAGATGGTAAGCCCTCCCGTATCGTAGTTATCTACACGACGGGGAGTCAGGCAACTATGGATGAA
CGAAATAGACAGATCGCTGAGATAGGTGCCTCACTGATTAAGCATTGGTAACTGTCAGACCAAGTTT
ACTCATATATACTTTAGATTGATTTAAAACTTCATTTTTAATTTAAAAGGATCTAGGTGAAGATCCTTTTT
GATAATCTCATGACCAAAATCCCTTAACGTGAGTTTTCGTTCCACTGAGCGTCAGACCCCGTAGAAA
AGATCAAAGGATCTTCTTGAGATCCTTTTTTTCTGCGCGTAATCTGCTGCTTGCAAACAAAAAAACCA
CCGCTACCAGCGGTGGTTTGTTTGCCGGATCAAGAGCTACCAACTCTTTTTCCGAAGGTAACTGGC
TTCAGCAGAGCGCAGATACCAAATACTGTTCTTCTAGTGTAGCCGTAGTTAGGCCACCACTTCAAGA
ACTCTGTAGCACCGCCTACATACCTCGCTCTGCTAATCCTGTTACCAGTGGCTGCTGCCAGTGGCGA
TAAGTCGTGTCTTACCGGGTTGGACTCAAGACGATAGTTACCGGATAAGGCGCAGCGGTCGGGCTG
AACGGGGGGTTCGTGCACACAGCCCAGCTTGGAGCGAACGACCTACACCGAACTGAGATACCTAC
AGCGTGAGCTATGAGAAAGCGCCACGCTTCCCGAAGGGAGAAAGGCGGACAGGTATCCGGTAAGC
GGCAGGGTCGGAACAGGAGAGCGCACGAGGGAGCTTCCAGGGGGAAACGCCTGGTATCTTTATAG
TCCTGTCGGGTTTCGCCACCTCTGACTTGAGCGTCGATTTTTGTGATGCTCGTCAGGGGGGCGGA
GCCTATGGAAAAACGCCAGCAACGCGGCCTTTTTACGGTTCCTGGCCTTTTGCTGGCCTTTTGCTC
ACATGTTCTTTCCTGCGTTATCCCCTGATTCTGTGGATAACCGTATTACCGCCTTTGAGTGAGCTGAT
ACCGCTCGCCGCAGCCGAACGACCGAGCGCAGCGAGTCAGTGAGCGAGGAAGCGGAAGGCGCC
CAATACGCAAACCGCCTCTCCCCGCGCGTTGGCCGATTCATTAATGCAGCTGGCACGACAGGTTTC
CCGACTGGAAAGCGGGCAGTGAGCGCAACGCAATTAAGGTGAGTTAGCTCACTCATTAGGCACCCC
AGGCTTTACACTTTATGCTTCCGGCTCGTATGTTGTGTGGAATTGTGAGCGGATAACAATTTCACACA
GATATCTAATACGACTCACTATAAGAGGAAATAAGAGAGAAAAGAAGAGTAAGAAGAAATATAAGAGC
CACCATGGTAAGTTTAAACGCTGCCTTCTGCGGGGCTTGCCTTCTGGCCATGCCCTTCTTCTCTCCC
TTGCACCTGTACCTCTTGGTCTTTGAATAAAGCCTGAGTAGGAAAAAAAAAAAAAAAAAAAAAAAAAA
AAAAAAAAAAAAAAAAAAAAAAAAAAAAAAAAAAGAAAACGAAGAGCCAGGTGGCACTTTTCGGGGA
AATGTGCGCGGAACCCCTATTTGTTTATTTTTCTAAATACATTCAAATATGTATCCGCTCATGAGACAAT
AACCCTGATAAATGCTTCAATAATATTGAAAAAGGAAGAGTATGAGTATTCAACATTTCCGTGTCGCCC
TTATTCCCTTTTTTGCGGCATTTTGCCTTCCTGTTTTTGCTCACCCAGAAA

P17:

>P2210 pIVT-T7-Nco-U3.Hba-A90(Gaaaac)

CGCTGGTGAAAGTAAAAGATGCTGAAGATCAGTTGGGTGCACGAGTGGGTTACATCGAACTGGATC
TCAACAGCGGTAAGATCCTTGAGAGTTTTCGCCCCGAAGAACGTTTTCCAATGATGAGCACTTTTAA
AGTTCTGCTATGTGGCGCGGTATTATCCCGTATTGACGCCGGGCAAGAGCAACTCGGTCGCCGCAT
ACACTATTCTCAGAATGACTTGGTTGAGTACTCACCAGTCACAGAAAAGCATCTTACGGATGGCATG
ACAGTAAGAGAATTATGCAGTGCTGCCATAACCATGAGTGATAACACTGCGGCCAACTTACTTCTGA
CAACGATCGGAGGACCGAAGGAGCTAACCGCTTTTTTGCACAACATGGGGGATCATGTAACTCGCC
TTGATCGTTGGGAACCGGAGCTGAATGAAGCCATACCAAACGACGAGCGTGACACCACGATGCCTG
TAGCAATGGCAACAACGTTGCGCAAACTATTAACTGGCGAACTACTTACTCTAGCTTCCCGGCAACA
ATTAATAGACTGGATGGAGGCGGATAAAGTTGCAGGACCACTTCTGCGCTCGGCCCTTCCGGCTGG
CTGGTTTATTGCTGATAAATCTGGAGCCGGTGAGCGTGGGTCTCGCGGTATCATTGCAGCACTGGG
GCCAGATGGTAAGCCCTCCCGTATCGTAGTTATCTACACGACGGGGAGTCAGGCAACTATGGATGAA
CGAAATAGACAGATCGCTGAGATAGGTGCCTCACTGATTAAGCATTGGTAACTGTCAGACCAAGTTT
ACTCATATATACTTTAGATTGATTTAAAACTTCATTTTTAATTTAAAAGGATCTAGGTGAAGATCCTTTTT
GATAATCTCATGACCAAAATCCCTTAACGTGAGTTTTCGTTCCACTGAGCGTCAGACCCCGTAGAAA
AGATCAAAGGATCTTCTTGAGATCCTTTTTTTCTGCGCGTAATCTGCTGCTTGCAAACAAAAAAACCA
CCGCTACCAGCGGTGGTTTGTTTGCCGGATCAAGAGCTACCAACTCTTTTTCCGAAGGTAACTGGC
TTCAGCAGAGCGCAGATACCAAATACTGTTCTTCTAGTGTAGCCGTAGTTAGGCCACCACTTCAAGA
ACTCTGTAGCACCGCCTACATACCTCGCTCTGCTAATCCTGTTACCAGTGGCTGCTGCCAGTGGCGA
TAAGTCGTGTCTTACCGGGTTGGACTCAAGACGATAGTTACCGGATAAGGCGCAGCGGTCGGGCTG
AACGGGGGGTTCGTGCACACAGCCCAGCTTGGAGCGAACGACCTACACCGAACTGAGATACCTAC
AGCGTGAGCTATGAGAAAGCGCCACGCTTCCCGAAGGGAGAAAGGCGGACAGGTATCCGGTAAGC
GGCAGGGTCGGAACAGGAGAGCGCACGAGGGAGCTTCCAGGGGGAAACGCCTGGTATCTTTATAG
TCCTGTCGGGTTTCGCCACCTCTGACTTGAGCGTCGATTTTTGTGATGCTCGTCAGGGGGGCGGA
GCCTATGGAAAAACGCCAGCAACGCGGCCTTTTTACGGTTCCTGGCCTTTTGCTGGCCTTTTGCTC
ACATGTTCTTTCCTGCGTTATCCCCTGATTCTGTGGATAACCGTATTACCGCCTTTGAGTGAGCTGAT
ACCGCTCGCCGCAGCCGAACGACCGAGCGCAGCGAGTCAGTGAGCGAGGAAGCGGAAGGCGCC
CAATACGCAAACCGCCTCTCCCCGCGCGTTGGCCGATTCATTAATGCAGCTGGCACGACAGGTTTC
CCGACTGGAAAGCGGGCAGTGAGCGCAACGCAATTAAGGTGAGTTAGCTCACTCATTAGGCACCCC
AGGCTTTACACTTTATGCTTCCGGCTCGTATGTTGTGTGGAATTGTGAGCGGATAACAATTTCACACA
GATATCTAATACGACTCACTATAAGAGGAAATAAGAGAGAAAAGAAGAGTAAGAAGAAATATAAGAGC
CACCATGGTAAGTTTAAACGCTGCCTTCTGCGGGGCTTGCCTTCTGGCCATGCCCTTCTTCTCTCCC
TTGCACCTGTACCTCTTGGTCTTTGAATAAAGCCTGAGTAGGAAAAAAAAAAAAAAAAAAAAAAAAAA
AAAAAAAAAAAAAAAAAAAAAAAAAAAAAAAAAAAAAAAAAAAAAAAAAAAAAAAAAAAAAAAAGAAAA
CGAAGAGCCAGGTGGCACTTTTCGGGGAAATGTGCGCGGAACCCCTATTTGTTTATTTTTCTAAATA
CATTCAAATATGTATCCGCTCATGAGACAATAACCCTGATAAATGCTTCAATAATATTGAAAAAGGAAG
AGTATGAGTATTCAACATTTCCGTGTCGCCCTTATTCCCTTTTTTGCGGCATTTTGCCTTCCTGTTTTT
GCTCACCCAGAAA

P18:

>P2210 pIVT-T7-Nco-U3.Hba-A120(Gaaaac)

CGCTGGTGAAAGTAAAAGATGCTGAAGATCAGTTGGGTGCACGAGTGGGTTACATCGAACTGGATC
TCAACAGCGGTAAGATCCTTGAGAGTTTTCGCCCCGAAGAACGTTTTCCAATGATGAGCACTTTTAA
AGTTCTGCTATGTGGCGCGGTATTATCCCGTATTGACGCCGGGCAAGAGCAACTCGGTCGCCGCAT
ACACTATTCTCAGAATGACTTGGTTGAGTACTCACCAGTCACAGAAAAGCATCTTACGGATGGCATG
ACAGTAAGAGAATTATGCAGTGCTGCCATAACCATGAGTGATAACACTGCGGCCAACTTACTTCTGA
CAACGATCGGAGGACCGAAGGAGCTAACCGCTTTTTTGCACAACATGGGGGATCATGTAACTCGCC
TTGATCGTTGGGAACCGGAGCTGAATGAAGCCATACCAAACGACGAGCGTGACACCACGATGCCTG
TAGCAATGGCAACAACGTTGCGCAAACTATTAACTGGCGAACTACTTACTCTAGCTTCCCGGCAACA
ATTAATAGACTGGATGGAGGCGGATAAAGTTGCAGGACCACTTCTGCGCTCGGCCCTTCCGGCTGG
CTGGTTTATTGCTGATAAATCTGGAGCCGGTGAGCGTGGGTCTCGCGGTATCATTGCAGCACTGGG
GCCAGATGGTAAGCCCTCCCGTATCGTAGTTATCTACACGACGGGGAGTCAGGCAACTATGGATGAA
CGAAATAGACAGATCGCTGAGATAGGTGCCTCACTGATTAAGCATTGGTAACTGTCAGACCAAGTTT
ACTCATATATACTTTAGATTGATTTAAAACTTCATTTTTAATTTAAAAGGATCTAGGTGAAGATCCTTTTT
GATAATCTCATGACCAAAATCCCTTAACGTGAGTTTTCGTTCCACTGAGCGTCAGACCCCGTAGAAA
AGATCAAAGGATCTTCTTGAGATCCTTTTTTTCTGCGCGTAATCTGCTGCTTGCAAACAAAAAAACCA
CCGCTACCAGCGGTGGTTTGTTTGCCGGATCAAGAGCTACCAACTCTTTTTCCGAAGGTAACTGGC
TTCAGCAGAGCGCAGATACCAAATACTGTTCTTCTAGTGTAGCCGTAGTTAGGCCACCACTTCAAGA
ACTCTGTAGCACCGCCTACATACCTCGCTCTGCTAATCCTGTTACCAGTGGCTGCTGCCAGTGGCGA
TAAGTCGTGTCTTACCGGGTTGGACTCAAGACGATAGTTACCGGATAAGGCGCAGCGGTCGGGCTG
AACGGGGGGTTCGTGCACACAGCCCAGCTTGGAGCGAACGACCTACACCGAACTGAGATACCTAC
AGCGTGAGCTATGAGAAAGCGCCACGCTTCCCGAAGGGAGAAAGGCGGACAGGTATCCGGTAAGC
GGCAGGGTCGGAACAGGAGAGCGCACGAGGGAGCTTCCAGGGGGAAACGCCTGGTATCTTTATAG
TCCTGTCGGGTTTCGCCACCTCTGACTTGAGCGTCGATTTTTGTGATGCTCGTCAGGGGGGCGGA
GCCTATGGAAAAACGCCAGCAACGCGGCCTTTTTACGGTTCCTGGCCTTTTGCTGGCCTTTTGCTC
ACATGTTCTTTCCTGCGTTATCCCCTGATTCTGTGGATAACCGTATTACCGCCTTTGAGTGAGCTGAT
ACCGCTCGCCGCAGCCGAACGACCGAGCGCAGCGAGTCAGTGAGCGAGGAAGCGGAAGGCGCC
CAATACGCAAACCGCCTCTCCCCGCGCGTTGGCCGATTCATTAATGCAGCTGGCACGACAGGTTTC
CCGACTGGAAAGCGGGCAGTGAGCGCAACGCAATTAAGGTGAGTTAGCTCACTCATTAGGCACCCC
AGGCTTTACACTTTATGCTTCCGGCTCGTATGTTGTGTGGAATTGTGAGCGGATAACAATTTCACACA
GATATCTAATACGACTCACTATAAGAGGAAATAAGAGAGAAAAGAAGAGTAAGAAGAAATATAAGAGC
CACCATGGTAAGTTTAAACGCTGCCTTCTGCGGGGCTTGCCTTCTGGCCATGCCCTTCTTCTCTCCC
TTGCACCTGTACCTCTTGGTCTTTGAATAAAGCCTGAGTAGGAAAAAAAAAAAAAAAAAAAAAAAAAA
AAAAAAAAAAAAAAAAAAAAAAAAAAAAAAAAAAAAAAAAAAAAAAAAAAAAAAAAAAAAAAAAAAAAA
AAAAAAAAAAAAAAAAAAAAAAAAAGAAAACGAAGAGCCAGGTGGCACTTTTCGGGGAAATGTGCG
CGGAACCCCTATTTGTTTATTTTTCTAAATACATTCAAATATGTATCCGCTCATGAGACAATAACCCTGA
TAAATGCTTCAATAATATTGAAAAAGGAAGAGTATGAGTATTCAACATTTCCGTGTCGCCCTTATTCCC
TTTTTTGCGGCATTTTGCCTTCCTGTTTTTGCTCACCCAGAAA

P19:

>P121 pAdHM4-CMV-M2K

TTGAGTGAGCTGATACCGCTCGCCGCAGCCGAACGACCGAGCGCAGCGAGTCAGTGAGCGAGGA
AGCGGAAGAGCGCCCAATACGCAAACCGCCTCTCCCCGCGCGTTGGCCGATTCATTAATGCAGCTG
GCACGACAGGTTTCCCGACTGGAAAGCGGGCAGTGAGCGCAACGCAATTAATGTGAGTTAGCTCA
CTCATTAGGCACCCCAGGCTTTACACTTTATGCTTCCGGCTCGTATGTTGTGTGGAATTGTGAGCGG
ATAACAATTTCACACAGGAAACAGCTATGACCATGATTACGCCAAGCTATTTAGGTGACACTATAGAAT
ACTCAAGCTAGTTAATTAACGTTAATTAACATCATCAATAATATACCTTATTTTGGATTGAAGCCAATATG
ATAATGAGGGGGTGGAGTTTGTGACGTGGCGCGGGGCGTGGGAACGGGGCGGGTGACGTAGTAG
TGTGGCGGAAGTGTGATGTTGCAAGTGTGGCGGAACACATGTAAGCGACGGATGTGGCAAAAGTG
ACGTTTTTGGTGTGCGCCGGTGTACACAGGAAGTGACAATTTTCGCGCGGTTTTAGGCGGATGTTG
TAGTAAATTTGGGCGTAACCGAGTAAGATTTGGCCATTTTCGCGGGAAAACTGAATAAGAGGAAGTG
AAATCTGAATAATTTTGTGTTACTCATAGCGCGTAATCTCTAGCATCGTAACTATAACGGTCCTAAGGT
AGCGAGAATTATTTAAATATCTATGTCGGGTGCGGAGAAAGAGGTAATGAAATGGCATCGACTCGAA
GATCTGGGCGTGGTTAAGGGTGGGAAAGAATATATAAGGTGGGGGTCTTATGTAGTTTTGTATCTGTT
TTGCAGCAGCCGCCGCCGCCATGAGCACCAACTCGTTTGATGGAAGCATTGTGAGCTCATATTTGA
CAACGCGCATGCCCCCATGGGCCGGGGTGCGTCAGAATGTGATGGGCTCCAGCATTGATGGTCGC
CCCGTCCTGCCCGCAAACTCTACTACCTTGACCTACGAGACCGTGTCTGGAACGCCGTTGGAGACT
GCAGCCTCCGCCGCCGCTTCAGCCGCTGCAGCCACCGCCCGCGGGATTGTGACTGACTTTGCTTT
CCTGAGCCCGCTTGCAAGCAGTGCAGCTTCCCGTTCATCCGCCCGCGATGACAAGTTGACGGCTC
TTTTGGCACAATTGGATTCTTTGACCCGGGAACTTAATGTCGTTTCTCAGCAGCTGTTGGATCTGCG
CCAGCAGGTTTCTGCCCTGAAGGCTTCCTCCCCTCCCAATGCGGTTTAAAACATAAATAAAAAACCA
GACTCTGTTTGGATTTGGATCAAGCAAGTGTCTTGCTGTCTTTATTTAGGGGTTTTGCGCGCGCGGT
AGGCCCGGGACCAGCGGTCTCGGTCGTTGAGGGTCCTGTGTATTTTTTCCAGGACGTGGTAAAGG
TGACTCTGGATGTTCAGATACATGGGCATAAGCCCGTCTCTGGGGTGGAGGTAGCACCACTGCAGA
GCTTCATGCTGCGGGGTGGTGTTGTAGATGATCCAGTCGTAGCAGGAGCGCTGGGCGTGGTGCCT
AAAAATGTCTTTCAGTAGCAAGCTGATTGCCAGGGGCAGGCCCTTGGTGTAAGTGTTTACAAAGCG
GTTAAGCTGGGATGGGTGCATACGTGGGGATATGAGATGCATCTTGGACTGTATTTTTAGGTTGGCTA
TGTTCCCAGCCATATCCCTCCGGGGATTCATGTTGTGCAGAACCACCAGCACAGTGTATCCGGTGC
ACTTGGGAAATTTGTCATGTAGCTTAGAAGGAAATGCGTGGAAGAACTTGGAGACGCCCTTGTGAC
CTCCAAGATTTTCCATGCATTCGTCCATAATGATGGCAATGGGCCCACGGGCGGCGGCCTGGGCGA
AGATATTTCTGGGATCACTAACGTCATAGTTGTGTTCCAGGATGAGATCGTCATAGGCCATTTTTACAA
AGCGCGGGCGGAGGGTGCCAGACTGCGGTATAATGGTTCCATCCGGCCCAGGGGCGTAGTTACCC
TCACAGATTTGCATTTCCCACGCTTTGAGTTCAGATGGGGGGATCATGTCTACCTGCGGGGCGATG
AAGAAAACGGTTTCCGGGGTAGGGGAGATCAGCTGGGAAGAAAGCAGGTTCCTGAGCAGCTGCGA
CTTACCGCAGCCGGTGGGCCCGTAAATCACACCTATTACCGGCTGCAACTGGTAGTTAAGAGAGCT
GCAGCTGCCGTCATCCCTGAGCAGGGGGGCCACTTCGTTAAGCATGTCCCTGACTCGCATGTTTTC
CCTGACCAAATCCGCCAGAAGGCGCTCGCCGCCCAGCGATAGCAGTTCTTGCAAGGAAGCAAAGT
TTTTCAACGGTTTGAGACCGTCCGCCGTAGGCATGCTTTTGAGCGTTTGACCAAGCAGTTCCAGGC
GGTCCCACAGCTCGGTCACCTGCTCTACGGCATCTCGATCCAGCATATCTCCTCGTTTCGCGGGTT
GGGGCGGCTTTCGCTGTACGGCAGTAGTCGGTGCTCGTCCAGACGGGCCAGGGTCATGTCTTTCC
ACGGGCGCAGGGTCCTCGTCAGCGTAGTCTGGGTCACGGTGAAGGGGTGCGCTCCGGGCTGCGC
GCTGGCCAGGGTGCGCTTGAGGCTGGTCCTGCTGGTGCTGAAGCGCTGCCGGTCTTCGCCCTGC
GCGTCGGCCAGGTAGCATTTGACCATGGTGTCATAGTCCAGCCCCTCCGCGGCGTGGCCCTTGGC

GCGCAGCTTGCCCTTGGAGGAGGCGCCGCACGAGGGGCAGTGCAGACTTTTGAGGGCGTAGAGC
TTGGGCGCGAGAAATACCGATTCCGGGGAGTAGGCATCCGCGCCGCAGGCCCCGCAGACGGTCT
CGCATTCCACGAGCCAGGTGAGCTCTGGCCGTTCGGGGTCAAAAACCAGGTTTCCCCCATGCTTTT
TGATGCGTTTCTTACCTCTGGTTTCCATGAGCCGGTGTCCACGCTCGGTGACGAAAAGGCTGTCCG
TGTCCCCGTATACAGACTTGAGAGGCCTGTCCTCGAGCGGTGTTCCGCGGTCCTCCTCGTATAGAA
ACTCGGACCACTCTGAGACAAAGGCTCGCGTCCAGGCCAGCACGAAGGAGGCTAAGTGGGAGGG
GTAGCGGTCGTTGTCCACTAGGGGGTCCACTCGCTCCAGGGTGTGAAGACACATGTCGCCCTCTT
CGGCATCAAGGAAGGTGATTGGTTTGTAGGTGTAGGCCACGTGACCGGGTGTTCCTGAAGGGGGG
CTATAAAAGGGGGTGGGGGCGCGTTCGTCCTCACTCTCTTCCGCATCGCTGTCTGCGAGGGCCAG
CTGTTGGGGTGAGTACTCCCTCTGAAAAGCGGGCATGACTTCTGCGCTAAGATTGTCAGTTTCCAA
AAACGAGGAGGATTTGATATTCACCTGGCCCGCGGTGATGCCTTTGAGGGTGGCCGCATCCATCTG
GTCAGAAAAGACAATCTTTTTGTTGTCAAGCTTGGTGGCAAACGACCCGTAGAGGGCGTTGGACAG
CAACTTGGCGATGGAGCGCAGGGTTTGGTTTTTGTCGCGATCGGCGCGCTCCTTGGCCGCGATGT
TTAGCTGCACGTATTCGCGCGCAACGCACCGCCATTCGGGAAAGACGGTGGTGCGCTCGTCGGGC
ACCAGGTGCACGCGCCAACCGCGGTTGTGCAGGGTGACAAGGTCAACGCTGGTGGCTACCTCTC
CGCGTAGGCGCTCGTTGGTCCAGCAGAGGCGGCCGCCCTTGCGCGAGCAGAATGGCGGTAGGGG
GTCTAGCTGCGTCTCGTCCGGGGGGTCTGCGTCCACGGTAAAGACCCCGGGCAGCAGGCGCGCG
TCGAAGTAGTCTATCTTGCATCCTTGCAAGTCTAGCGCCTGCTGCCATGCGCGGGCGGCAAGCGCG
CGCTCGTATGGGTTGAGTGGGGGACCCCATGGCATGGGGTGGGTGAGCGCGGAGGCGTACATGC
CGCAAATGTCGTAAACGTAGAGGGGCTCTCTGAGTATTCCAAGATATGTAGGGTAGCATCTTCCACC
GCGGATGCTGGCGCGCACGTAATCGTATAGTTCGTGCGAGGGAGCGAGGAGGTCGGGACCGAGG
TTGCTACGGGCGGGCTGCTCTGCTCGGAAGACTATCTGCCTGAAGATGGCATGTGAGTTGGATGAT
ATGGTTGGACGCTGGAAGACGTTGAAGCTGGCGTCTGTGAGACCTACCGCGTCACGCACGAAGGA
GGCGTAGGAGTCGCGCAGCTTGTTGACCAGCTCGGCGGTGACCTGCACGTCTAGGGCGCAGTAGT
CCAGGGTTTCCTTGATGATGTCATACTTATCCTGTCCCTTTTTTTTCCACAGCTCGCGGTTGAGGACA
AACTCTTCGCGGTCTTTCCAGTACTCTTGGATCGGAAACCCGTCGGCCTCCGAACGGTAAGAGCCT
AGCATGTAGAACTGGTTGACGGCCTGGTAGGCGCAGCATCCCTTTTCTACGGGTAGCGCGTATGCC
TGCGCGGCCTTCCGGAGCGAGGTGTGGGTGAGCGCAAAGGTGTCCCTGACCATGACTTTGAGGTA
CTGGTATTTGAAGTCAGTGTCGTCGCATCCGCCCTGCTCCCAGAGCAAAAAGTCCGTGCGCTTTTT
GGAACGCGGATTTGGCAGGGCGAAGGTGACATCGTTGAAGAGTATCTTTCCCGCGCGAGGCATAAA
GTTGCGTGTGATGCGGAAGGGTCCCGGCACCTCGGAACGGTTGTTAATTACCTGGGCGGCGAGCA
CGATCTCGTCAAAGCCGTTGATGTTGTGGCCCACAATGTAAAGTTCCAAGAAGCGCGGGATGCCCT
TGATGGAAGGCAATTTTTTAAGTTCCTCGTAGGTGAGCTCTTCAGGGGAGCTGAGCCCGTGCTCTG
AAAGGGCCCAGTCTGCAAGATGAGGGTTGGAAGCGACGAATGAGCTCCACAGGTCACGGGCCATT
AGCATTTGCAGGTGGTCGCGAAAGGTCCTAAACTGGCGACCTATGGCCATTTTTTCTGGGGTGATG
CAGTAGAAGGTAAGCGGGTCTTGTTCCCAGCGGTCCCATCCAAGGTTCGCGGCTAGGTCTCGCGC
GGCAGTCACTAGAGGCTCATCTCCGCCGAACTTCATGACCAGCATGAAGGGCACGAGCTGCTTCC
CAAAGGCCCCCATCCAAGTATAGGTCTCTACATCGTAGGTGACAAAGAGACGCTCGGTGCGAGGAT
GCGAGCCGATCGGGAAGAACTGGATCTCCCGCCACCAATTGGAGGAGTGGCTATTGATGTGGTGA
AAGTAGAAGTCCCTGCGACGGGCCGAACACTCGTGCTGGCTTTTGTAAAAACGTGCGCAGTACTGG
CAGCGGTGCACGGGCTGTACATCCTGCACGAGGTTGACCTGACGACCGCGCACAAGGAAGCAGA
GTGGGAATTTGAGCCCCTCGCCTGGCGGGTTTGGCTGGTGGTCTTCTACTTCGGCTGCTTGTCCTT
GACCGTCTGGCTGCTCGAGGGGAGTTACGGTGGATCGGACCACCACGCCGCGCGAGCCCAAAGT
CCAGATGTCCGCGCGCGGCGGTCGGAGCTTGATGACAACATCGCGCAGATGGGAGCTGTCCATGG

TCTGGAGCTCCCGCGGCGTCAGGTCAGGCGGGAGCTCCTGCAGGTTTACCTCGCATAGACGGGTC
AGGGCGCGGGCTAGATCCAGGTGATACCTAATTTCCAGGGGCTGGTTGGTGGCGGCGTCGATGGC
TTGCAAGAGGCCGCATCCCCGCGGCGCGACTACGGTACCGCGCGGCGGGCGGTGGGCCGCGGG
GGTGTCCTTGGATGATGCATCTAAAAGCGGTGACGCGGGCGAGCCCCCGGAGGTAGGGGGGGCT
CCGGACCCGCCGGGAGAGGGGGCAGGGGCACGTCGGCGCCGCGCGCGGGCAGGAGCTGGTGC
TGCGCGCGTAGGTTGCTGGCGAACGCGACGACGCGGCGGTTGATCTCCTGAATCTGGCGCCTCTG
CGTGAAGACGACGGGCCCGGTGAGCTTGAACCTGAAAGAGAGTTCGACAGAATCAATTTCGGTGT
CGTTGACGGCGGCCTGGCGCAAAATCTCCTGCACGTCTCCTGAGTTGTCTTGATAGGCGATCTCGG
CCATGAACTGCTCGATCTCTTCCTCCTGGAGATCTCCGCGTCCGGCTCGCTCCACGGTGGCGGCG
AGGTCGTTGGAAATGCGGGCCATGAGCTGCGAGAAGGCGTTGAGGCCTCCCTCGTTCCAGACGCG
GCTGTAGACCACGCCCCCTTCGGCATCGCGGGCGCGCATGACCACCTGCGCGAGATTGAGCTCCA
CGTGCCGGGCGAAGACGGCGTAGTTTCGCAGGCGCTGAAAGAGGTAGTTGAGGGTGGTGGCGGT
GTGTTCTGCCACGAAGAAGTACATAACCCAGCGTCGCAACGTGGATTCGTTGATATCCCCCAAGGC
CTCAAGGCGCTCCATGGCCTCGTAGAAGTCCACGGCGAAGTTGAAAAACTGGGAGTTGCGCGCCG
ACACGGTTAACTCCTCCTCCAGAAGACGGATGAGCTCGGCGACAGTGTCGCGCACCTCGCGCTCA
AAGGCTACAGGGGCCTCTTCTTCTTCTTCAATCTCCTCTTCCATAAGGGCCTCCCCTTCTTCTTCTTC
TGGCGGCGGTGGGGGAGGGGGGACACGGCGGCGACGACGGCGCACCGGGAGGCGGTCGACAA
AGCGCTCGATCATCTCCCCGCGGCGACGGCGCATGGTCTCGGTGACGGCGCGGCCGTTCTCGCG
GGGGCGCAGTTGGAAGACGCCGCCCGTCATGTCCCGGTTATGGGTTGGCGGGGGGCTGCCATGC
GGCAGGGATACGGCGCTAACGATGCATCTCAACAATTGTTGTGTAGGTACTCCGCCGCCGAGGGAC
CTGAGCGAGTCCGCATCGACCGGATCGGAAAACCTCTCGAGAAAGGCGTCTAACCAGTCACAGTC
GCAAGGTAGGCTGAGCACCGTGGCGGGCGGCAGCGGGCGGCGGTCGGGGTTGTTTCTGGCGGA
GGTGCTGCTGATGATGTAATTAAAGTAGGCGGTCTTGAGACGGCGGATGGTCGACAGAAGCACCAT
GTCCTTGGGTCCGGCCTGCTGAATGCGCAGGCGGTCGGCCATGCCCCAGGCTTCGTTTTGACATC
GGCGCAGGTCTTTGTAGTAGTCTTGCATGAGCCTTTCTACCGGCACTTCTTCTTCTCCTTCCTCTTG
TCCTGCATCTCTTGCATCTATCGCTGCGGCGGCGGCGGAGTTTGGCCGTAGGTGGCGCCCTCTTC
CTCCCATGCGTGTGACCCCGAAGCCCCTCATCGGCTGAAGCAGGGCTAGGTCGGCGACAACGCG
CTCGGCTAATATGGCCTGCTGCACCTGCGTGAGGGTAGACTGGAAGTCATCCATGTCCACAAAGCG
GTGGTATGCGCCCGTGTTGATGGTGTAAGTGCAGTTGGCCATAACGGACCAGTTAACGGTCTGGTG
ACCCGGCTGCGAGAGCTCGGTGTACCTGAGACGCGAGTAAGCCCTCGAGTCAAATACGTAGTCGTT
GCAAGTCCGCACCAGGTACTGGTATCCCACCAAAAAGTGCGGCGGCGGCTGGCGGTAGAGGGGC
CAGCGTAGGGTGGCCGGGGCTCCGGGGGCGAGATCTTCCAACATAAGGCGATGATATCCGTAGAT
GTACCTGGACATCCAGGTGATGCCGGCGGCGGTGGTGGAGGCGCGCGGAAAGTCGCGGACGCG
GTTCCAGATGTTGCGCAGCGGCAAAAAGTGCTCCATGGTCGGGACGCTCTGGCCGGTCAGGCGC
GCGCAATCGTTGACGCTCTAGCGTGCAAAAGGAGAGCCTGTAAGCGGGCACTCTTCCGTGGTCTG
GTGGATAAATTCGCAAGGGTATCATGGCGGACGACCGGGGTTCGAGCCCCGTATCCGGCCGTCCG
CCGTGATCCATGCGGTTACCGCCCGCGTGTCGAACCCAGGTGTGCGACGTCAGACAACGGGGGA
GTGCTCCTTTTGGCTTCCTTCCAGGCGCGGCGGCTGCTGCGCTAGCTTTTTTGGCCACTGGCCGC
GCGCAGCGTAAGCGGTTAGGCTGGAAAGCGAAAGCATTAAGTGGCTCGCTCCCTGTAGCCGGAGG
GTTATTTTCCAAGGGTTGAGTCGCGGGACCCCCGGTTCGAGTCTCGGACCGGCCGGACTGCGGCG
AACGGGGGTTTGCCTCCCCGTCATGCAAGACCCCGCTTGCAAATTCCTCCGGAAACAGGGACGAG
CCCCTTTTTTGCTTTTCCCAGATGCATCCGGTGCTGCGGCAGATGCGCCCCCCTCCTCAGCAGCGG
CAAGAGCAAGAGCAGCGGCAGACATGCAGGGCACCCTCCCCTCCTCCTACCGCGTCAGGAGGGG
CGACATCCGCGGTTGACGCGGCAGCAGATGGTGATTACGAACCCCCGCGGCGCCGGGCCCGGCA

CTACCTGGACTTGGAGGAGGGCGAGGGCCTGGCGCGGCTAGGAGCGCCCTCTCCTGAGCGGCAC
CCAAGGGTGCAGCTGAAGCGTGATACGCGTGAGGCGTACGTGCCGCGGCAGAACCTGTTTCGCGA
CCGCGAGGGAGAGGAGCCCGAGGAGATGCGGGATCGAAAGTTCCACGCAGGGCGCGAGCTGCG
GCATGGCCTGAATCGCGAGCGGTTGCTGCGCGAGGAGGACTTTGAGCCCGACGCGCGAACCGGG
ATTAGTCCCGCGCGCGCACACGTGGCGGCCGCCGACCTGGTAACCGCATACGAGCAGACGGTGAA
CCAGGAGATTAACTTTCAAAAAAGCTTTAACAACCACGTGCGTACGCTTGTGGCGCGCGAGGAGGT
GGCTATAGGACTGATGCATCTGTGGGACTTTGTAAGCGCGCTGGAGCAAAACCCAAATAGCAAGCC
GCTCATGGCGCAGCTGTTCCTTATAGTGCAGCACAGCAGGGACAACGAGGCATTCAGGGATGCGCT
GCTAAACATAGTAGAGCCCGAGGGCCGCTGGCTGCTCGATTTGATAAACATCCTGCAGAGCATAGT
GGTGCAGGAGCGCAGCTTGAGCCTGGCTGACAAGGTGGCCGCCATCAACTATTCCATGCTTAGCCT
GGGCAAGTTTTACGCCCGCAAGATATACCATACCCCTTACGTTCCCATAGACAAGGAGGTAAAGATC
GAGGGGTTCTACATGCGCATGGCGCTGAAGGTGCTTACCTTGAGCGACGACCTGGGCGTTTATCGC
AACGAGCGCATCCACAAGGCCGTGAGCGTGAGCCGGCGGCGCGAGCTCAGCGACCGCGAGCTGA
TGCACAGCCTGCAAAGGGCCCTGGCTGGCACGGGCAGCGGCGATAGAGAGGCCGAGTCCTACTT
TGACGCGGGCGCTGACCTGCGCTGGGCCCCAAGCCGACGCGCCCTGGAGGCAGCTGGGGCCGG
ACCTGGGCTGGCGGTGGCACCCGCGCGCGCTGGCAACGTCGGCGGCGTGGAGGAATATGACGAG
GACGATGAGTACGAGCCAGAGGACGGCGAGTACTAAGCGGTGATGTTTCTGATCAGATGATGCAAG
ACGCAACGGACCCGGCGGTGCGGGCGGCGCTGCAGAGCCAGCCGTCCGGCCTTAACTCCACGG
ACGACTGGCGCCAGGTCATGGACCGCATCATGTCGCTGACTGCGCGCAATCCTGACGCGTTCCGG
CAGCAGCCGCAGGCCAACCGGCTCTCCGCAATTCTGGAAGCGGTGGTCCCGGCGCGCGCAAACC
CCACGCACGAGAAGGTGCTGGCGATCGTAAACGCGCTGGCCGAAAACAGGGCCATCCGGCCCGA
CGAGGCCGGCCTGGTCTACGACGCGCTGCTTCAGCGCGTGGCTCGTTACAACAGCGGCAACGTG
CAGACCAACCTGGACCGGCTGGTGGGGGATGTGCGCGAGGCCGTGGCGCAGCGTGAGCGCGCG
CAGCAGCAGGGCAACCTGGGCTCCATGGTTGCACTAAACGCCTTCCTGAGTACACAGCCCGCCAA
CGTGCCGCGGGGACAGGAGGACTACACCAACTTTGTGAGCGCACTGCGGCTAATGGTGACTGAGA
CACCGCAAAGTGAGGTGTACCAGTCTGGGCCAGACTATTTTTTCCAGACCAGTAGACAAGGCCTGC
AGACCGTAAACCTGAGCCAGGCTTTCAAAAACTTGCAGGGGCTGTGGGGGGTGCGGGCTCCCACA
GGCGACCGCGCGACCGTGTCTAGCTTGCTGACGCCCAACTCGCGCCTGTTGCTGCTGCTAATAGC
GCCCTTCACGGACAGTGGCAGCGTGTCCCGGGACACATACCTAGGTCACTTGCTGACACTGTACC
GCGAGGCCATAGGTCAGGCGCATGTGGACGAGCATACTTTCCAGGAGATTACAAGTGTCAGCCGC
GCGCTGGGGCAGGAGGACACGGGCAGCCTGGAGGCAACCCTAAACTACCTGCTGACCAACCGGC
GGCAGAAGATCCCCTCGTTGCACAGTTTAAACAGCGAGGAGGAGCGCATTTTGCGCTACGTGCAG
CAGAGCGTGAGCCTTAACCTGATGCGCGACGGGGTAACGCCCAGCGTGGCGCTGGACATGACCG
CGCGCAACATGGAACCGGGCATGTATGCCTCAAACCGGCCGTTTATCAACCGCCTAATGGACTACTT
GCATCGCGCGGCCGCCGTGAACCCCGAGTATTTCACCAATGCCATCTTGAACCCGCACTGGCTACC
GCCCCCTGGTTTCTACACCGGGGGATTCGAGGTGCCCGAGGGTAACGATGGATTCCTCTGGGACG
ACATAGACGACAGCGTGTTTTCCCCGCAACCGCAGACCCTGCTAGAGTTGCAACAGCGCGAGCAG
GCAGAGGCGGCGCTGCGAAAGGAAAGCTTCCGCAGGCCAAGCAGCTTGTCCGATCTAGGCGCTG
CGGCCCCGCGGTCAGATGCTAGTAGCCCATTTCCAAGCTTGATAGGGTCTCTTACCAGCACTCGCA
CCACCCGCCCGCGCCTGCTGGGCGAGGAGGAGTACCTAAACAACTCGCTGCTGCAGCCGCAGCG
CGAAAAAAACCTGCCTCCGGCATTTCCCAACAACGGGATAGAGAGCCTAGTGGACAAGATGAGTAG
ATGGAAGACGTACGCGCAGGAGCACAGGGACGTGCCAGGCCCGCGCCCGCCCACCCGTCGTCAA
AGGCACGACCGTCAGCGGGGTCTGGTGTGGGAGGACGATGACTCGGCAGACGACAGCAGCGTCC
TGGATTTGGGAGGGAGTGGCAACCCGTTTGCGCACCTTCGCCCCAGGCTGGGGAGAATGTTTTAA

AAAAAAAAAAAGCATGATGCAAAATAAAAAACTCACCAAGGCCATGGCACCGAGCGTTGGTTTTCTT
GTATTCCCCTTAGTATGCGGCGCGCGGCGATGTATGAGGAAGGTCCTCCTCCCTCCTACGAGAGTG
TGGTGAGCGCGGCGCCAGTGGCGGCGGCGCTGGGTTCTCCCTTCGATGCTCCCCTGGACCCGCC
GTTTGTGCCTCCGCGGTACCTGCGGCCTACCGGGGGGAGAAACAGCATCCGTTACTCTGAGTTGG
CACCCCTATTCGACACCACCCGTGTGTACCTGGTGGACAACAAGTCAACGGATGTGGCATCCCTGA
ACTACCAGAACGACCACAGCAACTTTCTGACCACGGTCATTCAAAACAATGACTACAGCCCGGGGG
AGGCAAGCACACAGACCATCAATCTTGACGACCGGTCGCACTGGGGCGGCGACCTGAAAACCATC
CTGCATACCAACATGCCAAATGTGAACGAGTTCATGTTTACCAATAAGTTTAAGGCGCGGGTGATGG
TGTCGCGCTTGCCTACTAAGGACAATCAGGTGGAGCTGAAATACGAGTGGGTGGAGTTCACGCTGC
CCGAGGGCAACTACTCCGAGACCATGACCATAGACCTTATGAACAACGCGATCGTGGAGCACTACT
TGAAAGTGGGCAGACAGAACGGGGTTCTGGAAAGCGACATCGGGGTAAAGTTTGACACCCGCAAC
TTCAGACTGGGGTTTGACCCCGTCACTGGTCTTGTCATGCCTGGGGTATATACAAACGAAGCCTTCC
ATCCAGACATCATTTTGCTGCCAGGATGCGGGGTGGACTTCACCCACAGCCGCCTGAGCAACTTGT
TGGGCATCCGCAAGCGGCAACCCTTCCAGGAGGGCTTTAGGATCACCTACGATGATCTGGAGGGT
GGTAACATTCCCGCACTGTTGGATGTGGACGCCTACCAGGCGAGCTTGAAAGATGACACCGAACAG
GGCGGGGGTGGCGCAGGCGGCAGCAACAGCAGTGGCAGCGGCGCGGAAGAGAACTCCAACGCG
GCAGCCGCGGCAATGCAGCCGGTGGAGGACATGAACGATCATGCCATTCGCGGCGACACCTTTGC
CACACGGGCTGAGGAGAAGCGCGCTGAGGCCGAAGCAGCGGCCGAAGCTGCCGCCCCCGCTGC
GCAACCCGAGGTCGAGAAGCCTCAGAAGAAACCGGTGATCAAACCCCTGACAGAGGACAGCAAGA
AACGCAGTTACAACCTAATAAGCAATGACAGCACCTTCACCCAGTACCGCAGCTGGTACCTTGCATA
CAACTACGGCGACCCTCAGACCGGAATCCGCTCATGGACCCTGCTTTGCACTCCTGACGTAACCTG
CGGCTCGGAGCAGGTCTACTGGTCGTTGCCAGACATGATGCAAGACCCCGTGACCTTCCGCTCCA
CGCGCCAGATCAGCAACTTTCCGGTGGTGGGCGCCGAGCTGTTGCCCGTGCACTCCAAGAGCTTC
TACAACGACCAGGCCGTCTACTCCCAACTCATCCGCCAGTTTACCTCTCTGACCCACGTGTTCAATC
GCTTTCCCGAGAACCAGATTTTGGCGCGCCCGCCAGCCCCCACCATCACCACCGTCAGTGAAAAC
GTTCCTGCTCTCACAGATCACGGGACGCTACCGCTGCGCAACAGCATCGGAGGAGTCCAGCGAGT
GACCATTACTGACGCCAGACGCCGCACCTGCCCCTACGTTTACAAGGCCCTGGGCATAGTCTCGCC
GCGCGTCCTATCGAGCCGCACTTTTTGAGCAAGCATGTCCATCCTTATATCGCCCAGCAATAACACA
GGCTGGGGCCTGCGCTTCCCAAGCAAGATGTTTGGCGGGGCCAAGAAGCGCTCCGACCAACACC
CAGTGCGCGTGCGCGGGCACTACCGCGCGCCCTGGGGCGCGCACAAACGCGGCCGCACTGGGC
GCACCACCGTCGATGACGCCATCGACGCGGTGGTGGAGGAGGCGCGCAACTACACGCCCACGCC
GCCACCAGTGTCCACAGTGGACGCGGCCATTCAGACCGTGGTGCGCGGAGCCCGGCGCTATGCT
AAAATGAAGAGACGGCGGAGGCGCGTAGCACGTCGCCACCGCCGCCGACCCGGCACTGCCGCCC
AACGCGCGGCGGCGGCCCTGCTTAACCGCGCACGTCGCACCGGCCGACGGGCGGCCATGCGGG
CCGCTCGAAGGCTGGCCGCGGGTATTGTCACTGTGCCCCCCAGGTCCAGGCGACGAGCGGCCGC
CGCAGCAGCCGCGGCCATTAGTGCTATGACTCAGGGTCGCAGGGGCAACGTGTATTGGGTGCGCG
ACTCGGTTAGCGGCCTGCGCGTGCCCGTGCGCACCCGCCCCCCGCGCAACTAGATTGCAAGAAAA
AACTACTTAGACTCGTACTGTTGTATGTATCCAGCGGCGGCGGCGCGCAACGAAGCTATGTCCAAG
CGCAAAATCAAAGAAGAGATGCTCCAGGTCATCGCGCCGGAGATCTATGGCCCCCCGAAGAAGGAA
GAGCAGGATTACAAGCCCCGAAAGCTAAAGCGGGTCAAAAAGAAAAAGAAAGATGATGATGATGAA
CTTGACGACGAGGTGGAACTGCTGCACGCTACCGCGCCCAGGCGACGGGTACAGTGGAAAGGTC
GACGCGTAAAACGTGTTTTGCGACCCGGCACCACCGTAGTCTTTACGCCCGGTGAGCGCTCCACC
CGCACCTACAAGCGCGTGTATGATGAGGTGTACGGCGACGAGGACCTGCTTGAGCAGGCCAACGA
GCGCCTCGGGGAGTTTGCCTACGGAAAGCGGCATAAGGACATGCTGGCGTTGCCGCTGGACGAG

GGCAACCCAACACCTAGCCTAAAGCCCGTAACACTGCAGCAGGTGCTGCCCGCGCTTGCACCGTC
CGAAGAAAAGCGCGGCCTAAAGCGCGAGTCTGGTGACTTGGCACCCACCGTGCAGCTGATGGTAC
CCAAGCGCCAGCGACTGGAAGATGTCTTGGAAAAAATGACCGTGGAACCTGGGCTGGAGCCCGAG
GTCCGCGTGCGGCCAATCAAGCAGGTGGCGCCGGGACTGGGCGTGCAGACCGTGGACGTTCAGA
TACCCACTACCAGTAGCACCAGTATTGCCACCGCCACAGAGGGCATGGAGACACAAACGTCCCCGG
TTGCCTCAGCGGTGGCGGATGCCGCGGTGCAGGCGGTCGCTGCGGCCGCGTCCAAGACCTCTAC
GGAGGTGCAAACGGACCCGTGGATGTTTCGCGTTTCAGCCCCCCGGCGCCCGCGCCGTTCGAGG
AAGTACGGCGCCGCCAGCGCGCTACTGCCCGAATATGCCCTACATCCTTCCATTGCGCCTACCCCC
GGCTATCGTGGCTACACCTACCGCCCCAGAAGACGAGCAACTACCCGACGCCGAACCACCACTGG
AACCCGCCGCCGCCGTCGCCGTCGCCAGCCCGTGCTGGCCCCGATTTCCGTGCGCAGGGTGGCT
CGCGAAGGAGGCAGGACCCTGGTGCTGCCAACAGCGCGCTACCACCCCAGCATCGTTTAAAAGCC
GGTCTTTGTGGTTCTTGCAGATATGGCCCTCACCTGCCGCCTCCGTTTCCCGGTGCCGGGATTCCG
AGGAAGAATGCACCGTAGGAGGGGCATGGCCGGCCACGGCCTGACGGGCGGCATGCGTCGTGCG
CACCACCGGCGGCGGCGCGCGTCGCACCGTCGCATGCGCGGCGGTATCCTGCCCCTCCTTATTCC
ACTGATCGCCGCGGCGATTGGCGCCGTGCCCGGAATTGCATCCGTGGCCTTGCAGGCGCAGAGA
CACTGATTAAAAACAAGTTGCATGTGGAAAAATCAAAATAAAAAGTCTGGACTCTCACGCTCGCTTG
GTCCTGTAACTATTTTGTAGAATGGAAGACATCAACTTTGCGTCTCTGGCCCCGCGACACGGCTCGC
GCCCGTTCATGGGAAACTGGCAAGATATCGGCACCAGCAATATGAGCGGTGGCGCCTTCAGCTGG
GGCTCGCTGTGGAGCGGCATTAAAAATTTCGGTTCCACCGTTAAGAACTATGGCAGCAAGGCCTGG
AACAGCAGCACAGGCCAGATGCTGAGGGATAAGTTGAAAGAGCAAAATTTCCAACAAAAGGTGGTA
GATGGCCTGGCCTCTGGCATTAGCGGGGTGGTGGACCTGGCCAACCAGGCAGTGCAAAATAAGAT
TAACAGTAAGCTTGATCCCCGCCCTCCCGTAGAGGAGCCTCCACCGGCCGTGGAGACAGTGTCTC
CAGAGGGGCGTGGCGAAAAGCGTCCGCGCCCCGACAGGGAAGAAACTCTGGTGACGCAAATAGA
CGAGCCTCCCTCGTACGAGGAGGCACTAAAGCAAGGCCTGCCCACCACCCGTCCCATCGCGCCCA
TGGCTACCGGAGTGCTGGGCCAGCACACACCCGTAACGCTGGACCTGCCTCCCCCCGCCGACAC
CCAGCAGAAACCTGTGCTGCCAGGCCCGACCGCCGTTGTTGTAACCCGTCCTAGCCGCGCGTCCC
TGCGCCGCGCCGCCAGCGGTCCGCGATCGTTGCGGCCCGTAGCCAGTGGCAACTGGCAAAGCAC
ACTGAACAGCATCGTGGGTCTGGGGGTGCAATCCCTGAAGCGCCGACGATGCTTCTGATAGCTAAC
GTGTCGTATGTGTGTCATGTATGCGTCCATGTCGCCGCCAGAGGAGCTGCTGAGCCGCCGCGCGC
CCGCTTTCCAAGATGGCTACCCCTTCGATGATGCCGCAGTGGTCTTACATGCACATCTCGGGCCAG
GACGCCTCGGAGTACCTGAGCCCCGGGCTGGTGCAGTTTGCCCGCGCCACCGAGACGTACTTCA
GCCTGAATAACAAGTTTAGAAACCCCACGGTGGCGCCTACGCACGACGTGACCACAGACCGGTCC
CAGCGTTTGACGCTGCGGTTCATCCCTGTGGACCGTGAGGATACTGCGTACTCGTACAAGGCGCG
GTTCACCCTAGCTGTGGGTGATAACCGTGTGCTGGACATGGCTTCCACGTACTTTGACATCCGCGG
CGTGCTGGACAGGGGCCCTACTTTTAAGCCCTACTCTGGCACTGCCTACAACGCCCTGGCTCCCAA
GGGTGCCCCAAATCCTTGCGAATGGGATGAAGCTGCTACTGCTCTTGAAATAAACCTAGAAGAAGA
GGACGATGACAACGAAGACGAAGTAGACGAGCAAGCTGAGCAGCAAAAAACTCACGTATTTGGGCA
GGCGCCTTATTCTGGTATAAATATTACAAAGGAGGGTATTCAAATAGGTGTCGAAGGTCAAACACCTA
AATATGCCGATAAAACATTTCAACCTGAACCTCAAATAGGAGAATCTCAGTGGTACGAAACAGAAATT
AATCATGCAGCTGGGAGAGTCCTAAAAAAGACTACCCCAATGAAACCATGTTACGGTTCATATGCAAA
ACCCACAAATGAAAATGGAGGGCAAGGCATTCTTGTAAAGCAACAAAATGGAAAGCTAGAAAGTCAA
GTGGAAATGCAATTTTTCTCAACTACTGAGGCAGCCGCAGGCAATGGTGATAACTTGACTCCTAAAG
TGGTATTGTACAGTGAAGATGTAGATATAGAAACCCCAGACACTCATATTTCTTACATGCCCACTATTA
AGGAAGGTAACTCACGAGAACTAATGGGCCAACAATCTATGCCCAACAGGCCTAATTACATTGCTTTT

AGGGACAATTTTATTGGTCTAATGTATTACAACAGCACGGGTAATATGGGTGTTCTGGCGGGCCAAG
CATCGCAGTTGAATGCTGTTGTAGATTTGCAAGACAGAAACACAGAGCTTTCATACCAGCTTTTGCTT
GATTCCATTGGTGATAGAACCAGGTACTTTTCTATGTGGAATCAGGCTGTTGACAGCTATGATCCAGA
TGTTAGAATTATTGAAAATCATGGAACTGAAGATGAACTTCCAAATTACTGCTTTCCACTGGGAGGTG
TGATTAATACAGAGACTCTTACCAAGGTAAAACCTAAAACAGGTCAGGAAAATGGATGGGAAAAAGAT
GCTACAGAATTTTCAGATAAAAATGAAATAAGAGTTGGAAATAATTTTGCCATGGAAATCAATCTAAAT
GCCAACCTGTGGAGAAATTTCCTGTACTCCAACATAGCGCTGTATTTGCCCGACAAGCTAAAGTACA
GTCCTTCCAACGTAAAAATTTCTGATAACCCAAACACCTACGACTACATGAACAAGCGAGTGGTGGC
TCCCGGGCTAGTGGACTGCTACATTAACCTTGGAGCACGCTGGTCCCTTGACTATATGGACAACGTC
AACCCATTTAACCACCACCGCAATGCTGGCCTGCGCTACCGCTCAATGTTGCTGGGCAATGGTCGC
TATGTGCCCTTCCACATCCAGGTGCCTCAGAAGTTCTTTGCCATTAAAAACCTCCTTCTCCTGCCGG
GCTCATACACCTACGAGTGGAACTTCAGGAAGGATGTTAACATGGTTCTGCAGAGCTCCCTAGGAAA
TGACCTAAGGGTTGACGGAGCCAGCATTAAGTTTGATAGCATTTGCCTTTACGCCACCTTCTTCCCC
ATGGCCCACAACACCGCCTCCACGCTTGAGGCCATGCTTAGAAACGACACCAACGACCAGTCCTTT
AACGACTATCTCTCCGCCGCCAACATGCTCTACCCTATACCCGCCAACGCTACCAACGTGCCCATAT
CCATCCCCTCCCGCAACTGGGCGGCTTTCCGCGGCTGGGCCTTCACGCGCCTTAAGACTAAGGAA
ACCCCATCACTGGGCTCGGGCTACGACCCTTATTACACCTACTCTGGCTCTATACCCTACCTAGATG
GAACCTTTTACCTCAACCACACCTTTAAGAAGGTGGCCATTACCTTTGACTCTTCTGTCAGCTGGCC
TGGCAATGACCGCCTGCTTACCCCCAACGAGTTTGAAATTAAGCGCTCAGTTGACGGGGAGGGTTA
CAACGTTGCCCAGTGTAACATGACCAAAGACTGGTTCCTGGTACAAATGCTAGCTAACTATAACATTG
GCTACCAGGGCTTCTATATCCCAGAGAGCTACAAGGACCGCATGTACTCCTTCTTTAGAAACTTCCA
GCCCATGAGCCGTCAGGTGGTGGATGATACTAAATACAAGGACTACCAACAGGTGGGCATCCTACA
CCAACACAACAACTCTGGATTTGTTGGCTACCTTGCCCCCACCATGCGCGAAGGACAGGCCTACCC
TGCTAACTTCCCCTATCCGCTTATAGGCAAGACCGCAGTTGACAGCATTACCCAGAAAAAGTTTCTTT
GCGATCGCACCCTTTGGCGCATCCCATTCTCCAGTAACTTTATGTCCATGGGCGCACTCACAGACCT
GGGCCAAAACCTTCTCTACGCCAACTCCGCCCACGCGCTAGACATGACTTTTGAGGTGGATCCCAT
GGACGAGCCCACCCTTCTTTATGTTTTGTTTGAAGTCTTTGACGTGGTCCGTGTGCACCAGCCGCA
CCGCGGCGTCATCGAAACCGTGTACCTGCGCACGCCCTTCTCGGCCGGCAACGCCACAACATAAA
GAAGCAAGCAACATCAACAACAGCTGCCGCCATGGGCTCCAGTGAGCAGGAACTGAAAGCCATTG
TCAAAGATCTTGGTTGTGGGCCATATTTTTTGGGCACCTATGACAAGCGCTTTCCAGGCTTTGTTTCT
CCACACAAGCTCGCCTGCGCCATAGTCAATACGGCCGGTCGCGAGACTGGGGGCGTACACTGGAT
GGCCTTTGCCTGGAACCCGCACTCAAAAACATGCTACCTCTTTGAGCCCTTTGGCTTTTCTGACCAG
CGACTCAAGCAGGTTTACCAGTTTGAGTACGAGTCACTCCTGCGCCGTAGCGCCATTGCTTCTTCC
CCCGACCGCTGTATAACGCTGGAAAAGTCCACCCAAAGCGTACAGGGGCCCAACTCGGCCGCCTG
TGGACTATTCTGCTGCATGTTTCTCCACGCCTTTGCCAACTGGCCCCAAACTCCCATGGATCACAAC
CCCACCATGAACCTTATTACCGGGGTACCCAACTCCATGCTCAACAGTCCCCAGGTACAGCCCACC
CTGCGTCGCAACCAGGAACAGCTCTACAGCTTCCTGGAGCGCCACTCGCCCTACTTCCGCAGCCA
CAGTGCGCAGATTAGGAGCGCCACTTCTTTTTGTCACTTGAAAAACATGTAAAAATAATGTACTAGAG
ACACTTTCAATAAAGGCAAATGCTTTTATTTGTACACTCTCGGGTGATTATTTACCCCCACCCTTGCC
GTCTGCGCCGTTTAAAAATCAAAGGGGTTCTGCCGCGCATCGCTATGCGCCACTGGCAGGGACAC
GTTGCGATACTGGTGTTTAGTGCTCCACTTAAACTCAGGCACAACCATCCGCGGCAGCTCGGTGAA
GTTTTCACTCCACAGGCTGCGCACCATCACCAACGCGTTTAGCAGGTCGGGCGCCGATATCTTGAA
GTCGCAGTTGGGGCCTCCGCCCTGCGCGCGCGAGTTGCGATACACAGGGTTGCAGCACTGGAAC
ACTATCAGCGCCGGGTGGTGCACGCTGGCCAGCACGCTCTTGTCGGAGATCAGATCCGCGTCCAG

GTCCTCCGCGTTGCTCAGGGCGAACGGAGTCAACTTTGGTAGCTGCCTTCCCAAAAAGGGCGCGT
GCCCAGGCTTTGAGTTGCACTCGCACCGTAGTGGCATCAAAAGGTGACCGTGCCCGGTCTGGGCG
TTAGGATACAGCGCCTGCATAAAAGCCTTGATCTGCTTAAAAGCCACCTGAGCCTTTGCGCCTTCAG
AGAAGAACATGCCGCAAGACTTGCCGGAAAACTGATTGGCCGGACAGGCCGCGTCGTGCACGCAG
CACCTTGCGTCGGTGTTGGAGATCTGCACCACATTTCGGCCCCACCGGTTCTTCACGATCTTGGCC
TTGCTAGACTGCTCCTTCAGCGCGCGCTGCCCGTTTTCGCTCGTCACATCCATTTCAATCACGTGCT
CCTTATTTATCATAATGCTTCCGTGTAGACACTTAAGCTCGCCTTCGATCTCAGCGCAGCGGTGCAG
CCACAACGCGCAGCCCGTGGGCTCGTGATGCTTGTAGGTCACCTCTGCAAACGACTGCAGGTACG
CCTGCAGGAATCGCCCCATCATCGTCACAAAGGTCTTGTTGCTGGTGAAGGTCAGCTGCAACCCGC
GGTGCTCCTCGTTCAGCCAGGTCTTGCATACGGCCGCCAGAGCTTCCACTTGGTCAGGCAGTAGTT
TGAAGTTCGCCTTTAGATCGTTATCCACGTGGTACTTGTCCATCAGCGCGCGCGCAGCCTCCATGC
CCTTCTCCCACGCAGACACGATCGGCACACTCAGCGGGTTCATCACCGTAATTTCACTTTCCGCTTC
GCTGGGCTCTTCCTCTTCCTCTTGCGTCCGCATACCACGCGCCACTGGGTCGTCTTCATTCAGCCG
CCGCACTGTGCGCTTACCTCCTTTGCCATGCTTGATTAGCACCGGTGGGTTGCTGAAACCCACCATT
TGTAGCGCCACATCTTCTCTTTCTTCCTCGCTGTCCACGATTACCTCTGGTGATGGCGGGCGCTCG
GGCTTGGGAGAAGGGCGCTTCTTTTTCTTCTTGGGCGCAATGGCCAAATCCGCCGCCGAGGTCGA
TGGCCGCGGGCTGGGTGTGCGCGGCACCAGCGCGTCTTGTGATGAGTCTTCCTCGTCCTCGGAC
TCGATACGCCGCCTCATCCGCTTTTTTGGGGGCGCCCGGGGAGGCGGCGGCGACGGGGACGGG
GACGACACGTCCTCCATGGTTGGGGGACGTCGCGCCGCACCGCGTCCGCGCTCGGGGGTGGTTT
CGCGCTGCTCCTCTTCCCGACTGGCCATTTCCTTCTCCTATAGGCAGAAAAAGATCATGGAGTCAGT
CGAGAAGAAGGACAGCCTAACCGCCCCCTCTGAGTTCGCCACCACCGCCTCCACCGATGCCGCCA
ACGCGCCTACCACCTTCCCCGTCGAGGCACCCCCGCTTGAGGAGGAGGAAGTGATTATCGAGCAG
GACCCAGGTTTTGTAAGCGAAGACGACGAGGACCGCTCAGTACCAACAGAGGATAAAAAGCAAGAC
CAGGACAACGCAGAGGCAAACGAGGAACAAGTCGGGCGGGGGGACGAAAGGCATGGCGACTACC
TAGATGTGGGAGACGACGTGCTGTTGAAGCATCTGCAGCGCCAGTGCGCCATTATCTGCGACGCGT
TGCAAGAGCGCAGCGATGTGCCCCTCGCCATAGCGGATGTCAGCCTTGCCTACGAACGCCACCTAT
TCTCACCGCGCGTACCCCCCAAACGCCAAGAAAACGGCACATGCGAGCCCAACCCGCGCCTCAAC
TTCTACCCCGTATTTGCCGTGCCAGAGGTGCTTGCCACCTATCACATCTTTTTCCAAAACTGCAAGAT
ACCCCTATCCTGCCGTGCCAACCGCAGCCGAGCGGACAAGCAGCTGGCCTTGCGGCAGGGCGCT
GTCATACCTGATATCGCCTCGCTCAACGAAGTGCCAAAAATCTTTGAGGGTCTTGGACGCGACGAG
AAGCGCGCGGCAAACGCTCTGCAACAGGAAAACAGCGAAAATGAAAGTCACTCTGGAGTGTTGGT
GGAACTCGAGGGTGACAACGCGCGCCTAGCCGTACTAAAACGCAGCATCGAGGTCACCCACTTTG
CCTACCCGGCACTTAACCTACCCCCCAAGGTCATGAGCACAGTCATGAGTGAGCTGATCGTGCGCC
GTGCGCAGCCCCTGGAGAGGGATGCAAATTTGCAAGAACAAACAGAGGAGGGCCTACCCGCAGTT
GGCGACGAGCAGCTAGCGCGCTGGCTTCAAACGCGCGAGCCTGCCGACTTGGAGGAGCGACGCA
AACTAATGATGGCCGCAGTGCTCGTTACCGTGGAGCTTGAGTGCATGCAGCGGTTCTTTGCTGACC
CGGAGATGCAGCGCAAGCTAGAGGAAACATTGCACTACACCTTTCGACAGGGCTACGTACGCCAGG
CCTGCAAGATCTCCAACGTGGAGCTCTGCAACCTGGTCTCCTACCTTGGAATTTTGCACGAAAACC
GCCTTGGGCAAAACGTGCTTCATTCCACGCTCAAGGGCGAGGCGCGCCGCGACTACGTCCGCGAC
TGCGTTTACTTATTTCTATGCTACACCTGGCAGACGGCCATGGGCGTTTGGCAGCAGTGCTTGGAG
GAGTGCAACCTCAAGGAGCTGCAGAAACTGCTAAAGCAAAACTTGAAGGACCTATGGACGGCCTTC
AACGAGCGCTCCGTGGCCGCGCACCTGGCGGACATCATTTTCCCCGAACGCCTGCTTAAAACCCT
GCAACAGGGTCTGCCAGACTTCACCAGTCAAAGCATGTTGCAGAACTTTAGGAACTTTATCCTAGAG
CGCTCAGGAATCTTGCCCGCCACCTGCTGTGCACTTCCTAGCGACTTTGTGCCCATTAAGTACCGC

GAATGCCCTCCGCCGCTTTGGGGCCACTGCTACCTTCTGCAGCTAGCCAACTACCTTGCCTACCAC
TCTGACATAATGGAAGACGTGAGCGGTGACGGTCTACTGGAGTGTCACTGTCGCTGCAACCTATGC
ACCCCGCACCGCTCCCTGGTTTGCAATTCGCAGCTGCTTAACGAAAGTCAAATTATCGGTACCTTTG
AGCTGCAGGGTCCCTCGCCTGACGAAAAGTCCGCGGCTCCGGGGTTGAAACTCACTCCGGGGCT
GTGGACGTCGGCTTACCTTCGCAAATTTGTACCTGAGGACTACCACGCCCACGAGATTAGGTTCTAC
GAAGACCAATCCCGCCCGCCTAATGCGGAGCTTACCGCCTGCGTCATTACCCAGGGCCACATTCTT
GGCCAATTGCAAGCCATCAACAAAGCCCGCCAAGAGTTTCTGCTACGAAAGGGACGGGGGGTTTA
CTTGGACCCCCAGTCCGGCGAGGAGCTCAACCCAATCCCCCCGCCGCCGCAGCCCTATCAGCAGC
AGCCGCGGGCCCTTGCTTCCCAGGATGGCACCCAAAAAGAAGCTGCAGCTGCCGCCGCCACCCA
CGGACGAGGAGGAATACTGGGACAGTCAGGCAGAGGAGGTTTTGGACGAGGAGGAGGAGGACAT
GATGGAAGACTGGGAGAGCCTAGACGAGGAAGCTTCCGAGGTCGAAGAGGTGTCAGACGAAACAC
CGTCACCCTCGGTCGCATTCCCCTCGCCGGCGCCCCAGAAATCGGCAACCGGTTCCAGCATGGCT
ACAACCTCCGCTCCTCAGGCGCCGCCGGCACTGCCCGTTCGCCGACCCAACCGTAGATGGGACAC
CACTGGAACCAGGGCCGGTAAGTCCAAGCAGCCGCCGCCGTTAGCCCAAGAGCAACAACAGCGC
CAAGGCTACCGCTCATGGCGCGGGCACAAGAACGCCATAGTTGCTTGCTTGCAAGACTGTGGGGG
CAACATCTCCTTCGCCCGCCGCTTTCTTCTCTACCATCACGGCGTGGCCTTCCCCCGTAACATCCTG
CATTACTACCGTCATCTCTACAGCCCATACTGCACCGGCGGCAGCGGCAGCAACAGCAGCGGCCAC
ACAGAAGCAAAGGCGACCGGATAGCAAGACTCTGACAAAGCCCAAGAAATCCACAGCGGCGGCAG
CAGCAGGAGGAGGAGCGCTGCGTCTGGCGCCCAACGAACCCGTATCGACCCGCGAGCTTAGAAA
CAGGATTTTTCCCACTCTGTATGCTATATTTCAACAGAGCAGGGGCCAAGAACAAGAGCTGAAAATAA
AAAACAGGTCTCTGCGATCCCTCACCCGCAGCTGCCTGTATCACAAAAGCGAAGATCAGCTTCGGC
GCACGCTGGAAGACGCGGAGGCTCTCTTCAGTAAATACTGCGCGCTGACTCTTAAGGACTAGTTTC
GCGCCCTTTCTCAAATTTAAGCGCGAAAACTACGTCATCTCCAGCGGCCACACCCGGCGCCAGCAC
CTGTTGTCAGCGCCATTATGAGCAAGGAAATTCCCACGCCCTACATGTGGAGTTACCAGCCACAAAT
GGGACTTGCGGCTGGAGCTGCCCAAGACTACTCAACCCGAATAAACTACATGAGCGCGGGACCCC
ACATGATATCCCGGGTCAACGGAATACGCGCCCACCGAAACCGAATTCTCCTGGAACAGGCGGCTA
TTACCACCACACCTCGTAATAACCTTAATCCCCGTAGTTGGCCCGCTGCCCTGGTGTACCAGGAAAG
TCCCGCTCCCACCACTGTGGTACTTCCCAGAGACGCCCAGGCCGAAGTTCAGATGACTAACTCAGG
GGCGCAGCTTGCGGGCGGCTTTCGTCACAGGGTGCGGTCGCCCGGGCAGGGTATAACTCACCTG
ACAATCAGAGGGCGAGGTATTCAGCTCAACGACGAGTCGGTGAGCTCCTCGCTTGGTCTCCGTCC
GGACGGGACATTTCAGATCGGCGGCGCCGGCCGCTCTTCATTCACGCCTCGTCAGGCAATCCTAA
CTCTGCAGACCTCGTCCTCTGAGCCGCGCTCTGGAGGCATTGGAACTCTGCAATTTATTGAGGAGT
TTGTGCCATCGGTCTACTTTAACCCCTTCTCGGGACCTCCCGGCCACTATCCGGATCAATTTATTCCT
AACTTTGACGCGGTAAAGGACTCGGCGGACGGCTACGACTGAATGTTAAGTGGAGAGGCAGAGCA
ACTGCGCCTGAAACACCTGGTCCACTGTCGCCGCCACAAGTGCTTTGCCCGCGACTCCGGTGAGT
TTTGCTACTTTGAATTGCCCGAGGATCATATCGAGGGCCCGGCGCACGGCGTCCGGCTTACCGCCC
AGGGAGAGCTTGCCCGTAGCCTGATTCGGGAGTTTACCCAGCGCCCCCTGCTAGTTGAGCGGGAC
AGGGGACCCTGTGTTCTCACTGTGATTTGCAACTGTCCTAACCCTGGATTACATCAAGATCCTCTAG
TTAATGGATCCAATCGATATCGCGAAGTCGATCGACGGATCGGGAGATCTCCCGATCCCCTATGGTG
CACTCTCAGTACAATCTGCTCTGATGCCGCATAGTTAAGCCAGTATCTGCTCCCTGCTTGTGTGTTG
GAGGTCGCTGAGTAGTGCGCGAGCAAAATTTAAGCTACAACAAGGCAAGGCTTGACCGACAATTGC
ATGAAGAATCTGCTTAGGGTTAGGCGTTTTGCGCTGCTTCGCGATGTACGGGCCAGATATACGCGTT
GACATTGATTATTGACTAGTTATTAATAGTAATCAATTACGGGGTCATTAGTTCATAGCCCATATATGGA
GTTCCGCGTTACATAACTTACGGTAAATGGCCCGCCTGGCTGACCGCCCAACGACCCCCGCCCATT

GACGTCAATAATGACGTATGTTCCCATAGTAACGCCAATAGGGACTTTCCATTGACGTCAATGGGTGG
AGTATTTACGGTAAACTGCCCACTTGGCAGTACATCAAGTGTATCATATGCCAAGTACGCCCCCTATT
GACGTCAATGACGGTAAATGGCCCGCCTGGCATTATGCCCAGTACATGACCTTATGGGACTTTCCTA
CTTGGCAGTACATCTACGTATTAGTCATCGCTATTACCATGGTGATGCGGTTTTGGCAGTACATCAAT
GGGCGTGGATAGCGGTTTGACTCACGGGGATTTCCAAGTCTCCACCCCATTGACGTCAATGGGAGT
TTGTTTTGGCACCAAAATCAACGGGACTTTCCAAAATGTCGTAACAACTCCGCCCCATTGACGCAAA
TGGGCGGTAGGCGTGTACGGTGGGAGGTCTATATAAGCAGAGCTCTCTGGCTAACTAGAGAACCCA
CTGCTTACTGGCTTATCGAAATTAATACGACTCACTATAGGGAGACCCAAGCTGGCTAGCGCGATCG
CCATGGATTTTTTTCGGGTAGTGGAAAACCAGCAGCCTCCCGCGACGATGCCCCTCAACGTTAGCT
TCACCAACAGGAACTATGACCTCGACTACGACTCGGTGCAGCCGTATTTCTACTGCGACGAGGAGG
AGAACTTCTACCAGCAGCAGCAGCAGAGCGAGCTGCAGCCCCCGGCGCCCAGCGAGGATATCTGG
AAGAAATTCGAGCTGCTGCCCACCCCGCCCCTGTCCCCTAGCCGCCGCTCCGGGCTCTGCTCGCC
CTCCTACGTTGCGGTCACACCCTTCTCCCTTCGGGGAGACAACGACGGCGGTGGCGGGAGCTTCT
CCACGGCCGACCAGCTGGAGATGGTGACCGAGCTGCTGGGAGGAGACATGGTGAACCAGAGTTT
CATCTGCGACCCGGACGACGAGACCTTCATCAAAAACATCATCATCCAGGACTGTATGTGGAGCGG
CTTCTCGGCCGCCGCCAAGCTCGTCTCAGAGAAGCTGGCCTCCTACCAGGCTGCGCGCAAAGACA
GCGGCAGCCCGAACCCCGCCCGCGGCCACAGCGTCTGCTCCACCTCCAGCTTGTACCTGCAGGA
TCTGAGCGCCGCCGCCTCAGAGTGCATCGACCCCTCGGTGGTCTTCCCCTACCCTCTCAACGACA
GCAGCTCGCCCAAGTCCTGCGCCTCGCAAGACTCCAGCGCCTTCTCTCCGTCCTCGGATTCTCTG
CTCTCCTCGACGGAGTCCTCCCCGCAGGGCAGCCCCGAGCCCCTGGTGCTCCATGAGGAGACAC
CGCCCACCACCAGCAGCGACTCTGAGGAGGAACAAGAAGATGAGGAAGAAATCGATGTTGTTTCTG
TGGAAAAGAGGCAGGCTCCTGGCAAAAGGTCAGAGTCTGGATCACCTTCTGCTGGAGGCCACAGC
AAACCTCCTCACAGCCCACTGGTCCTCAAGAGGTGCCACGTCTCCACACATCAGCACAACTACGCA
GCGCCTCCCTCCACTCGGAAGGACTATCCTGCTGCCAAGAGGGTCAAGTTGGACAGTGTCAGAGT
CCTGAGACAGATCAGCAACAACCGAAAATGCACCAGCCCCAGGTCCTCGGACACCGAGGAGAATG
TCAAGAGGCGAACACACAACGTCTTGGAGCGCCAGAGGAGGAACGAGCTAAAACGGAGCTTTTTT
GCCCTGCGTGACCAGATCCCGGAGTTGGAAAACAATGAAAAGGCCCCCAAGGTAGTTATCCTTAAA
AAAGCCACAGCATACATCCTGTCCGTCCAAGCAGAGGAGCAAAAGCTCATTTCTGAAGAGGACTTG
TTGCGGAAACGACGAGAACAGTTGAAACACAAACTTGAACAGCTACGGAACTCTTGTGCGCAATGT
ACTAACTACGCTTTGTTGAAACTCGCTGGCGATGTTGAAAGTAACCCCGGTCCTATGAGGCAGCCAC
CTGGCGAGTCTGACATGGCTGTCAGCGACGCGCTGCTCCCATCTTTCTCCACGTTCGCGTCTGGC
CCGGCGGGAAGGGAGAAGACACTGCGTCAAGCAGGTGCCCCGAATAACCGCTGGCGGGAGGAGC
TCTCCCACATGAAGCGACTTCCCCCAGTGCTTCCCGGCCGCCCCTATGACCTGGCGGCGGCGACC
GTGGCCACAGACCTGGAGAGCGGCGGAGCCGGTGCGGCTTGCGGCGGTAGCAACCTGGCGCCC
CTACCTCGGAGAGAGACCGAGGAGTTCAACGATCTCCTGGACCTGGACTTTATTCTCTCCAATTCGC
TGACCCATCCTCCGGAGTCAGTGGCCGCCACCGTGTCCTCGTCAGCGTCAGCCTCCTCTTCGTCG
TCGCCGTCGAGCAGCGGCCCTGCCAGCGCGCCCTCCACCTGCAGCTTCACCTATCCGATCCGGGC
CGGGAACGACCCGGGCGTGGCGCCGGGCGGCACGGGCGGAGGCCTCCTCTATGGCAGGGAGTC
CGCTCCCCCTCCGACGGCTCCCTTCAACCTGGCGGACATCAACGACGTGAGCCCCTCGGGCGGC
TTCGTGGCCGAGCTCCTGCGGCCAGAATTGGACCCGGTGTACATTCCGCCGCAGCAGCCGCAGCC
GCCAGGTGGCGGGCTGATGGGCAAGTTCGTGCTGAAGGCGTCGCTGAGCGCCCCTGGCAGCGAG
TACGGCAGCCCGTCGGTCATCAGCGTCAGCAAAGGCAGCCCTGACGGCAGCCACCCGGTGGTGG
TGGCGCCCTACAACGGCGGGCCGCCGCGCACGTGCCCCAAGATCAAGCAGGAGGCGGTCTCTTC
GTGCACCCACTTGGGCGCTGGACCCCCTCTCAGCAATGGCCACCGGCCGGCTGCACACGACTTC

CCCCTGGGGCGGCAGCTCCCCAGCAGGACTACCCCGACCCTGGGTCTTGAGGAAGTGCTGAGCA
GCAGGGACTGTCACCCTGCCCTGCCGCTTCCTCCCGGCTTCCATCCCCACCCGGGGCCCAATTAC
CCATCCTTCCTGCCCGATCAGATGCAGCCGCAAGTCCCGCCGCTCCATTACCAAGAGCTCATGCCA
CCCGGTTCCTGCATGCCAGAGGAGCCCAAGCCAAAGAGGGGAAGACGATCGTGGCCCCGGAAAA
GGACCGCCACCCACACTTGTGATTACGCGGGCTGCGGCAAAACCTACACAAAGAGTTCCCATCTCA
AGGCACACCTGCGAACCCACACAGGTGAGAAACCTTACCACTGTGACTGGGACGGCTGTGGATGG
AAATTCGCCCGCTCAGATGAACTGACCAGGCACTACCGTAAACACACGGGGCACCGCCCGTTCCA
GTGCCAAAAATGCGACCGAGCATTTTCCAGGTCGGACCACCTCGCCTTACACATGAAGAGGCATTTT
ACGCGTACGCGGCCGCTCGAGCAGAAACTCATCTCAGAAGAGGATCTGGCAGCAAATGATATCCTG
GATTACAAGGATGACGACGATAAGGTTTAAACTCGAGTCTAGAGGGCCCGTTTAAACCCGCTGATCA
GCCTCGACTGTGCCTTCTAGTTGCCAGCCATCTGTTGTTTGCCCCTCCCCCGTGCCTTCCTTGACC
CTGGAAGGTGCCACTCCCACTGTCCTTTCCTAATAAAATGAGGAAATTGCATCGCATTGTCTGAGTA
GGTGTCATTCTATTCTGGGGGGTGGGGTGGGGCAGGACAGCAAGGGGGAGGATTGGGAAGACAAT
AGCAGGCATGCTGGGGATGCGGTGGGCTCTATGGCTTCTGAGGCGGAAAGAACCAGCTGGGGCT
CTAGGGGGTATCCCCACGCGCCCTGTAGCGGCGCATTAAGCGCGGCGGGTGTGGTGGTTACGCGC
AGCGTGACCGCTACACTTGCCAGCGCCCTAGCGCCCGCTCCTTTCGCTTTCTTCCCTTCCTTTCTC
GCCACGTTCGCCGGCTTTCCCCGTCAAGCTCTAAATCGGGGGCTCCCTTTAGGGTTCCGATTTAGT
GCTTTACGGCACCTCGACCCCAAAAAACTTGATTAGGGTGATGGTTCACATCGAATTCCTGCAGCCC
GGGGGATCCATTAACTAATAAAAAAAAATAATAAAGCATCACTTACTTAAAATCAGTTAGCAAATTTCTG
TCCAGTTTATTCAGCAGCACCTCCTTGCCCTCCTCCCAGCTCTGGTATTGCAGCTTCCTCCTGGCTG
CAAACTTTCTCCACAATCTAAATGGAATGTCAGTTTCCTCCTGTTCCTGTCCATCCGCACCCACTATC
TTCATGTTGTTGCAGATGAAGCGCGCAAGACCGTCTGAAGATACCTTCAACCCCGTGTATCCATATG
ACACGGAAACCGGTCCTCCAACTGTGCCTTTTCTTACTCCTCCCTTTGTATCCCCCAATGGGTTTCA
AGAGAGTCCCCCTGGGGTACTCTCTTTGCGCCTATCCGAACCTCTAGTTACCTCCAATGGCATGCTT
GCGCTCAAAATGGGCAACGGCCTCTCTCTGGACGAGGCCGGCAACCTTACCTCCCAAAATGTAACC
ACTGTGAGCCCACCTCTCAAAAAAACCAAGTCAAACATAAACCTGGAAATATCTGCACCCCTCACAG
TTACCTCAGAAGCCCTAACTGTGGCTGCCGCCGCACCTCTAATGGTCGCGGGCAACACACTCACCA
TGCAATCACAGGCCCCGCTAACCGTGCACGACTCCAAACTTAGCATTGCCACCCAAGGACCCCTCA
CAGTGTCAGAAGGAAAGCTAGCCCTGCAAACATCAGGCCCCCTCACCACCACCGATAGCAGTACCC
TTACTATCACTGCCTCACCCCCTCTAACTACTGCCACTGGTAGCTTGGGCATTGACTTGAAAGAGCC
CATTTATACACAAAATGGAAAACTAGGACTAAAGTACGGGGCTCCTTTGCATGTAACAGACGACCTAA
ACACTTTGACCGTAGCAACTGGTCCAGGTGTGACTATTAATAATACTTCCTTGCAAACTAAAGTTACT
GGAGCCTTGGGTTTTGATTCACAAGGCAATATGCAACTTAATGTAGCAGGAGGACTAAGGATTGATT
CTCAAAACAGACGCCTTATACTTGATGTTAGTTATCCGTTTGATGCTCAAAACCAACTAAATCTAAGAC
TAGGACAGGGCCCTCTTTTTATAAACTCAGCCCACAACTTGGATATTAACTACAACAAAGGCCTTTAC
TTGTTTACAGCTTCAAACAATTCCAAAAAGCTTGAGGTTAACCTAAGCACTGCCAAGGGGTTGATGTT
TGACGCTACAGCCATAGCCATTAATGCAGGAGATGGGCTTGAATTTGGTTCACCTAATGCACCAAAC
ACAAATCCCCTCAAAACAAAAATTGGCCATGGCCTAGAATTTGATTCAAACAAGGCTATGGTTCCTAA
ACTAGGAACTGGCCTTAGTTTTGACAGCACAGGTGCCATTACAGTAGGAAACAAAAATAATGATAAGC
TAACTTTGTGGACCACACCAGCTCCATCTCCTAACTGTAGACTAAATGCAGAGAAAGATGCTAAACTC
ACTTTGGTCTTAACAAAATGTGGCAGTCAAATACTTGCTACAGTTTCAGTTTTGGCTGTTAAAGGCAG
TTTGGCTCCAATATCTGGAACAGTTCAAAGTGCTCATCTTATTATAAGATTTGACGAAAATGGAGTGCT
ACTAAACAATTCCTTCCTGGACCCAGAATATTGGAACTTTAGAAATGGAGATCTTACTGAAGGCACAG
CCTATACAAACGCTGTTGGATTTATGCCTAACCTATCAGCTTATCCAAAATCTCACGGTAAAACTGCCA

AAAGTAACATTGTCAGTCAAGTTTACTTAAACGGAGACAAAACTAAACCTGTAACACTAACCATTACA
CTAAACGGTACACAGGAAACAGGAGACACAACTCCAAGTGCATACTCTATGTCATTTTCATGGGACT
GGTCTGGCCACAACTACATTAATGAAATATTTGCCACATCCTCTTACACTTTTTCATACATTGCCCAAG
AATAAAGAATCGTTTGTGTTATGTTTCAACGTGTTTATTTTTCAATTGCAGAAAATTTCAAGTCATTTTT
CATTCAGTAGTATAGCCCCACCACCACATAGCTTATACAGATCACCGTACCTTAATCAAACTCACAGAA
CCCTAGTATTCAACCTGCCACCTCCCTCCCAACACACAGAGTACACAGTCCTTTCTCCCCGGCTGG
CCTTAAAAAGCATCATATCATGGGTAACAGACATATTCTTAGGTGTTATATTCCACACGGTTTCCTGTC
GAGCCAAACGCTCATCAGTGATATTAATAAACTCCCCGGGCAGCTCACTTAAGTTCATGTCGCTGTC
CAGCTGCTGAGCCACAGGCTGCTGTCCAACTTGCGGTTGCTTAACGGGCGGCGAAGGAGAAGTCC
ACGCCTACATGGGGGTAGAGTCATAATCGTGCATCAGGATAGGGCGGTGGTGCTGCAGCAGCGCG
CGAATAAACTGCTGCCGCCGCCGCTCCGTCCTGCAGGAATACAACATGGCAGTGGTCTCCTCAGCG
ATGATTCGCACCGCCCGCAGCATAAGGCGCCTTGTCCTCCGGGCACAGCAGCGCACCCTGATCTC
ACTTAAATCAGCACAGTAACTGCAGCACAGCACCACAATATTGTTCAAAATCCCACAGTGCAAGGCG
CTGTATCCAAAGCTCATGGCGGGGACCACAGAACCCACGTGGCCATCATACCACAAGCGCAGGTAG
ATTAAGTGGCGACCCCTCATAAACACGCTGGACATAAACATTACCTCTTTTGGCATGTTGTAATTCAC
CACCTCCCGGTACCATATAAACCTCTGATTAAACATGGCGCCATCCACCACCATCCTAAACCAGCTG
GCCAAAACCTGCCCGCCGGCTATACACTGCAGGGAACCGGGACTGGAACAATGACAGTGGAGAGC
CCAGGACTCGTAACCATGGATCATCATGCTCGTCATGATATCAATGTTGGCACAACACAGGCACACG
TGCATACACTTCCTCAGGATTACAAGCTCCTCCCGCGTTAGAACCATATCCCAGGGAACAACCCATT
CCTGAATCAGCGTAAATCCCACACTGCAGGGAAGACCTCGCACGTAACTCACGTTGTGCATTGTCA
AAGTGTTACATTCGGGCAGCAGCGGATGATCCTCCAGTATGGTAGCGCGGGTTTCTGTCTCAAAAG
GAGGTAGACGATCCCTACTGTACGGAGTGCGCCGAGACAACCGAGATCGTGTTGGTCGTAGTGTCA
TGCCAAATGGAACGCCGGACGTAGTCATATTTCCTGAAGCAAAACCAGGTGCGGGCGTGACAAACA
GATCTGCGTCTCCGGTCTCGCCGCTTAGATCGCTCTGTGTAGTAGTTGTAGTATATCCACTCTCTCAA
AGCATCCAGGCGCCCCCTGGCTTCGGGTTCTATGTAAACTCCTTCATGCGCCGCTGCCCTGATAAC
ATCCACCACCGCAGAATAAGCCACACCCAGCCAACCTACACATTCGTTCTGCGAGTCACACACGGG
AGGAGCGGGAAGAGCTGGAAGAACCATGTTTTTTTTTTATTCCAAAAGATTATCCAAAACCTCAAAAT
GAAGATCTATTAAGTGAACGCGCTCCCCTCCGGTGGCGTGGTCAAACTCTACAGCCAAAGAACAGA
TAATGGCATTTGTAAGATGTTGCACAATGGCTTCCAAAAGGCAAACGGCCCTCACGTCCAAGTGGAC
GTAAAGGCTAAACCCTTCAGGGTGAATCTCCTCTATAAACATTCCAGCACCTTCAACCATGCCCAAAT
AATTCTCATCTCGCCACCTTCTCAATATATCTCTAAGCAAATCCCGAATATTAAGTCCGGCCATTGTAA
AAATCTGCTCCAGAGCGCCCTCCACCTTCAGCCTCAAGCAGCGAATCATGATTGCAAAAATTCAGGT
TCCTCACAGACCTGTATAAGATTCAAAAGCGGAACATTAACAAAAATACCGCGATCCCGTAGGTCCCT
TCGCAGGGCCAGCTGAACATAATCGTGCAGGTCTGCACGGACCAGCGCGGCCACTTCCCCGCCAG
GAACCATGACAAAAGAACCCACACTGATTATGACACGCATACTCGGAGCTATGCTAACCAGCGTAGC
CCCGATGTAAGCTTGTTGCATGGGCGGCGATATAAAATGCAAGGTGCTGCTCAAAAAATCAGGCAAA
GCCTCGCGCAAAAAAGAAAGCACATCGTAGTCATGCTCATGCAGATAAAGGCAGGTAAGCTCCGGA
ACCACCACAGAAAAAGACACCATTTTTCTCTCAAACATGTCTGCGGGTTTCTGCATAAACACAAAATA
AAATAACAAAAAAACATTTAAACATTAGAAGCCTGTCTTACAACAGGAAAAACAACCCTTATAAGCATA
AGACGGACTACGGCCATGCCGGCGTGACCGTAAAAAAACTGGTCACCGTGATTAAAAAGCACCACC
GACAGCTCCTCGGTCATGTCCGGAGTCATAATGTAAGACTCGGTAAACACATCAGGTTGATTCACAT
CGGTCAGTGCTAAAAAGCGACCGAAATAGCCCGGGGGAATACATACCCGCAGGCGTAGAGACAACA
TTACAGCCCCCATAGGAGGTATAACAAAATTAATAGGAGAGAAAAACACATAAACACCTGAAAAACCC
TCCTGCCTAGGCAAAATAGCACCCTCCCGCTCCAGAACAACATACAGCGCTTCCACAGCGGCAGCC

ATAACAGTCAGCCTTACCAGTAAAAAAGAAAACCTATTAAAAAAACACCACTCGACACGGCACCAGC
TCAATCAGTCACAGTGTAAAAAAGGGCCAAGTGCAGAGCGAGTATATATAGGACTAAAAAATGACGTA
ACGGTTAAAGTCCACAAAAAACACCCAGAAAACCGCACGCGAACCTACGCCCAGAAACGAAAGCCA
AAAAACCCACAACTTCCTCAAATCGTCACTTCCGTTTTCCCACGTTACGTCACTTCCCATTTTAAGAA
AACTACAATTCCCAACACATACAAGTTACTCCGCCCTAAAACCTACGTCACCCGCCCCGTTCCCACG
CCCCGCGCCACGTCACAAACTCCACCCCCTCATTATCATATTGGCTTCAATCCAAAATAAGGTATATTA
TTGATGATGTTACATCGTTAATTAACGATTTCGAACCCGGGGTACCGAATTCCTCGAGTCTAGAGGAG
CATGCGACGTCGCAATTCGCCCTATAGTGAGTCGTATTACAATTCACTGGCCGTCGTTTTACAACGTC
GTGACTGGGAAAACCCTGGCGTTACCCAACTTAATCGCCTTGCAGCACATCCCCCTTTCGCCAGCT
GGCGTAATAGCGAAGAGGCCCGCACCGATCGCCCTTCCCAACAGTTGCGCAGCCTGAATGGCGAA
TGGAAATTGTAAGCGTTAATATTTTGTTAAAATTCGCGTTAAATTTTTGTTAAATCAGCTCATTTTTTAAC
CAATAGGCCGAAATCGGCAAAATCCCTTATAAATCAAAAGAATAGACCGAGATAGGGTTGAGTGTTGT
TCCAGTTTGGAACAAGAGTCCACTATTAAAGAACGTGGACTCCAACGTCAAAGGGCGAAAAACCGT
CTATCAGGGCGATGGCCCACTACGTGAACCATCACCCTAATCAAGTTTTTTGGGGTCGAGGTGCCGT
AAAGCACTAAATCGGAACCCTAAAGGGAGCCCCCGATTTAGAGCTTGACGGGGAAAGCCGGCGAA
CGTGGCGAGAAAGGAAGGGAAGAAAGCGAAAGGAGCGGGCGCTAGGGCGCTGGCAAGTGTAGCG
GTCACGCTGCGCGTAACCACCACACCCGCCGCGCTTAATGCGCCGCTACAGGGCGCGTCCTGATG
CGGTATTTTCTCCTTACGCATCTGTGCGGTATTTCACACCGCATACAGGTGGCACTTTTCGGGGAAA
TGTGCGCGGAACCCCTATTTGTTTATTTTTCTAAATACATTCAAATATGTATCCGCTCATGAGACAATAA
CCCTGATAAATGCTTCAATAATATTGAAAAAGGAAGAGTATGAGTATTCAACATTTCCGTGTCGCCCTT
ATTCCCTTTTTTGCGGCATTTTGCCTTCCTGTTTTTGCTCACCCAGAAACGCTGGTGAAAGTAAAAGA
TGCTGAAGATCAGTTGGGTGCACGAGTGGGTTACATCGAACTGGATCTCAACAGCGGTAAGATCCT
TGAGAGTTTTCGCCCCGAAGAACGTTTTCCAATGATGAGCACTTTTAAAGTTCTGCTATGTGGCGCG
GTATTATCCCGTATTGACGCCGGGCAAGAGCAACTCGGTCGCCGCATACACTATTCTCAGAATGACT
TGGTTGAGTACTCACCAGTCACAGAAAAGCATCTTACGGATGGCATGACAGTAAGAGAATTATGCAG
TGCTGCCATAACCATGAGTGATAACACTGCGGCCAACTTACTTCTGACAACGATCGGAGGACCGAA
GGAGCTAACCGCTTTTTTGCACAACATGGGGGATCATGTAACTCGCCTTGATCGTTGGGAACCGGA
GCTGAATGAAGCCATACCAAACGACGAGCGTGACACCACGATGCCTGTAGCAATGGCAACAACGTT
GCGCAAACTATTAACTGGCGAACTACTTACTCTAGCTTCCCGGCAACAATTAATAGACTGGATGGAG
GCGGATAAAGTTGCAGGACCACTTCTGCGCTCGGCCCTTCCGGCTGGCTGGTTTATTGCTGATAAA
TCTGGAGCCGGTGAGCGTGGGTCTCGCGGTATCATTGCAGCACTGGGGCCAGATGGTAAGCCCTC
CCGTATCGTAGTTATCTACACGACGGGGAGTCAGGCAACTATGGATGAACGAAATAGACAGATCGCT
GAGATAGGTGCCTCACTGATTAAGCATTGGTAACTGTCAGACCAAGTTTACTCATATATACTTTAGATT
GATTTAAAACTTCATTTTTAATTTAAAAGGATCTAGGTGAAGATCCTTTTTGATAATCTCATGACCAAAA
TCCCTTAACGTGAGTTTTCGTTCCACTGAGCGTCAGACCCCGTAGAAAAGATCAAAGGATCTTCTTG
AGATCCTTTTTTTCTGCGCGTAATCTGCTGCTTGCAAACAAAAAAACCACCGCTACCAGCGGTGGTT
TGTTTGCCGGATCAAGAGCTACCAACTCTTTTTCCGAAGGTAACTGGCTTCAGCAGAGCGCAGATAC
CAAATACTGTTCTTCTAGTGTAGCCGTAGTTAGGCCACCACTTCAAGAACTCTGTAGCACCGCCTACA
TACCTCGCTCTGCTAATCCTGTTACCAGTGGCTGCTGCCAGTGGCGATAAGTCGTGTCTTACCGGGT
TGGACTCAAGACGATAGTTACCGGATAAGGCGCAGCGGTCGGGCTGAACGGGGGGTTCGTGCACA
CAGCCCAGCTTGGAGCGAACGACCTACACCGAACTGAGATACCTACAGCGTGAGCTATGAGAAAGC
GCCACGCTTCCCGAAGGGAGAAAGGCGGACAGGTATCCGGTAAGCGGCAGGGTCGGAACAGGAG
AGCGCACGAGGGAGCTTCCAGGGGGAAACGCCTGGTATCTTTATAGTCCTGTCGGGTTTCGCCACC
TCTGACTTGAGCGTCGATTTTTGTGATGCTCGTCAGGGGGGCGGAGCCTATGGAAAAACGCCAGCA

ACGCGGCCTTTTTACGGTTCCTGGCCTTTTGCTGGCCTTTTGCTCACATGTTCTTTCCTGCGTTATC
CCCTGATTCTGTGGATAACCGTATTACCGCCT

P20:

>P143 pAdHM4-SFFV-Oct4Cherry

CGCTGGTGAAAGTAAAAGATGCTGAAGATCAGTTGGGTGCACGAGTGGGTTACATCGAACTGGATC
TCAACAGCGGTAAGATCCTTGAGAGTTTTCGCCCCGAAGAACGTTTTCCAATGATGAGCACTTTTAA
AGTTCTGCTATGTGGCGCGGTATTATCCCGTATTGACGCCGGGCAAGAGCAACTCGGTCGCCGCAT
ACACTATTCTCAGAATGACTTGGTTGAGTACTCACCAGTCACAGAAAAGCATCTTACGGATGGCATG
ACAGTAAGAGAATTATGCAGTGCTGCCATAACCATGAGTGATAACACTGCGGCCAACTTACTTCTGA
CAACGATCGGAGGACCGAAGGAGCTAACCGCTTTTTTGCACAACATGGGGGATCATGTAACTCGCC
TTGATCGTTGGGAACCGGAGCTGAATGAAGCCATACCAAACGACGAGCGTGACACCACGATGCCTG
TAGCAATGGCAACAACGTTGCGCAAACTATTAACTGGCGAACTACTTACTCTAGCTTCCCGGCAACA
ATTAATAGACTGGATGGAGGCGGATAAAGTTGCAGGACCACTTCTGCGCTCGGCCCTTCCGGCTGG
CTGGTTTATTGCTGATAAATCTGGAGCCGGTGAGCGTGGGTCTCGCGGTATCATTGCAGCACTGGG
GCCAGATGGTAAGCCCTCCCGTATCGTAGTTATCTACACGACGGGGAGTCAGGCAACTATGGATGAA
CGAAATAGACAGATCGCTGAGATAGGTGCCTCACTGATTAAGCATTGGTAACTGTCAGACCAAGTTT
ACTCATATATACTTTAGATTGATTTAAAACTTCATTTTTAATTTAAAAGGATCTAGGTGAAGATCCTTTTT
GATAATCTCATGACCAAAATCCCTTAACGTGAGTTTTCGTTCCACTGAGCGTCAGACCCCGTAGAAA
AGATCAAAGGATCTTCTTGAGATCCTTTTTTTCTGCGCGTAATCTGCTGCTTGCAAACAAAAAAACCA
CCGCTACCAGCGGTGGTTTGTTTGCCGGATCAAGAGCTACCAACTCTTTTTCCGAAGGTAACTGGC
TTCAGCAGAGCGCAGATACCAAATACTGTTCTTCTAGTGTAGCCGTAGTTAGGCCACCACTTCAAGA
ACTCTGTAGCACCGCCTACATACCTCGCTCTGCTAATCCTGTTACCAGTGGCTGCTGCCAGTGGCGA
TAAGTCGTGTCTTACCGGGTTGGACTCAAGACGATAGTTACCGGATAAGGCGCAGCGGTCGGGCTG
AACGGGGGGTTCGTGCACACAGCCCAGCTTGGAGCGAACGACCTACACCGAACTGAGATACCTAC
AGCGTGAGCTATGAGAAAGCGCCACGCTTCCCGAAGGGAGAAAGGCGGACAGGTATCCGGTAAGC
GGCAGGGTCGGAACAGGAGAGCGCACGAGGGAGCTTCCAGGGGGAAACGCCTGGTATCTTTATAG
TCCTGTCGGGTTTCGCCACCTCTGACTTGAGCGTCGATTTTTGTGATGCTCGTCAGGGGGGCGGA
GCCTATGGAAAAACGCCAGCAACGCGGCCTTTTTACGGTTCCTGGCCTTTTGCTGGCCTTTTGCTC
ACATGTTCTTTCCTGCGTTATCCCCTGATTCTGTGGATAACCGTATTACCGCCTTTGAGTGAGCTGAT
ACCGCTCGCCGCAGCCGAACGACCGAGCGCAGCGAGTCAGTGAGCGAGGAAGCGGAAGAGCGC
CCAATACGCAAACCGCCTCTCCCCGCGCGTTGGCCGATTCATTAATGCAGCTGGCACGACAGGTTT
CCCGACTGGAAAGCGGGCAGTGAGCGCAACGCAATTAATGTGAGTTAGCTCACTCATTAGGCACCC
CAGGCTTTACACTTTATGCTTCCGGCTCGTATGTTGTGTGGAATTGTGAGCGGATAACAATTTCACAC
AGGAAACAGCTATGACCATGATTACGCCAAGCTATTTAGGTGACACTATAGAATACTCAAGCTAGTTAA
TTAACGTTAATTAACATCATCAATAATATACCTTATTTTGGATTGAAGCCAATATGATAATGAGGGGGTG
GAGTTTGTGACGTGGCGCGGGGCGTGGGAACGGGGCGGGTGACGTAGTAGTGTGGCGGAAGTGT
GATGTTGCAAGTGTGGCGGAACACATGTAAGCGACGGATGTGGCAAAAGTGACGTTTTTGGTGTGC
GCCGGTGTACACAGGAAGTGACAATTTTCGCGCGGTTTTAGGCGGATGTTGTAGTAAATTTGGGCGT
AACCGAGTAAGATTTGGCCATTTTCGCGGGAAAACTGAATAAGAGGAAGTGAAATCTGAATAATTTTG
TGTTACTCATAGCGCGTAATCTCTAGCATCGTAACTATAACGGTCCTAAGGTAGCGAGAATTATTTAAA
TATCTATGTCGGGTGCGGAGAAAGAGGTAATGAAATGGCATCGACTCGAAGATCTGGGCGTGGTTAA
GGGTGGGAAAGAATATATAAGGTGGGGGTCTTATGTAGTTTTGTATCTGTTTTGCAGCAGCCGCCGC
CGCCATGAGCACCAACTCGTTTGATGGAAGCATTGTGAGCTCATATTTGACAACGCGCATGCCCCCA
TGGGCCGGGGTGCGTCAGAATGTGATGGGCTCCAGCATTGATGGTCGCCCCGTCCTGCCCGCAAA
CTCTACTACCTTGACCTACGAGACCGTGTCTGGAACGCCGTTGGAGACTGCAGCCTCCGCCGCCG
CTTCAGCCGCTGCAGCCACCGCCCGCGGGATTGTGACTGACTTTGCTTTCCTGAGCCCGCTTGCA

AGCAGTGCAGCTTCCCGTTCATCCGCCCGCGATGACAAGTTGACGGCTCTTTTGGCACAATTGGAT
TCTTTGACCCGGGAACTTAATGTCGTTTCTCAGCAGCTGTTGGATCTGCGCCAGCAGGTTTCTGCC
CTGAAGGCTTCCTCCCCTCCCAATGCGGTTTAAAACATAAATAAAAAACCAGACTCTGTTTGGATTTG
GATCAAGCAAGTGTCTTGCTGTCTTTATTTAGGGGTTTTGCGCGCGCGGTAGGCCCGGGACCAGCG
GTCTCGGTCGTTGAGGGTCCTGTGTATTTTTTCCAGGACGTGGTAAAGGTGACTCTGGATGTTCAGA
TACATGGGCATAAGCCCGTCTCTGGGGTGGAGGTAGCACCACTGCAGAGCTTCATGCTGCGGGGT
GGTGTTGTAGATGATCCAGTCGTAGCAGGAGCGCTGGGCGTGGTGCCTAAAAATGTCTTTCAGTAG
CAAGCTGATTGCCAGGGGCAGGCCCTTGGTGTAAGTGTTTACAAAGCGGTTAAGCTGGGATGGGTG
CATACGTGGGGATATGAGATGCATCTTGGACTGTATTTTTAGGTTGGCTATGTTCCCAGCCATATCCC
TCCGGGGATTCATGTTGTGCAGAACCACCAGCACAGTGTATCCGGTGCACTTGGGAAATTTGTCAT
GTAGCTTAGAAGGAAATGCGTGGAAGAACTTGGAGACGCCCTTGTGACCTCCAAGATTTTCCATGCA
TTCGTCCATAATGATGGCAATGGGCCCACGGGCGGCGGCCTGGGCGAAGATATTTCTGGGATCACT
AACGTCATAGTTGTGTTCCAGGATGAGATCGTCATAGGCCATTTTTACAAAGCGCGGGCGGAGGGT
GCCAGACTGCGGTATAATGGTTCCATCCGGCCCAGGGGCGTAGTTACCCTCACAGATTTGCATTTCC
CACGCTTTGAGTTCAGATGGGGGGATCATGTCTACCTGCGGGGCGATGAAGAAAACGGTTTCCGG
GGTAGGGGAGATCAGCTGGGAAGAAAGCAGGTTCCTGAGCAGCTGCGACTTACCGCAGCCGGTG
GGCCCGTAAATCACACCTATTACCGGCTGCAACTGGTAGTTAAGAGAGCTGCAGCTGCCGTCATCC
CTGAGCAGGGGGGCCACTTCGTTAAGCATGTCCCTGACTCGCATGTTTTCCCTGACCAAATCCGCC
AGAAGGCGCTCGCCGCCCAGCGATAGCAGTTCTTGCAAGGAAGCAAAGTTTTTCAACGGTTTGAGA
CCGTCCGCCGTAGGCATGCTTTTGAGCGTTTGACCAAGCAGTTCCAGGCGGTCCCACAGCTCGGT
CACCTGCTCTACGGCATCTCGATCCAGCATATCTCCTCGTTTCGCGGGTTGGGGCGGCTTTCGCTG
TACGGCAGTAGTCGGTGCTCGTCCAGACGGGCCAGGGTCATGTCTTTCCACGGGCGCAGGGTCCT
CGTCAGCGTAGTCTGGGTCACGGTGAAGGGGTGCGCTCCGGGCTGCGCGCTGGCCAGGGTGCG
CTTGAGGCTGGTCCTGCTGGTGCTGAAGCGCTGCCGGTCTTCGCCCTGCGCGTCGGCCAGGTAG
CATTTGACCATGGTGTCATAGTCCAGCCCCTCCGCGGCGTGGCCCTTGGCGCGCAGCTTGCCCTT
GGAGGAGGCGCCGCACGAGGGGCAGTGCAGACTTTTGAGGGCGTAGAGCTTGGGCGCGAGAAAT
ACCGATTCCGGGGAGTAGGCATCCGCGCCGCAGGCCCCGCAGACGGTCTCGCATTCCACGAGCC
AGGTGAGCTCTGGCCGTTCGGGGTCAAAAACCAGGTTTCCCCCATGCTTTTTGATGCGTTTCTTAC
CTCTGGTTTCCATGAGCCGGTGTCCACGCTCGGTGACGAAAAGGCTGTCCGTGTCCCCGTATACAG
ACTTGAGAGGCCTGTCCTCGAGCGGTGTTCCGCGGTCCTCCTCGTATAGAAACTCGGACCACTCTG
AGACAAAGGCTCGCGTCCAGGCCAGCACGAAGGAGGCTAAGTGGGAGGGGTAGCGGTCGTTGTC
CACTAGGGGGTCCACTCGCTCCAGGGTGTGAAGACACATGTCGCCCTCTTCGGCATCAAGGAAGG
TGATTGGTTTGTAGGTGTAGGCCACGTGACCGGGTGTTCCTGAAGGGGGGCTATAAAAGGGGGTG
GGGGCGCGTTCGTCCTCACTCTCTTCCGCATCGCTGTCTGCGAGGGCCAGCTGTTGGGGTGAGTA
CTCCCTCTGAAAAGCGGGCATGACTTCTGCGCTAAGATTGTCAGTTTCCAAAAACGAGGAGGATTT
GATATTCACCTGGCCCGCGGTGATGCCTTTGAGGGTGGCCGCATCCATCTGGTCAGAAAAGACAAT
CTTTTTGTTGTCAAGCTTGGTGGCAAACGACCCGTAGAGGGCGTTGGACAGCAACTTGGCGATGGA
GCGCAGGGTTTGGTTTTTGTCGCGATCGGCGCGCTCCTTGGCCGCGATGTTTAGCTGCACGTATTC
GCGCGCAACGCACCGCCATTCGGGAAAGACGGTGGTGCGCTCGTCGGGCACCAGGTGCACGCGC
CAACCGCGGTTGTGCAGGGTGACAAGGTCAACGCTGGTGGCTACCTCTCCGCGTAGGCGCTCGTT
GGTCCAGCAGAGGCGGCCGCCCTTGCGCGAGCAGAATGGCGGTAGGGGGTCTAGCTGCGTCTCG
TCCGGGGGGTCTGCGTCCACGGTAAAGACCCCGGGCAGCAGGCGCGCGTCGAAGTAGTCTATCTT
GCATCCTTGCAAGTCTAGCGCCTGCTGCCATGCGCGGGCGGCAAGCGCGCGCTCGTATGGGTTGA
GTGGGGGACCCCATGGCATGGGGTGGGTGAGCGCGGAGGCGTACATGCCGCAAATGTCGTAAAC

GTAGAGGGGCTCTCTGAGTATTCCAAGATATGTAGGGTAGCATCTTCCACCGCGGATGCTGGCGCG
CACGTAATCGTATAGTTCGTGCGAGGGAGCGAGGAGGTCGGGACCGAGGTTGCTACGGGCGGGCT
GCTCTGCTCGGAAGACTATCTGCCTGAAGATGGCATGTGAGTTGGATGATATGGTTGGACGCTGGA
AGACGTTGAAGCTGGCGTCTGTGAGACCTACCGCGTCACGCACGAAGGAGGCGTAGGAGTCGCG
CAGCTTGTTGACCAGCTCGGCGGTGACCTGCACGTCTAGGGCGCAGTAGTCCAGGGTTTCCTTGA
TGATGTCATACTTATCCTGTCCCTTTTTTTTCCACAGCTCGCGGTTGAGGACAAACTCTTCGCGGTCT
TTCCAGTACTCTTGGATCGGAAACCCGTCGGCCTCCGAACGGTAAGAGCCTAGCATGTAGAACTGG
TTGACGGCCTGGTAGGCGCAGCATCCCTTTTCTACGGGTAGCGCGTATGCCTGCGCGGCCTTCCG
GAGCGAGGTGTGGGTGAGCGCAAAGGTGTCCCTGACCATGACTTTGAGGTACTGGTATTTGAAGTC
AGTGTCGTCGCATCCGCCCTGCTCCCAGAGCAAAAAGTCCGTGCGCTTTTTGGAACGCGGATTTGG
CAGGGCGAAGGTGACATCGTTGAAGAGTATCTTTCCCGCGCGAGGCATAAAGTTGCGTGTGATGCG
GAAGGGTCCCGGCACCTCGGAACGGTTGTTAATTACCTGGGCGGCGAGCACGATCTCGTCAAAGC
CGTTGATGTTGTGGCCCACAATGTAAAGTTCCAAGAAGCGCGGGATGCCCTTGATGGAAGGCAATT
TTTTAAGTTCCTCGTAGGTGAGCTCTTCAGGGGAGCTGAGCCCGTGCTCTGAAAGGGCCCAGTCTG
CAAGATGAGGGTTGGAAGCGACGAATGAGCTCCACAGGTCACGGGCCATTAGCATTTGCAGGTGG
TCGCGAAAGGTCCTAAACTGGCGACCTATGGCCATTTTTTCTGGGGTGATGCAGTAGAAGGTAAGC
GGGTCTTGTTCCCAGCGGTCCCATCCAAGGTTCGCGGCTAGGTCTCGCGCGGCAGTCACTAGAGG
CTCATCTCCGCCGAACTTCATGACCAGCATGAAGGGCACGAGCTGCTTCCCAAAGGCCCCCATCCA
AGTATAGGTCTCTACATCGTAGGTGACAAAGAGACGCTCGGTGCGAGGATGCGAGCCGATCGGGAA
GAACTGGATCTCCCGCCACCAATTGGAGGAGTGGCTATTGATGTGGTGAAAGTAGAAGTCCCTGCG
ACGGGCCGAACACTCGTGCTGGCTTTTGTAAAAACGTGCGCAGTACTGGCAGCGGTGCACGGGCT
GTACATCCTGCACGAGGTTGACCTGACGACCGCGCACAAGGAAGCAGAGTGGGAATTTGAGCCCC
TCGCCTGGCGGGTTTGGCTGGTGGTCTTCTACTTCGGCTGCTTGTCCTTGACCGTCTGGCTGCTCG
AGGGGAGTTACGGTGGATCGGACCACCACGCCGCGCGAGCCCAAAGTCCAGATGTCCGCGCGCG
GCGGTCGGAGCTTGATGACAACATCGCGCAGATGGGAGCTGTCCATGGTCTGGAGCTCCCGCGGC
GTCAGGTCAGGCGGGAGCTCCTGCAGGTTTACCTCGCATAGACGGGTCAGGGCGCGGGCTAGATC
CAGGTGATACCTAATTTCCAGGGGCTGGTTGGTGGCGGCGTCGATGGCTTGCAAGAGGCCGCATC
CCCGCGGCGCGACTACGGTACCGCGCGGCGGGCGGTGGGCCGCGGGGGTGTCCTTGGATGATG
CATCTAAAAGCGGTGACGCGGGCGAGCCCCCGGAGGTAGGGGGGGCTCCGGACCCGCCGGGAG
AGGGGGCAGGGGCACGTCGGCGCCGCGCGCGGGCAGGAGCTGGTGCTGCGCGCGTAGGTTGCT
GGCGAACGCGACGACGCGGCGGTTGATCTCCTGAATCTGGCGCCTCTGCGTGAAGACGACGGGC
CCGGTGAGCTTGAACCTGAAAGAGAGTTCGACAGAATCAATTTCGGTGTCGTTGACGGCGGCCTG
GCGCAAAATCTCCTGCACGTCTCCTGAGTTGTCTTGATAGGCGATCTCGGCCATGAACTGCTCGATC
TCTTCCTCCTGGAGATCTCCGCGTCCGGCTCGCTCCACGGTGGCGGCGAGGTCGTTGGAAATGCG
GGCCATGAGCTGCGAGAAGGCGTTGAGGCCTCCCTCGTTCCAGACGCGGCTGTAGACCACGCCC
CCTTCGGCATCGCGGGCGCGCATGACCACCTGCGCGAGATTGAGCTCCACGTGCCGGGCGAAGA
CGGCGTAGTTTCGCAGGCGCTGAAAGAGGTAGTTGAGGGTGGTGGCGGTGTGTTCTGCCACGAAG
AAGTACATAACCCAGCGTCGCAACGTGGATTCGTTGATATCCCCCAAGGCCTCAAGGCGCTCCATG
GCCTCGTAGAAGTCCACGGCGAAGTTGAAAAACTGGGAGTTGCGCGCCGACACGGTTAACTCCTC
CTCCAGAAGACGGATGAGCTCGGCGACAGTGTCGCGCACCTCGCGCTCAAAGGCTACAGGGGCC
TCTTCTTCTTCTTCAATCTCCTCTTCCATAAGGGCCTCCCCTTCTTCTTCTTCTGGCGGCGGTGGGG
GAGGGGGGACACGGCGGCGACGACGGCGCACCGGGAGGCGGTCGACAAAGCGCTCGATCATCT
CCCCGCGGCGACGGCGCATGGTCTCGGTGACGGCGCGGCCGTTCTCGCGGGGGCGCAGTTGGA
AGACGCCGCCCGTCATGTCCCGGTTATGGGTTGGCGGGGGGCTGCCATGCGGCAGGGATACGGC

GCTAACGATGCATCTCAACAATTGTTGTGTAGGTACTCCGCCGCCGAGGGACCTGAGCGAGTCCGC
ATCGACCGGATCGGAAAACCTCTCGAGAAAGGCGTCTAACCAGTCACAGTCGCAAGGTAGGCTGA
GCACCGTGGCGGGCGGCAGCGGGCGGCGGTCGGGGTTGTTTCTGGCGGAGGTGCTGCTGATGAT
GTAATTAAAGTAGGCGGTCTTGAGACGGCGGATGGTCGACAGAAGCACCATGTCCTTGGGTCCGGC
CTGCTGAATGCGCAGGCGGTCGGCCATGCCCCAGGCTTCGTTTTGACATCGGCGCAGGTCTTTGT
AGTAGTCTTGCATGAGCCTTTCTACCGGCACTTCTTCTTCTCCTTCCTCTTGTCCTGCATCTCTTGCA
TCTATCGCTGCGGCGGCGGCGGAGTTTGGCCGTAGGTGGCGCCCTCTTCCTCCCATGCGTGTGAC
CCCGAAGCCCCTCATCGGCTGAAGCAGGGCTAGGTCGGCGACAACGCGCTCGGCTAATATGGCCT
GCTGCACCTGCGTGAGGGTAGACTGGAAGTCATCCATGTCCACAAAGCGGTGGTATGCGCCCGTG
TTGATGGTGTAAGTGCAGTTGGCCATAACGGACCAGTTAACGGTCTGGTGACCCGGCTGCGAGAGC
TCGGTGTACCTGAGACGCGAGTAAGCCCTCGAGTCAAATACGTAGTCGTTGCAAGTCCGCACCAGG
TACTGGTATCCCACCAAAAAGTGCGGCGGCGGCTGGCGGTAGAGGGGCCAGCGTAGGGTGGCCG
GGGCTCCGGGGGCGAGATCTTCCAACATAAGGCGATGATATCCGTAGATGTACCTGGACATCCAGG
TGATGCCGGCGGCGGTGGTGGAGGCGCGCGGAAAGTCGCGGACGCGGTTCCAGATGTTGCGCAG
CGGCAAAAAGTGCTCCATGGTCGGGACGCTCTGGCCGGTCAGGCGCGCGCAATCGTTGACGCTCT
AGCGTGCAAAAGGAGAGCCTGTAAGCGGGCACTCTTCCGTGGTCTGGTGGATAAATTCGCAAGGGT
ATCATGGCGGACGACCGGGGTTCGAGCCCCGTATCCGGCCGTCCGCCGTGATCCATGCGGTTACC
GCCCGCGTGTCGAACCCAGGTGTGCGACGTCAGACAACGGGGGAGTGCTCCTTTTGGCTTCCTTC
CAGGCGCGGCGGCTGCTGCGCTAGCTTTTTTGGCCACTGGCCGCGCGCAGCGTAAGCGGTTAGG
CTGGAAAGCGAAAGCATTAAGTGGCTCGCTCCCTGTAGCCGGAGGGTTATTTTCCAAGGGTTGAGT
CGCGGGACCCCCGGTTCGAGTCTCGGACCGGCCGGACTGCGGCGAACGGGGGTTTGCCTCCCC
GTCATGCAAGACCCCGCTTGCAAATTCCTCCGGAAACAGGGACGAGCCCCTTTTTTGCTTTTCCCA
GATGCATCCGGTGCTGCGGCAGATGCGCCCCCCTCCTCAGCAGCGGCAAGAGCAAGAGCAGCGG
CAGACATGCAGGGCACCCTCCCCTCCTCCTACCGCGTCAGGAGGGGCGACATCCGCGGTTGACG
CGGCAGCAGATGGTGATTACGAACCCCCGCGGCGCCGGGCCCGGCACTACCTGGACTTGGAGGA
GGGCGAGGGCCTGGCGCGGCTAGGAGCGCCCTCTCCTGAGCGGCACCCAAGGGTGCAGCTGAA
GCGTGATACGCGTGAGGCGTACGTGCCGCGGCAGAACCTGTTTCGCGACCGCGAGGGAGAGGAG
CCCGAGGAGATGCGGGATCGAAAGTTCCACGCAGGGCGCGAGCTGCGGCATGGCCTGAATCGCG
AGCGGTTGCTGCGCGAGGAGGACTTTGAGCCCGACGCGCGAACCGGGATTAGTCCCGCGCGCGC
ACACGTGGCGGCCGCCGACCTGGTAACCGCATACGAGCAGACGGTGAACCAGGAGATTAACTTTC
AAAAAAGCTTTAACAACCACGTGCGTACGCTTGTGGCGCGCGAGGAGGTGGCTATAGGACTGATGC
ATCTGTGGGACTTTGTAAGCGCGCTGGAGCAAAACCCAAATAGCAAGCCGCTCATGGCGCAGCTGT
TCCTTATAGTGCAGCACAGCAGGGACAACGAGGCATTCAGGGATGCGCTGCTAAACATAGTAGAGC
CCGAGGGCCGCTGGCTGCTCGATTTGATAAACATCCTGCAGAGCATAGTGGTGCAGGAGCGCAGC
TTGAGCCTGGCTGACAAGGTGGCCGCCATCAACTATTCCATGCTTAGCCTGGGCAAGTTTTACGCC
CGCAAGATATACCATACCCCTTACGTTCCCATAGACAAGGAGGTAAAGATCGAGGGGTTCTACATGC
GCATGGCGCTGAAGGTGCTTACCTTGAGCGACGACCTGGGCGTTTATCGCAACGAGCGCATCCAC
AAGGCCGTGAGCGTGAGCCGGCGGCGCGAGCTCAGCGACCGCGAGCTGATGCACAGCCTGCAAA
GGGCCCTGGCTGGCACGGGCAGCGGCGATAGAGAGGCCGAGTCCTACTTTGACGCGGGCGCTGA
CCTGCGCTGGGCCCCAAGCCGACGCGCCCTGGAGGCAGCTGGGGCCGGACCTGGGCTGGCGGT
GGCACCCGCGCGCGCTGGCAACGTCGGCGGCGTGGAGGAATATGACGAGGACGATGAGTACGAG
CCAGAGGACGGCGAGTACTAAGCGGTGATGTTTCTGATCAGATGATGCAAGACGCAACGGACCCG
GCGGTGCGGGCGGCGCTGCAGAGCCAGCCGTCCGGCCTTAACTCCACGGACGACTGGCGCCAG
GTCATGGACCGCATCATGTCGCTGACTGCGCGCAATCCTGACGCGTTCCGGCAGCAGCCGCAGGC

CAACCGGCTCTCCGCAATTCTGGAAGCGGTGGTCCCGGCGCGCGCAAACCCCACGCACGAGAAG
GTGCTGGCGATCGTAAACGCGCTGGCCGAAAACAGGGCCATCCGGCCCGACGAGGCCGGCCTGG
TCTACGACGCGCTGCTTCAGCGCGTGGCTCGTTACAACAGCGGCAACGTGCAGACCAACCTGGAC
CGGCTGGTGGGGGATGTGCGCGAGGCCGTGGCGCAGCGTGAGCGCGCGCAGCAGCAGGGCAAC
CTGGGCTCCATGGTTGCACTAAACGCCTTCCTGAGTACACAGCCCGCCAACGTGCCGCGGGGACA
GGAGGACTACACCAACTTTGTGAGCGCACTGCGGCTAATGGTGACTGAGACACCGCAAAGTGAGG
TGTACCAGTCTGGGCCAGACTATTTTTTCCAGACCAGTAGACAAGGCCTGCAGACCGTAAACCTGA
GCCAGGCTTTCAAAAACTTGCAGGGGCTGTGGGGGGTGCGGGCTCCCACAGGCGACCGCGCGAC
CGTGTCTAGCTTGCTGACGCCCAACTCGCGCCTGTTGCTGCTGCTAATAGCGCCCTTCACGGACAG
TGGCAGCGTGTCCCGGGACACATACCTAGGTCACTTGCTGACACTGTACCGCGAGGCCATAGGTCA
GGCGCATGTGGACGAGCATACTTTCCAGGAGATTACAAGTGTCAGCCGCGCGCTGGGGCAGGAGG
ACACGGGCAGCCTGGAGGCAACCCTAAACTACCTGCTGACCAACCGGCGGCAGAAGATCCCCTCG
TTGCACAGTTTAAACAGCGAGGAGGAGCGCATTTTGCGCTACGTGCAGCAGAGCGTGAGCCTTAAC
CTGATGCGCGACGGGGTAACGCCCAGCGTGGCGCTGGACATGACCGCGCGCAACATGGAACCGG
GCATGTATGCCTCAAACCGGCCGTTTATCAACCGCCTAATGGACTACTTGCATCGCGCGGCCGCCG
TGAACCCCGAGTATTTCACCAATGCCATCTTGAACCCGCACTGGCTACCGCCCCCTGGTTTCTACAC
CGGGGGATTCGAGGTGCCCGAGGGTAACGATGGATTCCTCTGGGACGACATAGACGACAGCGTGT
TTTCCCCGCAACCGCAGACCCTGCTAGAGTTGCAACAGCGCGAGCAGGCAGAGGCGGCGCTGCG
AAAGGAAAGCTTCCGCAGGCCAAGCAGCTTGTCCGATCTAGGCGCTGCGGCCCCGCGGTCAGATG
CTAGTAGCCCATTTCCAAGCTTGATAGGGTCTCTTACCAGCACTCGCACCACCCGCCCGCGCCTGC
TGGGCGAGGAGGAGTACCTAAACAACTCGCTGCTGCAGCCGCAGCGCGAAAAAAACCTGCCTCCG
GCATTTCCCAACAACGGGATAGAGAGCCTAGTGGACAAGATGAGTAGATGGAAGACGTACGCGCAG
GAGCACAGGGACGTGCCAGGCCCGCGCCCGCCCACCCGTCGTCAAAGGCACGACCGTCAGCGG
GGTCTGGTGTGGGAGGACGATGACTCGGCAGACGACAGCAGCGTCCTGGATTTGGGAGGGAGTG
GCAACCCGTTTGCGCACCTTCGCCCCAGGCTGGGGAGAATGTTTTAAAAAAAAAGCATGATGCAAA
ATAAAAAACTCACCAAGGCCATGGCACCGAGCGTTGGTTTTCTTGTATTCCCCTTAGTATGCGGCGC
GCGGCGATGTATGAGGAAGGTCCTCCTCCCTCCTACGAGAGTGTGGTGAGCGCGGCGCCAGTGG
CGGCGGCGCTGGGTTCTCCCTTCGATGCTCCCCTGGACCCGCCGTTTGTGCCTCCGCGGTACCTG
CGGCCTACCGGGGGGAGAAACAGCATCCGTTACTCTGAGTTGGCACCCCTATTCGACACCACCCGT
GTGTACCTGGTGGACAACAAGTCAACGGATGTGGCATCCCTGAACTACCAGAACGACCACAGCAAC
TTTCTGACCACGGTCATTCAAAACAATGACTACAGCCCGGGGGAGGCAAGCACACAGACCATCAAT
CTTGACGACCGGTCGCACTGGGGCGGCGACCTGAAAACCATCCTGCATACCAACATGCCAAATGTG
AACGAGTTCATGTTTACCAATAAGTTTAAGGCGCGGGTGATGGTGTCGCGCTTGCCTACTAAGGACA
ATCAGGTGGAGCTGAAATACGAGTGGGTGGAGTTCACGCTGCCCGAGGGCAACTACTCCGAGACC
ATGACCATAGACCTTATGAACAACGCGATCGTGGAGCACTACTTGAAAGTGGGCAGACAGAACGGG
GTTCTGGAAAGCGACATCGGGGTAAAGTTTGACACCCGCAACTTCAGACTGGGGTTTGACCCCGTC
ACTGGTCTTGTCATGCCTGGGGTATATACAAACGAAGCCTTCCATCCAGACATCATTTTGCTGCCAG
GATGCGGGGTGGACTTCACCCACAGCCGCCTGAGCAACTTGTTGGGCATCCGCAAGCGGCAACCC
TTCCAGGAGGGCTTTAGGATCACCTACGATGATCTGGAGGGTGGTAACATTCCCGCACTGTTGGAT
GTGGACGCCTACCAGGCGAGCTTGAAAGATGACACCGAACAGGGCGGGGGTGGCGCAGGCGGCA
GCAACAGCAGTGGCAGCGGCGCGGAAGAGAACTCCAACGCGGCAGCCGCGGCAATGCAGCCGG
TGGAGGACATGAACGATCATGCCATTCGCGGCGACACCTTTGCCACACGGGCTGAGGAGAAGCGC
GCTGAGGCCGAAGCAGCGGCCGAAGCTGCCGCCCCCGCTGCGCAACCCGAGGTCGAGAAGCCT
CAGAAGAAACCGGTGATCAAACCCCTGACAGAGGACAGCAAGAAACGCAGTTACAACCTAATAAGC

AATGACAGCACCTTCACCCAGTACCGCAGCTGGTACCTTGCATACAACTACGGCGACCCTCAGACC
GGAATCCGCTCATGGACCCTGCTTTGCACTCCTGACGTAACCTGCGGCTCGGAGCAGGTCTACTG
GTCGTTGCCAGACATGATGCAAGACCCCGTGACCTTCCGCTCCACGCGCCAGATCAGCAACTTTCC
GGTGGTGGGCGCCGAGCTGTTGCCCGTGCACTCCAAGAGCTTCTACAACGACCAGGCCGTCTACT
CCCAACTCATCCGCCAGTTTACCTCTCTGACCCACGTGTTCAATCGCTTTCCCGAGAACCAGATTTT
GGCGCGCCCGCCAGCCCCCACCATCACCACCGTCAGTGAAAACGTTCCTGCTCTCACAGATCACG
GGACGCTACCGCTGCGCAACAGCATCGGAGGAGTCCAGCGAGTGACCATTACTGACGCCAGACGC
CGCACCTGCCCCTACGTTTACAAGGCCCTGGGCATAGTCTCGCCGCGCGTCCTATCGAGCCGCACT
TTTTGAGCAAGCATGTCCATCCTTATATCGCCCAGCAATAACACAGGCTGGGGCCTGCGCTTCCCAA
GCAAGATGTTTGGCGGGGCCAAGAAGCGCTCCGACCAACACCCAGTGCGCGTGCGCGGGCACTA
CCGCGCGCCCTGGGGCGCGCACAAACGCGGCCGCACTGGGCGCACCACCGTCGATGACGCCATC
GACGCGGTGGTGGAGGAGGCGCGCAACTACACGCCCACGCCGCCACCAGTGTCCACAGTGGACG
CGGCCATTCAGACCGTGGTGCGCGGAGCCCGGCGCTATGCTAAAATGAAGAGACGGCGGAGGCG
CGTAGCACGTCGCCACCGCCGCCGACCCGGCACTGCCGCCCAACGCGCGGCGGCGGCCCTGCTT
AACCGCGCACGTCGCACCGGCCGACGGGCGGCCATGCGGGCCGCTCGAAGGCTGGCCGCGGGT
ATTGTCACTGTGCCCCCCAGGTCCAGGCGACGAGCGGCCGCCGCAGCAGCCGCGGCCATTAGTG
CTATGACTCAGGGTCGCAGGGGCAACGTGTATTGGGTGCGCGACTCGGTTAGCGGCCTGCGCGTG
CCCGTGCGCACCCGCCCCCCGCGCAACTAGATTGCAAGAAAAAACTACTTAGACTCGTACTGTTGT
ATGTATCCAGCGGCGGCGGCGCGCAACGAAGCTATGTCCAAGCGCAAAATCAAAGAAGAGATGCTC
CAGGTCATCGCGCCGGAGATCTATGGCCCCCCGAAGAAGGAAGAGCAGGATTACAAGCCCCGAAA
GCTAAAGCGGGTCAAAAAGAAAAAGAAAGATGATGATGATGAACTTGACGACGAGGTGGAACTGCT
GCACGCTACCGCGCCCAGGCGACGGGTACAGTGGAAAGGTCGACGCGTAAAACGTGTTTTGCGAC
CCGGCACCACCGTAGTCTTTACGCCCGGTGAGCGCTCCACCCGCACCTACAAGCGCGTGTATGAT
GAGGTGTACGGCGACGAGGACCTGCTTGAGCAGGCCAACGAGCGCCTCGGGGAGTTTGCCTACG
GAAAGCGGCATAAGGACATGCTGGCGTTGCCGCTGGACGAGGGCAACCCAACACCTAGCCTAAAG
CCCGTAACACTGCAGCAGGTGCTGCCCGCGCTTGCACCGTCCGAAGAAAAGCGCGGCCTAAAGCG
CGAGTCTGGTGACTTGGCACCCACCGTGCAGCTGATGGTACCCAAGCGCCAGCGACTGGAAGATG
TCTTGGAAAAAATGACCGTGGAACCTGGGCTGGAGCCCGAGGTCCGCGTGCGGCCAATCAAGCAG
GTGGCGCCGGGACTGGGCGTGCAGACCGTGGACGTTCAGATACCCACTACCAGTAGCACCAGTAT
TGCCACCGCCACAGAGGGCATGGAGACACAAACGTCCCCGGTTGCCTCAGCGGTGGCGGATGCC
GCGGTGCAGGCGGTCGCTGCGGCCGCGTCCAAGACCTCTACGGAGGTGCAAACGGACCCGTGGA
TGTTTCGCGTTTCAGCCCCCCGGCGCCCGCGCCGTTCGAGGAAGTACGGCGCCGCCAGCGCGCT
ACTGCCCGAATATGCCCTACATCCTTCCATTGCGCCTACCCCCGGCTATCGTGGCTACACCTACCGC
CCCAGAAGACGAGCAACTACCCGACGCCGAACCACCACTGGAACCCGCCGCCGCCGTCGCCGTC
GCCAGCCCGTGCTGGCCCCGATTTCCGTGCGCAGGGTGGCTCGCGAAGGAGGCAGGACCCTGGT
GCTGCCAACAGCGCGCTACCACCCCAGCATCGTTTAAAAGCCGGTCTTTGTGGTTCTTGCAGATAT
GGCCCTCACCTGCCGCCTCCGTTTCCCGGTGCCGGGATTCCGAGGAAGAATGCACCGTAGGAGG
GGCATGGCCGGCCACGGCCTGACGGGCGGCATGCGTCGTGCGCACCACCGGCGGCGGCGCGCG
TCGCACCGTCGCATGCGCGGCGGTATCCTGCCCCTCCTTATTCCACTGATCGCCGCGGCGATTGGC
GCCGTGCCCGGAATTGCATCCGTGGCCTTGCAGGCGCAGAGACACTGATTAAAAACAAGTTGCATG
TGGAAAAATCAAAATAAAAAGTCTGGACTCTCACGCTCGCTTGGTCCTGTAACTATTTTGTAGAATGG
AAGACATCAACTTTGCGTCTCTGGCCCCGCGACACGGCTCGCGCCCGTTCATGGGAAACTGGCAA
GATATCGGCACCAGCAATATGAGCGGTGGCGCCTTCAGCTGGGGCTCGCTGTGGAGCGGCATTAAA
AATTTCGGTTCCACCGTTAAGAACTATGGCAGCAAGGCCTGGAACAGCAGCACAGGCCAGATGCTG

AGGGATAAGTTGAAAGAGCAAAATTTCCAACAAAAGGTGGTAGATGGCCTGGCCTCTGGCATTAGC
GGGGTGGTGGACCTGGCCAACCAGGCAGTGCAAAATAAGATTAACAGTAAGCTTGATCCCCGCCCT
CCCGTAGAGGAGCCTCCACCGGCCGTGGAGACAGTGTCTCCAGAGGGGCGTGGCGAAAAGCGTC
CGCGCCCCGACAGGGAAGAAACTCTGGTGACGCAAATAGACGAGCCTCCCTCGTACGAGGAGGCA
CTAAAGCAAGGCCTGCCCACCACCCGTCCCATCGCGCCCATGGCTACCGGAGTGCTGGGCCAGCA
CACACCCGTAACGCTGGACCTGCCTCCCCCCGCCGACACCCAGCAGAAACCTGTGCTGCCAGGC
CCGACCGCCGTTGTTGTAACCCGTCCTAGCCGCGCGTCCCTGCGCCGCGCCGCCAGCGGTCCGC
GATCGTTGCGGCCCGTAGCCAGTGGCAACTGGCAAAGCACACTGAACAGCATCGTGGGTCTGGGG
GTGCAATCCCTGAAGCGCCGACGATGCTTCTGATAGCTAACGTGTCGTATGTGTGTCATGTATGCGT
CCATGTCGCCGCCAGAGGAGCTGCTGAGCCGCCGCGCGCCCGCTTTCCAAGATGGCTACCCCTTC
GATGATGCCGCAGTGGTCTTACATGCACATCTCGGGCCAGGACGCCTCGGAGTACCTGAGCCCCG
GGCTGGTGCAGTTTGCCCGCGCCACCGAGACGTACTTCAGCCTGAATAACAAGTTTAGAAACCCCA
CGGTGGCGCCTACGCACGACGTGACCACAGACCGGTCCCAGCGTTTGACGCTGCGGTTCATCCCT
GTGGACCGTGAGGATACTGCGTACTCGTACAAGGCGCGGTTCACCCTAGCTGTGGGTGATAACCGT
GTGCTGGACATGGCTTCCACGTACTTTGACATCCGCGGCGTGCTGGACAGGGGCCCTACTTTTAAG
CCCTACTCTGGCACTGCCTACAACGCCCTGGCTCCCAAGGGTGCCCCAAATCCTTGCGAATGGGAT
GAAGCTGCTACTGCTCTTGAAATAAACCTAGAAGAAGAGGACGATGACAACGAAGACGAAGTAGAC
GAGCAAGCTGAGCAGCAAAAAACTCACGTATTTGGGCAGGCGCCTTATTCTGGTATAAATATTACAAA
GGAGGGTATTCAAATAGGTGTCGAAGGTCAAACACCTAAATATGCCGATAAAACATTTCAACCTGAAC
CTCAAATAGGAGAATCTCAGTGGTACGAAACAGAAATTAATCATGCAGCTGGGAGAGTCCTAAAAAA
GACTACCCCAATGAAACCATGTTACGGTTCATATGCAAAACCCACAAATGAAAATGGAGGGCAAGGC
ATTCTTGTAAAGCAACAAAATGGAAAGCTAGAAAGTCAAGTGGAAATGCAATTTTTCTCAACTACTGA
GGCAGCCGCAGGCAATGGTGATAACTTGACTCCTAAAGTGGTATTGTACAGTGAAGATGTAGATATA
GAAACCCCAGACACTCATATTTCTTACATGCCCACTATTAAGGAAGGTAACTCACGAGAACTAATGGG
CCAACAATCTATGCCCAACAGGCCTAATTACATTGCTTTTAGGGACAATTTTATTGGTCTAATGTATTAC
AACAGCACGGGTAATATGGGTGTTCTGGCGGGCCAAGCATCGCAGTTGAATGCTGTTGTAGATTTG
CAAGACAGAAACACAGAGCTTTCATACCAGCTTTTGCTTGATTCCATTGGTGATAGAACCAGGTACTT
TTCTATGTGGAATCAGGCTGTTGACAGCTATGATCCAGATGTTAGAATTATTGAAAATCATGGAACTGA
AGATGAACTTCCAAATTACTGCTTTCCACTGGGAGGTGTGATTAATACAGAGACTCTTACCAAGGTAA
AACCTAAAACAGGTCAGGAAAATGGATGGGAAAAAGATGCTACAGAATTTTCAGATAAAAATGAAATA
AGAGTTGGAAATAATTTTGCCATGGAAATCAATCTAAATGCCAACCTGTGGAGAAATTTCCTGTACTC
CAACATAGCGCTGTATTTGCCCGACAAGCTAAAGTACAGTCCTTCCAACGTAAAAATTTCTGATAACC
CAAACACCTACGACTACATGAACAAGCGAGTGGTGGCTCCCGGGCTAGTGGACTGCTACATTAACC
TTGGAGCACGCTGGTCCCTTGACTATATGGACAACGTCAACCCATTTAACCACCACCGCAATGCTGG
CCTGCGCTACCGCTCAATGTTGCTGGGCAATGGTCGCTATGTGCCCTTCCACATCCAGGTGCCTCA
GAAGTTCTTTGCCATTAAAAACCTCCTTCTCCTGCCGGGCTCATACACCTACGAGTGGAACTTCAGG
AAGGATGTTAACATGGTTCTGCAGAGCTCCCTAGGAAATGACCTAAGGGTTGACGGAGCCAGCATTA
AGTTTGATAGCATTTGCCTTTACGCCACCTTCTTCCCCATGGCCCACAACACCGCCTCCACGCTTGA
GGCCATGCTTAGAAACGACACCAACGACCAGTCCTTTAACGACTATCTCTCCGCCGCCAACATGCTC
TACCCTATACCCGCCAACGCTACCAACGTGCCCATATCCATCCCCTCCCGCAACTGGGCGGCTTTCC
GCGGCTGGGCCTTCACGCGCCTTAAGACTAAGGAAACCCCATCACTGGGCTCGGGCTACGACCCT
TATTACACCTACTCTGGCTCTATACCCTACCTAGATGGAACCTTTTACCTCAACCACACCTTTAAGAAG
GTGGCCATTACCTTTGACTCTTCTGTCAGCTGGCCTGGCAATGACCGCCTGCTTACCCCCAACGAG
TTTGAAATTAAGCGCTCAGTTGACGGGGAGGGTTACAACGTTGCCCAGTGTAACATGACCAAAGACT

GGTTCCTGGTACAAATGCTAGCTAACTATAACATTGGCTACCAGGGCTTCTATATCCCAGAGAGCTAC
AAGGACCGCATGTACTCCTTCTTTAGAAACTTCCAGCCCATGAGCCGTCAGGTGGTGGATGATACTA
AATACAAGGACTACCAACAGGTGGGCATCCTACACCAACACAACAACTCTGGATTTGTTGGCTACCT
TGCCCCCACCATGCGCGAAGGACAGGCCTACCCTGCTAACTTCCCCTATCCGCTTATAGGCAAGAC
CGCAGTTGACAGCATTACCCAGAAAAAGTTTCTTTGCGATCGCACCCTTTGGCGCATCCCATTCTCC
AGTAACTTTATGTCCATGGGCGCACTCACAGACCTGGGCCAAAACCTTCTCTACGCCAACTCCGCC
CACGCGCTAGACATGACTTTTGAGGTGGATCCCATGGACGAGCCCACCCTTCTTTATGTTTTGTTTG
AAGTCTTTGACGTGGTCCGTGTGCACCAGCCGCACCGCGGCGTCATCGAAACCGTGTACCTGCGC
ACGCCCTTCTCGGCCGGCAACGCCACAACATAAAGAAGCAAGCAACATCAACAACAGCTGCCGCCA
TGGGCTCCAGTGAGCAGGAACTGAAAGCCATTGTCAAAGATCTTGGTTGTGGGCCATATTTTTTGGG
CACCTATGACAAGCGCTTTCCAGGCTTTGTTTCTCCACACAAGCTCGCCTGCGCCATAGTCAATACG
GCCGGTCGCGAGACTGGGGGCGTACACTGGATGGCCTTTGCCTGGAACCCGCACTCAAAAACATG
CTACCTCTTTGAGCCCTTTGGCTTTTCTGACCAGCGACTCAAGCAGGTTTACCAGTTTGAGTACGAG
TCACTCCTGCGCCGTAGCGCCATTGCTTCTTCCCCCGACCGCTGTATAACGCTGGAAAAGTCCACC
CAAAGCGTACAGGGGCCCAACTCGGCCGCCTGTGGACTATTCTGCTGCATGTTTCTCCACGCCTTT
GCCAACTGGCCCCAAACTCCCATGGATCACAACCCCACCATGAACCTTATTACCGGGGTACCCAAC
TCCATGCTCAACAGTCCCCAGGTACAGCCCACCCTGCGTCGCAACCAGGAACAGCTCTACAGCTTC
CTGGAGCGCCACTCGCCCTACTTCCGCAGCCACAGTGCGCAGATTAGGAGCGCCACTTCTTTTTGT
CACTTGAAAAACATGTAAAAATAATGTACTAGAGACACTTTCAATAAAGGCAAATGCTTTTATTTGTACA
CTCTCGGGTGATTATTTACCCCCACCCTTGCCGTCTGCGCCGTTTAAAAATCAAAGGGGTTCTGCCG
CGCATCGCTATGCGCCACTGGCAGGGACACGTTGCGATACTGGTGTTTAGTGCTCCACTTAAACTC
AGGCACAACCATCCGCGGCAGCTCGGTGAAGTTTTCACTCCACAGGCTGCGCACCATCACCAACG
CGTTTAGCAGGTCGGGCGCCGATATCTTGAAGTCGCAGTTGGGGCCTCCGCCCTGCGCGCGCGAG
TTGCGATACACAGGGTTGCAGCACTGGAACACTATCAGCGCCGGGTGGTGCACGCTGGCCAGCAC
GCTCTTGTCGGAGATCAGATCCGCGTCCAGGTCCTCCGCGTTGCTCAGGGCGAACGGAGTCAACT
TTGGTAGCTGCCTTCCCAAAAAGGGCGCGTGCCCAGGCTTTGAGTTGCACTCGCACCGTAGTGGC
ATCAAAAGGTGACCGTGCCCGGTCTGGGCGTTAGGATACAGCGCCTGCATAAAAGCCTTGATCTGC
TTAAAAGCCACCTGAGCCTTTGCGCCTTCAGAGAAGAACATGCCGCAAGACTTGCCGGAAAACTGA
TTGGCCGGACAGGCCGCGTCGTGCACGCAGCACCTTGCGTCGGTGTTGGAGATCTGCACCACATT
TCGGCCCCACCGGTTCTTCACGATCTTGGCCTTGCTAGACTGCTCCTTCAGCGCGCGCTGCCCGTT
TTCGCTCGTCACATCCATTTCAATCACGTGCTCCTTATTTATCATAATGCTTCCGTGTAGACACTTAAG
CTCGCCTTCGATCTCAGCGCAGCGGTGCAGCCACAACGCGCAGCCCGTGGGCTCGTGATGCTTGT
AGGTCACCTCTGCAAACGACTGCAGGTACGCCTGCAGGAATCGCCCCATCATCGTCACAAAGGTCT
TGTTGCTGGTGAAGGTCAGCTGCAACCCGCGGTGCTCCTCGTTCAGCCAGGTCTTGCATACGGCC
GCCAGAGCTTCCACTTGGTCAGGCAGTAGTTTGAAGTTCGCCTTTAGATCGTTATCCACGTGGTACT
TGTCCATCAGCGCGCGCGCAGCCTCCATGCCCTTCTCCCACGCAGACACGATCGGCACACTCAGC
GGGTTCATCACCGTAATTTCACTTTCCGCTTCGCTGGGCTCTTCCTCTTCCTCTTGCGTCCGCATAC
CACGCGCCACTGGGTCGTCTTCATTCAGCCGCCGCACTGTGCGCTTACCTCCTTTGCCATGCTTGA
TTAGCACCGGTGGGTTGCTGAAACCCACCATTTGTAGCGCCACATCTTCTCTTTCTTCCTCGCTGTC
CACGATTACCTCTGGTGATGGCGGGCGCTCGGGCTTGGGAGAAGGGCGCTTCTTTTTCTTCTTGG
GCGCAATGGCCAAATCCGCCGCCGAGGTCGATGGCCGCGGGCTGGGTGTGCGCGGCACCAGCG
CGTCTTGTGATGAGTCTTCCTCGTCCTCGGACTCGATACGCCGCCTCATCCGCTTTTTTGGGGGCG
CCCGGGGAGGCGGCGGCGACGGGGACGGGGACGACACGTCCTCCATGGTTGGGGGACGTCGCG
CCGCACCGCGTCCGCGCTCGGGGGTGGTTTCGCGCTGCTCCTCTTCCCGACTGGCCATTTCCTTC

TCCTATAGGCAGAAAAAGATCATGGAGTCAGTCGAGAAGAAGGACAGCCTAACCGCCCCCTCTGAG
TTCGCCACCACCGCCTCCACCGATGCCGCCAACGCGCCTACCACCTTCCCCGTCGAGGCACCCCC
GCTTGAGGAGGAGGAAGTGATTATCGAGCAGGACCCAGGTTTTGTAAGCGAAGACGACGAGGACC
GCTCAGTACCAACAGAGGATAAAAAGCAAGACCAGGACAACGCAGAGGCAAACGAGGAACAAGTC
GGGCGGGGGGACGAAAGGCATGGCGACTACCTAGATGTGGGAGACGACGTGCTGTTGAAGCATCT
GCAGCGCCAGTGCGCCATTATCTGCGACGCGTTGCAAGAGCGCAGCGATGTGCCCCTCGCCATAG
CGGATGTCAGCCTTGCCTACGAACGCCACCTATTCTCACCGCGCGTACCCCCCAAACGCCAAGAAA
ACGGCACATGCGAGCCCAACCCGCGCCTCAACTTCTACCCCGTATTTGCCGTGCCAGAGGTGCTT
GCCACCTATCACATCTTTTTCCAAAACTGCAAGATACCCCTATCCTGCCGTGCCAACCGCAGCCGAG
CGGACAAGCAGCTGGCCTTGCGGCAGGGCGCTGTCATACCTGATATCGCCTCGCTCAACGAAGTG
CCAAAAATCTTTGAGGGTCTTGGACGCGACGAGAAGCGCGCGGCAAACGCTCTGCAACAGGAAAA
CAGCGAAAATGAAAGTCACTCTGGAGTGTTGGTGGAACTCGAGGGTGACAACGCGCGCCTAGCCG
TACTAAAACGCAGCATCGAGGTCACCCACTTTGCCTACCCGGCACTTAACCTACCCCCCAAGGTCAT
GAGCACAGTCATGAGTGAGCTGATCGTGCGCCGTGCGCAGCCCCTGGAGAGGGATGCAAATTTGC
AAGAACAAACAGAGGAGGGCCTACCCGCAGTTGGCGACGAGCAGCTAGCGCGCTGGCTTCAAACG
CGCGAGCCTGCCGACTTGGAGGAGCGACGCAAACTAATGATGGCCGCAGTGCTCGTTACCGTGGA
GCTTGAGTGCATGCAGCGGTTCTTTGCTGACCCGGAGATGCAGCGCAAGCTAGAGGAAACATTGCA
CTACACCTTTCGACAGGGCTACGTACGCCAGGCCTGCAAGATCTCCAACGTGGAGCTCTGCAACCT
GGTCTCCTACCTTGGAATTTTGCACGAAAACCGCCTTGGGCAAAACGTGCTTCATTCCACGCTCAA
GGGCGAGGCGCGCCGCGACTACGTCCGCGACTGCGTTTACTTATTTCTATGCTACACCTGGCAGAC
GGCCATGGGCGTTTGGCAGCAGTGCTTGGAGGAGTGCAACCTCAAGGAGCTGCAGAAACTGCTAA
AGCAAAACTTGAAGGACCTATGGACGGCCTTCAACGAGCGCTCCGTGGCCGCGCACCTGGCGGAC
ATCATTTTCCCCGAACGCCTGCTTAAAACCCTGCAACAGGGTCTGCCAGACTTCACCAGTCAAAGCA
TGTTGCAGAACTTTAGGAACTTTATCCTAGAGCGCTCAGGAATCTTGCCCGCCACCTGCTGTGCACT
TCCTAGCGACTTTGTGCCCATTAAGTACCGCGAATGCCCTCCGCCGCTTTGGGGCCACTGCTACCT
TCTGCAGCTAGCCAACTACCTTGCCTACCACTCTGACATAATGGAAGACGTGAGCGGTGACGGTCT
ACTGGAGTGTCACTGTCGCTGCAACCTATGCACCCCGCACCGCTCCCTGGTTTGCAATTCGCAGCT
GCTTAACGAAAGTCAAATTATCGGTACCTTTGAGCTGCAGGGTCCCTCGCCTGACGAAAAGTCCGC
GGCTCCGGGGTTGAAACTCACTCCGGGGCTGTGGACGTCGGCTTACCTTCGCAAATTTGTACCTGA
GGACTACCACGCCCACGAGATTAGGTTCTACGAAGACCAATCCCGCCCGCCTAATGCGGAGCTTAC
CGCCTGCGTCATTACCCAGGGCCACATTCTTGGCCAATTGCAAGCCATCAACAAAGCCCGCCAAGA
GTTTCTGCTACGAAAGGGACGGGGGGTTTACTTGGACCCCCAGTCCGGCGAGGAGCTCAACCCAA
TCCCCCCGCCGCCGCAGCCCTATCAGCAGCAGCCGCGGGCCCTTGCTTCCCAGGATGGCACCCA
AAAAGAAGCTGCAGCTGCCGCCGCCACCCACGGACGAGGAGGAATACTGGGACAGTCAGGCAGA
GGAGGTTTTGGACGAGGAGGAGGAGGACATGATGGAAGACTGGGAGAGCCTAGACGAGGAAGCT
TCCGAGGTCGAAGAGGTGTCAGACGAAACACCGTCACCCTCGGTCGCATTCCCCTCGCCGGCGCC
CCAGAAATCGGCAACCGGTTCCAGCATGGCTACAACCTCCGCTCCTCAGGCGCCGCCGGCACTGC
CCGTTCGCCGACCCAACCGTAGATGGGACACCACTGGAACCAGGGCCGGTAAGTCCAAGCAGCCG
CCGCCGTTAGCCCAAGAGCAACAACAGCGCCAAGGCTACCGCTCATGGCGCGGGCACAAGAACG
CCATAGTTGCTTGCTTGCAAGACTGTGGGGGCAACATCTCCTTCGCCCGCCGCTTTCTTCTCTACCA
TCACGGCGTGGCCTTCCCCCGTAACATCCTGCATTACTACCGTCATCTCTACAGCCCATACTGCACC
GGCGGCAGCGGCAGCAACAGCAGCGGCCACACAGAAGCAAAGGCGACCGGATAGCAAGACTCTG
ACAAAGCCCAAGAAATCCACAGCGGCGGCAGCAGCAGGAGGAGGAGCGCTGCGTCTGGCGCCCA
ACGAACCCGTATCGACCCGCGAGCTTAGAAACAGGATTTTTCCCACTCTGTATGCTATATTTCAACAG

AGCAGGGGCCAAGAACAAGAGCTGAAAATAAAAAACAGGTCTCTGCGATCCCTCACCCGCAGCTG
CCTGTATCACAAAAGCGAAGATCAGCTTCGGCGCACGCTGGAAGACGCGGAGGCTCTCTTCAGTAA
ATACTGCGCGCTGACTCTTAAGGACTAGTTTCGCGCCCTTTCTCAAATTTAAGCGCGAAAACTACGT
CATCTCCAGCGGCCACACCCGGCGCCAGCACCTGTTGTCAGCGCCATTATGAGCAAGGAAATTCCC
ACGCCCTACATGTGGAGTTACCAGCCACAAATGGGACTTGCGGCTGGAGCTGCCCAAGACTACTCA
ACCCGAATAAACTACATGAGCGCGGGACCCCACATGATATCCCGGGTCAACGGAATACGCGCCCAC
CGAAACCGAATTCTCCTGGAACAGGCGGCTATTACCACCACACCTCGTAATAACCTTAATCCCCGTA
GTTGGCCCGCTGCCCTGGTGTACCAGGAAAGTCCCGCTCCCACCACTGTGGTACTTCCCAGAGAC
GCCCAGGCCGAAGTTCAGATGACTAACTCAGGGGCGCAGCTTGCGGGCGGCTTTCGTCACAGGGT
GCGGTCGCCCGGGCAGGGTATAACTCACCTGACAATCAGAGGGCGAGGTATTCAGCTCAACGACG
AGTCGGTGAGCTCCTCGCTTGGTCTCCGTCCGGACGGGACATTTCAGATCGGCGGCGCCGGCCG
CTCTTCATTCACGCCTCGTCAGGCAATCCTAACTCTGCAGACCTCGTCCTCTGAGCCGCGCTCTGG
AGGCATTGGAACTCTGCAATTTATTGAGGAGTTTGTGCCATCGGTCTACTTTAACCCCTTCTCGGGA
CCTCCCGGCCACTATCCGGATCAATTTATTCCTAACTTTGACGCGGTAAAGGACTCGGCGGACGGCT
ACGACTGAATGTTAAGTGGAGAGGCAGAGCAACTGCGCCTGAAACACCTGGTCCACTGTCGCCGC
CACAAGTGCTTTGCCCGCGACTCCGGTGAGTTTTGCTACTTTGAATTGCCCGAGGATCATATCGAGG
GCCCGGCGCACGGCGTCCGGCTTACCGCCCAGGGAGAGCTTGCCCGTAGCCTGATTCGGGAGTT
TACCCAGCGCCCCCTGCTAGTTGAGCGGGACAGGGGACCCTGTGTTCTCACTGTGATTTGCAACTG
TCCTAACCCTGGATTACATCAAGATCCTCTAGTTAATGGATCCAATCGATATCGCGAAGTCGATCGAC
GGATCGGGAGATCTCCCGATCCCCTATGGTGCACTCTCAGTACAATCTGCTCTGATGCCGCATAGTT
AAGCCAGTATCTGCTCCCTGCTTGTGTGTTGGAGGTCGCTGAGTAGTGCGCGAGCAAAATTTAAGC
TACAACAAGGCAAGGCTTGACCGACAATTGCATGAAGAATCTGCTTAGGGTTAGGCGTTTTGCGCTG
CTTCGGCTAGCTGCAGTAACGCCATTTTGCAAGGCATGGAAAAATACCAAACCAAGAATAGAGAAGT
TCAGATCAAGGGCGGGTACATGAAAATAGCTAACGTTGGGCCAAACAGGATATCTGCGGTGAGCAG
TTTCGGCCCCGGCCCGGGGCCAAGAACAGATGGTCACCGCAGTTTCGGCCCCGGCCCGAGGCCA
AGAACAGATGGTCCCCAGATATGGCCCAACCCTCAGCAGTTTCTTAAGACCCATCAGATGTTTCCAG
GCTCCCCCAAGGACCTGAAATGACCCTGCGCCTTATTTGAATTAACCAATCAGCCTGCTTCTCGCTT
CTGTTCGCGCGCTTCTGCTTCCCGAGCTCTATAAAAGAGCTCACAACCCCTCACTCGGCGCGCCAG
TCCTCCGACAGACTGAGTCGCCCGGGTACCGCGATCGCCATGGCGGGACACCTGGCTTCGGATTT
CGCCTTCTCGCCCCCTCCAGGTGGTGGAGGTGATGGGCCAGGGGGGCCGGAGCCGGGCTGGGT
TGATCCTCGGACCTGGCTAAGCTTCCAAGGCCCTCCTGGAGGGCCAGGAATCGGGCCGGGGGTT
GGGCCAGGCTCTGAGGTGTGGGGGATTCCCCCATGCCCCCCGCCGTATGAGTTCTGTGGGGGGAT
GGCGTACTGTGGGCCCCAGGTTGGAGTGGGGCTAGTGCCCCAAGGCGGCTTGGAGACCTCTCAG
CCTGAGGGTGAAGCAGGAGTCGGGGTGGAGAGCAACTCCGATGGGGCCTCCCCGGAGCCCTGCA
CCGTCACCCCTGGTGCCGTGAAGCTGGAGAAGGAGAAGCTGGAGCAAAACCCGGAGGAGTCCCA
GGACATCAAAGCTCTGCAGAAAGAACTCGAGCAATTTGCCAAGCTCCTGAAGCAGAAGAGGATCAC
CCTGGGATATACACAGGCCGATGTGGGGCTCACCCTGGGGGTTCTATTTGGGAAGGTATTCAGCCA
AACGACCATCTGCCGCTTTGAGGCTCTGCAGCTTAGCTTCAAGAACATGTGTAAGCTGCGGCCCTT
GCTGCAGAAGTGGGTGGAGGAAGCTGACAACAATGAAAATCTTCAGGAGATATGCAAAGCAGAAAC
CCTCGTGCAGGCCCGAAAGAGAAAGCGAACCAGTATCGAGAACCGAGTGAGAGGCAACCTGGAGA
ATTTGTTCCTGCAGTGCCCGAAACCCACACTGCAGCAGATCAGCCACATCGCCCAGCAGCTTGGGC
TCGAGAAGGATGTGGTCCGAGTGTGGTTCTGTAACCGGCGCCAGAAGGGCAAGCGATCAAGCAGC
GACTATGCACAACGAGAGGATTTTGAGGCTGCTGGGTCTCCTTTCTCAGGGGGACCAGTGTCCTTT
CCTCTGGCCCCAGGGCCCCATTTTGGTACCCCAGGCTATGGGAGCCCTCACTTCACTGCACTGTAC

TCCTCGGTCCCTTTCCCTGAGGGGGAAGCCTTTCCCCCTGTCTCCGTCACCACTCTGGGCTCTCC
CATGCATTCAAACGGGGGAGGAGGCTCCATGGTGAGCAAGGGCGAGGAGGATAACATGGCCATCA
TCAAGGAGTTCATGCGCTTCAAGGTGCACATGGAGGGCTCCGTGAACGGCCACGAGTTCGAGATC
GAGGGCGAGGGCGAGGGCCGCCCCTACGAGGGCACCCAGACCGCCAAGCTGAAGGTGACCAAG
GGTGGCCCCCTGCCCTTCGCCTGGGACATCCTGTCCCCTCAGTTCATGTACGGCTCCAAGGCCTA
CGTGAAGCACCCCGCCGACATCCCCGACTACTTGAAGCTGTCCTTCCCCGAGGGCTTCAAGTGGG
AGCGCGTGATGAACTTCGAGGACGGCGGCGTGGTGACCGTGACCCAGGACTCCTCCCTGCAGGA
CGGCGAGTTCATCTACAAGGTGAAGCTGCGCGGCACCAACTTCCCCTCCGACGGCCCCGTAATGC
AGAAGAAGACCATGGGCTGGGAGGCCTCCTCCGAGCGGATGTACCCCGAGGACGGCGCCCTGAA
GGGCGAGATCAAGCAGAGGCTGAAGCTGAAGGACGGCGGCCACTACGACGCTGAGGTCAAGACC
ACCTACAAGGCCAAGAAGCCCGTGCAGCTGCCCGGCGCCTACAACGTCAACATCAAGTTGGACAT
CACCTCCCACAACGAGGACTACACCATCGTGGAACAGTACGAACGCGCCGAGGGCCGCCACTCCA
CCGGCGGCATGGACGAGCTGTACAAGTAACTCGAGTCTAGAGGGCCCGTTTAAACCCGCTGATCAG
CCTCGACTGTGCCTTCTAGTTGCCAGCCATCTGTTGTTTGCCCCTCCCCCGTGCCTTCCTTGACCC
TGGAAGGTGCCACTCCCACTGTCCTTTCCTAATAAAATGAGGAAATTGCATCGCATTGTCTGAGTAG
GTGTCATTCTATTCTGGGGGGTGGGGTGGGGCAGGACAGCAAGGGGGAGGATTGGGAAGACAATA
GCAGGCATGCTGGGGATGCGGTGGGCTCTATGGCTTCTGAGGCGGAAAGAACCAGCTGGGGCTCT
AGGGGGTATCCCCACGCGCCCTGTAGCGGCGCATTAAGCGCGGCGGGTGTGGTGGTTACGCGCA
GCGTGACCGCTACACTTGCCAGCGCCCTAGCGCCCGCTCCTTTCGCTTTCTTCCCTTCCTTTCTCG
CCACGTTCGCCGGCTTTCCCCGTCAAGCTCTAAATCGGGGGCTCCCTTTAGGGTTCCGATTTAGTG
CTTTACGGCACCTCGACCCCAAAAAACTTGATTAGGGTGATGGTTCACATCGAATTCCTGCAGCCCG
GGGGATCCATTAACTAATAAAAAAAAATAATAAAGCATCACTTACTTAAAATCAGTTAGCAAATTTCTGT
CCAGTTTATTCAGCAGCACCTCCTTGCCCTCCTCCCAGCTCTGGTATTGCAGCTTCCTCCTGGCTGC
AAACTTTCTCCACAATCTAAATGGAATGTCAGTTTCCTCCTGTTCCTGTCCATCCGCACCCACTATCT
TCATGTTGTTGCAGATGAAGCGCGCAAGACCGTCTGAAGATACCTTCAACCCCGTGTATCCATATGA
CACGGAAACCGGTCCTCCAACTGTGCCTTTTCTTACTCCTCCCTTTGTATCCCCCAATGGGTTTCAA
GAGAGTCCCCCTGGAGTTCTTACTTTAAAATGTTTAACCCCACTAACAACCACAGGCGGATCTCTAC
AGCTAAAAGTGGGAGGGGGACTTACAGTGGATGACACTGATGGTACCTTACAAGAAAACATACGTG
CTACAGCACCCATTACTAAAAATAATCACTCTGTAGAACTATCCATTGGAAATGGATTAGAAACTCAAA
ACAATAAACTATGTGCCAAATTGGGAAATGGGTTAAAATTTAACAACGGTGACATTTGTATAAAGGATA
GTATTAACACCTTATGGACTGGAATAAACCCTCCACCTAACTGTCAAATTGTGGAAAACACTAATACAA
ATGATGGCAAACTTACTTTAGTATTAGTAAAAAACGGAGGGCTTGTTAATGGCTACGTGTCTCTAGTT
GGTGTATCAGACACTGTGAACCAAATGTTCACACAAAAGACAGCAAACATCCAATTAAGATTATATTTT
GACTCTTCTGGAAATCTATTAACTGATGAATCAGACTTAAAAATTCCACTTAAAAATAAATCTTCTACAG
CGACCAGTGAAACTGTAGCCAGCAGCAAAGCCTTTATGCCAAGTACTACAGCTTATCCCTTCAACAC
CACTACTAGGGATAGTGAAAACTACATTCATGGAATATGTTACTACATGACTAGTTATGATAGAAGTCTA
TTTCCCTTGAACATTTCTATAATGCTAAACAGCCGTATGATTTCTTCCAATGTTGCCTATGCCATACAAT
TTGAATGGAATCTAAATGCAAGTGAATCTCCAGAAAGCAACATAGCTACGCTGACCACATCCCCCTTT
TTCTTTTCTTACATTACAGAAGACGACAACTAAAATGCCCAAGAATAAAGAATCGTTTGTGTTATGTTT
CAACGTGTTTATTTTTCAATTGCAGAAAATTTCAAGTCATTTTTCATTCAGTAGTATAGCCCCACCACC
ACATAGCTTATACAGATCACCGTACCTTAATCAAACTCACAGAACCCTAGTATTCAACCTGCCACCTC
CCTCCCAACACACAGAGTACACAGTCCTTTCTCCCCGGCTGGCCTTAAAAAGCATCATATCATGGGT
AACAGACATATTCTTAGGTGTTATATTCCACACGGTTTCCTGTCGAGCCAAACGCTCATCAGTGATATT
AATAAACTCCCCGGGCAGCTCACTTAAGTTCATGTCGCTGTCCAGCTGCTGAGCCACAGGCTGCTG

TCCAACTTGCGGTTGCTTAACGGGCGGCGAAGGAGAAGTCCACGCCTACATGGGGGTAGAGTCATA
ATCGTGCATCAGGATAGGGCGGTGGTGCTGCAGCAGCGCGCGAATAAACTGCTGCCGCCGCCGCT
CCGTCCTGCAGGAATACAACATGGCAGTGGTCTCCTCAGCGATGATTCGCACCGCCCGCAGCATAA
GGCGCCTTGTCCTCCGGGCACAGCAGCGCACCCTGATCTCACTTAAATCAGCACAGTAACTGCAGC
ACAGCACCACAATATTGTTCAAAATCCCACAGTGCAAGGCGCTGTATCCAAAGCTCATGGCGGGGA
CCACAGAACCCACGTGGCCATCATACCACAAGCGCAGGTAGATTAAGTGGCGACCCCTCATAAACA
CGCTGGACATAAACATTACCTCTTTTGGCATGTTGTAATTCACCACCTCCCGGTACCATATAAACCTCT
GATTAAACATGGCGCCATCCACCACCATCCTAAACCAGCTGGCCAAAACCTGCCCGCCGGCTATACA
CTGCAGGGAACCGGGACTGGAACAATGACAGTGGAGAGCCCAGGACTCGTAACCATGGATCATCAT
GCTCGTCATGATATCAATGTTGGCACAACACAGGCACACGTGCATACACTTCCTCAGGATTACAAGC
TCCTCCCGCGTTAGAACCATATCCCAGGGAACAACCCATTCCTGAATCAGCGTAAATCCCACACTGC
AGGGAAGACCTCGCACGTAACTCACGTTGTGCATTGTCAAAGTGTTACATTCGGGCAGCAGCGGAT
GATCCTCCAGTATGGTAGCGCGGGTTTCTGTCTCAAAAGGAGGTAGACGATCCCTACTGTACGGAG
TGCGCCGAGACAACCGAGATCGTGTTGGTCGTAGTGTCATGCCAAATGGAACGCCGGACGTAGTCA
TATTTCCTGAAGCAAAACCAGGTGCGGGCGTGACAAACAGATCTGCGTCTCCGGTCTCGCCGCTTA
GATCGCTCTGTGTAGTAGTTGTAGTATATCCACTCTCTCAAAGCATCCAGGCGCCCCCTGGCTTCGG
GTTCTATGTAAACTCCTTCATGCGCCGCTGCCCTGATAACATCCACCACCGCAGAATAAGCCACACC
CAGCCAACCTACACATTCGTTCTGCGAGTCACACACGGGAGGAGCGGGAAGAGCTGGAAGAACCA
TGTTTTTTTTTTATTCCAAAAGATTATCCAAAACCTCAAAATGAAGATCTATTAAGTGAACGCGCTCCC
CTCCGGTGGCGTGGTCAAACTCTACAGCCAAAGAACAGATAATGGCATTTGTAAGATGTTGCACAAT
GGCTTCCAAAAGGCAAACGGCCCTCACGTCCAAGTGGACGTAAAGGCTAAACCCTTCAGGGTGAAT
CTCCTCTATAAACATTCCAGCACCTTCAACCATGCCCAAATAATTCTCATCTCGCCACCTTCTCAATAT
ATCTCTAAGCAAATCCCGAATATTAAGTCCGGCCATTGTAAAAATCTGCTCCAGAGCGCCCTCCACCT
TCAGCCTCAAGCAGCGAATCATGATTGCAAAAATTCAGGTTCCTCACAGACCTGTATAAGATTCAAAA
GCGGAACATTAACAAAAATACCGCGATCCCGTAGGTCCCTTCGCAGGGCCAGCTGAACATAATCGT
GCAGGTCTGCACGGACCAGCGCGGCCACTTCCCCGCCAGGAACCATGACAAAAGAACCCACACTG
ATTATGACACGCATACTCGGAGCTATGCTAACCAGCGTAGCCCCGATGTAAGCTTGTTGCATGGGCG
GCGATATAAAATGCAAGGTGCTGCTCAAAAAATCAGGCAAAGCCTCGCGCAAAAAAGAAAGCACATC
GTAGTCATGCTCATGCAGATAAAGGCAGGTAAGCTCCGGAACCACCACAGAAAAAGACACCATTTTT
CTCTCAAACATGTCTGCGGGTTTCTGCATAAACACAAAATAAAATAACAAAAAAACATTTAAACATTAG
AAGCCTGTCTTACAACAGGAAAAACAACCCTTATAAGCATAAGACGGACTACGGCCATGCCGGCGTG
ACCGTAAAAAAACTGGTCACCGTGATTAAAAAGCACCACCGACAGCTCCTCGGTCATGTCCGGAGT
CATAATGTAAGACTCGGTAAACACATCAGGTTGATTCACATCGGTCAGTGCTAAAAAGCGACCGAAAT
AGCCCGGGGGAATACATACCCGCAGGCGTAGAGACAACATTACAGCCCCCATAGGAGGTATAACAA
AATTAATAGGAGAGAAAAACACATAAACACCTGAAAAACCCTCCTGCCTAGGCAAAATAGCACCCTC
CCGCTCCAGAACAACATACAGCGCTTCCACAGCGGCAGCCATAACAGTCAGCCTTACCAGTAAAAA
AGAAAACCTATTAAAAAAACACCACTCGACACGGCACCAGCTCAATCAGTCACAGTGTAAAAAAGGG
CCAAGTGCAGAGCGAGTATATATAGGACTAAAAAATGACGTAACGGTTAAAGTCCACAAAAAACACCC
AGAAAACCGCACGCGAACCTACGCCCAGAAACGAAAGCCAAAAAACCCACAACTTCCTCAAATCGT
CACTTCCGTTTTCCCACGTTACGTCACTTCCCATTTTAAGAAAACTACAATTCCCAACACATACAAGTT
ACTCCGCCCTAAAACCTACGTCACCCGCCCCGTTCCCACGCCCCGCGCCACGTCACAAACTCCAC
CCCCTCATTATCATATTGGCTTCAATCCAAAATAAGGTATATTATTGATGATGTTACATCGTTAATTAACG
ATTTCGAACCCGGGGTACCGAATTCCTCGAGTCTAGAGGAGCATGCGACGTCGCAATTCGCCCTATA
GTGAGTCGTATTACAATTCACTGGCCGTCGTTTTACAACGTCGTGACTGGGAAAACCCTGGCGTTAC

CCAACTTAATCGCCTTGCAGCACATCCCCCTTTCGCCAGCTGGCGTAATAGCGAAGAGGCCCGCAC
CGATCGCCCTTCCCAACAGTTGCGCAGCCTGAATGGCGAATGGAAATTGTAAGCGTTAATATTTTGTT
AAAATTCGCGTTAAATTTTTGTTAAATCAGCTCATTTTTTAACCAATAGGCCGAAATCGGCAAAATCCC
TTATAAATCAAAAGAATAGACCGAGATAGGGTTGAGTGTTGTTCCAGTTTGGAACAAGAGTCCACTAT
TAAAGAACGTGGACTCCAACGTCAAAGGGCGAAAAACCGTCTATCAGGGCGATGGCCCACTACGTG
AACCATCACCCTAATCAAGTTTTTTGGGGTCGAGGTGCCGTAAAGCACTAAATCGGAACCCTAAAGG
GAGCCCCCGATTTAGAGCTTGACGGGGAAAGCCGGCGAACGTGGCGAGAAAGGAAGGGAAGAAA
GCGAAAGGAGCGGGCGCTAGGGCGCTGGCAAGTGTAGCGGTCACGCTGCGCGTAACCACCACAC
CCGCCGCGCTTAATGCGCCGCTACAGGGCGCGTCCTGATGCGGTATTTTCTCCTTACGCATCTGTG
CGGTATTTCACACCGCATACAGGTGGCACTTTTCGGGGAAATGTGCGCGGAACCCCTATTTGTTTAT
TTTTCTAAATACATTCAAATATGTATCCGCTCATGAGACAATAACCCTGATAAATGCTTCAATAATATTGA
AAAAGGAAGAGTATGAGTATTCAACATTTCCGTGTCGCCCTTATTCCCTTTTTTGCGGCATTTTGCCT
TCCTGTTTTTGCTCACCCAGAAA

P21

>AP001582.4 Homo sapiens genomic DNA, chromosome 11q, clone:RP11-215H18, complete sequence

CTGCCTAAGTCATTACAGCTAATTAGCCGTTAGCTCTTGTGTGGAAAAAATGGCAGAGGAGCATGAA
GCAAGTGCTGAAGATGCCTCAGAATTGTCAAGAAATGTTATAAATAATCTCTTTCTCTGGTAAATAAGG
GTAACTTTTTCGACCACATTAGCTTTTTTTTTTTTTTTTTTTTTTTGAGACGAAGTCTCGCTCTGTCAC
CCAGGTTGAAGTGCAGTGGCACAATCTCAGCTCTCTGCAACCTCTGCCTCCCGAGTCCAAGCAATT
CTCCTGCCTCAGCCTCCTGCCTCCCAAGTAGCTGGGATTACAGGCACCTGCCGCCATGCCTGGCTA
ATTTTCGTGTTTTAGTAGAGATGGGGTTTCACCATGTTGGCCAGGCTGGTCTTGAACTCCTGACCTC
AGGTGATCCACCTGCCTCAGCCTCCCAGGGTGCTGGGATTGCAGGCATGAGCCACTGCACCTGGC
CCACATTAGCTTTTGTTGATATGATCCTGTGAGGAGTCTTGCCTCAATAAACCAAAGGTAGACCAAAT
GTTCTTTTTTTCTTCCTTTATACCCCTACTCCCGCTATTTCTAAGGTCTCTGTCCTTATAGCAGTTCAA
GATCCTTATGAACAAAACATTCATACCCTTTGTCTCTTCCCACCCTTTTCAGTGTTGTATTTAATATTTC
TATGTCATGGCCTTAAAATTTCAGAAAGGGGAGAAAGCCTTTCAGGGAGCCCTGTTTATTCTGTTGT
GCCACTGAGTTCACTTGGTCTTGAATGAGGAAATGTAATCAGAAGAGTGTGAATGCAGGCTCAGCC
AGCTCTACTAAACACCTTTGAGGTATCTATATTTCATATTACACAGTAGACCTGACCGTTGGAATTCGT
GTTTCTTCAGTATTATTTTAAGAAATCAGGTAAACAGTGAAGGTTAGATATAGCATGAAGCAAAGACCA
TTTAATTTTTAGTGGAAGAAATATCAGTATTTTTAAGCCTAGAAATAAATAGATTGCTTAATCTATGTTAA
AAACCCACAACATACAGCTGAACTGCTTCCAGTCCTGATTTGCCAGAGTCCAAATCTAAGATATGGG
CTTTTACTTAATTTATATGTAGCTGAGAAATATCCATCCCCAGCCCTAAGTCATAGAAGTCCCTTGTCC
TCACCCCAGTGAATGAGAACTAGACTGCCAGCTTAATTCTTGGGAAATCCATAATTCTTGTCTTATTAT
TTTTCAGTGCCACATTGTTATTACAATTAGTAGGTGAGTTCTCTCTTTGACTTCAATAATTCTTTCTACT
TAACCTGTTTTCTTAAAGGTATTGCAAATAAAAGTATATCAACAGATTAATGCATTCCAGCTGAACGTTT
TAGCCTTTTTAGTTTCAGCTATCCACAAATCAAGCCACCGAGTCCAGCTGGTAGGTTTGTAACCATTT
TGCCAAAAATGTCTTGAGTTTGGGGACAGGAATTGGAAAATCTTAACCCAAAGTATTTTAAAATGTATT
GTATTTGAGAAATGGGAAAAGAAACAACCACAGGTATATTACAGCTATTGTGTCATCTGCCCTCTTGT
GTCTAATACCTGTCCTTTGAACAGCCTCTCCCACTAGGTCTGGATGGAGGATACCTTAAAGTGAAAT
GACAGACCAGGAGAATAACAACAACATCTCAAGTAACCCCTTTGCTGCTCTTTTTGGCTCCCTGGCT
GATGCCAAACAGTTTGCGGCAATCCAAAAAGAGCAGCTGAAGCAACAATCTGGTAAGTGGAGGAGC
CAATAGCAGCAAATAAGATGACCTAGAATGGGGTCTTCCCAAATTAAGTATTCTTTGAGTCCTTAATTG
CAATTGGAATTTAAGTTTGGAACAAAAGTTGGTTTTTGTTTGTTTGTTGAGAGCAGAAATTGGGGTTT
TTTGTTTGTTTGCTTTTATTGTTGGTTTTGTTGTGAGTCCCTCCCCTCCGCTGTCCTCTCCACTGTAC
AAAAAGAGACCAGGTAACAGGTGGTTCCATAGTAACAGATGCCAGTCGTGATCCCAAAGCTGTCAG
GCTGTTTTCTTTAAAAGAAAAAAAAAATCACCTAGTAAAAGCTATTTAAATAAATGTTTTTAACATGGGA
CACAGAGGTGGTAAGTGTAGGTGTAGAAAGCAAGTCATGTGCAGCTATAGAAAGCAGATGTAAAAGG
AAGAACTATAAAAGCTTTTTGACCTGTAGGCAAATGAGACTTGTTCTTAAGCTCAGCACTTCAAAGGC
CCTCCCAAGCAAAGTGCTTCTTCTGTTCCGATGAGTGGCATCATTTGTAGCTTAGTCCTGTTAGGAC
AGACTTTTTTAAAAAATCTGATCCTTAGCCTTACTATGAAAAACATCAGAATAACATCTCAGTAATCAGA
AAAAATGGTTTTCTTTCAGGCACCGAATTTAAGATCAAGTCCAGCTTCCTCCTATAAGGAGACATATAC
ACTAGAAGAATTAGGATTTTTCTCAAATATGTTAGTCTTCTGAAGATATTCACAATCTTTCAAACCAGAA
ATTGCTCCCCTTCTCCTCTCTCCGAAAAGAATAATAGTGTTTTTCTTTTGAGAGAAGGAGAAGTGTTT
GCATTTTCCCTTTATTTTGTTTATTTAAAAAAAAAAAAAACACTCAGGTGCATAGCTGACTTTTGGCCT
TTCAGTTAGAAAGGGCTGAAGAAAAAAGAGGTAAAGGCAAGACAGATAAGTGATTGATGCAAGGGA
AAAACAAAATCAATGTTGATTTTTTTTAACATGGTATTTTCAGTCTCTGAAATCTCTCCCAGATTTCTAT
GATACGGGGCCAGAATGGGAAACTCTGGTTTAGAAGATTGTAGGGGAAAAAGCAGAGTGAAAAATA

GGAAAAATAAGGAATGAGAAAGTATGATTTATGAGTATGTGGTTGAGGGAAATTTTGCTAAACAGATT
CTTAGCTTCTGTCACCATTGGAGGTCAGAGAAAAACTCAACAGTATAGCAGCTGAGCAACAGAACGC
TTGTCCTTCTGGCTATAAACAGACATAAATCTGGCTAGAGTTGGGAGCATTGCAGGCACATGCCCTT
CTGTTACTATTTTGGCACCTCATCATCCTAGTTTGTTTACACTGAGTTTATCATGCCACCAGGTCCAAC
CTGTTTCCACTAGCAATTAGTACTTTGCCAAAACCTTTGGCTTCCTATGTTGAGCTTATACTCACCACA
CTGAAACAACTCTTACTTTGAGAGTTTTACCTGGACTTTCAGACTAGCCATTCTGTCAATAGAGAAAA
GTACATATAGACTCATATGTGGCATATAGTGAATAACACTACTTTTATGTAGTCATTCAACATGAATGAT
GGGGCTGGGCGCAGTGGCTCGCGCCTGTAATCCCAGCTCCTTGGGAGGCCTAGGTGGGCAGATC
ACCGGAGGTCAGGAGTTCGAGACCAGCCTGGCCAACATGACAAAACCCCGTCTCTACTAAAAACAC
AAAAATTAGTCGGGCATGATGACAGGCGCCTATAATCCCAGCACTCTGGGAGGCCAAGGTGGGCGG
ATCACCTGAGGTCAGGAGTTTGAGACCAGCCTGGCCAACATGGTGAAACCCCATCTCTACTAAAAAT
ACAAAAATTAGCCGGGCGTGGTGGTGGGCGCCTGTAGTCCCAGCTACTTGGCTGAGGCAGGAGAA
TCACTTGAACCTGGGAGGTAGAGGTTGCGGTGAGCCGAGATCACACCACTGCACTCCAGCCTGGG
CGACAGAGTGAGAGTCCATCTCAAAAAAAAAAAATCAAAGGATGGGAGCAGATTTTGAGGACAGAAG
TTTAAAAAAAAAAAAAATCGAAGGAATTTCATCAATAAGAAAAAAAATAACAAGAAAGAATAAAAAGGTAC
AAAAGATGTGACATGCAACTTATAACCAAATGATTAAAAAATTTATTCTCATCAATCATTAGAGAAGTGC
CAATTAAAACAACCATTAAATTGTTTTTTTACCTATGAAATTTGTGAAGATTCAAACAGGCACTCTCTCA
TATACTGTTGGGTTTGCATGTAAATCAGACAGTCTCTTTGAGGGTAACTTGGTAGTAAATGTTAACTTT
TTTTTTTTTTTTTTTTTGAGACGAAGTCTTGCTCTGTCACCGAGGCTGGAGTGCAGTGCAACCTCTG
CCTCCCGGGTTTAAGCAATTCTCCTGCCTCAGCCTCCCGAGTAGCTGGGACTACAGGTGTGTGCCA
CCACGCCTGGCTAATTTTTTGTATTAGTAGGGATGGGGTTTCACCATGTTAGCCAGGATGGTCTCAAT
CTCCTGACCTCGTGATCCGCCTGCCTCAGCCTCCCAAAGTGCTGGGATTACAGGTGTGAGCCACC
GTGCCAGGCAGTAAATGTTAACTTTTTAAATGTTCAAGCTCTTTGATCCAGCAGTTCTGCTACTGTGA
ATTTATCCTATAGAAATACTCATACAAGTAAACAAAAAGAGTTAAGTTGTATTGTATTCAGTGCAGTATTT
AAAATCATGAAATATTAGAAACAACCCTATTGTTCATCAGTTGAAAAGTAGCTACATAAACCACAGGGC
ATGCAAAAGATGTGATACAATACAGCTATTAAAAATAGTGAGGTAGGCCTCTAAATGGCCTAACAGGG
AAAGGTGTTGGTGCTGATTAAAAGGCAAGCTGAAAAGAGCTATGTATAGTATCCCATTTTTTAAAAAAA
TTAAAAAGATAGTAATGAATATTAGCACCAGAAAAGAAAAAAATTCTTGAACACTGTTAAGTGTCTTTA
GGGAATTGGTTCAGGAGACTTTCTATACTATATTTTTCTGTACAGTTAAGAACTATTAAAACAAATGTTA
TCTCTTAAAATCAGGGAGAAAAGCCATACATTTAAAATAATTAAAGGCATTTTGATCTATTAGGTAATTC
TTTGTTCACTAGGATTCTGTAGAGAAGATAACTTTGCTATTTCTGGCAGTGTGGAGTATAATTATGGAT
AGAAATAAAACTTTTGCTTTGTAAAATGTGGTAGGATTTATAGCCAGGAATTTCAAAACCCTATATTTAA
CTTCCCTGACTTGAGATGTTCACTAATTAGATGATTAGCTTGTTTTACATCTGAAATTATCATCTTCTTG
GTTACATTTATTATTTTTGGTAGCATGCTTGGTTCTCTCTTAAAGGGGTGTAACATTTAACAGTATACAT
TTTCTGGCAGATGAACTCCCAGCTAGCCCAGATGACTCGGATAATAGCGTGTCAGAGAGCCTGGAT
GAATTCGATTACTCTGTGGCTGAGATTAGCCGCTCATTCCGATCACAGCAGGAAATATGTGAGCAAC
TCAACATCAATCACATGATCCAAAGGATCTTCCTTATTACTCTGGACAACAGTGAGTTACAAACTTATA
GCATTATTCTACCCTTCCTATTTGAGCTTGGGCTAGGGGCTTGCTCTTTGAAACTGGTTACAGCTAAT
GGTTAGATACAAACCTACTCACTGCACTCCCTGGAACTCAGGGTTTATATTTCAGTGAACCTTTTGGT
ATGACCTAAAAACAGTATTTAGATGCAGATCTAACTTTTGTTCCCATTTCAATACTGACTGCACTATATA
GCTACTTGCCTTTACTACCTTAAAAAGGAAATGGGAAGGGTAGCCAAACCCATTTTCTTCTCAGAGGT
AACCTAAATATTCAGATGGAAGCCTTCAGAGGCATCAGAGAGTGTTTGCCCTCTGTGCCCTGTCTGT
GTTTAAACCCCAAGGTTGGTACTAATGTCTGGGATTTACTCTTTAAATGTCATGATGCTAGGTTTATAA
CAAATCCCATTCTCTGTGACTAGAGGGGGAAAAAAAAAGATTTGAATCTTTAGATGTTGCTTTGTGTA

TCAGGATCATTCTGGTATTACTTGAGTCAGATTTGCAGTTCAAGAGTGATGCTGTCAAGATCAGTTGT
CTGTATAAAACAGAAGCATTATCCTTAACCTATCATATTAAAACCATTTCCCAGGTGATCCCAGCTTGA
AAAGCGGGAATGGCATCCCTAGCCGTTGTGTGTATTTGGAAGAAATGGCAGTAGAGCTAGAAGATCA
AGACTGGCTTGATATGAGCAATGTTGAGCAGGTAATATTCTTACGTTCCTAATGTGTGCCCTTAGCAA
AAATTAGCTATGCGGCTGAATTCTGCTCACTGTTCTCCAACTTATGCTTGTGTCCATATGTCCATCCCT
CTCTCTTTTTTTTTTTTTTTTTTTACCAGTAGAAGGCCAAGTAACCTGGATGTGCATCTCTAACATTGT
GGTGTTCCTCTTTTGGAATCTTTTCATCTACCATGGCCACATGCCCAAATCTACCCATCCTCTAAGAA
CCAGTTAAAATGCCTTCTTAATTACAAAGCCTTGGCCAGGCACATTGACTCACGCCTGTAATCCCAG
CACTTTGGGAGGCTGAGGCAGGTGGATCACGAGGTCAGGAGATCAAGACCATCTGGCTAACATGG
TGAAACGCCGTCTCTACTAAAAATACAAAAAATTAGCTGGGCATGGTGGTGGGCGCCTGTAGTCCCA
ACTACTTGGGAGGCTGAGGCAGGAGAATGGTTTAAACCCAGGAGGCGGAGGTTGCAATGAGCCGA
GATCATACCACTGCACTCCAGCCTGGACAACAGAGCAAGACTCCGTCTCAAAAAAAAAAAAGAAAAA
ATTACAAAGCCTTGCCAAATAAAAGTTGTTACTCCCTCCTCTGGGTACCTAGAATAATCAGCCTCTGA
ATCTCTTGTGGGAATTGTCACTTTCTGTCTCATGTTGCGGCTTTCTTATGTACCTGTCTTATCTCCCCT
ATTAGACTTAAGTTCTTAAGGGCAAGGCCCATCGCTTTTTCTTCTTTATATCCTTTCTATCTTAGTCTGT
TTTGTGTTGCTATAAAAGAATACCTGTGACTAGGCACAGTGGCTCACATCTGTAATCCCAGCACTTTG
GGAGGCCAAGTTGGGAGGATCTCCAGGCCAGGAGTTACCAGCCCTGGCAACATAACAAGACCATG
TTTCTACAAAAAATACAAAGAATTAACTGGGTGTGATGGCATGTACCTGAAGTCCTAGCTACTCAGGA
GGCTGAGGTGGGAGGATTGCTTGAGCCTGGGAGGCAGAGGCTGCAGTGAGCCATAGTCACACGA
CTGCACTCCAGCCTGGGCGACAGAGCAAGAAAGACCCATCTAAAAAAAAAAAAAAGAACACCCAAG
CAGTAATTTATAAAGAAAAGTTTATTTAGCTCACAGTTCTGCAGGCTGAGAAGATTCAGGGTATGTCC
TTGGCCTCTGGTGAGGGCTTTTGTGCTGCCTCACAATGTGGTGGAGGAAGTCACCAAAGGACACAT
ACCATGAGGACGCATGCAAGGAGGAGAAAACCTGAGAGGTGTCCTGGCTTAATAAGAACCCGCTCT
GGAGGGAATTTGTCCATTCCTATGAGAATTAATCCAGTCTTGTGAGAGCAAGAACTCATTCAGCACCT
TGAGAAAAGCACCAAGCAATTCAGGAGGGATCCGCCTCCATACCCAGACACCTCCCACTAGCCCCA
CCTCCAACAGTGCCACATGGAGATCAAATTTCAACATGAATTTTGTTGGAGACAAACATATAAACCAC
AGCACTTTAAAAATCAAAACAGTGCGTTGTACCTTGCAGGTGCCTGAGTAACATTAAAAGAATAAAAG
TTTGGTAGGAAAGACAGTTGACAGCTTGCAGTTTAACTTCACATTTGGAGTATCTTTATATAAGTCTGA
ATGAGATCAGCAATCACACAGAATAGTAGATTTCACATCTAAGTTTAATCACTGATCTCAGCACCATTC
AACAAATAGTCTTTTAATGCCTACTCTATGCTGGGGCTATACTAGACACTTGAGGCTTGATTTAGGATA
ACTTTATATATGCCATAATTTTCTAAAATTTTCTAAAGTAAAAATATACAGGGCCCTTTTTTTCTCCTTCA
CTCTACATTCAAATCTGTGATTCTTGCTAATAACCTGTGTCAGAATTACCTTTTACATCTTAATGTTCTC
AGATTAATTTTGCAATAGTTTAGTATTTTTGTTTGGTTTTCTTTATAGGCCCTCTTCGCTCGCTTATTACT
TCAAGATCCAGGCAACCACTTAATTAACATGACTTCTTCTACAACGCTAAATCTCTCTGCTGATCGAG
ATGCAGGAGAGAGGCACATTTTTTGTTACCTTTACTCCTGCTTCCAGAGAGCCAAGGAAGAGGTAAA
GGAATAATCCCCATTGAATACTGTATTGTCCTCTTGGGTATGACTTCAGCATGTGGGCCAGAGTTGGA
TCAGTCTTTATTTCAATTTATATATGTCATAGTATGTCTAGAAGTGAATATCAGGGACACTAAAGCTTTTA
TATCAATTTATATATGTCATAGTATGTTTAGAACACTGGATATCAGATGCACTAAAGCTTTAGTCTCCTTT
CCGTATCTTGGGGTCCACCCTTCTCCCCAGTTAACCAATGCCTCCTATTCTCTCTAAAACCTCTTTAG
AAATCACATCCTTAAAAAAATCTCATGAGGCTGGGCACAGTGGCTCACTCCTGTAATCCCAGCACTT
TGGGAGGCCGATGCAGGCGGATCACATGAGGCCAGGAGTTCGAGACCAGCCTGGCCAACATGGC
AAAACCTTTACTGAAAATACAAAAATTAGCCGGGCATGGTGGCACACATTTATAATCCCAGCTACTCA
GGAGGCTGAGGCAGGAGAATCACTTGAACCTGGGAGGTGGAGGTTGCAGTGAGCTGAGATCACAC
CACTGCACTCCAGCCTGGGTGACAGAGCGAGACTCTGTCTCAAAAATCAGAAAAGGGCTCATGAAT

TCATTTCCACTGAGATGTAAATATTAGAGTAGGCTAAGCCGGATATTTCCAGGTTTGGAATTTTAAATA
GGCAGATAGATGATTCCTTAAGCCAAGGAAGGAATCTCTAATTAATTTTCAAGCATCCACAGGAGACC
AGTATGTCTCTTCTATTGTATTCACTATGTCTACTTTCGTTCCAGATTACAGAGTTAGACTATTCCCTCT
TTTCTTCATGCTGTTTGCAGATTACCAAAGTTCCAGAGAACCTGCTACCCTTTGCAGTGCAGTGCAG
AAACCTCACTGTGTCCAATACCCGAACAGTTCTTCTCACCCCAGAGATCTATGTTGACCAAAACATC
CATGAGCAACTGGTAGATTTGATGTTAGAAGCCATCCAGGGAGCCCGTGAGTACATGAACAAGATCT
GTAAGCTTCTACATTTTGATTTTCAGATTTAAAATTTCACTAATTGCATTAGAAAGTCTCAGAGCCATTA
TGGTTCTGGTATAAATTAAGGAGGAGGGCTCACATCTTGATCTTTGAGGTTGAGTACCAATAGAATAA
AGCTGGGCATGGTAGCACACCCCTGTAGTCCCATCTACTCTGGAGGCTGAGGCAGAAGGATCCCTT
GAGCCCAGGAGTTTGAGGCTGTAGTGTGCTATAATCACACCTGTGAATAGCCACTACACTCCAGGCT
GAGCAAAAAAACAAGACCCCCAGCTCTAAAAAAAAAAAAAAAAGAATTATTGAAAGTAAAGAGCTAGT
GTGTAAAGGCATATAAAAATTCTTGTGCCCTCCTTTCTCTCTCATTTGTCTTCTGTTAGATTTTGAAGA
TGTAACTGAGTTTCTGGAAGAGGTCATTGAAGCCTTGATATTGGATGAGGAAGTTAGAACATTTCCAG
AAGTCATGATTCCAGTGTTTGATATTTTATTGGGCCGAATAAAAGATCTAGAGCTCTGTCAGATCCTTT
TGTATGCATATCTGGATATTCTTCTCTATTTCACTAGGCAAAAAGATATGGCAAAGGTAGGTCTGAAAG
ATGATATGTATTCAGTTGAGCTAGATGTGTAATACATGATGCTTCCCTATCCTGAAGACACTATAATACT
ACCTATAAAATAAGAAAAAGATAGCTTGGTTGTTGTATCAACTTAACTGTTTTGAGGCCCTCCCACCC
CATCACCTTATCTCACCATGAAGATGAATAAGCAGAACTTGTCTTCTCTTACAGGTTTTTGTAGAATAC
ATTCAGCCCAAGGACCCTACCAATGGGCAAATGTACCAGAAGACCTTGCTGGGAGTAATTCTGAGTA
TCTCCTGCTTATTAAAGACTCCGGGTGTTGTAGAAAATCATGGCTACTTTTTGAATCCATCTCGTTCC
AGCCCCCAGGAGATCAAAGTACAGGAGGCCAACATCCATCAGGTGGAACTGTTTACCAGGGTATCC
CAGCCCCCTGATCTTAAAAGAGTTTTCTCAGAAAGCAGATACAGATAAGGCTGCTCAAGCCTCTCTG
GGAAAGCAAATTGATCTAGTTGGCTTTTACATCACATAAATAAATTAAAACTGGGCATGGTAGCTCAAG
CCTGTAATCCCGGCACTTTGGGAGGCCAAGGTGGGAGGATCGCTTGAGCCCAGGAGTTCAATTTG
AAACCAGCCTAGGCAACACAGTGAGACCCCCATGTCTACAAAAAAATACAAAACTTAGGTGGGCATG
GTGGCAAGCATCTGTAGTCCCACCTACTCGGGAGGCTGAGGTGGGAGGATCACTTGAGCCTGGGA
GGTCAAGACTTCAGTAAGCCATGATTGCATCATCACGCTCCGGCCTGGATGACAGAAGAAGACTCT
GTCTCAAAAAAAAAAAAATCCCCTCTTTGTAGCATATGCTAAGAACAGCACGTGTGGGGTAAGAATGA
GCTGCTGCTACTCATTACCCTTTGCTGTAGCACTGCTACAGCTACTTCTCCCCAACCCTCATTTTTCT
CACTGAGCTATAACTTTAGCTCTGAAAATCTTTTGACTGAACTCTAGCTTATAAACCCAGTTTTTCTTC
CTTTTCGTGTCTCCACCTATGCCCTCTCTTTGGGCTTCTTCACATAAATTATAGTAGAGGTCTGGCCA
ACATGAGGCATTGAATGATTGAATTGGGGATAGTTTAAGTCCCCTAATCATTAATCATTACAACTCTCT
AAGGCAAGGCATGGCAAACTAGCAGTGGCCAAATCTGGCCTGCCACCTGTTTTTGTAAATAAAGGTT
TATTGAACATAGCCATGCGCATTAGTATATGTATTGTTTCTGGTTTGAATTCATACTACAGTGATAGAGT
TGAATAATTGCTACAGACACTGTATGGCAAGTAATGCCAGAAATCCTTATTGTCTAGCCCTTTGCAGA
AAAAATTTACAGACCTGTGCTCTAATACCGTGTAATGCCATCTGCAGTTGGGCAGCTGATCAGAAAAT
CTCTTATAAGTGCCCAGAAATATATTTTAAAATCAAATTAGGGGATGAGGCACAGAGAGTGAGGGGAA
AGATTAGGATTAGCATATTTTTGGAAACTGCCAGTTGCTAAGGTTTGGGAATAAAAACTATTAGGATTG
TGAAACGTTCTGTGGTTGTGTTTTCTGGTTGAACAGTTCATGGCTCAGTTCCACGAAAAGATCTACC
AGATGCTGAAGAACTTACTCCAGCTCTCTCCAGAAACCAAACACTGTATCTTGTCCTGGCTTGGAAA
CTGTTTGCATGCAAATGCAGGCCGCACCAAGATTTGGGCCAATCAGATGCCAGAAATCTTTTTCCAA
ATGTATGCCTCAGATGCTTTCTTTCTGAATCTGGGTGCTGCTCTCCTGAAGCTATGCCAGCCATTTTG
CAAACCCAGATCCTCTCGGCTCCTCACCTTTAATCCCACATACTGTGCCCTCAAGGAGTTGAATGAT
GAAGAACGAAAAATTAAAAATGTACACATGAGAGGTAGGAGAGAACCAGGCTTCTCAAAACTGTGTG

TGTGTGTGTGTGTAACGTGTAAAGCATTCACTCCCTTATCCTTTTTCACAGAGTTCCTTTCTCAAAAA
AAGATGAGGCAGAAACGAGCTATTTCGATTTTTTTTTCTTTTTTTTTGAGATGGAGTCTTACTCTGTCA
CTCAGGCTGTAGTGCAGTGACGCTATCTTGGCTCACCGCAATCTCTGCCTCCCGTGTTCCAGCGAT
TCTCCTGCCTCAGCCTCCCGAGTAGCTGGGACTACAGGCACGCGCCACCACGCCCAGCTAATTTTG
TGTATTTTTAATAGAGACGGAGTTTCACCATGTTGGCCAGGATAGTCTCGATCTCTTGACCTCATGAT
CCATCCGCCTCAGCCTCTCAAAGTGCTGGGATTACAGGCGTGAGCCACCACGCCCAGCCAGAATTT
TTAAAAGACAGATGTTTGAGAGAAAAAAAAAAACACCTGGTAGCTTTTAAATGTTGTACTTTAGTGAG
ACCTTCTCTCTGGATTTGGCAGCCTTGCATATTCATTCATTCATAGTTTATAAAAATGAATAAAGCAGA
GTCCTCATCCTTTTTGAACTCAGGCTCCAGAAGGCAGATATATTAACTGATAATTATAATATGGTATAGG
GCTAAGCTATGGGAGCACAGAGGAGGGAGTAAATAATCAATTCTGCAGAAGTTGGAGGTCTTCATAG
CTGAGTCTTGAGAGACAAGTTAGGTTTTACCAGGCAAAGAAGCAAGAGGGTCCCTTTATACAGAGA
GAAAAGCAAGTGTAAAGGTACAAAGCTATGGAAAATCTTGACTAAGGAATGGTGAGAGGCTCAATGT
GGCTAGGGCAAGGGGTAGATGGGGTTGAGTTGGGGAGGAAATGGATCAAGGGAGGAAGATAGGTG
GGGCTGGATTGGAGAAAACAGTACAGAGAAATGAAGAGAAAATAAATGACTCACAGTCCTACCATTC
AAAGTGAAATCACTCTTAATTTTGGTATATTTCATTCCAGTTTTATAATTCTAATTGTAAACATGATAACT
GCTCAAAAGCAAATAATCAAATTCAGAAAAGCACAAAATAAATACAAATGATCTATTGCCTATCACTTG
AAACAATCTATAGTAGTATTCCTTTTAATTTTTGGTGATTTCATGAAAAAGATTTAACTGTATTTCTGTAG
ACGGTGAGAAACCAAAAGATATATTCAGACAAAGAACTAATATGACAAGATATGTTTTTAAGAAGATAT
CACTGCTAACAGTATGGAAATTGAATGAGTGTAAGAGGAGACTGGAGACCAAGACATTTACCCTCTT
TTTTTTTTTTAACTTCTTTGAGGTTTGGACAAAGAAACCTGTTTGATCCCAGCTGTGCAGGAGCCGAA
GTTTCCACAGAACTACAACCTTGTAACAGAGAACCTTGCTCTGACAGAGTACACCTTGTACTTGGGA
TTTCACAGGTAACTCCTCTGATGTCATTAGGAAAAAACAGTTTAGTTGTGTTGATAACTAGAAGGTAG
AAATTGAGTCTTTGGCCAGGCACAGTAGCTCACATCTTTAATCTCAGGATTTTGGGAGGCCAAGGCA
GAAGGATTGCTTGAGGCCAGAAGTTTAAGACCACCAACCTGGGCAAGAAAGTGAGACCCCCATCTC
TACCAAAAAAAAAAAAATTAGCCAGGCATGGTGGCATATGCTTGTGGTCTTAGCTGCTTCAGAGGCTG
AGGCAGGAGGATTGCTCTAGCCGAAGAGTTGGAGGCCACAGTGAGCTATGATTGTGCAACTGTACT
CCATCCTGGGTGACAGAGTGATAACCTGTTTCTTAAAAAAAAAAGAGAGGAAGAGTCTTACAGATATA
ATCTGAGCTTATTCTAAAGGTTCACATTAGAATCTAATAGAAAAGGGGCATTGTATGAATTTACTTGTG
TTTCATGAATCAGCTTTGTCCAGATTCCTTAAATTTGTTTTTAACTTTTATCGTTGTCGTTGTTGTTGAA
ATGGATTTTCGCTCTTGTGAATTCCTGAGCTCAAGCAATCCACCTGCCTCGGCCTCCCAAAGTGCTA
GGATTAGAGGTGTGAGGCACCATGCCCAGTGGATCAATAGCATTTTAAAGCAAAAAAGTATTTTTAAT
TAAGGTATGCACATTGTTTTTTAGACATAATGCTATTACACATTTAGAAGACTACAGTATAGAGTAAACA
TAACTTTTATAGGCACTGGGAAACCAAAGAATTTGTGTGACTCACTTTATTGCAGTACTCACTTTATTG
TGGTGGTTTGGAACTGAACCTGCAATATCTCTGAAGTATGATTGTATAGGCTTCTTAGTTTTTCCATTA
AAATGAACAATTTAAAATTAAATACTCAGGCTCAGACTTCAATCTACTCTTTATTCAATCAACTCTCCAT
TAGTCAAACTTTATAGATAAATTAATTAATGGGAAAGCAAACAGAAATACCAGATTTTAGGTTAAACATG
ACAAAGTGAACACACATGTATTTCGTTCTTTTTATCTCCTAGAAAGCCCACTCAAATGGCATATACCCA
TAAAAGACAAGGACAATATAAGGATAATAGCAGACAAGAAATGCAATAAAAATTTGGAAGCTGAAAAA
CAAGTGAATGAGTGATAAATACAATAGAATGCCAGAAAAAAACTAAATTCTTCCAGAAAATAGAAGAAA
TAAAATGTTCCTCAACTCATTTATTGAGGCCAGTATAACACTGATACAAAAGAAAATAAGATAAGGACA
CTTAAGAAAAAAATTCCAAGCCAATTTCTCTCATGAAAATAAAGACAACAATTCTAAACAAAATATTAGC
AAATTGACTTCAAGTGGGTAAAAATTTTAAAAAGATAACACACCACTACCAAGTAGGACTTATTTCAGT
TATATAACAATGGTTTAATATCAGGAAATCTATAAATGTAATTCATCATATAAACAGCTTAAAAGAGAAAA
ATCATAGGAACCTGTCTATAGAACCGGAAAAAGTATTGGATAAAATCTCATGCCTATTCAGGATAAAAG

CTCTTTATAAATCAGTATTAGAAGGGAATTCCTTAGCCTGATAAAGGGTAACTATTAAACCCCCACAGC
AAATATTGTTCTAAGTAGAGACATATTAGATCATTCCCAATATTCAATATAATTCAATAATTAAAGAAGAAT
GCCCAGTATCTTCGCTGCTATTCAGTGTTATATTGGAGGCCCATCTGGTACAGTACAAGAAAAACAAA
TTAGAAGCTTACAGAACAAAAAAAGAGAAAATAAAGCTGTAATTGTTTGCAGATGACATGATTTTTTAC
ATTGACAAAGCAAAAGAATATAGACAAATGTTTAAGAATAATAGAATTTAGCAATGAAACTGAATATTAC
ACCAATATACATAATATTTAAAAACATTTCCATAAGGCAACAACAATCAACAACAAAAAAATAACTTTTTT
TTTTTTTAAACGGAGTCTCCCTGTGCCGGCCAAGGCTGGAGTGCAGTGGTGCAATCTCGGCTCACT
GCAACCTCCACCTCCTGGGTTCAAGTGATTCTCTTGCCTCAGCCTCCTGAGTAACTGGGATTACAG
GTGTAAGCCACCACACCCGGTTTTTTGTATTTTTAGTAGAGATGGGGTTTCCCCATGTTGGCCAGGC
TGGTCTTGAACTCCTGACCTCAAGTGATCCGCCTGCATCAGCCTCCCAAAGTGTTGGGATTACAGG
CGTCAGGCACCGCACCCAGCCAAAAATACATAACTTTTAAAAAGATACTACATACTATAGTATCAGAGT
ATATCTCATACCTAAGAACAAATCTAATAAAAAAAAAGTATGACCTACACAACGGAAATTATAAAACTTT
AGTGACATTTAAAAATACCTTCATAGACCAGGCATGGTGGCTCACATCTGTAATCCCAACACTTTGGG
AGGCCAAGGCGGGCGGATCACTTGCTCCTGGGAGTTTCAGACCAGCCTAGGCAACATAATGAAACT
TCTCCTCTTCAAAAAATACAAAAAATTAGCTGGGCATGGTGGAGTGCACCTGCAGTGTTAGCCACTT
TGGAGGCCGAAGTGGGAGGATCGCTTAAGCCCCAGAGGTTGAGGCTGCAGTGAGCTGAGATCGTG
CCACTGCACTCCAGCCTGGGCAACAGAGTGAGACCTTGTCTCAAAAAAATAGCAATAATAACAGAGA
GCTATGTCATGTTCATGTATTTTAAACAATATAATAATATCATAAAGATATAAATTCCCCCCAAATTGATCT
ATAAATTTGTTTCTTTTCCTATCAGAATCACAGCAGGGTTTTTCATGAAACTTAAGATTCTAAAATTTATA
TGGAGAAGTAAAAAGCCAAGAATAGCCAGAAAAAGGAAGAGCAGGGCAGGGAACTGACTCTATGCC
AGGATGTATTGTAAATCTATAATAGTTAAGATGATGTGGTATTGATGTAAGGATAGAAAAATTAACTGAT
GAGACAATATATAGCCTATAAGTCAGGCCTTTATGCATATGTTATGCATATAAAAATTTGGTATATGATGG
AGGTGGCATAGCAGATCAGTGGGGAAAGGAAAATATGTCCAAATTTGGAGCTTGGAAAAATGCCATA
GGGGAAAAAATGAAATTGAATCTCTAGTTCACATTGTACACAAAATCAACTCCAGATGGTTAAGGGGT
GGAGGAGGCAGGCAGGAGATTGTTGGTTTTCATTATAAACTTTACAATACTATTAGACTGTTATGTTAT
TTAGATTAAAATACAAACTACTTTTTAAGAAAGAAAAAAATTCCCAGGGTTAACAAAGATGTAATGAAA
GAGTAACATTTATACCCTGTGTGAATATAAATTGGTATGACTTTTTGGAAGGGTAATTTACAAATATCTG
TCTAAATTTTAAGTACATGTATCCTTTGGTTCAGCAATCTCTCTTCTAGTAGCTTATCCTGCAGCTATAG
TTGCTGAAGAGATATGTACACAGATGTTTCTTGCAGCTTTGTGGGTGTATTTGTCCATTCTCAAATAG
CTACAAAGAAATACCTGAGACTGGGTAATTTATTAAGAAAAGAGGTTTAATTGGCTCATGGTTCTGCA
AGCTGCACAGGAAGCATGATGCTGGCATCTGCTTGGCTTCTGGAGAGGCCTCAGGAAACTTACAAT
CATGGTAGAAGACAAAGGGCAAGCGGGCACATCATATGGCCAGAGCAGGAGAAAGAGAGAGCCAG
CGAGGTGCCACACTTTTTTTTTTTTTTTTTTTTTTTGAGACAGAGTCTCGCTGTCACCCAGGCTGGA
GTGCAATGGCACAATCTCGGCTCACTGCAACCTCCGCCTCCTGGGTTCAAGCAATTCTCATGCCTC
AGCCTCCCGAGTAGCTGGGATTACAGGCATGCACCACCATGCCCGCTAATTTTTGTATTTTTAGTAGA
GATGGGGTTTCACCATGTGGGCCAGGCTGGTCTCAAACTCAACTTCAGGTGATCTGCCCACCTCGG
CCTCCCACAGTGCTGGGATTACAGGTGTGAAACACCGCGCCTGGCTGAGGTGCCACCCTTTTAAAC
AGCCAGATCTTGAGAGAACTCACTCACTATCACGAGGACAGCACCAAGAAAATGGTGTGAAACCATT
CATGAGAAATCCACCCCCATGATCCATGACCCACCTCCAACATTGGGGATTACAATTCAACGTAGGAT
TTGGGCAGGGACACACATCCAAACTATATCAGAGGGGATAGGGGAATGTTTAATCGGTATACAGTTT
CAGTTTTGCAAGATAAAAAAAATCCTGGATATTGATTGCACAAAATTGTGAATGGACTTAACACTACTC
AACTGTATACTTAAAATGGTTAAGATGATAAATTTTGTTATTTGCATTTCACCACAATTTTTAAAAAAATA
ACAATCCTGAAAAAAAAAATGGGGCCACTGCTCGAGGACTGCAAGTTTGCTAGAACCTGAGAGGAG
ATAGGAATCATTTGGGAGGCCAGAGAGGAGAACATGATTAAGTAGAGTGCCAGAAATCAGAGCAGG

CTTCAAGCTGGTCCGGAGGCTCGTGGTGGAGGTTGAGTGAGGGCAAAGTTCAGAGAAGTAGGACA
CCTCATGCCAAGTATATTCGTGTCCATAATACTGATTTTCCAGTCAGTTGCTGCCTCCTTCTGTTCATT
TTGGGAGGGATCGCGATACTAGGTTCTGCAAACCGAGGTCGGAGGAAGGCTCTAAACAGTGATTCT
TAACTGAGATGGCACGGTTCCCCGGAAGATGTTTTGGAAATGTGTGGGACTGTTTTTGTGAGATGCT
AGAGGCATTTAGTGGTTGGGGGCTAGAAATGCTGGCATCCTGCAATGCATGAGAACAGTTCAGCAA
AACAAAGAATTGTCCTACATCCCACACGACTTTCAAATGGCACATGGGACATTAATGAAGCTGGAAAA
CCTAAATGTGTAATATCTGATCCTAGACCGTAGCTCCATTTTACACAATATGTTTTGCATGGTTTTAATAT
ATGTTGAATTTTCCAGGCATGCAACTACCATATAATTTAAAGGAAGATTGTTTTGTTCAGAGCTTTATC
AATCGTTGTTTACTGCTTTTGAAAATTACATCACTAAAGCACCCGTGGTATTTGAGTCACCGGCACCA
CCCAATAGCATACATTTGAAGCTGAGCATTTACATGGCTAATACATACACAGGCAAGAATCTGACTCTT
ATTCTGTCTTCTCATGGAATCATGCCAAAGGATTTATTTCATCATACACTTCTTGTATTTCTTCTTTACA
AGGAGCATTGTATTGATTTGAGGGAAAATTGTGTGTGGAGATTCGCTTTCTTTTCTGTGATTTCCTTT
CAGCGTAGTAAAGGGGATGTTACAAAATAGTTGTTATAGGAAGGGGCGTGTGTCTTAGGAGTTGAGA
AGACTACATTACACTAAGGTGGCCAGTTGTTGTTCTGGGCTGTGCCATCTCAACTAATTAAGAACTG
GATGGTGCCGGAAAGGGAAGGACTTCAAGCCAAGGAGGGGACCTAGGAACAGAAGCATGAATGAT
ATGAGGACTGAGTCAGAAGAAGCAAAAATTAACAAATAAATGCATAAATAAGTAAGACACGTGAGCGC
TAGGAAAAGCAGGATGGGTATGTCCACATGCTGATATCAAAAATACATGCACGTTAGGCAGGGTGCA
GTGGCTCATTCCTGTAATCCCAGCACTTTGGGAGGCCAAGGCAGGAGGATCGCTTGAGCCCAGGA
GTTCAAGACCAGCCTTGACCACATAGGGAGACTTCATCTTTACAAAAAATAAAAATAAAAATAGCTCG
GTGTTGTAGCACACACCTGTGGTCCTACCTACTTGGGAAGCTGAGGCAGGAGGATCATTTGAGCTC
AGAAGGATGAGGCTGCAGTGAGCCATGTTCTTGCCACTGCACCCCAGCTTGAGTGGCAGAGTGAG
ACTGTTTCAAAAATATATCTATATTTCACATTAGATTCATACCTATTTGAAAATTAGCATCAATTATTCTTT
TCTTTTTTAAATGTTTATCTTTAGTTATATAAGTAATAAATATTCATTACAGAAAATTTCAAACGTAGAAAA
GTATACGAAGAAAATTTAAATTGTTCTATTTATATTCATGTACCATCTTTGTAGGCATTTTGCTGTAGTTT
TTTCCAGTCACTTTCATAAATATACATTTGCATTGTACATGGTAAAGATTATACTGTGCAATTCTGTCTTT
TTTGTATGATTAAATTGAATTGGTTTTGAATTTTTTTAGATTTAACTTGTCAATTCCTTTTGGTTCTATAG
TTGTATAAAAGATATAAGCATATTGAGTTCTTGCCTAGTTTTATGTATCTATCATTAAAAAAAGAATTCCC
ATGAAATCACATCCCCTCCAACCAAAGCCCCATTCTCTGCTCTCTTTCACTGAAAAACTTCTCAAAAG
AGATGTCTACATTCTGACTCTACCTCTTTACCTCTCATTTATGCCTAATTTTTTTTAGTGAAGAAGTATA
AGGACATTGTTTTAAAAAATCAGTTTGCTAGGCCGGGCACAGTGGCTCATGCCTGTAATCCCAGCAC
TTTGGGCGGCTGAGGCCGGTGGATTGCTTGAGCTCAGGCGTTTGAGACCAGCCTGGGCAACATAG
CAAGACTCTATATAACATTACAAGAATTAGCTGGGCATGGTGCGTGCCTGTAGTCCCAGCTACCTGG
GAGGCTCAGGAATGAGCCTCGACCGCGTCACTCTACTCCAGCCTGGGCGACAGAGCAAGACCCTG
TCTTAGAAAAAAAAAAAAAATCAATTTGCTGAAGGCTTAGAATGAAAAAACAACCAACCAATCCTTGC
CCCTCCCCTTTTACTCTATGATCTATTACTTTGACATTGTTCTGTAGTCTTTCTCCCATGTGTGTATAGG
TGTCTATTTGTACATTCTTTTTTTGAGGCTTGATTTACATACCATAAAATGCACCCTTTTAAATATACAGT
TCAAGAATTTTGACAAATGTATGCACTCGTGTACCACCACCCCAGTTAAGATACAAAACATTCCTGTC
ACCCCTAAAGGTTCCCTCAAGGCTCTTTGCAATCAATCTTCCACCTACCTCCAGCCCCAAACAGCCA
ATAATTTTCTTTCTGTCACTGTAAATTAAATATTACTGTTCTAAGCCAGGCATAGTGGCTCACACATGTA
ATCTCAGCATGTTGGGAGGCCGAGGCAAGAGGATCATTTGAGTCCAGGAGTCCAAAAAAGCCTGG
GTAACCTGGTGAGATCCTGTCTCTACAAAAAAGAAAATAAATTAAAAAATAATAATAAATACACTTTACT
GTTTTAGAATTTCATATAAAAGAAACCATACAGCAATACACTTTTGTGTCTGGATTCTTTGCTTAAAATA
AAGTTTTTGAGATGTAGTTTGCTACATCAGTAGTTTGCTCCTTTTTATTGCTTTAGTAGTATGCTCTGAA
TGTACTGCAATTTATTTATTCATTCATCTCTTGTTGAACATTTGGGCTGTTTCCAGGTTTTTGCCATTAT

TATTTTTTTAAAACACTGCTGTGATCATCATATACATGTCTTTTTGTGGATACATGTTTTCGTTTATCTCA
TTAACCTGGGTATGGTATTGCTGAGTCGTAACATAAAAGTATATTAACTTCATAAACAAAACTGACAAC
CATTTTTCCATTGTTATTTCAAGTGGTTGTACCATTCCACACTCCCACCAGCAAAGTAAAAAAGTTCC
AATTGTTCCACATCCTCTTCAACACTTACTATTGTCAGTCATCCTAATTTTAGCCATTTTAGAATGTGTA
TAGTAGTATCTCATTGTGATTTGAATTTGCATTTCCCTGACAACTAATGATGTTGAGAATATTTTCATGT
GCTTCTTGGCTATTTGTATGTCTTCTTTTATGAAGTGCCTATTTATATCCTTTGCCCAATTTTTTATTGG
GTTGCCTTCTTACTAAGTTGTAAGAGTTACTTATATATTTTGAATACAAGTCCCCTGTCAGATACACATA
TTGTGAATTATTTCTCCTAGTCTGTGACTTACCTCTTCATTTTAATAGAGTCTTTCAAAAAGCAGGAAA
GTTTTTAATTTTGACAAGGTCTAACTTACTCTTTTTTACCTTTTATTATTTATTAAAAATTACTATTTTTGTA
TCCTAAAATATGTTTGCCTACTCCAAAGCTATAGTGATATTATCCTATGTTTTCTTCCACAAGTTTTATAG
TTTTAGCTTTTACATTTAGGTCCTTATTTCAAGTTCTCTCTCTCTCTTTTTTTTTTTTTTTTTTTTTTTTTTG
CATAGTTTGAGGTAAGGATTGAAATTCATTATTTTCCTTATGGATTTCTATCTAGTTTTTCCTAGACCATT
TGTTGAAAAGACTATCATTTCTCCCGTTGTTTACTTTCCTTGAAAATCAATTGTTTATATATGTTAGGGT
CTTTTTCTGGGCTCTCTACTCTGTTTTATTGAATTATGTCTATTCTTAATGCCAATTCCAGCAATACACT
TCAAAATACCCCATGAGTCAAAGATAAATCAATGGAAATTAGAATGTATTTTAACTGAATAAGTAAAATA
CAACATATTAAAATTTGCGGAACGCAGCCCAAGCAACAACATATAAGATTTTCAGAAACCCAATGTTAA
CCAAAAGACACCAAACACAAAGTATGAGTGATATGAGTGATTTCATTTACATAAAACTCATTAAAAGGA
AATGGAAATTAGTGTTTAGAGATGCATGCTGTAGCAGGTGTTGCTGGTGCCCTGTCGCATCCCCTTC
ACCTGCTCAGTGCACCCATCCCCTGGCTTCTGTGAGTATTGACTGCTAATAGCTTCACAGCAGCACA
CTTCGCCTGAGAATTGCCCTCCACCTCCAGGAAGCTGCCTTACCTGGCATCCTGCAGCCAATGACT
GACAGATAGGGGAATACAGAAGTTCAGTCTCCTGCCTCTAGACGGGACAACTCCGCGGGGGCATTC
TTGCTCTAGAGCTCCCCTGGAGATCAGATTAAGGATAGACCTCAGCTGAACCCACATTTTGAACTAG
CTTTTTCTCCTCTTCTAATGTGCCATCCAGTGCCATCCTTACTCCCTTACAGAACTCATCTGAAGGCA
CTCCCTCACCAAATCACTTGAACAAGAATTGCAATCTCAAGCTCTGCTCTTGGGGGATCAATCCTAA
ACTAGTTGGTTCTAGAAGTGGTCCTAAAAAGTAGATTTTAAAGATGGGATTCTGGAGTTGGATAACAG
ATGGCAATGAAGACCCCATCATCAATGGTAGGAGGAGCTACAGCCCCTGTGATGACCTGGGCCAAG
ATTAAGGTGGAAGGAATACACTGGCTTACGAATGCTCCTGGCATGTGAGAAGTTCAGGTAAATAGTA
ACTATGAAGATTGTGAAATTGGGTGGCTCTTGCTTAGAACCACTGATATGTTAAAGAAAGAAAATGAC
AAGCTCATATCAATTAATCACCGATTTAAAACAAAGTGAGTCAGAGGCTGGCTTCCTTAGCAGCATTT
AAAGAGCACCTCATCTCCTGCAGCTAGAGGGCACCCTCTGTAAAACTGGAAGAATTTCAGAGGAGG
AATTCTTAGCCTAGGTAAGTTGCTTATGCCAAAATCAGGGCCATGATAAGGAAGATATGGGACCCTGA
GGCTGAGCAAATGTGATTGAGACCCCTGAACTTCCAGTTCCCCCTGAACCCACTGGGCCTAGGCTT
CAAATAAGAAGGTACCTTACAAGACATCATTAGCCCATCTCAGAATCTGCCCTCACCTCCCCTCCTG
GCCATCAGATCAATCACTACAGTCAAATCTCAAGATGACCCAACTGAGGAAGTGCTGGGGCTGCTG
AGAGAAGAGAGAGATTGATATGCCAAGGCCGCTGCAGGACCCCACTGATGTGGACTGGCAAGACC
TGGGGCATTTGCCTGTGGGAGGCATGTGGATCAAGTGGGGTGGAATGTAAAGCTGAATGGAAGAG
GCTTTGTCAATGTGGGGGCACTCTCATGACACGAGATTTAATACCTGGTATGGGCCCTAGGAGACTG
ATTTAATCTGCTGCTAGGAAGGCTCTTGAAATTTTGGTAAGAGTGGTGGCCCACATTAAATGAAGTAG
AGTGTCTGAACTGCTGTGGCAGACTAGGGAGAAAGGACCAAAAAGGATCAGAGAGTGAGCATGCTA
GAATGAATCTACTCTCAAATACCAGAAAACCCACCAGTTGACTCTATCCCTCGGGGGGCCTAGAGGA
TTTGCCAACACATTCTAATAAATGAGCACCAGCATTATTTTTTTTAAGTTTTTTTTTTAAATTTTTGTGGG
CACATAGTAAGTGTATATGTTTATGGGGCACATGAGATGTTTTGATAAAGGCATGCAGTGTGAAATAAA
CACAGCATAAAGAATGGGGTATCCATTCCCTCAAGCATTTATCCATGAGTTGCAAGCAGTTCAATTAC
ACTCTTTACATTTTCATTTTTTGGTTTTTGGTTGTTTTGTTCTGTTTTGTTCTGTTTTTTGAGACAGAGT

CTTGCTCAGTTGCCCAGGCTGGAGTACAATGGCGCGCTCTCAGCTCACCGCAATTTCTGCCACCCG
GGCTCAAGCGATTCTCCTGCCTCAGCCTCCTGAGTAGCTGGGATTACAGGTGCCCACTGCCATGCC
TGGCTAATTTTTGTATTTTTAGTAAGACAGGGTTTCGCCATGTTGGCCAGGCTGGTCTTGAACTCCTG
ACCTCAGGTGATCCACCCACCTCGGCCTCCCAAAGTGCTGGGATTACAGGCGTAAGCCACTGTGC
CTGGCCTCTTTAAGTTACTTAAAAATGTACAGTTAAGTTATTATTGACTATAGTCACCCAGTTGTGCTAT
CAAACAGTAGGTCTTATGCATTCTAATTTTTTTTTAACCAGTTAACCATCCCCACCCCAGTCTTTCACT
ACCCTTCCCAGCCTCTGTTAACCATCTTTCTACTCTCTAAGTCCATGAGTTCAACTGTTGCACCAGCA
TTATTGAGACAGTTAATATTAGTCATTTTCTGTAGGCCTGGGCTAATGGTATCATAGACCTGGTTCCTA
GGTAGTAATGAGAATGAGAAGGTCCCGGAGAAGCAGAGGCCAGCTGGCGGCATTTATCTTCCAAAG
CAAGGTGGCTATAAGGTCAAAACTGCCAGTCAGGAGGCCTGATTCACAGAGATCTATGGCAATGACT
GATGGAATATGTTTTTCCTAAGGTGAGTTTGCCAGATTTAGCAAATAAAAATATAGGCTGCCCAATTAA
ATTTAAATTTCAGATAAATAACAAAGAATTTTTTAGTGTCAGTATGTCCCATACACATCTGACAATATGT
GTGTGTATTTTTATTTCCCCAAAATTGATCCAAGAATGAATGAACAGATGGCCAAGTTGAGCTTACTAA
TGGAAAGCCATAATTCCCTTCTGAGTTTCCAGATCTTAGCCGATTCTCATACCCAGAACCAATCAACT
GAAAGAAAGGCCAAGTCCCCATAAGAAAGGACATTACAACACCATGGCAAGTATATAAATTAGTGATT
CCCACAGTCCTTCCCCACAAGGACTCATTTATTCGTGGAAAGGGGAATACCTGGACCTTTCAAGGG
CTGTTGGATACAGGGCCCAAGTTGATGATTATAACCAGGGTCCCAAAGTGCTGTTTATGGCTGTTAG
AGGAGAGGCACATAAGAGTCAGGTGGTAAATGGGGTCCTTGCCCAGGTCCATCTCACTGACCCACC
CAGTCACCATCTACCCAGTTCCCAAAAGTATGATAATGGATATAGCAGTTGGCAGAACCCTCACCTTG
GCTCCTTAACGTGTAAATAAGAGCTATTGCTTTAGGAAAAGCCAGTGGAAGCCCTTGTAACTTTCCAT
GACTCCCTGCCAAGACAGTAAATAAAAAATAATATTGCATCCTAGAGAGAATGGTAGAGATTAGTGTC
ATTTTCAAAGACTTAAAGAATTGTGGTCCCATTATACACCCTATTTAATTCACTGATTTGGCCTTTGCAA
AAATTGAGGAATCATTGTGGATGAGAGTGGACTACTGCAAATTTAAACAAGTAGTAGCTCCAACTGCA
GCAGTTATGCCAAATGTAATATTTTTAATCAGAGCATTCACTCATAGCCAAGAGACAATATACTTTCCT
GGTCTTGCCTGGGGCTGTGTCAACTTGCCAGCTCTCTATCATAACATAGTCTGAAGGGACCTGGAC
AATCTGGACGTTCTGCAGAACATCACATTGGTCCGCTATATCAATGACATTGTGTTAATCACATCAAAT
GAGCAAGAAATTGCGAGTATTTGGAGGCCTTGGAGAATAGGAGATAAACTCTATGAAGATTCTGGGG
CCTGCCACGTAAGTGTGTTTTTGGTTTTTTTGGTTTTTTTTTTTTTAGATGGAGTTTCGCTCTTGTCACC
CAGGCTGGAGTGCAATGGCACTATCTCGGCTCACCGCAACCTCCACCTTCCGGGTTCAAGCAATTC
TCCTGACTCAGCCTCCCAGGTAGCTGGGATTACAGGCATGCACCACCATGCCTGGCTAATTTTTTTT
GTATTTTTAGTAGAGACGGGGTGTCTCCATGTTGGTCAGGCTGGTCTTGAACTCCCAACCTCAGGTG
GTCCACCTACCTCAGCCTCCCAAAGTGTTGGGATTACAGGCATGAGCCACCGTGCCTGGCCAGGT
GTCCTTTTTAAGAGTGCAAGGTCTTGGACCATCCAGGTCATTTCTTCCAAGTAGAGGCTATGTTATTT
TACCTTGCACCATCCCCTCAGTTAAGAAAGGGAATACCTTATAGGCCTCTTCATTTCCAAGTGACATA
AAACTCTTACAGCTTTGAGTGGAGCCCAGAGCAGTAAAGTGCTCTGCAGCAAGTCCATGCTGTGGT
ACAAGCAGCTCTGCAGCTTGGGCCACATACCAGCAACCCTGATGGCAGTTAGAAGTATCTGTGGTG
GGAAAAGATGCCTTTTGGAGTCTATGGCAAGTTCCAGCAGGAGAATCAAAATGTTGATACCTAGGGT
TCAGGAACAAAGCCATTCTACCTACAGCAGTGAATTATATACCATTTGAAAAGCAGCTCTTGGCATGC
TACTAGAGTAGATACACAGGAAGAAGTTGAGTAACTGACAATGGGACATCAGCTGACCATTAAACTT
GATCTGCCCATTATAAGCTAAGCAGTGTCAGACCCTCCAAGTCACAAGTTTGGGTGGGCGCAACAAT
CCACTGTAAGAAGGAAGGAATGCAGTCTGGCTGAGGCATGAATAGTGCCAGAGGACACAAGCAGC
AACTGCAGGAGCACATGGCCTCCATCATCCACCGCTACTACACTAATAATTCACCTTCAGTTCAACCT
GTGGCTACCTGATGGGCTATTCTTGAAGTCCAGCTGACAGAGGAGAATAAAGCCTGAGCTTCCTTG
ACAGATGTCATCTTAGTATGTGGATGCAAGCCAAAAATGGAATGCTGCAGTCCTGCAGTCCTGCATT

CCCACTAGGGAACGAGCTTGAAAGAGTAGTAAAGGGAACCGGGTGTGGTGGCTCACACCTATAATC
CCAGCACTTTTGGAGGCCAAGGCGGGTGGATTGCCTGAGCTTGGGAGTTCAAGACCACCCTTGGC
AACATGGTGAAACCTTGTCTCTACTAAAATACAAAAAATTAGCTGGGAGTGTTAGCACGTGACTGTAG
TCCCAGCTACTCGGGAGGCTGAGACACAAGCATCGCTTGAGCACCAGAAGCAGAGGTTGCAGTGA
GCCGAGATGGTGCCACAGCACTCCAGCCTGGGCGACAGAGTGAGACTGTGTCTCAAAAAAAAAAA
AGAAAAGAAAAGAAAAAGAAAGAGGAGTAAAGGGAAATCCTCTCAAAGGGCAGAGCTTCTGCCATT
GTACGTGGTTATTCACTTTATGTGGAAAGAGAAGTGGCCTAATGTTAGAATATACAGAGATCCATGGG
CAGTGGTGAGTGGCTTGATTTATTGATTAGGGGCCTGGAACAAGTAAGAGTAGAAGATTGGGGACA
GAAAAGTCTGGGGAAGATGCACGTGTATGGGCCCACGGGAGTCGTTACCAGATGTGAAGATCTTTG
TGTCACATGTGAACACCAAGCAGAGAACATCCGTCGTGGGAGAAGCACCACAACACAAAGTAGAAG
AAAATGACTTGGCCCGTTAAAGTCAACTAGCCTCTGTCCATGGCCATCTCAAGACTCAAACAGTGGG
GTTATGAATGTAGTAGCTATGGTGGCAATGGTAAATGCTAGACCTCGGCACAACAGGGTGAGCTGTC
ACTCACAAAGGTTGATCCAGCTACTGCCACTGCCAAAACAGAGTTCAGTGCAGAGCCTCCAATTTG
GCATCATCGTTGAAGGAAACCAACCAGTCATTTCTTGACAAGAGGATTACATTGGACCCTTTTCAAAC
CAAAATCTGGAAAGGGCTCAATTCATTGACAGGACAAATCAATTCATTTGACATACTCTGGGTATGAG
TTTGATTCTCTTACCTGCAGTGCCTGAGCTAGCCCTACTATCCAAGAGTTCACATACTGTCTGATCCG
TCAGCATTAGATCTTCCATAGCATCACAACAGACCATGGGACCCACTTTAAAACAAGGATGTGTCCGT
GGACATATGACCATGGGATCCACTGGCCCTATCATATACCACAACACCCAGAAGCTGCCAGCCTGAT
GGATCCATGGAATAGTCTTTTCAAGACATAGCTAAGGTATCAACTTAGAGATGACACCCTGCTAGGAT
TGGGCACTGTCCTTCAGGATGTGGTGTACATATTAAGCCAATGACTGTTATATGATACTATGTCTCCAA
AAGGTAAAACGCATGTGTGCAAGAGCTAAAGGTTAGAAGTAGGGATATTCATGCTTACCATTACTTCC
AGTGACCCACTTAGGAAATCTGTGCTTCCTGTCCCGCTAACTCTAGGCTCTGAGCTTCTAGGGGCCA
TGGTTTCCAGAGACAGAACACTTTCACCACAGCACCCATTAAACTTTAAGCTACACCACTGCCAAGA
GACCAGCATGCCAAAAAAAAAAAAAAAAATGATTTGCTCCCCCAGCAGAGTGAATCTTTCCATATCTGA
TGATAAGGAGGAGGCAGGGCTGCAGGGACAACTCAGATAGAATCCAAGGGCTCCCCTGGGGCATT
TTTTAGTGCAATTTTGATGGTAAATGGGCAAGTGCAGCAGCCACAACCCGAAAAGACCACGACGGC
CAGGGGCTCAGACTCCTCAAGCTATGTCTTCCCTGAGGTCAGCAGAGGTGCTAACTGAGGCTAGGA
GCAATCCAGAAAAGGGAACAAGCATTGCCTATCAGATGCGCCCTGGGGCAACTGTGCTAATGGGGC
TGTAATTTGTCCCATTAGGAAACTTTCCTTTTATAAGCTTCTCCAAAAAAAGAGATAACCAGAATCCCG
AAAAGGCTGTTTAAGGACAGGATGAAATTATCACATGAAGCGAATGGCTCCAAGCAGTGCAAGTGTT
AAACTGCATGGATGCTGTGGCTGCACCCAGGTCATCACTCCAGGGCTGCTCCCTCCCCCAACTGCT
GGGATTCTTGTACTCAGAGGACTCACAGTTGAGCTCTTTCAAAGACATTGCCTTCAGCAGAAGCAG
CTGCCTCACCCAACTTCGCAGCCCCCTCACCCAACTTCGCAGCCCCTCAATTCATAATACCTGTATT
TTTCATGGCCCCTTTGCTCAATTCAGGATAACTCAGGATAACTTGGGAAGGTGATCCCAGGCTAAGA
ACTCCCCATGGCTCTGGCTGAGGCTCCTGATATGTCTCATCGCAGTTCACCTCTCTCTGCCAGATGC
TGCCCCTCTCTCCCTTACAGATGATGCTCCTGTGCTTCCTCAGAGAGCCACCTGCCCAAATCTCCAT
CTTTTTTTCCCAGAAAACCCGATGGATAGCAATGCACATTTACAGACACGAAGGTAACCACCAGGAA
AAAAAAAAAAAAGTTGAAAAACTTAAAAGTTGTTTTCTCTAGGGAGAAGTGTAGGGGTGAAGCAAGG
TAGAACAGGGGCTATTGTTATACAGCCCTTCTATATGCATTGTTTTTAACATGCACGTATTACTTCCATC
ATTAAAACTTTTTTCATTTTGGCCAGGCATGGTGGCTCATGTCTGTAATCCCAGCACTTCAGGAGGC
CAAGGTGGGCAGATCGCTTGAGCTCAGGAGTTCGAGACCAGCCTGGGCAACATGGTGAAACTCTA
TCTCTACCAAAAATACAAAAATTAGTTGGGCGTGGTGGCAGGCACATGCAATTCCAGCTACCTGGGA
GGATGAGACAGGAGGATCCCTTGAGCCCAGGAGGGGGAGGTTGCAGTGAGCCAAGATCGCACGA
TTGCATTCCAGCCTGAGCAACAAAGTGAGACTTCATCTAAAAAAAACCCCAACTTTTTCCATTTGAAT

ATTATAAGAATTTGATACTGTCACTCACCTTTTTAAAACTATTTGAATACTTCCTACAGCCCTAAGAAAA
TCCTTAAGAATCCCTAACATAGCTTGCAAATCCCTTCCCATCTGGCCCCTGCTGAGTTTCCTATCTCT
TTTATTTTTTTCATTTTTATTTTTTGACAGGGTCTTGCTCTATTGCCCAGGCTGGAGCATAGTAGCACA
ATCATAGCTGACTGCAGCCTCAACCTCCTGAGCTCAAGTGATCCTCCCACCTCAGCCTCCCGAGTA
GCTGGGACTACAGGTGAGCACCATCACATCCGGCTAATTTTTTGTATTTTTTTGTAGAGATAGGGTTT
CACCATGTTGCTCAGGCTGGTCTTGAGCTCCTGGCCTCAAGCAATCTGCCTGCTTTGGCCTCCCAA
AGTACCATATCTGGCGAATTTCTTATATTATTTCTTACACTCCTCGTTCTCCCTGTTACCTGCCGTGCT
GGATTTTCCATTTCTCTTCCTCATCTCTGGACTTCAAACATGCTGCTGCCTCTATCTGGAATGCTTTC
CACTCCACCTGGTTCATTGCTGTTCCTCCATCAAATTTCAGCTTAGATGTCACCTGCTCAGGGGGAC
CTTCTCTTCCCCATCCCACCCCAGACTATCTTAGATCCCTTCTATCACTCCCACAGCACCTGAACGTC
TTTTTTCTTAGCATCAATCTCCTGAAGTTATCTGGGGGGTTTTTTAGAGATGGGATCTCACTATGTTGC
CTAGGCTGGAGCGCAGTGGCTATTCACAGGCACAATCACACTCACTGCAGTCTCGAACCCCTGGG
CTCAGTCAGTCCTCCTGCCTCAGCCTCCTGAGTAGCTGGGAATACAGGTGTACACCATCATGCGGG
GCATGGTTATTGGTTATGAGGTTATCTGTTTAATCTCCACATTTCCTTTTTTATCCAAGAGGACTATTTA
TCTTGTTTCTCCTGTATTACTAGCACCCAGCATATAACTACAGTACAGTAGACAGAAAATATGTATTAAA
TAAGTGAATTTCATCAAAATGTTTTCTAGAGTCCCAGCTCTGTAATCTGCAACTTAGATAACCTGCCTG
AGCCTTAGTTTGCTCAAGTATAAAATGGAAATAAAAATAAAAATAATACTTACCTTAGAGGGTCGTTTTG
AGGATTAAATAAGATATATCAGAAAAGCTCTTAGAACGGTGCTTGGATATAGTAAGCATTTGATTAATG
CATGCTTAAACATAAAGAAAGGTTACATCAATTCACACTCATACCAGCAGACTTTGACACTATCACTCT
TTACCCTGTTCAACATTGAGTCTTGTGTTTTTAAATATTTTTGTAAAGAATATAGGTAAAAAGTGGCATT
TTTTCTTTGGATTTAATTCTTATGGATTTAAGTCAACATGTATTTTCAAGCCAACAAGTTTTGTTAATAA
GATGGCTGCACCCTGCTGCTCCATGCCAGATCCACCACACAGAAAGCAAATGTTCAGTGCATCTCC
CTCTTCCTGTCAGAGCTTATAGAGGAAGGAAGACCCCGCAATGTGGAGGCATATTGTATTACAATTAC
TTTTAATGGCAAAAACTGCAGTTACTTTTGTGCCAACCTACTACATGGTCTGGACAGCTAAATGTCAT
GTATTTTTCATGGCCCCTCCAGGTATTGTCAGAGTCCTCTTGTTTGGCCTTCTAGGAAGGCTGTGGG
ACCCAGCTTTCTTCAACCAGTCCAGGTGGAGGCCTCTGCCTTGAACGTTTCCAAGTGAGGTAAAAC
CCGCAGGCCCAGAGGCCTCTCTACTTCCTGTGTGGGGTTCAGAAACCCTCCTCCCCTCCCAGCCT
CAGGTGCCTGCTTCAGAAAATGGTGAGTCTCTCTCTTATAAAGCCCTCCTTTTTCATCCTAGCATTGG
GAACAATGGCCCCAGGGTCCTTATCTCTAGCAGATGTTTTGAAAAAGTCATCTGTTTTGCTTTTTTTC
CAGAAGTAGTAAGTCTGCTGGCCTCCGCCATCTTAGTAAAGTAACAGTCCCATGAAACAAAGATGCA
GTCGGGCACTCACTGGAGAGTTCTGGGCCTCTGCCTCTTATCAGGTGAGTAGGATGGAGTGGAAA
GGGTGGTGTGTCTCCAGACCGCTGGAAGGCTTACAGCCTTACCTGGCACTGCCTAGTGGCACCAA
GGAGCCTCATTTACCAGATGTAAGGAACTGTTTGTGCTATGTTAGGGTGAGGGATTAGAGCTGGGGA
CTAAAGAAAAAGATAGGCCACGGGTGCCTGGGAGAGCGTTCGGGGAGCAGGCAAAGAAGAGCAGT
TGGGGTGATCATAGCTATTGTGAGCAGAGAGGTCTCGCTACCTCTAAGTACGAGCTCATTCCAACTT
ACCCAGCCCTCCAGAACTAACCCAAAAGAGACTGGAAGAGCGAAGCTCCACTCCTTGTTTTGAAGA
GACCAGATACTTGCGTCCAAACTCTGCACAGGGCATATATAGCAATTCACTATCTTTGAGACCATAAA
ACGCCTCGTAATTTTTAGTCCTTTTCAAGTGACCAACAACTTTCAGTTTATTTCATTTTTTTGAAGCAA
GATGGATTATGAATTGATAAATAACCAAGAGCATTTCTGTATCTCATATGAGATAAATAATACCAAAAAA
AGTTGCCATTTATTGTCAGATACTGTGTAAAGAAAAAATTATTTAGACGTGTTAACTGGTTTAATCCTAC
TTCTGCCTAGGAAGGAAGGTGTTATATCCTCTTTTTAAAATTCTTTTTAATTTTGACTATATAAACTGATA
AGTCCTCTCTACTTCACAGATTAAGAAATTGATACTCAAAAAAGTTAAATAACTTGTTTTAAACCACATA
GTAAGTGCCGAAGCCAATCTGTGAGACCAGGACTGTTTGTACTCTAAATGGCTGCACCACATGAGG
CAAAATGGCTCGTGATGGTTTTATTTCAAAGACCTAGAAAACACTATCACAGCTGGTGCTCCCGTCT

CAGACCCACAGCAACGATGTCTCCACTTCCTGCTTCATCTTGGGTTTCTCACGTCTTGAATGTGCAC
ACAAATCACCTGGGGATCTTGTTAAAAATGCAGCTTCTGTTTCAAAAGACCCAGACTGGAAATGGAG
ATTCCGCATGTCTAGTACGCTCTCAAGTTTATTAATCTGCTGCTGGTCCTAGGAACATATTGAGTAGC
GAGGGGCAGGATGTGACTCCTGTAAGGAGTGGCCAGGCATTTTCTTAGAGACCTGTGTTATAAAGTA
TGCTTTTCCTTAAAAAAAAGAAGAAGGAGGAGGAGGAGAAGGGCCAGGTGCAGTGGCTTACGCCT
GTAAGCCCAGCATTTTGGGGGACCAAAGTGGGAAGATCACTTGAGCTCAGGAGTTCAAGACCAGC
CTGGGCAAAGTAGTGAAACCCCATTTCTACAAAAAATTAAAAATTAGCCAGGCAAGGTGGCACACAC
CTATAGTTCCAGCTACTTGGGAGGCTGAGGTGGAAGGATCACTTGAGCCCAGGAGGTCGAGGCTG
CAGTAAGCCATGATCAAGCCACTGCACTCCAGCCTGGAGTGTCTCAAAATAGATAAATAAATAAATAA
ATAAATAAATAAATAAATAAATAAAAGAGACAGTATCAAAGACCCAATCACCTCTAGACATCTGGCATCA
TAGGAATGTGCCCAGTCTGCTCTGGGGATAGGAAAGTGGGGATCCTGTCTCCCCCTGTGTAGAGGT
TTCAGTAAAAGAAAGGCCTAGGTGTGCAGAAAGCTTTCAGGCAATGCCAGGGAAACTGATCATTGTA
ATGAATCCAGGGTATTGCTGAGTGAGGGCATCCTGGAGGGCCCGGTGGAAATGTGGTCAGGCTCTT
CAATGCACAGGCCCTAGTTGATGAGTAATCAGGGTTTCAAATATTTCCATCTCTGTCTCAAGCAGAAA
ACAAATGGAAAAACTGAACCACCAGAAAAGCAGAGCCAGAGATGGAACAAGAATCCCAGTGTTTGT
ACCCAACCAAGAGCGTGTTTTTCTTCCACAGACACCAATGTTCAAAATGGAGGCTTGGGGGCAAAA
TTCTTTTGCTATGTCTCTAGTCGTCCAAAAAATGGTCCTAACTTTTTCTGACTCCTGCTTGTCAAAAAT
TGTGGGCTCATAGTTAATGCTAGATGCTTCCTTCCTCTATTTCCCCCCAAATTTCCTGGGAACCCCTG
GTCAATACCAGCAGTAAGTTCCACTGTTCTAGGGTGTAGAAATGGCTGTGACCCAGCAGCAAGAGG
GAAGGACATCAGATGTCATCAGTGGTCATACTGCAACACAGCCCTTTTTCTGTTTAGGAATGCAGGT
ACCCACAACATTTACTAACACTTTTTTTTTCTTATTTATTTTCTAGTTGGCGTTTGGGGGCAAGATGGT
GAGATATGCTTTCTTTCTTTCTTTTTTATGAAATCACCCCATCATTCTTTGTAGTTATGAATGGAGCTTT
CTCTTAGGCCTCCCACAGAACTTCCACAGAGGTCAGGAAAAGGAGTTTCTGCCATCTACCCCTTTG
ACTTTCCTCACAAGTCTGGAGATATTTCTAGCCCAGAAGAGGGAAGCAACAGAGGCAGGAAATAATG
AGTCTTAACCATACAAAAGAAAAATTGAGACTTAAATGAAGTTGAAAGCACTAACAGTTTTCATTTGTT
TGCATTTCATATTTGATGTGAGATTCTGCAGAGGAGACGTAGCCAGAATGCATGCACAGGGTTACTC
TGGATAAGCTGCTGGGGCAACATTTGGATGTGTGTTCAGAATCACATGTCTGAATACTCTGAATATAT
GTGTGTACATGTGTATTTATGCAAGTGCACATGCATATGAGTGTGCCCGGCCTGAACTTACTCTCTCA
ACCACAGCGGTAGAGTCAGGAGTGTTCCAACATTGGAAGCCCCTCTATTCAATCAGCTCTTCCAAAC
TGAGTGAACCAATGTTGTATTTAATGGCAACCATGGCTGGACACCATGGCTCACACCTGTAATCCCA
GCACTTTGGGAGGCCGAGGTGGGCAGATCACTTGAGGCCAGGAGTTCGAGACCAGCCTGGCCAA
CATGGCGAAACCCCGTCTCTACTAAAAATACCAAAATCAGCCAGACATGGTGGTGTACGCCTGTAGT
CCTAGCTACTCGAGAAGCTGAGGCAGGAGAATCGCTTGAACCTGGGAGGCAGAGGTTGTAGTGAG
CCGAGATCACACCACTGCACTCTAGCCTGGGTGACACATCGAGACTGTCTCAAAAATAAAATAAAGA
CAACCATTATGCCAGCCTAGATTCCGCCATGCTGCCTAATTTGTAGTGTCCTTAGGAGCCATTTTTGT
AAATAGTCATCAGATAAGATGTAAGGCCCATAACAGCTTTTTCTATGCAGCTGAGGGAATTGGAAGAT
CCATTGTTTCCTAAGAGTTGAGGGAAGAGTCCCAACCCACGGGAGCAGGGTCTGATCTTCATTGCC
GATAGAAACATTACTAATGGCTTCTTACTGTTTCCTTTTCAGGTAATGAAGAAATGGGTAAGAAGATTT
CCACTCTATCTAGCAAAAGTTTTCAAATATGGAATGAAATGCTCATAGAGTACAATCACAGTAACAAAC
CCTGAGAACTAAAACTATTAAAGGGAAAATACAAGTATCTTTCAATGGGATCCGTATGAAACTTGCCT
GTATTTGTTGCTAGCTGTCATGTCAGATTATAGCTGTGCATATATGTATCTCTGATCATACACATATGGA
TGTGGGTTGGAGCTACCATGTGTTTTTGTATAAGCCATGAAATCTTTGAAGGCAGACAGAGACAGTG
TCTCATTTACCTAGCCCAGTGTCTGGCACATAGTAGGTGCTCAATGAATATTTTTTGAATGAATAAATG
AACAAACATATGAACACATTGCTAATTACCTCCCCTCAAGAAGCTGATGGTCTTGTGTGAGAGACAAA

TAATTGAAAATATAGTGAGTTGCATGTTATAATATGGGTAGATACAGAGTAAAATGAAGTATAAAGAGGG
GAGTGGTCAACTCTACTGAGTGTCGTTGGGAAAGGTTCCCTGGGGGAGGTGGTCCTTGAGCTGAA
TTTTAAAGGATAAGCTTATGTTTTAGGGAAGAAAAATATTTTATGCAGAAGAGATAAAGCTGTATAGTAT
GAGGATAAGAGTCTAACTGAGCTAGATCAGAATGTTTGAATCTTGGCTCAACTCTCTACTTGCTGGGT
GTGTTTGAGTAATTTACCTAACTTTTCTGTGCCACAGCATCATCATGGTACAATGGAAATAATAGTGCT
ACCTAACTTGTAGGGTTATTATGAGGACCAAATGAGTAATTCATTTAAGGCACTTAGAACATTATCTGA
CATAAAAGGCAGTAGGAGGGCCGGGCATGGTGGCTCACACCTGTAATCCCAGCACTTTGAGAAGCC
GAGGTGGGAGGATCACCTGAGGTCAGGAGTTCGAGACCGGCCTGGCCAACATGGTGAAACTCCAT
CTCTGCTAAAAATACAAAAATTAGCCAGGCATGGTGGCAGGTGCCTGTAATCTCAGCTACTCAGGAG
GCTGAGGTAGGAGAATTGCTTGAACCTGGGAGGCGGAGGTTGGAGTGAGCTGAGATTGTGCCATT
GCACTCCAGCCTGGGCGACAGAGCAAGACTCTGTCTCAAAACAAACAAACAAACAAACAGACAGTA
GGTGAATTTTAGCTATTAATACATGGAAAGCATGCTGACTATAGATGATAAGCATTAAAGTTTACTGAG
CATGTATGTTTTAGGCATTGCTCTAAATATTTTACTTGAATTTCCTCATTTAATTCTTCCAACACCCCTA
CTGTACAGTTAAGGAAACAAAGCCTCAAATAAATACAGAAATAAACAAAAATAAGTAAACAATCCAGTC
CTGGGGATATAAATGCAGATTTAGGCCAAGTGCCATGGTTCATGCCTATAATCCCAACACTTTGGGAG
GCCAAGGCAGGAGGCTCGCTTGAGCTCAGAAGGTTGAGGCTGCATTGAGCAAAGATTGTGCCACT
GTACTCGAGCCTCTGTGGCAGAGAAAGACCCTGTCTCTGAAAAAATTAATAAATAGAAATTTAAAAAT
AAAAAAATTTTAATGCAGATTTATATGATACCGAAGTTCATTTTCTCAACCATTATGAAATACTGTTTCTG
GATATGTATAAAATCTTTGTGAGCACACATATCTTTTTTTAACTTAACTTTCATTTTAAATTCAGGGGTAC
ATGTGCAGGTTTGTTATATAGGTAAACTTGTGCCATGGGGGTTTGTTGTATAGATTATTTCATCACCCA
GGCATTAAGCCTGGTACCTGTTAGTTATTTTTCCTGATCCTCTCCCTCCTCCCACCCTCCACCTTCTG
AGAGGTCCCAGTGTGTGTCATTTCCCTCTGTGTTCATGTGTTCTCATCATTTAGCTCCCACTTCTAAA
TGAGAACATGTGGTATTTGGTTTTCTGTCACTGTGTTAGTTTGCTAAAGATAATGGCCTCCAGTCCCA
TCCATGTTCCTGCAAAGGACGTGATCTCATTCTTTTTTATGGCTGCGTAGTATTCCATGGTGTATATGC
AGCACATTTTTTTATCCAGTCTACCACTGACAGGCATTTAGGTTGATTCCATGTCTTTGCTATTGTGAA
TAGTGCACAATGAACATACGTGGAGCACATTTCTGTCTAAGCACAGACATCTAGACCCTTGTGTGAG
CATGAGTTAAGTCTAAGCTCTGCTACTGAATTTGTGCCAATAAAAGTTGTGAGCAATTTTCTTTACATT
TTTTTCAAACAAACACACCCAGCAGAGTATAATGTCTATGTACTTTATTTATGATTTCTAGTTCATTTAA
CATGTCTAAGAAACATCCGTGTTGAAAAATTATTTATAAATTAAAATAATATAAACTATCTACTGTCCTTAT
ACTCAACTCCCAATTATAAGCAGGTGGAAAAACCTGGAGAATGTTTTGTTTACATTCTGTGCAGTCTT
TGTCAGAGGGCTGCCTGAGCAACTGGGTCAGAGTTTAGTTCTGCTCTGGGAGTAGCAGGACCTCA
AGAAGGAAAGGAGGAAAGGAAGTAACTTTTTCTTGAGCACCTGCTATGTGTCATTCACTTTCACCTT
CATAATCCATTTAATTTTCGCAAAAACTTTGTGAGGTTGGTGTTTTATCTCCATTTCCCTGATAAAGAA
GTTGAGGTTCAGCAAAGTTAAATGACTTGCCCTCAGTCACACAGACTAGGGCAGATCCAGGATTCAA
ACTCAGGGCTTCTGACTCTTGAGTCCAGAGCTCTGTCCCTGACAGCAGCAGCACTGCCTCTCCTCT
CTTCCAGCTGTTATGTCCAGACTGTAGCAGAACCCAGTGTTCCAGCCACAAGTTTTCCAGGAAATAA
TAAAGGACTCCTAGCTCCACCTCCCAGGGCAAAAATGGCTGCTGTGGGAAACACAGGCTGGACCTA
CGAATGGCATTAGTGGTTTATTAGTTGATTTCAGTTGTCCACACTAATAGGCCTCCCTCTAACAAAAAT
AATTGAGAGCTGATTATGCTCAGATATAATGTAAAGTGAAGCCACTTTTTATTGGAAGAAGCATTCCCT
CAAAACGTGTAGAGTATTTCACATTATTTAAAGGCAAATAGAGAGAAAATTATATGGAATAAGAACAAA
GATGTTTCTTCTCTATTATGAGGGACTCAGTTCTGAGAAAGGATTTTAAATTGTAAGAAATAGGTAAGT
CCACGAATCAGTGATTCAGTGGTGTGGAGAGCTTTATTTCTGAGAAGGCCAGTAGCGCTCCCTTCT
GACAAGCAAATCTAAGACCTGGATGACAGATGACTTCCTGCATTTGGTTGGTTCTTTTGTCATTCATA
TCTATCTGTAATACAGTTCTGGCTAATTTAAGAGGATAAGCTTGAAGACCTCTGGAATTTTTCGGCTTT

AGGACTTTAAGGCTTTCTGAGCTTCAGTAGATCTAGATCTAGGAGCTCATGCTGGTATATTCTGAATC
CGATGTATCTGAGTTACATCTATGAGCTACTTAATAAATATATCTATGAGCTAAATCTCATAGGCTAAGC
ATGAACCTCACCTCCAAGACTCGGGGTTCCTAAATGGATGAGACCCTCTTTGGGAAGTCTTGTGGG
CAGTGTCTAATTCCACTAGAAAAGTTTTACCTACAATTTAAACTTAAACCATGATATTTTCTTACTGCTG
TTTCCTTTTTTCATTTTCAGGTGGTATTACACAGACACGTGAGTTTATTGGTCTTTTATTTATGCCCTGT
CTGAGGATGCAGATTGGTGGGTAGATGAGAAGGAACTGATTGAGAGAGATTAACCCCAAGAACTGAT
ATCTTCCCAGCATTGCATTCTCAACTCCATTTTAGAAAGGTTCCAAATAGGGACTTCTGTGGGTTTTT
CTTTACATCCATCTTACCCTTCCCAAGTCCCCATGTCCCTGCGTAAACCCTAAAGCCACCTCTCAAAA
GGTTCTCTAGTTCCCTTCAAGGTTCTCTAGTTCCCTTCATTCCACATATCTCCTCTTCCACACCCTCTA
GCCAGTAGAGCTCCCTTCTGACAAGCAAGTCTAAGATCTAGATGACAGATGACTTCCTGCATTTGGG
TGGTTCTTTTGTCACTAATTTGCCTTTTCTAAAATTGTCCTGGTTTCTTCTGCCAATTTCCCTTCTTTCT
CCCCAGCATATAAAGTCTCCATCTCTGGAACCACAGTAATATTGACATGCCCTCAGTATCCTGGATCT
GAAATACTATGGCAACACAATGATAAAAACATAGGCGGTGATGAGGATGATAAAAACATAGGCAGTGA
TGAGGATCACCTGTCACTGAAGGAATTTTCAGAATTGGAGCAAAGTGGTTATTATGTCTGCTACCCCA
GAGGAAGCAAACCAGAAGATGCGAACTTTTATCTCTACCTGAGGGCAAGAGGTAATCCAGGTCTCC
AGAACAGGTACCACCGGCTCTTTAGGGAGGACCATTCAAAAGGGCATTCTCAGTGATTTTCCCTAAC
CCAGCTCACAGTGCCCAGGCGTCTTTGCGCTTCCTCCCACACTCAATCCTGGGACTCTCTGGTACC
ACACGGCATCAGTGTTTTCTGGAATATAGATTAAACACCAATATGAGGCTTCTGGGTAACCCCAGTCT
GTGCGAGATCTAAAATAGCAACTCCCTAAGAGACAGGACTGGGTCATTTGCACCGCATCACACCCA
GGTTCATAGCACACCAACATGAGTTTATCTAATGCTTCCTCCAGAGATAAATTTTTCAGAAAGGTTTG
CAAAAAACACTCAAGGCCACTATAGTAAAATGGCATAAGCTAAGGTATAATAATAAAATAATAACAATAC
TTAACATTTATTGAGTGCTTATTAAGTCTCAAGCACTGTCTGTACCCAACACTTATCAAGGATTCTTTTT
CATGTAATCCTCTCAACAACTATATGGGTTAAGTATCATTTTATTCCCATGAGTAAAGGGATGAGGAAA
CAGAGGGTTTGTGAGTTGAAAACACATTTCACGCTTCTCACAGCTAGTGAGTAATAAAGCTGGGACT
CAAACCCAGGGCTGTTTGACTCCAGTGCCTCTACCCACGGCCACCACTCTTTGCTTGTCAATGTTG
TTCTAAACATATTGAAGGGGGGGCTCTGACCGTGGCAAGCGTGTGAGTAGTAAGGGGAGAATGGCC
TTCATGCACTCCCTCCTCACCTCCAGCGCCTTGTGTTTTCCTTGCTTAGTGATTTCCCCTCTCCCCA
CCCCACCCCCCACAGTGTGTGAGAACTGCATGGAGATGGATGTGATGTCGGTGGCCACAATTGTCA
TAGTGGACATCTGCATCACTGGGGGCTTGCTGCTGCTGGTTTACTACTGGAGCAAGAATAGAAAGG
CCAAGGCCAAGCCTGTGACACGAGGAGCGGGTGCTGGCGGCAGGCAAAGGGGTAAGGCTGTGGA
GTCCAGTCAGAGGAGATTCCTGCCAAGGGGGACGACCAGCCTGGGCCAGGGTGGGTGGCAAGTC
CACAGCTAGGTCAGAACAGCTTCTCTAGAGCTTCTATGCACAGCTTCTATTACTGTGATGACAAGATC
TCAACAGACGGTTTCAAATCTCACATCACTCCCCTCCTTCCCATCCTAGAAAAGTGCAAAAAAGTTTA
TGAAAGTGATGGGCTTCCTCACATACCTGTCAATGCCTGCAGTCATCCGATTCCGCCCCTAAGCTGT
GGGAAGAGAGACTTTGATATATTAGCTCCTGCCTTTTCCTTTCCCTTCCCCTATGGAGAGAAACAATG
GGAGGATCTTGAGCTGAGGAAAGTCACAAAATGATGAGAAGAGTGTAGGGTCCTTAGAGATGAATG
AAAGAAAAAAAAAGAGAAAGGACGTCTGAACAGAAAAGGGAGCGGTAGAGGAGAGAACAATGGGG
TTTGCCATTCTCTATCTGGGTCTCACTGGCACAGACAGTGCTGCAAGATTGGTTCCCTCATGGGAAT
GAAATGTTTCCCCTCCTTCCTCCGCAGGACAAAACAAGGAGAGGCCACCACCTGTTCCCAACCCAG
ACTATGAGGTAACGTGGGATAGAAATGGGCCAGGACGCTGGAGGGGATGTCCCTCCAGGGGGGAA
GGAAACAGATGGGATGGCCCATCTTGTCTGCCAGATGCCTCAAAGCCCCTCACTCAGGGCTTCCAT
TACAACCCTCTATGTGCCACCTCTGCGTCCTTCATGGTAAAACAGGACTGTCTCAAAGGCTGCATGG
CTTCCACAACCATGGAGAGGTGGAAGCTTGCAGGAGACATACTCCTCTTTCTCTGGCTTATTCATTG
ACTGGGATACAGCCATGGAGAATATTATATATGCAAATTCTAACACAATAAATTCTGGGCTGATATTCCA

CCAGCATGCACCAGTATAGCGAGTTATTGAAATATTAAAATTATATAAATATTATATAAAGTTATTGAAAT
ATTAAAATACTCATTGGGAAATAGCCCCAAACTTTGCTCACCCCAACCCACCCTTACACACACACATA
CACACACACACACACACACACACACACACACACACACACGTGACCAGACATCCCAGTCCCTCCCCT
ACCGGGCTGCCTCTTGAGTTGGGGTAACAAAGAGTTAATGCCTGGCATGGCAGAGGATCACCAGGA
TTGTTCTAGTTGATTGGTATGTGTGCACTCCTAGTTGTTAAATATTTTCACTATCACACCTGGATATACT
CAACAAATATTTGTTGAGCCAAATACTCAACACCAGCCAAACACGTAGTATTTACTTTAGCTTAAGCGA
ATTATTTAGCCCTGACAGAAGCCCTGGAATGTGGGTCTTTAAGTTCCTATTTTTGAGATGGGAAAGCT
GAGGCTCACGGAAGGAGGTGACCAGCTCAAGTCTCCTACCGTCCATGCCAAATTAGAATTCCAGCC
TGCCTCCTGACTTCAAGTCCAAAGTTCTTCCCACGCACTAAAGCTAGCTCTTCAGTGTCCTTTCTTA
GGAGGTACTTCCTCCCGCACCACTGACCGCCCCCTCTCTATTTCACCCCCAGCCCATCCGGAAAGG
CCAGCGGGACCTGTATTCTGGCCTGAATCAGAGACGCATCTGACCCTCTGGAGAACACTGCCTCCC
GCTGGCCCAGGTCTCCTCTCCAGTCCCCCTGCGACTCCCTGTTTCCTGGGCTAGTCTTGGACCCC
ACGAGAGAGAATCGTTCCTCAGCCTCATGGTGAACTCGCGCCCTCCAGCCTGATCCCCCGCTCCCT
CCTCCCTGCCTTCTCTGCTGGTACCCAGTCCTAAAATATTGCTGCTTCCTCTTCCTTTGAAGCATCAT
CAGTAGTCACACCCTCACAGCTGGCCTGCCCTCTTGCCAGGATATTTATTTGTGCTATTCACTCCCTT
CCCTTTGGATGTAACTTCTCCGTTCAGTTCCCTCCTTTTCTTGCATGTAAGTTGTCCCCCATCCCAAA
GTATTCCATCTACTTTTCTATCGCCGTCCCCTTTTGCAGCCCTCTCTGGGGATGGACTGGGTAAATGT
TGACAGAGGCCCTGCCCCGTTCACAGATCCTGGCCCTGAGCCAGCCCTGTGCTCCTCCCTCCCCC
AACACTCCCTACCAACCCCCTAATCCCCTACTCCCTCCACCCCCCCTCCACTGTAGGCCACTGGAT
GGTCATTTGCATCTCCGTAAATGTGCTCTGCTCCTCAGCTGAGAGAGAAAAAAATAAACTGTATTTGG
CTGCAAGAGTTGCTGTCCCTGTTTTCTGAGAGACCCTCAGAGGCTGCCTTAACTCCCAAGACCATT
CAGGTTCTCACTATCTTTGACTAAATTCTCAGCTATGACCCCAGATGCGAGCACTCTCTGTTTCCACA
GCTGGTAGGGCCGTTACCTGAAGCTCTCTTTAGAGAGGGCATAAGGGAATAAGAAAAAAAAGAGAG
AGAGAGAGTGAAGGGAGGGGAAAGAAAAGAAAAAGAGAGAGAGAAAGAAAAATAAAAGGAAAGGG
AAAGAAGGGAGGGATTGAGGAAGGGAAGAAAGAAAGAGGATGGGAGGGAGAGAAAGAAAGAAAG
TTTCACCAACTTCTCTTGGAGGAAAACATCAAACGTGGCAGAGAAAGACTGATGGGTCCCTGATGG
AGTGGTTGGTAGAAAATGGACAAGACAGAAATAGGCCTTGAAAGACAGATGAAGGTACGGAAGAGG
ACGGTGGCGGTGGAAGCCGGGCTTGGAGATGGGACACAGATTTCCACAAGCTGCCAGGAAAAGCT
GCGAGCCAGGGCTGGGGAAGTGAAGGAGGGAGGTGTCTCAAGCAGGCACACCCCCACCCTGAG
GCAGCCGCCTGCAGCCAGAGGCGGGCTGTGGTTAAGCAGTGCAGGATGTGGGCTGCACTGCTAA
GCGTGGCTTCTGGGAGTGAGGGTGGGAGAGGTACAGCGGCAGCTGGGCGGAGGCCCGTGTGAG
AGCGCTTTGTTCTCAGTCTCCCACAGCACACTCTGCTTGCAGAGAGAGGAGGCTGGAGCATCCAG
GAACTGTCCTAATCCAGCTCAAACCAGTGGGCTCTCTTCTCTCCCTCCCTGCACTAATAATTCACCC
TGCTGATTTATGAGTGCGACCACCACTAAAAAGGCCATTCGTGACCAGCAGCCTCCACATACTGCTC
CCCCTCAGCCTTGGGCTGTAGCTACAGCTTCTATTTGTTTTTGGTCGCTGTTGAAGAGTCACTTCCT
TCACTCTAAGCATCAACACTTAGGTTCAGAAGATGCCTCCATGATGTTGTCCATCACTAGCTGCTGGA
TGGCAGTTTTGTCCCAAGTATACTTGATTGTGCCCTGTCAAGGGGAGGCATGGCTCAGCGTAATACA
GGCAATAACAGCTGACATGTGTTGAGCATTTACTCAATACATGCTTCAGAAATTGTACGTAGCACTTTA
CCTGCATTATCTCATTTAATCTTTATGTTGACACAAGAAGTTAGATGTCATCCTCATTTTACAGACGAG
GATACTGAGGCTCAGAGAGGTCAAATCATTGCCCAAGTCACAGATGGGGTGGGGCAGAGCTGGAA
CTAGCAGCCAATTTTAATGGTCAGAGCATCTACTCATTACTGACTGTGATCATAAAGAATGTTCCCCAT
TGTCGCACACTGGGGCCTGTCGGGGGTGTGGGGGCCAAGGGCAGGAAGAGCATTAGGACAAATA
CCCAATGCATGAGGGGCTTAAAACCTAGACGATGGGTTGATAGGTGCAGCAGATCACCATGGCACA
TGTATACCTATGTAACAAACCTGCACGTTCTGCATATGCATCCCAGAACTTAAAGTAAAATTTAAAAATA

AAAATTTTAAAAAAGAATGTGTCGGGCACGGTGGCTTACGCCTGTAATCCTAGCACTATGGGAGGCC
GAGGCAGGCGGATCATGAGGTCAGGAGTTCGAGACCAGCCTGGCCAACATAGTGAAACCCTGTCT
CTACTAAAAATACAAAAAATTAGCCAGGCATGGTGGTGGGTGCCTATAATCCCAGCTATTTGGGAGGC
TGAGGCAGGAGAATCACTTGAACCCAGGAAGCAGAAGTTGCAGCAAGCGGAGATCGTGCCATTGC
ACTCCAGCCCAGGCAACAGTGCAAGACTCCGTCTCAAAAAAAAAAAAAAAAAAAAAAAGAATGTTTC
TCATTGTCCTGACTTAAAAGCTCTGTATTTAATTGTTTATGCATTCCTTCAATGAGTGCTTTTCTAATCA
CTCATGTGGCTATCAGCACTATGCTAACTGCTAAGAGGATATCAATAAACAGTTCTTGTCTTCAAGGA
GCTTGCAATTATATAAAAGCGGTCAAACATGCAAAAAGTTAAAGAGCAACACAAGAACAGATGATGTG
ATTAAGCACTCATGCGTGTTACAGAGTGTTGCAGGTGTTCAGAGGCAGAAAAAGCTCCACGGAGGA
AAGACTTTATCTAGGCCTTGAAGGGAAGGATGAATAGAACTTGGAGATAAAGAAGGCACTCCAGGCC
AGACACGGTGGCTCACACCTGCAATCCCAGCACTTTGGGAGGCCAAGGCAGGTGGATCATGAGGT
CAGGAGTTCAAGACCAGCCTGACCAACATAGTGAAAACTCGTTTCTACTAAAAATACAAAAATTGGC
CGGGCACGGTGGCATATGCCTGTAGTCCCAGTTACTTGGGAGGCTGAGGCAGGAGAATTGCTTGAA
CCCGGAAGGCGGGGGTTGTGTGGGTGAGCCGAGATTGTGCCACTGCACTCCAGCCTGGGCAATA
GAGAGAGACTTCATCTTCGAAAAAAAAAAAAAGAAGAAGAAGGCACTCCAGATTGCAGAGAAGGTTA
TGAACAAAGACTTCCAGGCAACCAGCTCCGCACTAGAGAAGCAAATCGACTTAAGGAAAGAAGAAG
GAAAGTGGCTATTTGAGGGCACATGCTATGCTCAGGTCCTTTCTCAAGTTCTAGCCACGCCTTACTT
ACTCACTCCTTAATGCATCATGCAAAGTAGGTATTATTTTATCAGTTTCGCAAAAGAGAAAACTGACTT
TCAGCAATAATAGGTAACTTCCTTAAGTTCCCAGAGCAAGGAAGTGAAGGCCCTGGAACCTAAACCC
AGATCCTCTAGCACACTCTCCCACAAATGTCTGATTCTGCTTTGCTGAAATACAGTGAATATGGCAGG
TCCCCAAAGTACATCCTAGGACCTTGTGGTTCCTGGCTGTGCCTGGCCACATCAAAGCAGGGAGGC
TGACAATCTCTGTCCCTTCTTGGCTCCCTCTTCCACTCCCCTGAGTGGTTGGGCCACACTTTCAGTG
CTCCTGTCAAAAGGCCACCCAATCCCTGCTCTCCTCACCGTGAGCAGCTGAGGGCAACTCCAGCTT
CACGGAGAACAGAGAATAGGCCCACGGGGCCTGCAGCCAACCCCTTACCTACTCCCTCATGTGCTT
CCTCACCTGCCCTTCCTGCCCCAAGGATGGGACTTTTCCCACTCACCTAAAACCTTGATAGCCTCCC
CTTTCTTCTGCTCCAGTGTCTCATGACATCACCTCTCCCTCTCCCCAGTATTACCCAACTGTCCCCTT
TAACTTGATCTTTCCCCTCCACTGCATGCTCACATCTCTCCCATGCTAAGATAAGCTCTCCCTTGACC
CTGCATGCCTCCTGCCCAGATACTTCCCTTTCTCTCTCTGCCTTTTTAAGTCCCAAAGAGAAGTCTAC
TAGGCCAACTTCCTCATTTCCCTCTGATCCTCAGCCCTTTACAGTCGGGCTTCCTCCTCTCCCACTC
CAGAAACTCTCAAAAACATCACAGTGCCCTCCTCCTTGCCAAATCAAAGAAGCACTCCATGAGCCTT
TCAAGAGCAGACCTCTTGGAAGTATGTGAAAGATTGACTGCTCACTTCCTTGGCTTCCATGACATCC
TCTGCCCCTCCTTACCTCTCTTGAAATTCCTCCTCCCTGCTCCTTAAACACTGGTGTTCTTCGAGGTT
CTGTTCTCATTCCAATACCCTTCACACTTCACCTATTTTTCTTACTTTCCCATATCTCTACCCCCAGAC
CTGACCAGATCAGATATCCTCTTAGGCTTTAGACATATTCTGGATGTATCCATTGGGAGCCCACAGAC
TTCACAGACATTTCAAATTTGAGATGTTCAAAACTGAAGTCACCATTCCACCCCCCACCCACCCCCA
ATCCCTCAAACCATCTCCTCAACCTGCATGTCCTTCCCCAGCGAAGGGTACCACCATCTACACAGTG
CTACTGGATCAGAAACCCTGGAATTGGCCTGGATGCCTCCTGCACCCTCATTTTCCACATCCAAATG
GCACTCCCTAAATAGCTCTTCAAACTGATTTCTCTCTGTTTCTGCCTCCAGAATTATGTGTGCCATATC
ACTTCCTGCTGGAAGCTCTTCAAGGATGCATTACCTCATACAGAAAAAGCCCATACCCCCTACTTAGC
CTTACACTCCAGGTCTTCCGCCTTTGGACCCTTACCACCTCTCCCTTATTTCTAACACTCTGCAACTT
GAACTTCATGCTCTAGTAAAACTAAACCTCTTGAAATTCTCCCTGCTCTCCACCTCCGTGGTGTCTCT
CCCCATCTTTGCTCATGCTTTCTTCTCTGCCTAGACTGTTCTTTCCTTGTCTCCACTTTTCTTTTCTTT
TTCTTTTTTCTTTTTTTTTAGTTAAGCAGACTTTATTTGAGCAAAGAAAATTTATGAATCGGGCAGCATC
CTGAACCAGTAAACGTTCAAAGTGCTGCACCCAGCAACATGGGCAGGCCGAATTTATTGGAAGTAA

CACACAGAAACAGTTTGATGGGGTACAGTTCAGCATTTGCCTTATATGGACATGGTCTGATCAGCCT
GTGACTGGTTAAAGCACAGTGGCTGTGACTTGCTGAGATTCAACTATTTGTTACAAGAATATAGTCTC
AGTTGGGAGGCAGTTTACTTAAATACTAAGTTAGGTTGCAGTTCAACAGATATGCCAGCAGCTTTGG
GCCAAATTTAATTTAAACATTTCTGCCTTTTGGTCAGCATCTCAATTTTGAGAGATTGAACAGAACCTT
GGCCCCTGACCCCATTTTCTGTCACCATCATAATGGACTTGTTTTGTTTCAATATTGAATTCACAATTC
ACAACTTCACCTTAGTTGGATGATTCTTTATGCTTTCTTCATGTTTTTGTTATACTAACCCTAATAGACC
AGTTGATGTATAAAGAATGGCTGCATAGGAGCACTTAAGACCTCCCAGGGGAAGCAACACGAGAGA
CAATTATTATTATTTTAATTTTTAAGATGGAGTCTCACCCTGTCACCCAGGCTGGAGTGCAGTGGCAC
AATCTTGGCTCCCTGCAACCTCCACCTCCTGGGTTCAAGCAATTCTCCTGCTTCAACCTCCTAAGTA
GCTGGGATTACAGGCACCCGCCACCACAGCCAGCTAATTTTTGTATTTTTAGTAGAGATGGGGTTTC
ACCATGTTGGCCAATCTGATCTTGAACTCTAGTCCTCAAGTGATCTGCCTGCCTCGGCCTCCCAAAG
TGCTAGAATTACAGGCATGAGCCACCGCGCCTGGCCAGGGAGACTATTATGATGACTATGAGTGGAC
AATGCCAAGAAACTGAAGTACACTCCTTATCAGGAGTTTCCACAAACAACTAGTCAAAATCAAACAAC
TCAAAGTCCGTGCAATAAAGATAGTTCAGTAGTATTTGAGTCCAGCTGGTCAGTCTTGGTTCAGGAG
ATGATGGTGATTAGGGCCACAGTTCACTACAAAAAGACCTGATTTAAACTCACTCCATTAAATGAGAG
AGCTATATTAGAGATGTCTCAGAGACCTCCACCCAGCAACAGTGGACAGCCAGTGTTTATAAACAGA
AAAAGGAAGTGGCATTGGATATAGCCTGATCGGTTACAGGTCACCCTTTGCCTAATTTAGACACATCT
GATCAGTCTGCAGTCTGCCCCTGGCTGAAAGCTGGCTGCTATGGTTGGAGGAGACTTCCTTTCTTC
TTACAAAAATATGCCCTCAAGTCAGGTTGTAGTTTGTTTACATATTAAATGAGGTTACATCCCAGTCAA
ATCCTCCCTACAAAGTATCCCCTATTCTGACCTCTATCACCATAGACTAGTTTTTCTGGTTTTTCAACT
TTAAATAAATGGAATCACACAATAAAAAAAAAAAAAGAAGAAGAAGAAGAGAAAAAAGGAGGTTG
CAGTTCACTGAATTGCAGCATTTTGGAACAAATTTAATTTTTTTTTTTTTTTTTTTTTTTGACGCGGTCT
TGCTCTGTCACCCAGGCTGGAGGAGTGCAGTGGCACAGTCTCGGCTCACTGCAACCTCCACCTCC
TGGGTTCAAGTGATTCTCCCTCCTCAGCCTCCCAAGTAGCTAGGACTACAGGCGCCCGCCACCATG
CCCAGCTAATTTTCGTATTTTCACTAGAGGCAGGGTTTCACCATGTTGGCCAGGCTGCTCTCGAACT
CCCCACTTCAGATGATCTGCCCACCTCAGCCTCCCAAAGTGTTGGGATTACAGGCGTGAGCCACCG
CGCCCGGCCTGGAACACATTTAATTTAATTTAGCAGAACTTACTCCCTGGGCCCAGAGGACGTAGAG
GCAAGAGTTGCTTCCCATGATGAGTGCACCTGATGAGAAGGCACCTCCAGATGACTCAAAGGAAAA
CGAAGAATCAGGAGAGAAAGAGAAGCGCAGTTCTTGCCTGCTTCAGGGGAGGACCCCAAACTCCT
GGTTAATAGTAGGGACCTGCTTCCCCGTGCACCCCAGCTCCCTCTCCACCGTACAGTCGCCTCCTA
GGACTGGCGACCACAACTTGAGGCCTCCTCTGAGGCCCAGAGTTCGCGAAGTCAGAAGACAACCA
GCTGAAGTGGTTTGGGCCTTGTGACAGCACCAGATCCAGGGCGAGGTCGTGCTGCTGGTCCTCAG
AGTACACGGGCCGGTAGTTCTGCAGCTGCAGTGCGCCGCCGCACACCTCCTGCACACGCAAAGG
GCAGCAGGTGGTGAAGACCTTGGTGAAGGGCAGCGCGTGGCGCTGGGCCTGGGAGACGTTGAG
CTCATTCCTAGACAAAGTCTAGAAGGCCTCAAGTGGCCAGCCGATCCCTGACCCATGGGTGATGTT
GTGGATCCAGGCCTCGAGCTGCGGCATGACTCACCTTGCCTCTACTTCTCACATGACTTCTATTCAT
GCATCAAAACTAAACAGATGATGATTTTCTCCAGGAAGCTTTCAAAATCCCTCCCAGATGAGGCTAAA
AGCCGCTACTCCTTGGTACCAAACAACTCTATAACTCTGTCATATTCCTACCATCATATTAGATGTATCT
GTATGTTTAATGAAAGGATGGATGATAAATGGATGGGTGGATGGGTGAACGGATGGATGGATGATGG
GTGGATGAATGGATGAATAGATGATGGATGGATGAATTTTTTCACTACTAGATAAGAGGACAGAATCT
GTATTTTTCTTTGATATACCAGCTCCTACACAGTTCAGACCCTAGTTCATAATATATCCTCAATAAATGT
TGACATGAACTGACCTTGAAGATGATGCTGATGGAATTTCCACAGAAGACCAATAAGATACTGATGG
GAAATCCAGATATTACTAACATAGATAGAAAAATAATCAAAATGAACATACTAAAAACTTGCAAGCACAG
CGTCCCTACTCAAAGGAACTTGCTATTCCTCTAACTCTCTGGCTTGGATTGATGTCTAGGAACAGCC

AGCTAAAATCAACCTCATCCTAGGCAAAGATACTTTGGTTGGCAGTAGGGAATAAGAGAAAAAGTTTG
GTGTGGAATGAGAAAAGGTAAGAGCTTTCTTTACTGGGGGAACCCATAGTTATAGATTATTCAGAAAG
GGAAGCTCTCTAAAAACCAAGAGCCCCTTTTCAGATATGTCGACTCTTAGTTCCTTCAGACATCCCT
GGTACAAGTCTTGGCCCCATCCCAGAGCTGGGACATGATTAGGAGACAGAGATCTGCCCCCCTGGC
ATTTGCCCATTCCCTGGTCATGTAAAGTACAAGCAACTAGATATAATGACAAAAATATTTCCCAACTTTT
TCCAATGAGTTCTTGAGCTATTTAACCCTGAGCACTTATTTTATTTAAGAATATCTCAAAATGGAAGAAT
TGGAGGTGAAGCTGAGAAGCCAGGGAAACCCTGATAACTCAGAGACAATTCCAACATTGCAGTGTA
ACTATCACTCCATCATCACACCCTGTTCACTCAGCAGGCTATGGACTCAGGGGTTCTATTTACCCTAA
TCCCATATTCCTCCTCCCAAATTTAAAGCCGACCAGAACAACAGGTCTCTTAATTCCCTCTTTAGCTA
AGAATTTCTAGCCCTGAGAAATGCAAAGCAAAACCACAATGCGATACCACCTTACTCCTGCAAGAAT
GACCATAATCAAAAAATCAAAAAAAAAAAAAAGATGTTGGCGTGGATGTGGTGAACAGGGAACACTTC
TACACTGCTGATGGGAATGTAAACTAGTACAACCACTATGGAAAACAGCATGGAGATTCCTTATAGAA
CTAAAAGTAGAACTGCCATTTGATCCAGCAATCCCTCTACTGGGTATCTACCCAGAAGAAAAGAAGTC
ATTATATGAAAAAGATACTTGCACACACATGTTTATAGCCACACAATTTGCAATTGCAAAAATGTGGAG
GCAACCCAAATGCCCATCAACTAACAAGTGGATAAAGAAAATGTGGGAGGGGTGTGTATATATGTATA
TATATACGGGGGTGTGTGTGTGTGTGTGTGTGTGTGTGTGTGTGTATACATATGATGGAATACTACTC
AGCCTTAAAAAGGAATGAATTATGAATTAATGGCATTCACAGCGACCTAGATGAGATTGGAGACTATTA
TTCTAAGTGAAGTAACTCAGGAATGGAAAACCAAACATCACATGTTCTCACTCATAAGTGGGAGCTAA
GCTATGAGGAAGCAAAGGCATAAGAATGACACAATGGACTTTGGGGACTCAGGGTTGGGGATAAAA
GACAACAAATTGGGTACAGTGTATACTGCTCAGGTGATAGGTGCACCAAAATCTCACAAATCACCACT
AAAGAACCTACTCATGGCCAGGCATGGTGGCTCAGGCCTGTAATCCCAGCACTTTGGGAGTCTGAG
GTGGGTGGATCACGAGGTCAGGAGTTCAAAACCAGCCTGGTGAAAATGGTGAAACACCGTCTCTAC
TAAAAATACAAAAAATTAGCCAGGTGCGGTGGCAGGCGCCTATAATCCCAGCTACTCAGGAGGCTGA
GGCAGGAGAATCGCTTGAATTTGGAGGGCGGAGGTTGCAGTGAGCTGAAATCGCGCCACTGCACT
CCAGCCTGGGTGACAGAGTGAGAATCTGTCTCAAAAAAAAAAAAAAAAGAACTTACTCATGTAACCAA
ACACCACCTGTTCCCCAATAACCTATGGAAATTAAAAAATTTTAAATAAAGAATTTCTAGCCCTGAACA
TTGGTTTTATTTGTGAAGTCTGATCCCACAGGTACCTGTTGCTTACTGACATCATGGGCAGAAATATC
CTAGCAAATGTCCTTGGAATGACAGGTGGAAGAGTTGGAAGTGGAGTTGAAATCTTGAGTAGGTTTC
CTACCTGGACAGAAGCAGAGTAGTGCAAGATAAATGTAGATGCTGAGGTGAGAAAAATAAGCAGGAT
GAGGAGGCTGTAGATTACATATGGGATGCAGAGGATAGGAGGGATACACATTATAGAAGGAACACCA
CGTTGCATTTGAATGCAGTGTATAGGTGGGAAACAGTGGAAAGGAGGAATACAGAATAAAGGAAAGA
TTAAGAGTGTAGAAGGCCCAAGTCAAACTTGTAGAGATGAAAACTATGATATCTGAGAGGGAAAAAAA
TGAATTGGCCATATCTGGCCAGGCACGGTGGCTCATGCCTGTAATCCCAGCACTTTGGGAGGCCAA
GGCAGGCAGATCACCTGAGGTCAGGAGTTCAAGACCAGCCTGGCCAACATGGCGAAACCCTGTCT
CTACTAAAAAATATAAAAACTAGCCGGGCGTGATGGTGGGCGCCTGTAATCCCAGCTACTCTGCAGG
CTGAGGCAGGAGAATTGCTTGAACCTGGGAGGCGGATGTTGCAGTGAGCCAAGATTGCTCCACTG
CACTCTAGCTTGGGCCAGAGAGGGAGACTCCATCTCAAAAAAAAAAAAAAGAAAAGAAAAAAAATGCA
TTGCCCATATCTAACCAGCAGATTTGATATTACAGAAGAAAAGATTAGATTAGTGGGCTTGAAGACATA
ACAATAAAAACACAAAGAGAAAATGAAACGCAAAGAAAAAAATTAAAGAATAGAAAAGAATATATCATG
CTAACACTAATGAAAAGAAAACCGTATATTAACATCAGCCAAAGACATACTTCAAAGACCAAAGAATGT
TACCTGGAATACAGAGAGATATCACGTCATGTCACAATGACAAAGGAGTCAAGAGAATATAGCAATCC
TAAATGTTTATGCACCTAGTAACAGGGATGCAAAATTGGCAGGCATGGTGGCTCACACCTGTAATCC
CAACACTTTGGGAGGCCAAGGTAGGCAGATCACTTGAGGCCAGGAGTTCAAGACCAGCCTGACCA
ACATGGTGAAACCCTGTCTCTATAAAAAATACAAAAATTAGCCAGATGGAGTGGTGGACATCTGTAAT

CCCAGCTACTCAGGAGTCTGAGGCAGGAGAATCACTTGAGCCCAGGAGGTGGAGACTGCAGTAAG
CCGAGATCACGCTACTGTACTCCAGCCTGAGCGATAGAGCGAGATTCTGTCTCCAAAAAAAAAAAAA
ATTGAAGCAACTCACCACAAATCAATGATAAGCATTTGAGGTAATGCATATATTAATTAGCTTGATCTAG
CCACTCCACAAGATATACATATTTCAAAACATCATACTGCACACCATAAATGTATACAATTTATATTAGCT
TATTAAAAGTAAATAAATAATAAATTTTTAAAATAAAAAAACCCCAATTACACAATTTAAAAATTGAAGCAA
TCCCAGCATTTTGGGAGGCTGAGGTGGGCAGATCACCTGAGGTCAGGAGTTTGAGACTAGCCTGG
CCAACATGGTGAAACCCCATCCCTACTAAAAATACAAAAATTAGCAGAGCATGATGGCAGGTGCCTG
TAATCCCAGCTACTCTGGAGGCTGAGGCAGGAGAATCGCTTCAACCCTAGAATCAGAGGTTGCAAT
GAGCTGAGATAACGCCATTGCACTCCAGTCTGGGCAACAAGAGCGAGACTCCCTCTCAAAAAAAAA
AAAAAAAAATTGAAGCAAAAACTAATGAAACTTCAAGGAATAATATACAAATTCACAACTGTAGTTAGA
GATTTCAACATCCCTTTTTCAAAAGGTAGTAAAACAAGTACAATCAGCCCTCCCTATCTGAGGGTTTC
ACATCCACGGATTTAATCAACCATCAATCAAAAATACTTAGAACAAAAGTGGATGGTTGCATCTGCAC
TGAATGTGTATGGACTTTTTTATCATTTTTTCCTAAACAATACAGTATAATAATTATTTACATGGCATTTAC
ATTGTATTAGGTGTTATAAATAATGTAGAGACGATTTAAAGTATATTAGACAATGTGCATAGATTATATGC
AAATACTATGGCATTTTGTATAAGAGACCTGAGCATCCATGGATTTTGGTATCTGAGGATTCTAGAACC
AATCTCCTGTGAATACCAAGGTACCACCATAGACCAAAAATCAGTAAGATACGTAAACCAAATTCACTT
GATTAACATTTATAAAACCCCACACCCACCACCAACAGAATACACAGCCTTTTGAAGTGCACACAGAA
CATTTATCAAAATTGACCATATTTGGGGTCATAAAACAAATCTCAATATATTTTAAAAGATTAGTCACATA
ATGGAATTAAATAAGAAATAAATAACAGAAAGATAAACTAGGGATGTTAGCCAGAATAATGCTTCCTTC
AAGTATGTCCACATCCTAATTCCTAGACCTAACAAGAGGTCACCTCACATGACTGAAGGGACTTTACA
GATGTAACTAAAGTTAGAGACTCCGTGATGGGAAGATTACCCTGGATTATCTGGGTGTGCCCAATCTA
ATCATATGAGTCTTTAAAAAGGAAGAGGAAGGCACAAGCATGAGTTGCAGAGAAGAAACAAAAGGA
GAAAAAGAAAAGATACAAAGTGTAAGAGGGCGTCAACTCACCATTTCTGGCTTTGTAGATGGAGGAA
GGAGGTCATGAGGCCAACGAATGTGGGTGACCTCTAGAAACTAAAAATGGCTCTCATTTTACAGCCA
TTGAGAAAACAGGGACTTCAGCCATACAACCACGATGAACAGAATTCTGCCAACAACTCAACTCGG
CAGGAAACGGAGTCTCCCCTAAAGCCTCCAAAAAGGAATGCAAGCTGCTGACACCTGAGTTTTAGC
TCAGTGAGATCCATACTCTACTTCTGACCTATAGAACGATAAGATAATAAATTTGTGTAGTTTTCAGCC
ACTAAACCTACGGTAATTCGTTACAGCAGTAATAGAAAACAAATACACTGGGAACTCTTGAAATATTTG
GAAAGCAAATAACACACTTCTAAGTAGCCTATGAACAAAAAACAAATCAATGGAAAAATTAGAAAGTAT
TTAAACTGAATGATAATGAGAACAAAACATCAAAATCTTGGGGTTCCTGCTAAAGCAGTATTTAGAAG
GAAACGTATAGACAGAATTAAATGTAGATTAGAAAAGAATAAAGGTCTTAAACAATGAGCACAGCTTC
CACCTTGAGAAACTAGAAAAAGAAGAGCAAATTAACTCAAAGTAGACCCCAAAAAAAGAAAGAAAGA
AAGAAATAGTGAAGATTACTATTCACAATAGCAAAGATGTGAAATCAACCTAAATGGCCATCAACAGTA
GACTGGATAAAGAAAATGAGGTACATACACACCAGGGAATACCATGCAGCCATAAAAAAGAATGAGAT
CATGTACTTTGCAACAATATGGATGGAACTGGAGGCCATCATCCTTAGCAAACTAATAAAGAAAACCA
AATACTTCATGTTCTCACTTATAAGTGGGAGCTAAATAATGAGAACACATGGACAGAAAGAGGGAAAC
AATAGGGGCCTACTTCAGGGTGGAGGGTGGGAGGAGGGAGAGGTTCAGAAAAAAAACTATTGGATA
CTATGCTTAGTACCTGGGTGATGAAATAATCTGTACACCAAATCCCCATGTCACAAGTTTACCTATATA
ACAAACCTGGACATGATCCCCTGAACCTAAAATAAAAGTTAAAATATTTTTTTAAATAAAACTAAACCAC
ATATAAATGAAAAATAATTTAAACAATTATAAACAAAGTTTAAACAAAAATAAACAATATGAGAATAACCA
TGAAAAATAAAATAAAGATCAGAGCTGACATCAATAAAATAGGAAACAGAAAAACAACAGAAAAAGCA
ATGAAACATTTCAGAAATAAATAATCTGTAAGTAAAAACAAAAATCAATGTGACCAAACAATGGTCCAG
AGACCAATAAAACTGATAAATTTCTAGCCAGATTGATCAGGAAAAAAATAGAGAAGATATAAATTATCAA
TATTAGGAATGAGAAATAACATCACAGGTCCTACAGGTATTAAAAGAATAAGAAGTCAGGCACTGTGG

CTCACGCCTGTAATCCCAGCACTTTAGGAGGCTGAGGCAGGCAGATCTCCTGAGGTCAGGAGTTTG
AGACCAGCCTGGCCAACAAGGCAAAACCCCATCTACTAAAATACAAAAATTAGCCCTACCTGGTGGC
TCATGCCTGTGATCCCAGCTACTGAGGAGGCTGAGGCAGGGGAATAACTTGAACCCGGGAGGCGG
AGGTTGCGGTGAGCTAAGATCACGCCATTGCACTCCAGCCCAGGCAACATAGTGAGACTCAGTCTC
AAAAAAAAAAAAACAATAAGAAAGGAATGTTATTGTCTTAGTCCATTCACCTTGCTATAACAAAAATAC
AGAATGGGTGGCTTGAACAACATTTATTTCTCAGTTCTGGAGGCTGGGAAATCCAAGATTTCCCAGA
TTTAGTATCTGGTAAAGACCCACTTCCTGGTTCATATCTTCTTACCATATCCTCACATGGTAGAAGGGG
CAAGGGGGCTCTCTGGGTCCTCTTTTGTGAGGGCACTAATGCCATTCATGAGGGCTCCACCCTCAT
GACCTAATCACCTCCCAAAGCCTCCACCTTCTTTTTTTTTTTTTTTTTCAGTTTTTTTGTTTGTTTGTTTG
TTTTATTTTTTATTTTTTTATTTTATTTATTTATTTATTTTTTTAATTTATTTTTTTATTGATAATTCTTGGGTG
TTTCTCACAGAGGGGGATTTGGCAGGGTCATGGGACAATAGTGGAGGGAAGGTCAGCAGATAAACA
AGTGAACAAAGGTCTCTGGTTTTCCTAGGCAGAGGACCCTGCGGCCTTCCGCAGTGTTTGTGTCCC
TGATTACTTGAGATTAGGGATTGGTGATGACTCTTAACGAGCATGCTGCCTTCAAGCATCTGTTTAAC
AAAGCACATCTTGCACCGCCCTTAATCCATTTAACCCTGAGTGGACACAGCACATGTTTCAGAGAGC
ACAGGGTTGGGGGTAAGGTCACAGATCAACAGGATCCCAAGGCAGAAGAATTTTTCTTAGTGCAGA
ACAAAATGAAAAGTCTCCCATGTCTACTTCTTTCTACACAGACACGGCAACCATCCGATTTCTCAATC
TTTTCCCCACCTTTCCCGCCCTTCTATTCCACAAAGCCGCCATTGTCATCCTGGCCCGTTCTCAATG
AGCTGTTGGGTACACCTCCCAGACGGGGTGGTGGCCGGGCAGAGGGGCTCCTCACTTCCCAGTA
GGGGCGGCCGGGCAGAGGCGCCCCTCACCTCCCGGACGGGGCGGCTGGCCGGGCGGGGGGCT
GACCCCCCACCTCCCTCCCCGACGGGGCGGCTGGCCGGGTGGGGGGCTGACCCCCCCACCTCC
CTCCCGGACAGGGCGGCTGGCCGGGCGGGGGGCTGACCCCCCCACCTCCCTCCCGGACGGGGC
GGCTGGCCGGGCAGAGGGGCTCCTCACTTCCCAGTAGGGGCGGCCGGGCAGAGGCGCCCCTCA
CCTCCCGGACGGGGCGGCTGGCCGGGCGGAGGGCTGACCCCCCCACCTCCCTCCCGGACGGGG
CGGCTGGCCGGGCGGGGGGCTGACCCCCCCACCTCCCTCCCGGACAGGGCGGCTGGCCGGGCA
GAGGGGCTCCTCACTTCCCAGTAGGGGCGGCCGGGCAGAGGCTTCCCTCACCTCCCAGACGGGG
CGGCTGGCCAGGAGGGGGGCTGACCCCCCCACCTCCCTCCCAGATGGGGCGGCTGGCCGGGCG
GGGGGCTGACCCCCCCACCTCCCTCCCGGATGGGGCAGCTGGCCAGGCGGGGGGCTGACCCCC
CCACCTCCCTCCCGGACAGGGCGGCTGGCCGGGCAGAGGGGCTCCTCACTTCCCAGTAAGGGCG
GCCGGGCAGAGGCGCCCCTCACCTCCCGGACGGGGCGGCTGGCCGGGCGGAGGGCTGACCCC
CCCACCTCCCTCCCGGACGGGGCGGCTGGCCGGGCGGGGGCTGACCCCCCCACCTCCCTCCCG
GACAGGGCGGCTGGCCGGGCAGAGGGGCTCCTCACTTCCCAGTAGGGGCGGCCGGGCAGAGGC
GCCCCTCACCTCCCGGACGGGGCGGCTGGCCGGGCGGGGGCTGACCCCCCCACCTCCCTCCCG
GACGGCGCGGCTGGCCGGGCGGGGGGCTGACCCTCCCACCTCCCTCCCGGACGGGGCGGCTG
GCCGGGCGGGGGGCTGACCCCCCACCTCCCTCCCGGATGGGGCGGCTGGCCGGGTGGGGGGC
TGACCCCCCCCACCTCCCTCCCGGACGGGGTGGCTGCTGGGCGGAGACGCTCCTCACTTCCCAG
CTGGGGTGGCTGCCGGGCGGAGAGGCTCCTCACTTCTCAGACGGGGCAGCTGCCAGGCGGAGG
GGCTCCTCACTTCTCAGACGGGGTGGTTGCCAGGCAGAGGGTCTCCTCACTTCTCAGATGGGGCG
GCCGGGCAGAGACGCTCCTCACCTCCCAGACGGGGTCGTGGCCGGGCAGAGGTGCTCCCCACAT
CTCAGACGATGGGCGGCCGGGCAGAGACGCTCCTCACTTCCTAGATGTGATGGCGGCTGGGAAGA
GGCGCTCCTCACTTCCTAGATGGGATGGCGGCCGGGCAGAGATGCTCCTCACTTTCCAGACTGGG
CAGCCAGGCAGAGGGGCTCCTCACATCCCAGACGATGGGCGGCCAGGCAGAGACACTCCTCACTT
CCCAGACGGGGTGGCGGCCGGGCAGAGGCTGCAATCTCGGCACTTTGGGAGGCCAAGGCAGAC
GGCTGGGAGGTGGAGGTTGTAGCGAGCCGAGATCACGCCACTGCACTCCAGCCTGGGCACCATT
GAGCACTGAGTGAACGAGACTCCGTCTGCAATCCCGGCACCTCAGGAGGCCGAGGCTGGCGGAT

CACTTGTGGTTAGGGGCTGGAGACTGGCCTGGCCAACACAGCGAAACCCCGTCTCCACCAAAAAA
ATACGAAAACCAGTCAGGCGTGGCGGCGCACGCCTGCAATTGCAGGCACTCCGCAGGCTGAGGCA
GGAGAATCAGGCAGGGAGGTTGCAGTGAGCCGAGATGGCAGCAGTACAGTCCAGCTTCGGCTCC
GCATGAGAGGGAGACCGTGGAAAGAGAGGGAGACCGTGGGGAGAGGGAGAGGGAGAGGGAGAG
GGGGAGGGGTGTTTGTTTGTTTTTAAGACAAGGTCTTGCTCTGTTGCCTGTGTTGAAGTGCAGTGA
CTTGATCATAGCTCACTGCAGCTTCTAACTCCTGGGCGCAAGTCTTTTTTTTTTTTTTTTTTGAGACT
CTCTCTGTCACCCAGGCTGGAGTGCAGTGGTACGATCTCGGCTCACTACAATCCCTGCCTCCTGAG
TTCAAGAGATTCTCCTGTCTCAGCATCCTGAATAGCTGGGATTACAAGTGCACACCACCACACTCAG
CTAATTTTTGTATTTTTAGTAGAGATGGGGTTTCACCCTGTTGCCCAGGATAGTCTCAAATTCCTGAC
CTCAAGTGATCTGCCCACCTCGGCCTCCCAAAGTGCTGGGGTTACAGGCGTGAGCCACCACACCC
CACCTTCTCCCAAAGTCCCCACCTTCTAATACCATCACATTCAAGGTTAGGAATTCAACACAGGAATT
TCGGGGGAACACAAACATTCAGCCCATAACAGTTATGAACAATCGTGCCAATAAATTTCACAACTCG
GGTAAAATGGACACAGTCCTTGAAAAAGCCCACTGATGATGAAATAGATAGCCCAAATAGCCCTTTAT
CTATTAAATAAATTGAATTTGTAGTTTAAAAAAAAATTCAGACCTAGGTAGCATCACTGGTGAAATCTAC
CAGACATTTAAAGAAGAAATAATACCAATTCTACACAAACACTTCAAGAAAATTGAAGAAGGAATTCTT
CCAACTTGTTCTGTGGAGACCAGATTTACCCTGTGTTGATTACCCTGGTTAATTTTATGTGCCAACTT
AGCTAGGCCCAGATATTTGGCCTAATACATCAGTATGTTGTTGTGAAAGTATTTTTATAGATGAGATTAA
CATTTAAATTAGTAGACTTTAAGTAAGGTAGATTATTCTCCATAACGTGAGTGGACCTCATCCAATCAC
TTGAAAGCTTTAAGAGAAAACAGACTGAGGTCCTGTGAGGAAGAGGAAAATCTGTCTCTAGATTGTC
TATGGACAGTTGCAACAGCAACTCCCTTGGTCTCCAACCTGCCAAAATACCCTGCAGATTTTAGACA
TGTCAGCTACATGAGCCAACTTCTCTAGAGAAACAGAACCAATAGGGTACATACACAGTTGTCCCAT
CCCTCGGTGGGGGATTAGTTCCAAGACCTCCCCATGGCTACCAAAAGCCATGGATGCTCAAGTCCC
TTATATAAAATGATGTAATATTTGCATGTAACCTGCACACATCCTCCTATATACTTTAAACCATCTCTAGA
TTACTTGTAATACCTAATACAACATAAATGCCATATAAATAGTTATTGTACTATATTGTTTAGGGGATAATG
ACAAGAAAAATTCTGTATCTGTTCAGTACAAGTGCAGTCATCCTTTTTTTTTTCCAAATATTTTTGATCT
TCAGTTGGTTAAATCCACAAATGCAGAACCCATGGATTCAGAGAAACAGACAAACTGAACCAATGGG
AGATATATATATATATATATGTGTGTGTGTGTGTGTATATATATATATATATATATAAAATTAAAGTAGACAAA
GAAATAATAAAGATTACTATTCACAATAGCAAAGTGGGATACACGCACACAAACACACACACACACATA
CACACACACACACACACACACACACACACACATATATATATCACATTGGTTCTCTCTCTCTGGAGAAC
CCTAATATACTGATACCAAAAACAAAGACATTACAAGAAAAGAAAACTATAGACAAATCCTCTCATGAC
TTAGAGGCAAAAATTCTTAACAAAATTTTAATAAATTTAATCTAATAACATATCAAAAGAATAATACCAAAT
GACTAGAGTTTTTTTCAGGAATGAAAGGTTGGTTTAACATATCGTGCCATCAATGTAATGTACCATATT
AACAGACTAAAAAAGTAAAACCATATTATCATCTCAATGGATGCAGGGTAAGCACTTGACGAGACCCA
ACATCTATTTTTGATAAAAATTCTCAGCAAACTAGAAAGAAAAGAACTAGCTCAATCTGTTAAAGGACA
TCTAAGAAAAACCCACAGCTAATGTACTTAAAGATGAAAGACAATGCTTTCCTCCTAAGATCATGAAC
AAGACAAGGATGTCCTCTCTCCACTTCTATTCAACATTGTACTGGAGGTTCTAGCCAGTGCAATAAGC
CAAGAAACAGAAACAAAAAGCACCCAGATAGGAAAGAAAACAAACTGTTTTTGTTCACAGACAACAT
GATTGTCTATGTAGAATACTCAACGGTCTTACAAAAAGTTACTAGAATTAATAAGTGAGTGTAGCAGTT
TTGCTTTTTGTTTTGATTAATATTCAAAAATTAATTGTATTTTTATATACTAGTAATAAACAATTGGAAATTT
AAAAAATTTTAATACCATATATAGCCTGGTACAGTGGCTCACACCTATAGTCCCAGCACTTTGGGAGG
TCAAGGTGGGAGGATCGCTTCAGTACAGGAGTCCAAGACCAGCCTAGGCAACATAGTGAGATCCCG
TGTCTACACAAAAATAAAAATATTCAGGCATAGTGGCACATGTCTGTAGTGCCAGAGCTACTTGAGAA
GCTGAGGTGGGAGGATCACTTAAGCCCAGAAGTTCGAGGCTGCAGTGAGCTATAATCATGCCACTG
TACTTCAGCCTGAGTGACACAGCAAGACCTTGCCTCTAAAAAACCAAAAAAAAAAAACAAAGCACAT

ACAATAGCATCAAAAATATAAAATATTTCGGAATAAATCTGGTAAAATATATGTTCATGTATGAAAAGACT
CAATAGTGTTAAGATGTTAATTCTCCCCAAATTGATCTATAAATGCAATACAATCTTGATCAAAATCCCA
GCAGGCATTTTTGTAGTAACTGAGGAGCCAATTCAAATGGAAGTGCAAAGGACTTAGAGTAGCTAAA
ACAACTTCCTTTAAAAGTTTTGGGCCAGGCATGGTGGCTCACGCCTGTAATCCCAGCAGTTTGGGA
GGCCAAGGCTGGCAGATCACCTGAAGTCAGGGGTTTAAGACCAGCATGGCCAATGTGGTGGAACC
CTGTCTCTACTAAAAATACAAAAATTAGCCAGGTATGGTGGTGGGCACCTGTAATCCCAGCTACTCAG
GATGCTGAGGCAGGAGAATCACTTGAACCTGGGAGGCAGAGTTTGCAGTGATCCCAGATCAGGCC
ACTGCACTTCAGCCTGGGCAACAGAGTGAGACTCCATCAAAAAAAAAAAAAAGTTTCACCTGGCCAG
GGGTGATGGTTCATGACTGTAATCCCAGCATTTTAGGAGGCTGAGGAGGAAGGATCACTTCAGACTT
AGGTAGCAACACTGGTGAAATCTATCAAACCCAGGAACTCAAGACCAGTCTGGGTAACAAAGCAAG
ACTCTGTTTCTACCAAAAAATTTTTAAAAATTTGCCAGGTGTGGAGATCACTACTCCATCCTTGAACTC
CTGGGCTCAAACAATCCTCTGGCCTCAGCCTCCCGAGTAGCTAGGACTACAGGTGCACACCAGCAC
ACCTAGCTAGCATGAGTGTTTTTTGTTTTTGTTTTGTTTTGTTGTTTTTTTGTTTGTTTTTTTATCATGT
GACATAAGATTTATTGACTTCATATCAGCATTTAAGTATTGTTAACTTTTTTTTAATTTTATTATTATTATAC
TTTAAGTTTTAGGGTACATGTTTGTTTTGTTTTTTGAGACAGGGTCTCACTCTGTCGCCCAGGCTGGA
GTGCAGTGGTGAAATCATGGCCCACTGTAGCCTTGACCTCCCCAGGCTCAGGTGATCCTCCCACCT
CAAACTCCCAAGTATCTGGGACTACAGGTGCGCATCACCACACCAGCTAATTTTTTTGTGTATTTTTG
TAAAGATGGAGTTTCTCCAATATTGCCCAGGCTGGTCTCGAACTCCTGAGCTCAAGTGATCCACCCA
CCTCAGCCTCCTAAAGTACTGGGATTACAGGCATAAGCCACTGCACCTGGCCAGGCACCAGTGTTT
TTAAAGGTCCCAGAAGATTCCAATGTGCAGCCAATGTTGAGAACTGGGGAATGAGGGATGTAAATGA
ATCCGAGGAATACAAGGTATTATGAATGTGTGGAACGCGAGAAGGATCCAAAGTTTTGGTTAGATACA
AGCCATGGAAGGGATGCCCAAGGTAGAGAATCCAGTGGGAAAGCAGGATCTTGGGTTAGGACGAC
TGCAGGAGTAGGGATAAAAAGGATAGGGATAACTCAGAACTATGAATGAAACATGTGAATTAGACATT
TGTTCTGTCTCTCATGAAATTAGGGTCTGAATGTATCTTGAGAAGATGGTGGGCAGAATCATCCAATT
TTCTGGCTATGTGGTTACTAATTCTTTTCTCCTTTGGGAACCCAGGTCTATGAAATTCTCAGTCATTAA
ACTTTCCAGAGGCTCTTTAAATTATTTGTGTAATCAGAGAACAAAAGAAGGAATCCCCTGCTCTGGGA
TCCAGGAACAGGCATAAAAGACATAAGAGAGAAAGTGGCCTCTATTCTCCCTCTAAACAAGAGACTC
TGAATTTGGGCTGTTTCAGAATCAACTGGAAACACCCTGCCTAGAGTGCGTCTCTAGCACTTTTATTT
TATTTTATTTTATTTTATTTTATTTTATTTTGAGACCAAGTCTCGCTCTGTCACCAGGCTGGAGTGCGGT
GGCGCAATCTTGACTCACTGCAACCTCCGCCTCCCGGGTTCAAGCAATTCTCCTGCCTCAGCCTCC
CGAGTAGCTGGGATTACAGGCGTGCACCACCATGCCCAGCTAATTTTTGTATTTTTAGTAGAAACGG
GGTTTCACCATGTTGGCCAGGATGATCTCAATCTCCTGATCTTGTGATCTGCCCGCCTAAACCTCCC
AAAGTGCTGGGATTACAGGTGTGAGCCACCATGCCTGGCTGCATTTATATTTTTAATAAATCCCTCAG
GCCTCCAAAGTTTCAGAACCATAGCTATTTTTTTTTTTAAGATAGGGTCTCACTCTGTAGCCCAGGCTG
GAGTGCAGTGGCACAATCATAACTCACTGCAGCCTCCAATTCCCAGCCTCATGCAATCCTCCCACCT
CAGCCTCCCAAATAGCTAGGACTACAGGTGCCCACCACTGCACTGAGCTGATTTTGTTTTTTTTTTTG
GTCGAGACGGGATGTTGCTATGTTGCCCAGGCTGGTTTTGAACTCCTAGGCTCAAGTGATCCTCTT
GCCTCTGCCTCCTAAAGTGCTGGGATTACATGCATGAGCCATTGTGCCCAGCCTAGAACCACTACCA
TTAGAAGTAATAGAAGATGACATCTATTGAAAATATCCCTGTCTCAGCCAGGTGCAGTGGCTCATGCC
TCTAATCCCAGCACTTTTGGAGGCTGAGTGAGTGGATCATTTGAGGTCAAGAGTTCGAGACCAGCC
TGGCCAACATGATGAAACCCTGTCTCTACTAAAAATACAAAAATTAGCTGAACTTGGTGACACATGCT
TGCAATCCCAGCTACTCAGGAGGCTGAGGCAGGAGAATCACTTGACTCCAGGAGACAGAGGTTGC
AGTAAGCCAAGATCATGCCACAGCACTCCAGCCTGAGCAACAGAGCAAGACTCCATCTCAAAAAAA
AAAAAAATCCCTGTCTCAAACTCCTGCTTTCCAGGTAGATGGAGCGGGATAATGTGCTTTAATTGGTG

AAGGGTTTCTGATCCCCATTTATCATGCAGAAGGCAGGAGCAAGCCAGAGACAGGTCCTGGCGAAA
GGAAGCTAAAATGGTGCCTTTGATACAGGGGAAAGGGATACGAAAGGGGCTTAGGTGAGACCTCCT
TACAGCTAGTGGCACCTGGTCAGCAGGGCCGTAGAGTTGAGGCCATCCCCTTAGCAGCAGTGGGA
CTTTGTTACACACACCCTCTGAGTCTTGGTGAGAAACTAGTCCCAGGGCCCCTTGTTGTCTCTGCCA
CTCCCATTCATGACATCATGGAAACTTCTGTGCGGCTGGGCTCCTGAACACAGCAGCCTGCCCTCG
GCTGGATGAGGCAGCCAAAACCCTAAAAGGGTTGTGATAGGGACTGGAGTCAACAAGTAGCTGTGG
GAACGTGGTGGCGCAGACCCTTCACCCAGGGAGCCCTGTGGGCTAGAAACACCAGACAGCCAGG
ATGTGGCGAGAGACTGCCCAACACACTGTCTGTGTGAGGCAGAGGAAGGAAGGAGATGAAGGAAG
AAACAGGTAGGTAGGAGGGCGAGATCCCAAGCAAGTCAGGACAACTCCCACGCCTTGCCCAGGAG
CCTTAAGAAGCCCAGGCACCTGCTGAGTGAAAGAGGATATATTTATTGGCTGAGCAAGAAGGGAAG
GTACAGTTGGTAATGGCTGCTTCTAGAAGCCACCAGTCTCAGGTTCACTTGTTCCGAGCCCAGTTTC
CTCCAAGGTGGCTGTACTGAGCATCATCTCGATCTCGGAGGGGCTAAGAGAGGAGAAGAGAAAAC
GGTCAGGAGGCAGGGTTAGAACTCTTCAAGGAAGGGCCCCAGCAGGCCCTGACGATGAGAGCTC
CTGCCTGACCTCTCCAGTCACACCCAGCAATGAACTCCCCAGTAGGACCCTTCCCACGTGCAAACA
GCATTCAGTGTGAGCCTCTCTATCCCCACTTACCCTCCCTTCATTCCTGCCTCCTTCCCCTCAACGC
TCACCTGATAGACCTGGTCATTCCTCAACAGAGCTTGTGTGTCGGCAGCTAGAAGAACCAGAGAGA
GACATCAATGGCCTAGCAGATGGGACTGTGAGATCCACCCTCCCACACCCTCAGAAGTCTGCATGA
GTGATATGAACACACAAGTTCTTTCAACAGTCAAAGTTCTAGGAGTCTTTAGGAGATTGCAAGAGTAA
CTCCCAGCTGAGACTAAACCTACCACCAGCCCCATTCCTTAGCTGGTGCATAAGCTCACTGGTACAC
ACACACACACACACACACACACACACACAAACACACTCTCATGCTCTGCTCTTCCACTAACCCCCAG
ACAGCCTTCCAGTCTCATGTCCAGCAAAGCAGAAGACTCCCAAAGCAAGGAGCAGAGTGGCAATGA
CATCAGTGACAATGATGCCAGCCACGGTGGCTGGATCCAGCTCCACACAGCTCTGGCACACTGTG
GGGGAAGGGAGGAGAGAGGAGAGGTTGAGAGCCTTTAAGATCAGGGAACCATCCTCTGCCTCCTA
GGGCAACCCTTAGATCTCTTATGCCAAAACCCAAACTTCAATAAGACCCTGGGAGAAAGGCCTGGT
GATGGGCTTGCCACACCTTCCTCCACCCTGCATCCTAAGGAATCCCTGGGAAGAGCCAAGAAGTAA
TCTCAGCTTTTGGGACCTTGAACAAGGTGGTGGGCCAAACCTCTCACCCAACCAAGGCTGCAGTAA
CATGACCCCTACTGCTCTGTCTTGCTGAGTTAAGCTCTCTCTTCCACTTTGAAGGTCCCCAAATCTG
GCTTGTACCATTACAATAGTGACCTCACTTTGTTTAGAGAACAGATGGGTTCACCATATCCCTCTCTA
GCCAGAAAGTTCTCACATCCAGAAGCCCTATCCATTCCAACCCAAAGGGTTCAGGAAGCACGTACTT
CGATAATGAACTTGCACGGTAGATTCTTTGTCCTTGTATATATCTGTCCCATTACACCTATATATTCCTC
GTGGGTCCAGGATGCGTTTTCCCAGGTCCAGTCTTGTAATGTCTGAGAGCAGTGTTCCCACCGTTC
CCTCTACCCATGTGATGCTGGTATTGCAATTCACAAACACTCTGTCCTCAAGTTCCTCTATAGGTATCT
TGAAGGGGCTCACTAAAGGGGAAAAAATATCACAGTTGGAGACAGCTCTTTGATCTGCACCAAGCC
CTTTGTTCTGCGGAAGCTCATACTTAACAGAGACCATTTTCCTGGTCCAGGACAGTTTATGGCTTCCA
TCAAGAGAGACAGAAGTCACAAGAAAAAGCCTTCAGAAAGTTCCCCACCAACTGCAGGGGTCAAG
GGGGACATGAGGATGCCATTCAAGCAGAGGACAGGTCTTGGGGCCTTGGTGCAAAAGAGGACCCC
TCAGAGCAGGATTGACCCAAGCACCTTCCTGGAAATGAATCCAGACCACTGATGAGGAGTAGGGGG
AGCACGGACCACTGAAGCACCTGGAAGATGTGGAAAGACAGAAGAACATTCCTCGATTGGAAATGT
CTGCATTTTTTCTTCAAGGAAACATCTAATTCCACTTCCCAGCCATCTACAACACTCCCACTTCAGCT
TCTTATCCTCATCCTTCCTCATTGCCCCTGCTCCATTGACAACCAAGAAAGCGGGGCTCTCAACACT
GAAGCCTTTCCCAGGGCCAGGGATGGCTGTGGGTGGAGACCAGCTGGTTTACCAGCCCCTGAATT
ATCAGCCAAGTGGTCCAGAACGGGACCAGGGCAAATCCCATGTACAGTTTTCCACCCTTGGTTAGA
AGGAGGAGAACAGGAAAAAAATTTTATTGAATCCATCCCTAGAGCTCCTCACAAGTCAAGTCTTGTG
GGAGACTTTTAGGGCTGGAGGTGAGTGCAGCAACATTCCAGATGCAGTGAGTTCCTCTGACAGCCT

GAGCACATCTCCACAGGCCACAGAGGCACTACAGTCTATGCCTCCAAACACAGGGAAAAGTGGAG
GCTACATTCATTCATCCTGGGCTTCACACTAAGTCCCAAATTTGGATACAAGAGCATCTTCTAGAAAA
CCCTGAAACAGCTGTTGCTCACACTTCTGAAGCAGGTTGGAAGTATATGCATGTATCCTCAGGGAGA
CACATGCACATCAAATGCTTCACGTCCTACAGTCGCGTCCTCTTCAGGGATCTGTCTCCAGTGGAAA
TCCTGAGTGCCCTAGTGCAGCCAACTATTAGGTGACCATTGGACCCAGTTTGCTTAGTGTTGAAGGG
GTTCCTCGGACATGGGACTTTCCATTTTAAAACTGAAATTGGCAAACTGAGATGAGTTAAAATCCTAC
CATGTAACAACCCCTCAAATCTTCCCTCCGTCCTGCTCAACCTAAAGTTAACTTCTCTTAAAGCATTC
ACATAAGTGCTAGGACATGCCTCCAGGGATGACATAATCATGGCCAAACAAACAAGAGTCCTGATTC
CAGAGGCCATCAGGCCTAAAAGGAGTAGTGCAGGAAGCTGTGCTCCCATGGCCAGTCCCAGATTCA
GGTACATACGTACTGAGCTATTTTCTGCAGATCTCTGGCCTAAGGCCTTCTGAGAGACATTCTAGGC
CCACATGCACCCATGGCTGGAGTCAGTCAAAGCCAAGAGCCTGTTTCCCAGACTCTATGCTACATC
CTGCCCCTGCCCTCCTGACACCCCTGGGGTGCCTGGTGAACTGAAGCTAGCACCGAGAAGCACTT
TTTTTTTTTTTTGAGATAAGGTCTCACAGGTTGCCTAGGTTGGAGGGCAGTGGCATGATCACAGCTC
ACTGAAGCCTTGAAATCCCGGGCTCAAGTGACCCTCCTGCCTCAGCCTCTCAAATAGCTGGGACTA
CAGTTGTGCACCACCATGCCTGGCTAATTCTTTTGTTTTTTGTAAAGATAGAGTCTCATCATGTTGCT
CAGACTGGTCTCAAACTCCTGGCCTCAAAGGATCCTCCCACTTCGGCCTCCCAAAGCTCTGGGATT
ACTGGTGTGAGCCACCGTGCCTGGCAAGAAACACTTTCAAGTGGGCCTCACTCCCATCAGTAATGT
CCCTCTCAGGTCCCTTCCCCCACCCACCTGGAGTAGCCTTACCTTGCGAGAGAAGGGTAGCCAGTA
CCAGGCCAGAGAGAAACGTGCTATGTTCCATCTCCCAGCGGAACTCATCCAGTAGATAAAGCCAGG
TCACCGAACTATCAGCCTGGGTGAGAGCTGCCCTCCCCTAGCTGACTCACAGGTACCGGAAAGAG
GAGAGTGGGGGAGGAACTAGAAGATGTCTGCTTCTCTGCTTTCTGCCGTTGATGGTGGGAAATTTT
CAGAGTGGGGGTGGGCTAGGGGTAGGGTAGGAAGGAAGCGGTGTAAATTGCTCTATCTCTGAGCA
GGGAGCTGGCAGAGAATATGGAAAAGGTGGCCTGAACTTTAGCGTCCCTATTGACAATGCAACCAC
ATTTACACACAACCTAAACACTGCCACATCTCGAAGCCCCTTGAGAGAAGCCGTCGGCCCCATAGC
GCAAGCCGTAGCAGCTAGATTTCTCATGGAGGCTGATCTTTCTCAGGACCCTTCACTAGGCAGCCA
GGGACACCAGATCTAGCAGCTTCTTGTCAGTGGGAGGTTGGGCTTTAGAGACCCCAGCCAGAGATT
TGAATCCTGGGTCCAATACTGCCTACCTGTGGGGCCTGGGCCAGCCATAAAATTTTTCAGAGTCTTA
TTCCATTAGTACCATTATTAGGATTCAAACAAGATATTTGCATGGTGCCTCACGCATCATATGTGCTCAT
TAAGGGGTAGTTATTAATAATAATATAATTGACTGACAGGCAATATTGAGCCTCCCGGTGAGACAAATG
GACCTTTTTCCCCTGTGGCCTACGAGGATCTGAAACTCTTCACGCTGCTGCAGTTAGACTGTCACTT
ACCTGGGGACAGAGTCATGCCTGTCTTGCTCACTGCTGTATCTTGTGCCTGGCACATAACGGGAGC
TCTGCACATTTTTGTTGGCTCACTGACTGACTGGCTGAGGGAGATAGGGGCCTGAGATCCTGGACA
TTCAGTCCGGGCTCTGGCCCCTGAAAATGTGCTGGCCTGTCCTCGGAATTGTTCCACCTATTGCCT
TCCAGGCGCCTCTTTCATGATCTCAAAAGAATAGTGAAACCAGGTGCCGTGTCTCACGCCTGTAATC
CCAACACTTTGGGAGGCTGAGGCAGGTGGATCACAAGGTCAGGAGTTCGAGACCAGCCTGACCAA
CAAGGTGAAACCCCGTCTCTACTAAAAATACAAAAATTAGCCGGGCTTGCTGGCACGCACCTGTAAT
CTCAGCTACTCAGGAGGCTGAGGCAGGAGAATCGCTTGAACCCCGGAGGTGGAGGTTGCAGTGAG
CTGAGATAGCGCCACTGCACTCCAGCCTGGGCAACAAAGTGAGACTCTGTCTCAAAAAAGAAAAAG
AAAAAAAAAGTGAAAAAAATTCCTGAATGAAGGCCTGGACTGAGGTGGCTTTCCATTTGGAGGTCCA
GCCCCAAGCATCTGAGAGTCCCTCCTAAATTCATTACCTACAGCAACAACAACTAGCAACAAGTAAC
AACTGGCTACGATCCTAACAACTAATGACAGGGACATTTATCTCCCTCATGAAGAAACGGTCCCAGG
ACCATCTCCCACCCAGCATCCATTGCGGTTCCCTGTGCAAGATGAGTCTCTGAGTGGGAATCCAGC
ACTCTCTCCCTCTTCTTCCCCACCACCTTCACCCTCCTTAACGGAAAAACAAAAGGCATCTGCACCT
GCAGCCCTGCTGAGGCCCCTGCTGCTCACACTTGCAGCAGAGGGTGGAGGCTCTGGGTTCTTGC

CTTCTCTCAAAGGCCCCAGCCCCAACAGTGATGGGTGGAGCCAGTCTAGCTGCTGCACAGGCTGG
CTGGCTGGCTGGCTGCTAAGGGCTGCTCCACGCTTTTGCCGGAGGACAGAGACTGACATGGAACA
GGGGAAGGGCCTGGCTGTCCTCATCCTGGCTATCATTCTTCTTCAAGGTAAGGGCCTACTAGGGGT
CTGGAAGCCTGGGGAAGGGCTCAAGGGAAGAGCCCATCACTAGTGAGACAGGAATATTGGTATCCC
TAACCTTCAGCCTACCTCTGCTGTCACCTTAGAGTTCAAAGAAGGGCAAAATGGAGGCTCTTAACTG
TTCTCTGCTAGAGAGAAACAGTGTCCCATGGAGGAGAAGGAATCCTTGTCTCTGAAAAATGCAAACA
GAGTACTTAAATGGCTGAAGAGAGGACCCTGTTACCGCCATCTTAGATTTGAATGCAGCCCAAAAGG
GCATAGGCCAAGAAACTAAAAGGAAAAAGTATATGTTCCCTACTTCAGAGCTGGGGGCTAGCAGTCG
ACCTAGGAAATGTCCATTCACTCAGTTGGGCAGTTGGCCAATTGGTCAACAAAGATGGGTGAGACC
CTAATATGTGATATGCCATGAGGAAATAAAAATGAATAAAACACTTATGCTCTACAGAACTCATAGTCTC
ATTGGAGAGATTAGTTTTCATGGTGCCAGACTGAGTGTACAGTATAGTATGGGTGCCTGCATGAATAC
TCCATTGGACTATCCAATAGGAAGTGGATGAACTGGGTTGATGCCTGGAGTCTGGACATCCTGGGAA
AGTTAAAACCATGGAATTTGCTGAAGTCACACAAGTACTATGTGAGGAAGAGGGAGCTGAAGATGGA
ACAGTGAGGGCACCAATTTTAAGGGGCAGTTAGAGAAAAAAAAACATTCCAGGAAAGAGATAAAAGG
AGAATCAAAGAGCTGAAATAATCAAATGTATGCACTACCTTGGAAACCAAGAGAGTAGAGAGCTTCTA
GAAGAAGAGAATTACCAAGAGTATAAAATGTTAAAGAGAGCCCACTAATATAAGAATATATTTAAAAGTT
TAAGTCTGCGGGATGTGGCAGAATATGCATCATGGGTGATTTAGCACAGTTTCAGCAGGGTATTGTG
AGTGGAAGCCAAGTGAAAATTGGTAGAAGGTGAAGGTCAGGTGATGAGGTCAGCAAGTGTAGACTA
TTCTTTAAAGACATTTGGATGAGGAGGGAGAAAAGACTGGGTGGTAGGTAGAGGAGGAAGTAGAGT
CTAAGAATAAATGTTTTTGAGGGTGAGAGAAACTTGAGCACATTTCTAGGCTAAAGGAGAAGATGATG
AGTAGAGCAAGGTCCTGGAGTTGATGGTGGAAGGACAGAGTCAAGAGTGGGGTCAAGAGCACAGA
GAGGAAGGCAACGTTACCCTCTAAGACCAGGAGAAAGGAAGTGCAAACGGCACTGGCCACAAATA
GGTTTATAAAGGGAAAGGAAGCTAAAGGAGAAAGGACAATAACAACTTCCGTGTTCTAAAAAATCCA
GGTGGGAGGCAATGCCACCTGCTGAAACTGAGCTGGGTGATGACCGGTTGGGAGACTTGAAGAGG
ACAGTGAGAGTTTAGAATAGAATTATTTCTGAGAGAAATGCAAGGAAAAGTCAGGTAAAGACAACACA
GGTTGAGGAGCAACTCTCAAGCCTGGGTGAGGTGGCTGCATATCATGGAAAGAGCATTTGAGTTCA
GCAGTAGAAAACCTGAGCTCTGGATCCAGCTTGGACACCAATGAGCTATGTGTACTTGAACAACTCC
TCCACTGTCTGGGAGTTCTCATGCCAGCCAATCCAGAAATGTGGCAGGGACGGGCATGGTGGCTCA
TGCCTGTAATCCCAGCACTTTGGGAGGTCGAGGCAGGTGGATCACCTGAGGTCAGGAGTTCGAGA
CCATCCTGGTCAACATGGTGAAACTCTATCTCTACTAAAAATATTTTAAAAATTAGTGGGGTGTGGTG
GTGGGTGCCTGTAATCCCAGCTACTCAGGAGGCTGAGGCAGGAGAATTGCTTGAACCCAGGAGAT
GGAGGTTGCAGTGAGCCAACACGGTGCCACTGCACTCCAGCCTGGGCGACAGAGTGAGACTCCG
TCTCAAAAAACAAAACAAAAAAAAAAGAAATGTGGCTATATGAACTCTTAAGTCCTAAAAGGAAACCT
CACTCAGATATGCAACATTAAAAGATGTCAATCAGCCAGGCACAGTGGCTCATGGCTATAATCCCAAC
ACTTTGGGAGGCTAGGGAGGGAGGATCATTTGAGCCCAGGAGTTCAAGACCAACCTGGGCAAAATA
GGGATACCCTGTGTCTACAAAAACTACGAAAATTAGCCAGGCATGTTGGCATGCACTTATGCCAGCT
ACTTGGGAGGCTGAAGTGGGAGGATTGCTTGAGCCCAGGAGCTTGAGGCTGCAGTGAGCCATGAT
CACACCACTGCACTTCAGCCTGAGTGACAGAGCAAGAAAAAAAAAAAAAAGATGTCTCAAAAAAAAAA
AAAAAAACTATGTCAATGATCAGTGCAATCTCCTTCCCTGAATCCAATTTAGAATGAGGGGTCTCTGT
CTTCTCTCGGTCTTCTGACCAGGTGGTCAGGAGAAAAACTAGTTAGAAACCTCTTTAACTGTTTGTC
TTGCCTTTTCTCCCATTTGCTCATTTATTCATTTGCACATTCATTCAACAAATAAAGTTCCTACCATGTT
CCAGACACTGCACTGGATACTGTTCAAAAATAATCCCGCCCTGCATGAGGTTCTAAGACAGTTCACT
TCTTGTATTTCCATTATACTTTGACTCATGACCGCCTCTCCCTATCACACACACTCCACCTCTAGCCAC
ACTACCTCTCTTGTCCTTTTCCAAAGCTGCATCTTATTCAGCTATTCTTTCTACTCAGAATGACTTTCT

CTATCTGGTATATTCTTGCTTACTCTTCAAGGCCCAGACAAATGTCGTTTCCTCTATTAAGCTTTCCCT
GATCCTCACCCTTGGACAAAATTAATAGCTTCTACCCTTTTTTCATATGCAGTCATATAAGCACTAAAA
GATAATGTATACATTCTATTAAATTAATTGCATATATGTTTTAATTTTTTCACTTCATTCATAAACTTTGGA
AGCAGCAGTTGATTCTTCATATCAGCACTCTTTTCTTTCATAATATCTAATTATTTTAGGTCATAATAGTC
CTCGTTAAATGTTTGGATTAAATCTAATTGAATTGAATTCAAGTGCCAGATCTCTGTAATATGTCTACCA
TGCAACTCTACTACCCTAAGGTTTTGTTTGTCTGTTTGTTTGTTTTTATTTTTTTGAGACAGCTTTAACA
CCTAGGCTGGAGTGCAGTGGTGCAATCTTGGCTCACTGCAACCTCCACCTCCCAGGCTCAACCATT
CCTCCCACCTCCTGAGTAGCTGGGACTACAGGTGCATACCACCATGCCCGGCTAATTTTTTTGTATT
GTCTGTAGAGACAGGGTTTCGCCATGTTGTCCAGGCTGGTGTCTAACTCCTGGGCTCAGGCAATCT
GCCCACCTCAGCATCCCAAAGTGCTGGGATTACAGGTGTGAGCCACTGCTGCCAGTCAACTCTGAG
TTATGAGACAGATTTGAGGACTAATTGAAAAGCTAACTTCCCTGGGACATCCAGGGTTTTTATAATAAA
AGGATCACCAAGGCTGAATATTTTATAAGGAACTCAGCCTAAAGGTTTTGGATAGTGCACACCCACTC
CTTGCCAGGGCATTAGCTGCTCAAGAAGCAGAGTGTTCTGGACTGGATAGGCAATATCCTTACTTAC
ATATACTACAATACAATTCTGATGCTAACCCCCCAAATTAGTGTCAGACCCTACAGGTTAAGAGCATGG
TTCCCAACCAGATTGCCCTCACTTTAAATGCCAGCTACAAGTTCAGGGTACCCCAGACCACTCACAT
TTCTGACAAACTGCCTGCAATCTTAAGGATTCCCATGACCCTCCTCATTGATAATTTGCTAGAATAACT
CACAGAATTTAAGAAAGTGCTGTACTTCCTATTTCAGTTTTATTAAAAAATAAAATAAAATCAGAACCAG
CTAAATGAAAAGGGCAAGGTCTAGGAGGGTCCCGAACACAGAGCCTCCGTGCCCTCTCCCCGTGG
AATTAGAACACATCGCCCCCAGCACTCTCCCACACTCTGCCAGCACATTGATGGGTTCACCAACCA
GGAAGCTCCACCAAGTTTTTATTGAGGTATCTTTACATAGGCATGATTGATTGAATCATTGGCCGCTT
GACTGATCTCAATCTCTAGGATCCCTTCCTGGGGCTGATACCACTAGTTTCAAAGCTGCAATCCTCTT
ACCATATGGTTGGTCTTGACCAGCACCATCCTGAGTCATTGCCATGCATAAACTCAGGTATGGTCTAA
GGATCCACCACAGATAACAAAGCCACTCCTGTCACTCATGAAATTCCAACGGTTAGAAACACCCTCC
CAGGATACCAGGACAAAGATGAGACAAATTGTTTATTATCACCTTGAATTCAATTGAGTGATTTAGTCT
ACAATCCGGAAAACTAAGTATAGATACTACCATTTTCATGGATTTGGATCTTTCTTCATCTTGGCCTCA
AATAACCATGGAAATACTTCAGGGCATCTGAACAACTCCATGCCCAGCTAATACTCTATCTCTCTTCT
GTCTTTACAGGTACTTTGGCCCAGTCAATCAAAGGTAGGAGAAATGGCTTCTTTCTATACTCAGACTC
AGAATATTGACGGAAATTTGGCTTCCTACAACAGTAGTCCTACAGGAGCGAACAGTTTAGAATGAATG
AAATGACGGGGATAGAGAGGTGATGTCTCTATTGTCAACCAAATCAGTGACCTGACATAACCTGTTC
CGGGCAGCTTGCCTGTAGCTAAGCATTTAACTGGTCTCTTACAGGAGAAGCAGGACCCTAGTAGCTA
GGGACACATCTCAAACTGTGACCCATGAACCAGTAGGTATTGGCGTCACCTGGGATCCTGACGTTA
GTCTCTGTCAATCCTTCTCTTTAGTTCATCTATTCTACCCAAAGTGATCTCATCATCTGGTATGCTGTT
AGCAGTTTCTTACCTGTATAGTATCTTCCAAATAACATGCCCCAAAATCCCAAAGTTTTACCCCTACTA
ATTACAGCAATGTCTCTTTTATTCTTCACCCCCTGACGCAGATATTGGCGTCACCCGAGAGCATGTTA
GTAATGCAGAATCTCCCCTCCCCAGAACTACTAAATAGCACCTGAAATTTTAACAAGATCCCCATGTG
ATTCATGTGCACATCAAAGTTTGAGAAACACTACTCTAATGATCTCCTGGTATGCAGAAGCAGGGAGA
ATTTCAGAGGCAAGATCCTTAATAGAACCACGGCTTTTCTCATTTCAGGAAACCACTTGGTTAAGGTG
TATGACTATCAAGAAGATGGTTCGGTACTTCTGACTTGTGATGCAGAAGCCAAAAATATCACATGGTT
TAAAGATGGGAAGATGATCGGCTTCCTAACTGAAGATAAAAAAAAATGGAATCTGGGAAGTAATGCCA
AGGACCCTCGAGGGATGTATCAGTGTAAAGGATCACAGAACAAGTCAAAACCACTCCAAGTGTATTA
CAGAAGTATGTAATCCCCTTTGGTCTGTTTGTTGTGAAATTAATCAGTATTTGCTGTTCTGGTGAGCTT
TTTATCTGGGGTGAAAGTGGAAATAGATCCTCAACAGTAATATTATCGCCTGTTCTCTTAATTTCAGCT
TGCCTCTTTTAAAATACTGTAAGATACTTCCCTCACCCTATTGAAAAACTACAGCCAGTCCTGTAAAAT
TTTGTTTACCTTTGGGTGGGCTCCATGGATTCAAGCAATTCAGCACTGAGTTGAATGAAGGGGTTGG

GGAGACAGAGGTGATCATAAGGTGAGCAGCAGTGAGAACTGGAGCGCAGTGGAGCATGAAGAGTA
TCTAGTCTTCCTTCTGTTCCAGATACTTCCTGGTAGTTAGACTACATGGGCTTCCCCAGGAATCCTGG
GGGACTTAAGAGCATCAAGATGCATTGAGTTTTGGCGCAAGGATCTTCCCTTGCCCTGCCACCCAC
AGAGGAAAAGCCTGCTGCCCTCCACAGCGGCATTATTGCAGACAGGCAGGAGAAAACGAACCAGG
AAAAACAACTTTCGCAACCTGAAGGTTTGTCTCTCCTTTTCCCTACAGTGTGTCAGAACTGCATTGA
ACTAAATGCAGCCACCATATCTGGCTTTCTCTTTGCTGAAATCGTCAGCATTTTCGTCCTTGCTGTTG
GGGTCTACTTCATTGCTGGACAGGATGGAGTTCGCCAGTCGAGAGGTAAAAGAATGCTCTTAGATG
AGAGATGGGACCACCTGAGACCCTCAGCTTTCCTCCTACCATTATGTACCCAATGGAAGAGACTGAG
TTGGTGCTTCTTGCTAGTGTGCATAGTTGGGTGAGGCTGTATTTCTCTGAGACAGGAGAAAGGATAC
TTGGTGTTATCACAGCATGTTCACCTGTCTCAAGATACAGCTCCCTCTGTGAAAAAAAAAAAAAACAA
AAACAGGCGCAGTGGCTCACGCCTGTAATCACAACACTTTGGGAGGCCGAGGCGGGCAGATCACG
AGGTCAGGAGATCGAGACCATCCTGGCTAGCACGGTGAAACCCCGTCTTTACTAAAAATACAAAAAA
TTAGCCGGGCGTGGTGGCGGGTGCCTGTAGTCCCAGCTACTCTGGAGGCTGAGGCAGGAGAATG
GCATGAACCCGGGAGGCGGAGCTTGCGTGAGCCGAGATCGTGCCACTGCACTCCAGCCTGGGCG
ACAGAGCAAGACTCCGTCTCAAAAAAAAAAAAAAAAAAAAACTATTTTAAGTAAACCCTTACATACAGA
AAAGTGAATCACAAGCATACAGCTCAAAAAAAAAATTCACAGAGCAAATACACCCATGCAGCCAGCA
CATAATTTAAGAAAGACAATATTACCAATGCCATTGAAGCCCCTCTTGTGGTCACTTCCAGCCACTAC
CCATCCTCAAGAGTGAGCATTGTCCTGACTTCTAACAACAAGGATTAACTTTGCTTGTTTTTGTCCTT
TATGTAAATGGACTCATATACTAAGTATTCTTGTGAGTAGCTTCTTTCACTCAACAACATGTTTATGTGC
TTTATCCATACTCTTGTGTATATATAAAACTCTTATTGTTATTTAGTATTCCATTTTGTGAATATAACCCAA
TTTATTATGCATTCTACCTTGTGTGGATTCTGTTTCTTTTTTGTGCAGCTTCAGACAAGCAGACTCTGT
TGCCCAATGACCAGCTCTACCAGGTAAGGGGATGAAGAATAAAAGAGACATTGCTGTAATTAGTGGG
GGTAAATCTTTGGGATTGAGGGGCATGTTATTTGGAAGATCCTATACAGGTAAGAAACTGCTAACAGC
ATACCAGATGATGAGATCAGTTTGGGTAGAATAAATGAACTAAAGAGAAGGATTGACAGAGACTGAG
GTCAGGATCTAGTGAGCACAAGTTGAAGAACACACTGAGAGGGACACACGAAGAAACTCTCGACAG
GCTGGGCACTCACTATAGACAGGCCATGGCTCATGCCTGTAATCCCAGCACTTTGGGAAGCCAAAA
CAGGTGAATCACCTGAGGTCAGGAATTCGAGACAAGCCTGGCCAACATGATGAAACTCTGTCTCTA
CTAAAAATACAAAAAAAATTAACCAGGCGTGGAGGCGCGCGCCTGTAGTCCCAGCTACTCAGGAGG
CTGAGGCAGGAGGACCACTTGAACCCAGGAGGTCGAGGTTGCAGTGAGCTGCGATTGTGCCACTG
CACTCCAGCCTGGGCAACAGAGAAAGACTCCGTCTCAAAAAAAAAAAAGAGAGAGAGAGAAAAAGA
AAAAAGACAGAGCCTCCATCTCCTTGTCCTCTTTCCATCCTCAGGACCATGAAGTACCCACTCCAAA
TTCTCACATATAAAAAACATTCAATAAACATGCATCAAATTAATTAATAGAGGATGGAAAAAATGACTTAT
GACTGTGCTGTCCTTTCCAGCCCCTCAAGGATCGAGAAGATGACCAGTACAGCCACCTTCAAGGAA
ACCAGTTGAGGAGGAATTGAACTCAGGACTCAGAGTAGGTGGGTTCTTCAATGCCAATTCTAATAAA
GGACCCTTGCATCAACTGCCCTCGCAATTGCTTCTAAGTCTAGCTCCCTTCCCTAAGCGGCTATAAG
CATCAGACTCTGGGGATCAGGGATTGGGACGTGGTTTGGGGTACTCTTTTCTAAAAATTCTGGGGC
CATACTGATTGTCTTGGCCTAGGTAAATATGAATTTTATGTATCTGTAAATCCTGTCAGAGCAGGGCCT
CAAGCCATAGAGATGCTGAATATTAATCTTAACCTACATTTGAATTTCTCATTATCTACACTATTAACATT
TTGGGCTAATTAATTATTTGTGATGAGGGGCTAGCCTGTGCATTGTAGGAGTTATGGAAGCATCCCTG
GCCTCTCTCCACCAGATGCTGGTAGATTGTCCAGTGTGACAATCAAAAATGTGTCCAGACATTACCA
AATGTGTCCAAACATCACCTCCAGGGCAAAATCACCCTTAGTTAAGAACCACTAACCCATATTAACCT
TCCAATCAATAAATCAATCAGTCAGAAGTTATGATTTAATTAATCTATCTGAAGTTTCTATCAGGAAGAC
AGGGTTGAAAGCATTATTTGTTTTTTTTGAACAAATTGCAATTTTTCTTTTTTCAGTCCAGGTGTTCTC
CTCCTATTCAGTTCCCAGAATCAAAGCAATGCATTTTGGAAAGCTCCTAGCAGAGAGACTTTCAGCC

CTAAATCTAGACTCAAGGTTCCCAGAGATGACAAATGGAGAAGAAAGGCCATCAGAGCAAATTTGGG
GGTTTCTCAAATAAAATAAAAATAAAAACAAATACTGTGTTTCAGAAGCGCCACCTATTGGGGAAAATT
GTAAAAGAAAAATGAAAAGATCAAATAACCCCCTGGATTTGAATATAATTTTTTGTGTTGTAATTTTTAT
TTCGTTTTTGTATAGGTTATAATTCACATGGCTCAAATATTCAGTGAAAGCTCTCCCTCCACCGCCATC
CCCTGCTACCCAGTGACCCTGTTGCCCTCTTCAGAGACAAATTAGTTTCTCTTTTTTTTTTTTTTTTT
TTTTTTTTGAGACAGTCTGGCTCTGTCACCCAGGCTGAAATGCAGTGGCACCATCTCGGCTCACTG
CAACCTCTGCCTCCTGGGTTCAAGCGATTCTCCTGCCTCAGCCTCCCGGGCAGCTGGGATTACAG
GCACACACTACCACACCTGGCTAATTTTTGTATTTTTAGTAGAGACAGGGTTTTGCTCTGTTGGCCAA
GCTGGTCTCGAACTCCTGACCTCAAGTGATCCGCCCGCCTCAGCCTCCCAAAGTGCTGGGATTACA
GGTGTGAGCCACCATGCCTGGTCTTAAAACCAGTTTCTTATATATCTCTCTGGAGGTATTCTAGGCAT
ATATGAGCACATTCTCAAGTACATATTATCCTCCCTTCCCCTATCTTTTAGACAAATGATATCAAACTATA
CATCTTGTGAGATTATTGCATACCATTATATGAAGATACCATTATATCCTTTTTAATGCAACCATATTGTA
CAAATAGACTATGATTTATTTAACCTGTTATCTATCAGTGGATATTTAAGTTGGTAGTTGGTTCCAATCT
TTTGCTCTTACAACAATTCTGCAATGACTAACATTGTATAAATATCATTTTTAAAAATAATTGCATTGAAG
CATAATGTACATGCCATAAAATCCACCCATCTTAAGTGATTTCACCTGTTCTCAGAAATTTTTAGTAAAT
TTAACTAATTGTACAGCCATTACCATAATCCAGCTTTAGGACATTTTCTTTTTTTTCTTTTCTTTTCTTTT
TTTTCTTTTTTTTTTTTTTTTTGAAGTGGAATCTTGCTCTGTGGCCCAGGCTGGAGTGCAGTGGCGCG
ATCTCAGCTCACTGCAACCTCCACCTCCTGGGTTCAAGCGATTCTCTTGCCTTGGCCTCCCGAGTA
GCTGAGACTACAGGCACATGCCACCACGCCCAGCTCATTTTTTGTGTATTTAGTATTTGTGTATCTAG
TATTTGTGTACTTAGTAGAGACAGGGTTTCACCATGTTGGCCAGGCTGGTCTCCAATTCCTGACCTC
AGGCGATCCACCCGCCTTGACCTCCCAAAGTGCTGGGATTACAGGTGTGAGCCACCGCGCCAGGC
CCGTAACTGTATTTTAATATAGCCATTCTATGGATTTAATATGGTATTTTATTATGGCCTTAATTTGCATTT
CCCTAGATACTAACCATGCTGAGTGTCCTGTCTTGTGTTTATTAACCATTCATATATTTTTAGTGAAATG
TGTATCAAATCTTTTGCCCATTTTTAAGTTGACTTATTTGTTTGTCTTCTTACTATTGGGTTGCATATGTT
TTTGATATAAGTCCTTTATCAGATATATGATTTGGAAATATTTTCTACCAATCTGTGGTTTGTTTTTCTTA
ATGGTGTCTTTTGAAGTGCAAAAGGTTTGAATTTTGAAGTACATTTTATTGATTTTTTCTTCTATATATTG
TGCTTTTGGTATCATGTCTAATAAATCTTTACCAAACCCACAGTTACAAAGATTTTCTCCTGTCTTCTTT
TTATACTTTTTACAGCTTTATGGTTTTAGCTCTAACAATAAATGTGATTTTGAACATACATAAGACTATTT
GTAACAAACACAAATAAATTGAATTGTTGGGCACTTGTATTTGCATTTGTAATTTTATTAACTATTGCCA
AATAGCAGTTGTAAACACGTACACACACACAAGTGTATATACACATATACATATATATACATATACATGTA
TGCATATATGCATATATACATATATGCATATATACACATATATACATATATACACATATACATATATGCATATAT
ACACATATATACACATATACATATATGCATATATACACATATATACATATATACACATATACATATATGTATATA
TACACATATACGTATATGTGTATATATACACATATACGTATATGTATATATACATATATGTATATGTATATATAC
ACATATACACATGTATATATACACATATACACATGTATATATACACATATGTATATATACACATATACACATAT
GTATATATACACATATACACATATATACACATATGTATATATACACATATACATATATGTATATATACACATAT
GTATATATACACATATACATATATGTATATATACACATATACATATATGTATATATACACATATACATATATGTA
TATATACACATATACACATATACACATACATATATGTGTATCTACACATATACACATATACACATATACGTAT
ACGTGTATATACGTAGATACACATATACATATATACACACGTGTGTATATACGTATATACACAAGTGTGTA
TATACGTATACACACATATATGTGTGTGTATACGCATACACACATATATGTGTGTGTATACGCATACACA
CATATACGTATATATGACAGGGTCTTGCTCTGTTGCTAAGGCTGGAGTGCAGTGGCACAACATTGGC
TCACTGCAGCCTCGACTGCCCAGGCTCAAGCAATCCTCCCACCTCAGCCTCCTGAGTAGCTGAGAT
TTCAGGCGTGTGCCACAATATCCAGCTAATTGCTTAATTTTTTTTATAGAGAGAGGGTCTCACTATGTT
GCCCAGGCCAGTCTCAAACTCTTGGGGTCAAGTGATCCTCCCATCTCATCCTCCCAAAGTGCTGGG
ATTACAGGTGTGAGCCACTGTGCCCAGCCAGGTTGTACCAATATACACTACCAATAACAACAAACAAA

AGTGCCTGTTACTCCATACATGTTCCAACAGTGTATTATTAAACTTTTGAATCATGCCAAACTGAATGG
CTTCTCACTATAATTTTAGTTTACATTTTTCTTATTAAAAATTGGAGGCTAGGTGTGGTGGCTCATACCT
GTAATCCCAACACTTCGGGAGGCCAAGGCAGGTGGATCACCTGAGGCCAGTAGTTTGAGACCAGC
CCAGCCAACATGGCAAAACCCTGTCTCTACTAAAAATACAAAAAATTAGCTGGGTGTGGTGGTGGGT
GCCTGTAAACCCAGCTACTCAGGAGGCTGAGGCAGGAGAATTGCTTGAACCTGGGAGGTGGAGGC
TGCAGTGAGCTAAGATTGCACCATTGCACTCTAGCCTGGACAACAAGAGTGAAACTCCGTCTCAAAA
AAAAAAAAAAAAAAAAAAAAACTGGAGTAAACCATTTGTTTTTCCTTCTCATGATACATAAGTTCACATC
TTTGGTCTTTTTCTTATTGACTTACACTAACCCCTTACATATTTGGGAAACTAGGTCTTTTGTGATATTA
GTTGCAAGTCTTTTTCCAACTTTATGTGTTTTTTAACTTTGCTTTAAAAAACACATTGGTTAGTTTTTCA
CGCACACTTTACAATTATTTGCATGTAGTATAATTCACTGGTCTCCTATGGTCTCTAGGTTACGTTTCAT
ATTTAGAAAGTCTTTCTCCATTTTAAAATTATCCTTTTTTAAAAGTTCCCCCATGAATTCCTCTAGCATTT
TTATGGTTTTATTTTATGCATTTATATCTTTGACCCATCTCGAATTGATTTTAGCATCAGATCTAATGTAG
TAACCAACTTTAAATACAGATAGTTACCCAGCTATTCCAGAAATATTATTGAATGATTCATTTTGCGTCC
ATTAATTTTAAATGCTGTTACAACCCCACTCGCCAGATTATAAGCTCCAAGAAGGAAAACACTTACGTT
TTTTCACTATGGTATCCTTAGGAACTAGAACAGTGGTAGGCACACAAAGATTGCCCAGGGCTTTTTGT
GTGTGTGTGGTAAAAGATATGTAACCTAAAATTTACCATTTTAACCTTTTTTTTTTTTTTTTTTGAGACA
GAGTCTCACTCTGTCATCCAGGCACTCTAGCGCAGTGGCACGATCTCGGCTCACTGCAACCTCTGC
CTCCCAAGTTCAAGCGATTCTCCTGCCTCAGCCTCCCAAGTAGCTGAGATTACAGGCGCACATCCA
CATGATCAGCTAATTTTTGTATTCTTAGTAGAGACAGGGTTTCACCATGTTGACCAGGCTGGTCTTGA
ACTCCTGACTTCAAGTGATCTGTCCACCTCGGCTTCCCATAGTGCTGGAATTACAGGCATGAGCCAC
TGCACCCAACCCCATTTTAACTATTTTTAAGTGTACCATTCAATGGCATTAAATATATCCACAATGTGTA
GCCATCATCACTACCCATTTCAGAAATTTTTCATCCTCCTAAACAGAAACTTGGTACCCATTAGACAAT
AACTCCCATTCGCCTCTCCCTCCAATCCCTGGTGACCTCTATTATACTTTCTCTCTCAAGGAATGTGC
CTATTCTGTGTACCACATATAAACTGAATCATACACTATTTAGTCCTTTCTGTCTGGCTCGTTTCAGTTA
GCATAACGTTTTCAAGGTTCATCAATGTTTCAGCGTGTATCAGAACTTCATTCTTTTTCAAGGCTGAAT
AAGTCATCTGTAGCTATATACCATATTTTGTTTATGCATTCATCTGTTAATGAACATTCTGGGTTATTTCT
ACCTTTTGGTTATTATGAATAGTGCTACTGTAAATATATGTATACAAGTATCTGAGTTCCTGCTTTCAGTT
ATTTTGAACATCTATATCAAAGCGGAATTGCTTGATCATGTGGTTATACTATGTTTAACTTTTTGAGGAA
TCATCAAACTGTTTTTCCAAAAAGTCTGGACTATTTAACATTGCCATCAGCAACGCATACGGTTTCAAT
TTCTCCACATCTGCCAACAATTACTTTCCTTTTTTTTTATAGTAGCCATCCTAGTAGGTATTAAGTGGTA
TCTCATGATTTTTTTTTTTCATTTCCCTAAAGACTAGTGGTGTTGAGCACTCTTTCATATGCTTATCCACC
ATTTGTACAGTTGGCCTTCCATATCTGTGGGCTCTGCATCTGTGGCTTCAACCAACCATGAATTGGAA
ATATTCCAAAACAAATTTCATCTGTATTGAACATGTACAGACTTTTTGTCATTATTCCCTAAACAATACA
ACACTATTTAGATTTATGTTGTATTAGGTATTATAAGTGATCTAGAGATGACTTAGTCATCTCTAGGATG
ACACAGGAGGATGTGCATAGGTTATATGCAAATACTGCATCATTTTATGTAAGGAACTTGAGCATCCTT
GGATTTTGGTACCCAAGGGGGTCCTGAAACAAATCCCCCACAGATACCAAGTGGATAGTCATTCTCT
TGGAATGTATCTTCTTTGGAGAAATGTCTATTCAAGTCCTTTGCCCTTTTTTGAACTGAACTGTTTGG
GGGTTTTTCGTTCTTGTAGTTCTTTACATATTCTAGAGATGAATCCTTTATCAGATATATGAGTTGCAAC
TATTTTCTCTCAATCTGTGAATTTTCACTTGCACAGCGCTTTTAACTCTTGAAAATTATGCTGCTTCTAT
TATTTATGTGTATCCCGATCTCTTGCCCTCGTGACCCGCCCGTCTCAGCCTCCCAAAGTGCTGGGAC
TACAAGGCGTGAGCCACTGTGCCTGGCCATCTATGTGTATCTCAAAACAACCTTGTAGGATGTATACT
GTCTCCACTTTACAGATAAGAAAACTGAAATCCAATCAAAGATAGCACACTCCAAAGATTGCTCCAAA
GACAGCACAACAATAACAAACTTGTAGTTGTTATTGCTGTTAGTGTTATCCCTTTGAATAATAGAGAGT
AACAGAAGCTGCTTCAAAAAGGTCTAGGACTGGTGGGAAAATAAGAGAGCAGACCCCCAACAGAG

GTTGCGAGCAGAGCCAGGGATCACAGAGTTTTATCTGCCCTCGGTACGCTGATTTCCAAAACCCAG
CCTCATATTCTATACTCCAAAGCGCACTGCCAGGTGGGCCAACTCCAGCCCCCACAATCCGATGCCA
AGGCCACTTCTTGCCACTTCCTGCCCGCCTCCAGCCCGCCCCTAACAAGCTAGCGCATGCGCTGTA
GGACTATTTGCGCGCCGTTCCCCCCGGTGAGCCTGCACACCGTGCTTCCGGCGCGGAGATGAACT
TCCACATGCCCCGGCGCTTGTTGATGACGTAACTTCCGGTTGCTGTGCTGAGTCGGAAGTGGGAAC
CCTTCGGCCGCTGAGATTCTGTCGTGTCGTCGCTGCTGGCACTTCAGGCTCTGGTAAGACAGGCAA
TAGCTTCAGGGGACGGGGTCGTTGACTTTGCGTTGTGATATTTTCTGCTGGTCCAATGCCACTTCAA
GCCTGTTCACTCTTATCTTCTCCAAATTCCTTCTCCTATGCTTTGCTAACCAGGGACCTTCTCCTTCTT
GAACTTCGACCCTGTCTGATTTGGAACCCCTTACAGTGCCCCAGCTTCTGATGTCTCTTTGTACATTT
CTGTCCCCATCTTTGCCAAGCCCCTCCTGGAAGAGGCCGGAGCGCTATCATTGCATTATCTTTTGAC
TTAACTTTCTCTAGAACGTGCCCCGGGCGGGTGGATCTCCCTCCTTCCCGCGAATCCCTTATGGAG
CATGGAAGGATAAAATATAAAGAGTAGATAACAAATAGGCAGATGCCAAGAAGATAAGATTAATCGTGA
GTCGTAAGCAAACCAGTTCAATGCATACGTCAGGGTGTTGATAATTCTTAAGGCAGGTGATGAGTTC
ACCACAGATAACTCTTGGGGCACTCTGCTTCATGTTTCCTTGGGTTTAGAAGACACTTCTAAAATTGA
GTGACCATGCCTCATGACCTTATATTTGATGAGCCTTCCTTTTAGGATCAGTTTTATGCCCTGACAGC
AAGCATGTGTTTAGTCTCGCTGTCATTTACTGTTTTGGTCTCACGTGGAAGCCCACCTGCGATAAAAT
GGGATCCAAGAAGGACTGCTGTCTGCTTTTTGACTTGAGATCCTTCTAGGCTGCAGATAGATGTGTT
GATGGAATTTGTTGCCACATTAGCATGGCCTGATAGTGTTAGGTGAAACATACGACAACGAAAATTGT
TTCTTAATTTCTATTTTAAATTGATATTTTAATTTCTATTTTATATTGATATATGTATTATTTGCCAGGACAC
AGACCTTATGGGAGTATGATTTGATAGAGCTTGATTTTGACGATCCCAGTTTTACAAAGTGTTCCAGAT
ATTTCTGTTGAAAAACAATACCAAGTCTGATGTTAACACACTTGTGTGTTCATGAGAGCTGACAGTGG
TCTTAAAGGATTAGTGGTACTATTCATAAACAAAAGCTTGTATTAGGAGGAAGTGACAGAATTTTGTAG
GACAGCTATTATAGATATAGATCTTTGCAGCAAGTGTATCAGAGTGAAACTTGTTTTCAAGGAGGATAT
ATTTTAAAAGAATCAAATCCAAAAGCTTCTAAGGTTGGTAGCTGTACCATCCCAAAGTGTCTGACAAC
AGGATAGTGTTGTATCTCTGTATGGCAGTTGAAAATAGGAAAAAAAATACATACATATATGTATGTATATA
TGTGTGTGTGTGTGTGTGTGTATTTTTTTTTTTTTTTTTTTTGAGACAGAGTCTGGCTCTGTCACCCAG
GCTCTGCTCTGGAGTGCAGTGGCACGATCTCAGCTCACGGCAACCTCCATCTCCCAGTTTAAAAGT
GATTCTCCTGCCTCAGCCTCCCAAGTAGTGGGATTACAGCCTGGCTAAGGTGTGGTGGTATGCGCC
TGTAATCGCTTTTAGTGGAGACGGGGTTTCACCACATTGGCCAGGCTGGTCTCAAACTCCTGACTTC
AAGTGATCTGTCTGCCTTGGCCTCCCAAAGTGCTGGGATTACAGGCATGACCTACTGCGCCCGGCG
CTGAAAATAGGAGATTATAAATCCCGAATGACATTATTGAATATTAAACAAAATATTTAGGTTTCATTTTA
TCGGACCTTTAGAGTGGGTGATTATCTGGGTACTCTTACTTTGGTAAGTCCATTGAACAAATCTAAAC
TGGCATCTCCACAAAGGTTCTCATCAAGTGATCAGTCCTCCTTTTTCCTGTAAAATTTTTCTATAAGTG
ACTACACAGTTTTAGCCCTTTTTTAATGAAAAAAAGTTTGTGCACAGATTTGACATACGTGTGAAAAA
GATGAGCTTAGTGAAAGATAGGTGAGAACCTTTTTGTTTTTTACAGATAGTAGAGATGACAACAGAGA
CTTATTTCCATCCACCATTCTACACATGCCAAGATACCAAAGAACCAGTGTCTAGCCTGGTAGCTTGA
CCTTCCACTGCAACCTAAGACAAGTAGTAAAGCCCTTACCTTGAAGTAGGGTCATGGGTGATGCTGT
TGATTACAAGGAGTTCAGAAAGAACTACATGTAAGATTGAAGCCATTCAATAACTACTTATTTAGGATT
TTGTGATTGTGATGTATAATGCCAGACACAGAAGAAAGAATAATCAGATATGAACATGTAATCAGAAGA
GCTTACAGCCTAGCAATGTAGAAGATCTGCTTAGAAATACAATGCAGAAAGTTTAAAATCCTAGTAAAA
ATAGAGGTAAAATGTAGAAAGCTAGTTTCTTTATACAGTGCAACAGTATAGTATTGAGAAAGAAGGAAG
TAATGCAGTTTATGGTGATAAATTCTTTTTGTTTCTCTTGATAGTCTTTTTGTTTGTTTGTTTGTTTGAG
ACAGTTTCCCTCTGTTGCCCAGGCTGGAGTGCAGTGGCAACGATCTTGTCTCACTGCAACCTCCGC
CTCCCAGGTTCAAGCAATTCTCCTGCCTCAGCCTCCTGAGTAGCTGAGATTCCAGGTGCACGCCAC

CATGCCTGGCTAATTTTTTGTGTGTTTTAAGTAGAGATGGGGTTTCTCCATGTTGGTCAGGCTGGTCT
TGAACTCCTGACCTCATGATCTGCCCGCCTTGGCCTCTCAAAGTGCTGGGATCATAAGCATGAACCA
CTGTGCCCGGCCTTCTTGATAGTCTTAAGAGCTTCTCTGTCATATGTCCTATATTCCTGGCTTTCAGC
TTCCTTTCTGCTTACGTGTTGCAGGTGATGCATGTCCTTTGTAATAATCTGGACACTTTTCAGTTGTG
TTTTAAAAAACATCAAACTGGTTTAGGCATTTAGGGAAATTTTATTGGTTTTTATAACTGAAAAGTCCAG
ACTTATCTTTCCATCTCTCAATTCTGTTTTCTAGTGTATGTTGGCTTGGTTTTAGGCTAGGCCTTTTCTA
AATGGTGTAAAGATGGCAACTCTATCTTACTGGCTTATCAACTCAATCAGGAGAAAACTCACTTTTCC
GTAGATTCAGCTGAAGTCCTGAAGCTGGTATTCATTGACTCTGTTAGGCCTCCATAGTGTCTTGTGC
CCACTCCTGGATCAGTTGCTTTGGCTGGAAAATGCTGTATTCTTTTTAGCCTGGCCAGACTTGTATGT
TCATTCCTGGGGCTAGTGGGTGAGGTCAACCCCTTCTGAACTGGTCAGCCTCTTCTGGATTGAGTG
GAGTGGATACCCAAAGGAAAAAAATGAATGTGATGTTTTAGAGAGAGAAATAACAAGAGCTCAATATA
CTTTTTGCATTCCTCTGGAATTTGTGGTATTTTAGGATGGAAGTAGAATGGTGGTCAGCTAACAATAA
GACAATTCTTGGTGTTTTTACATTAACTACAGGTAATAATTAAACTTGTGCGGTAATTACATGGGGAAA
ATCCCTCTCATTAATTTCAATACTAAATATGGTATCTTTCCCTTTTTTTGCATTATTAAGCTTACTGAAATA
TACATAGCAAAAAATAAAATACAGAAGGCCAGGCGCAGTGACTCATGCCTATAATTCCAGCACTTTGG
GAGGCCGCGGTGGGAGGATCACTTGAGGTCAGGAGTTCGAGACCAGCCTGGCCAACATGGTGATA
CCCCGTCTCTACTAAAAATCCAAATAACAGTTCAATTCCTTGAAGCAGTAAAGCTTTGAGCTGCACTG
AGAGGAGACTAGA
