## Supplementary material for "NPlasmid-seq: Cas9-encoded terminal signatures enable molecule-level nanopore quality control of vector and IVT-template plasmids": Supplementary File 2 including supplementary figures and legends V6.0.pdf

### Figure S1

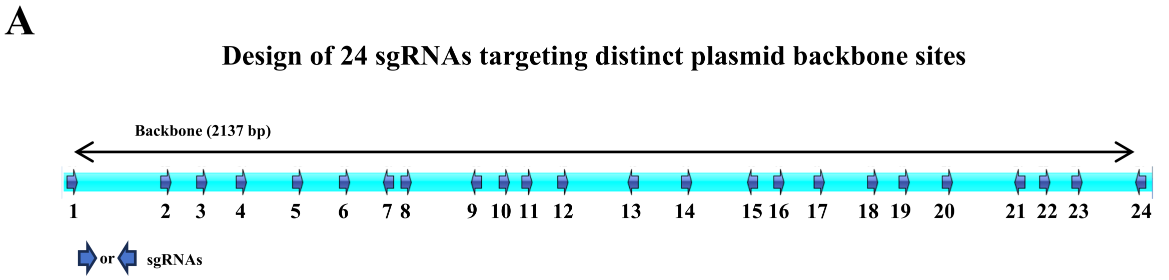


**Figure S1. Design and sequences of 24 sgRNAs targeting distinct sites across the plasmid backbone.**
(A) Schematic showing the design and distribution of 24 sgRNA target sites across the plasmid backbone. The circular plasmid map is expanded into a linear backbone representation, with numbered markers indicating the positions of sgRNAs 1-24. Numbers correspond to the sgRNA entries listed in Table S2.

### Figure S2

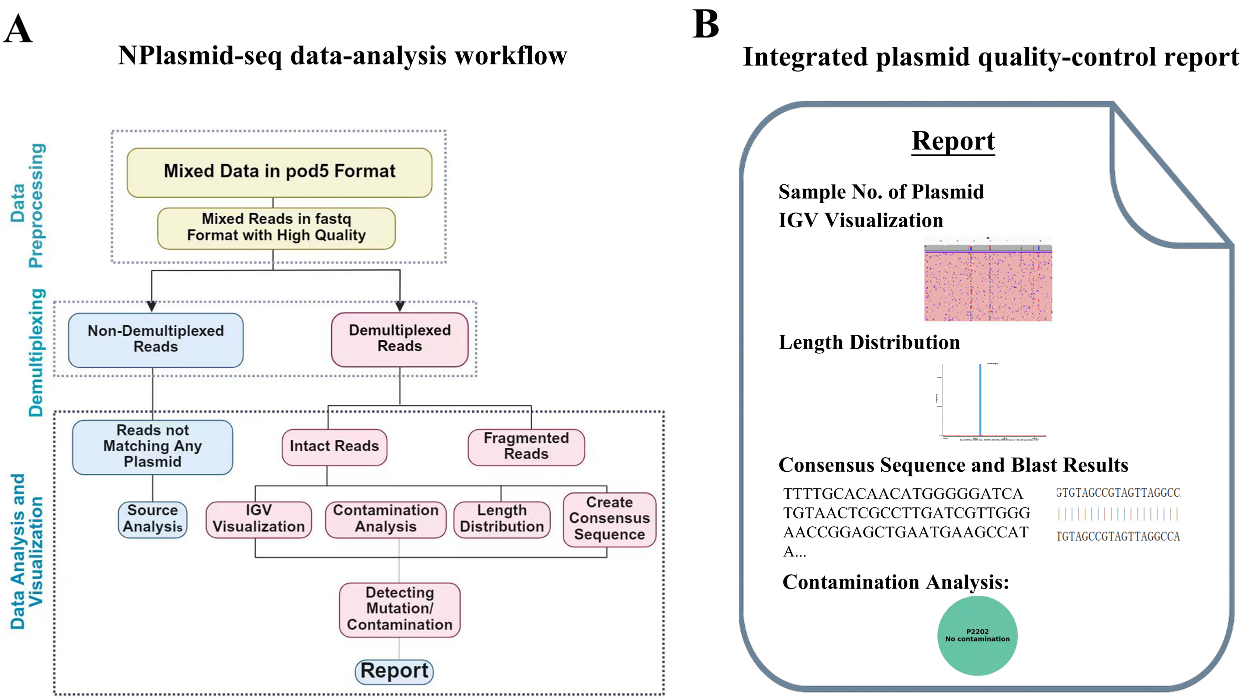


**Figure S2. NPlasmid-seq data-analysis workflow and integrated plasmid quality-control reporting.**
(A) Overview of the NPlasmid-seq data-analysis workflow. Pooled Oxford Nanopore sequencing data in POD5 format are basecalled and quality-filtered to generate high-quality pooled FASTQ reads. Reads are demultiplexed according to the plasmid-specific terminal sequence signatures generated by Cas9 cleavage and separated into assigned and unassigned reads. Unassigned reads undergo source analysis. Demultiplexed reads are further evaluated according to read integrity and subjected to reference alignment and IGV visualization, read-length distribution analysis, consensus-sequence reconstruction, BLAST-based sequence validation, and contamination analysis. (B) Schematic representation of the integrated quality-control report generated for each plasmid sample, summarizing the sample identifier, IGV alignment visualization, read-length distribution, reconstructed consensus sequence and corresponding BLAST results, and contamination assessment.

### Figure S3

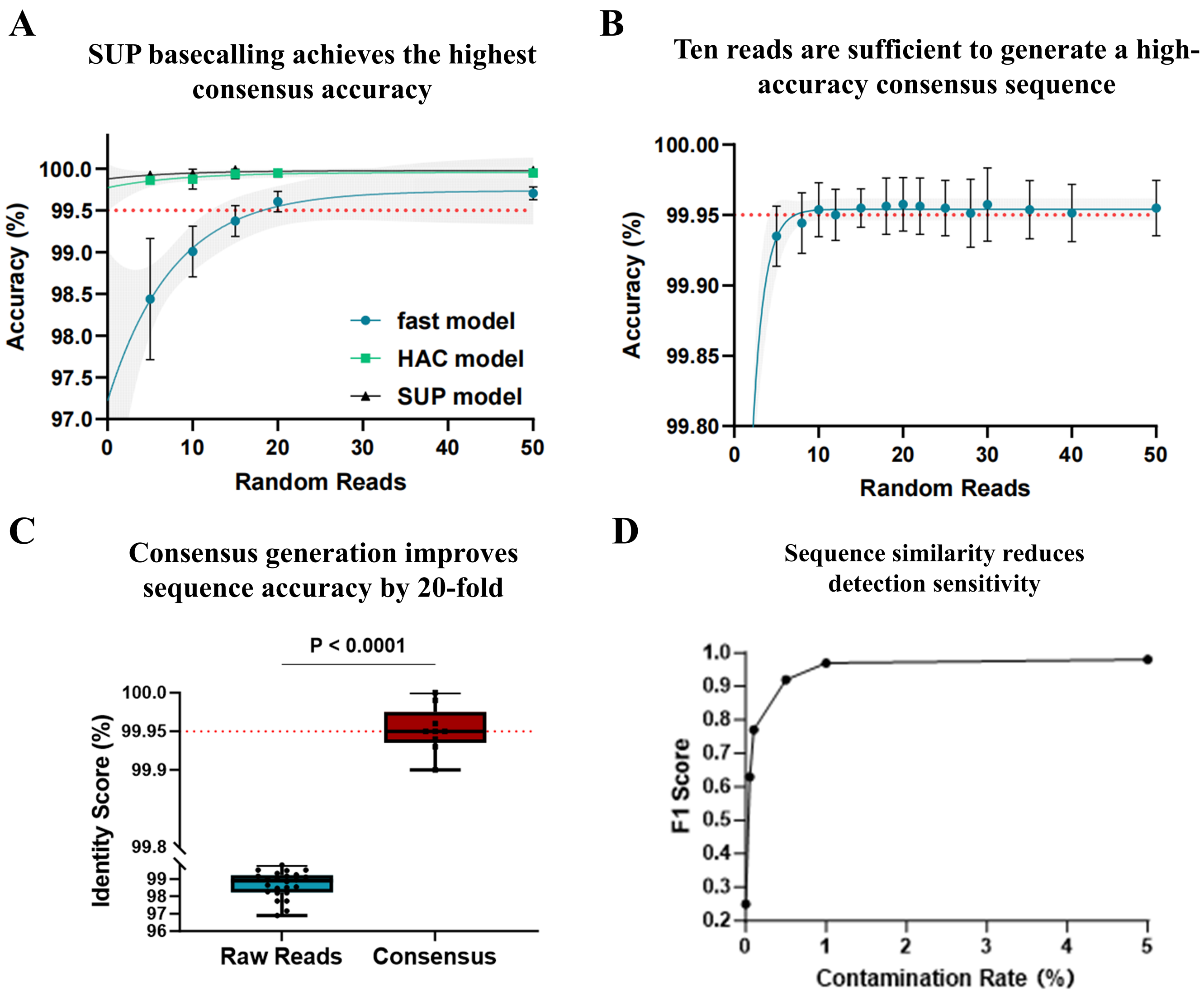


**Figure S3. Optimization of basecalling model and read depth for accurate plasmid consensus generation.**

(A) Comparison of consensus-sequence accuracy obtained using different Oxford Nanopore Technologies basecalling models. The super-accuracy (SUP) model achieved the highest consensus accuracy and was therefore selected for subsequent analyses.

(B) Effect of read number on consensus-sequence accuracy. Reads from each plasmid were randomly subsampled at the indicated depths and used to generate polished consensus sequences. Consensus accuracy reached a stable plateau at approximately 10 reads.

(C) Comparison of sequence accuracy between individual raw nanopore reads and the corresponding polished consensus sequences. Sequence accuracy was calculated as the percentage identity between each read or consensus sequence and its corresponding reference plasmid sequence. Error-reduction fold was calculated as (100 - raw-read accuracy)/(100 - consensus accuracy).

(D) Detection performance for contamination between sequence-similar plasmids at increasing contaminant abundances, analysed with the near-identical plasmid classification rule described in Methods.

### Figure S4

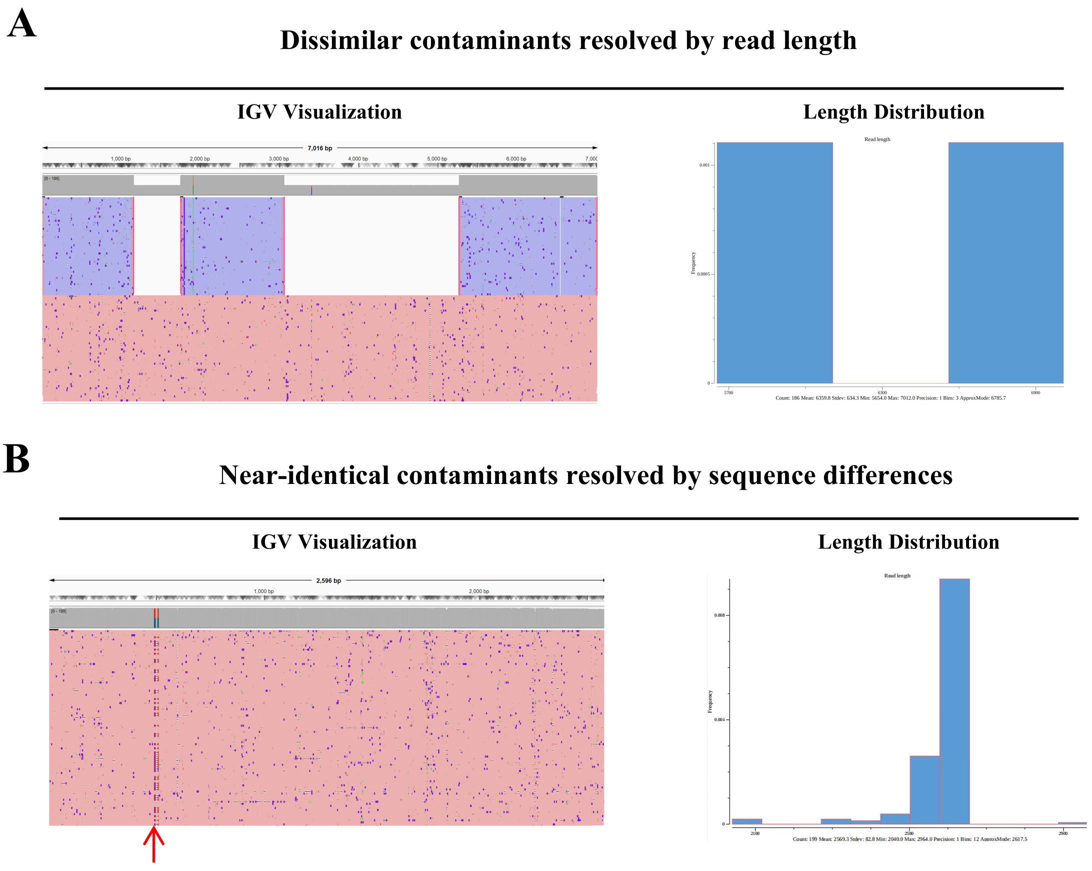


**Figure S4. Visualization and identification of plasmid contaminants and representative sequence variants.**

(A) Representative identification of a dissimilar plasmid contaminant. Read alignments visualized in IGV (left) exhibited distinct alignment patterns between the expected plasmid and the contaminating sequence, while the corresponding read-length distribution (right) showed two clearly separated populations.

(B) Representative identification of a near-identical plasmid contaminant. Because the contaminating sequence was highly similar in length and overall structure to the expected plasmid, the two could not be reliably distinguished by read-length distribution alone (right). IGV visualization (left) revealed a localized mismatch region, indicated by the red arrow, enabling the near-identical contaminant to be resolved on sequence-level differences.

### Figure S5

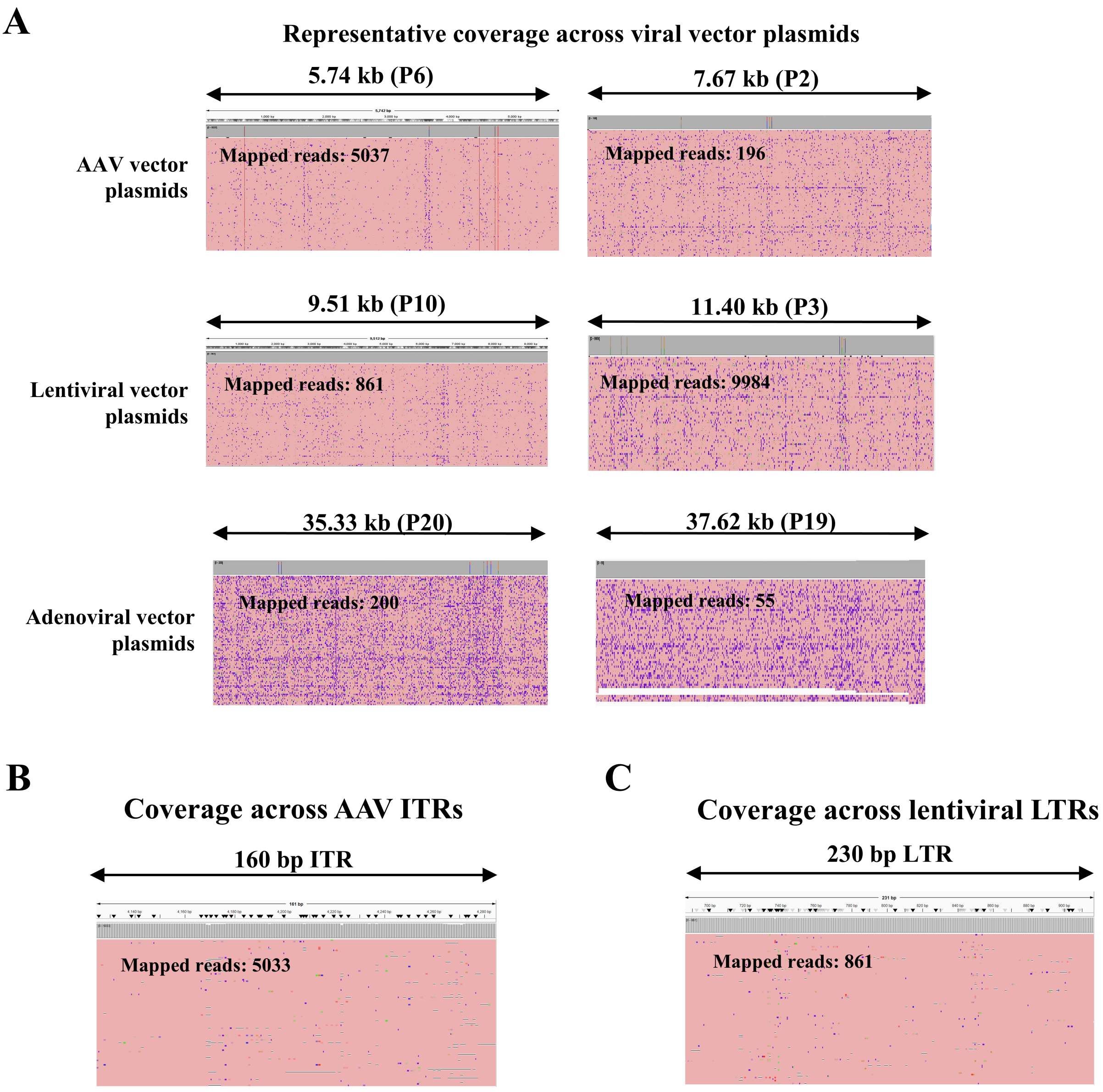


**Figure S5. IGV visualization of viral vector plasmids and repeat-rich terminal regions.**
(A) Representative IGV views of sequencing reads aligned to viral vector plasmid reference sequences, including AAV vector plasmids of 6 and 8 kb, lentiviral vector plasmids of 9 and 11 kb, and adenoviral vector plasmids of 35 and 38 kb.
(B) Enlarged IGV view of a representative 160-bp inverted terminal repeat (ITR) region within an AAV vector plasmid.
(C) Enlarged IGV view of a representative 230-bp long terminal repeat (LTR) region within a lentiviral vector plasmid.

### Figure S6

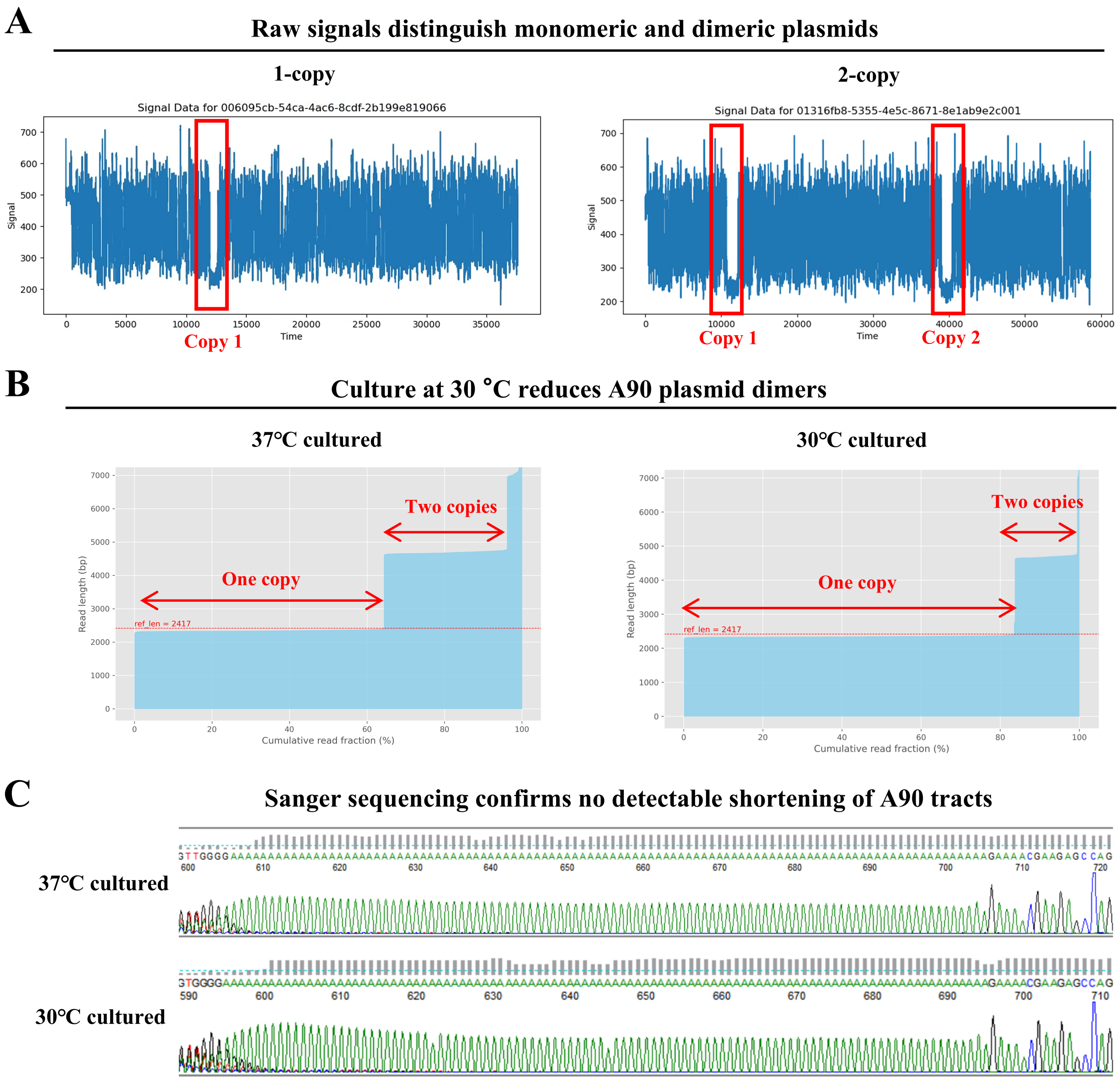


**Figure S6. Nanopore-based detection of A90 plasmid dimers and evaluation of poly(A)-tract stability at different culture temperatures.**

(A) Representative raw nanopore current traces of monomeric and dimeric plasmid molecules containing one or two poly(A) tracts, respectively. Signal shifts corresponding to individual poly(A) tracts are outlined by red boxes.

(B) Representative cumulative read-length distributions of the A90 plasmid propagated at 37 or 30 °C. Reads corresponding to approximately one plasmid length were classified as monomeric, and reads corresponding to approximately twice the plasmid length as dimeric.

(C) Representative Sanger chromatograms spanning the poly(A) tract of A90 plasmids propagated at 37 or 30 °C. The expected 90-nucleotide tract was retained under both conditions.

### Figure S7

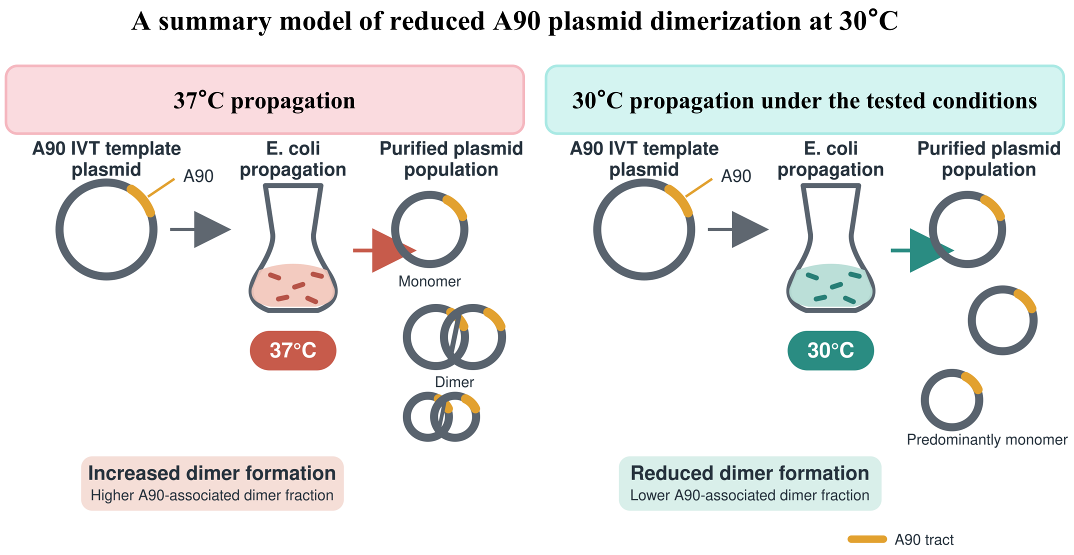


**Figure S7. Schematic summary of the reduction in A90-associated multimeric-read fraction at 30 °C.**

A90-containing IVT-template plasmids were propagated in Escherichia coli at 37 °C or 30 °C under matched conditions. Lower-temperature propagation yielded a lower multimeric-read fraction, as quantified in Fig. 4D,E. Orange segments denote A90 tracts. Monomer and dimer symbols schematically represent the observed shift in read populations; their numbers and sizes do not indicate measured molecular abundances or establish plasmid topology.

### Figure S8

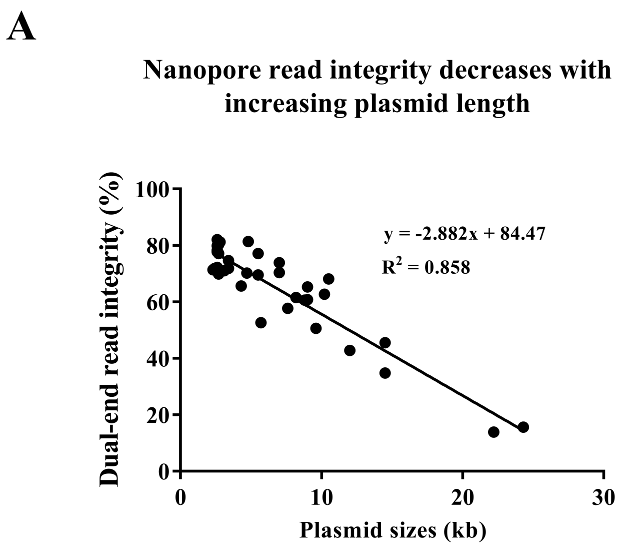


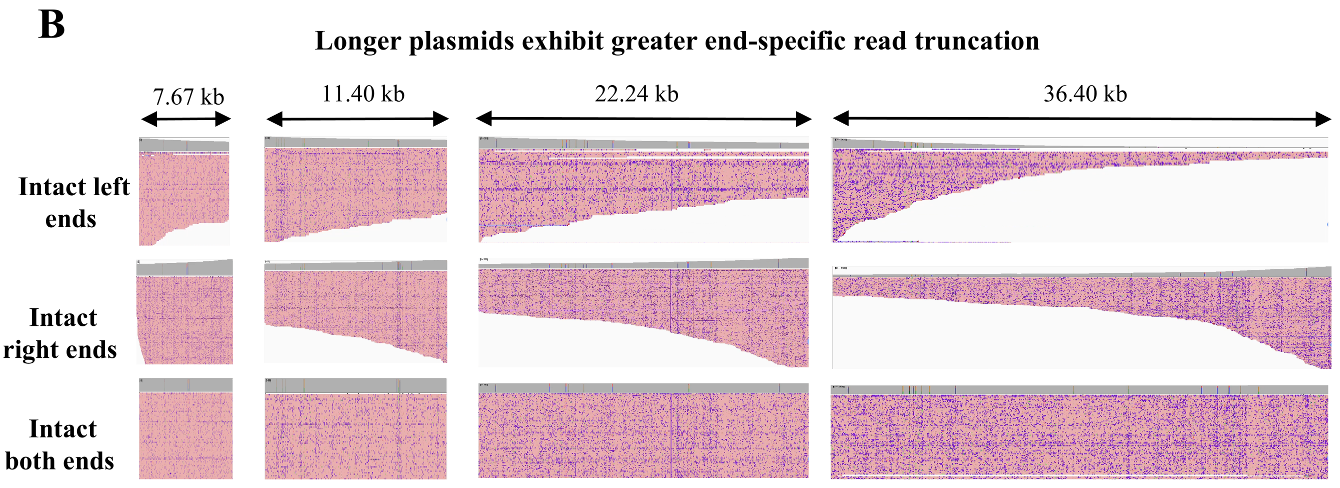


**Figure S8. Dual-end read integrity decreases with increasing plasmid length.**
(A) Relationship between plasmid length and nanopore dual-end read integrity across 37 independently sequenced plasmids spanning 2.28-24.30 kb. Each point represents one plasmid dataset. The solid line indicates the linear regression fit ($y = -2.882x + 84.47$; $R^2 = 0.858$), where x is plasmid length in kilobases and y is dual-end read integrity expressed as a percentage. The 36.40-kb plasmid shown in panel B was not included in this regression analysis.
(B) Representative read-alignment profiles for plasmids of 7.67, 11.40, 22.24 and 36.40 kb. Rows show reads retaining an intact left end, an intact right end or both intact ends. Each horizontal alignment represents an individual read. Double-headed arrows indicate the full reference length of each plasmid.
