## Supplementary material for "NPlasmid-seq: Cas9-encoded terminal signatures enable molecule-level nanopore quality control of vector and IVT-template plasmids": Supplementary File 3 including supplementary tables V6.0.pdf

#### Table S1. Plasmids included in this study.

| ID | Plasmid information | Length (kb) |
|---|---|---|
| P1 | pU6-sgKLRC1b-gN21(Sa-v2) | 2.60 |
| P2 | pAAV-Alb2(SA85-0)-P2A-BDDF8-N6-mir122T-15(S1002FF) | 7.67 |
| P3 | pRSC-SFFV-Cas9NG-E2A-Puro-wpre(Sp-NG)-(SpG) | 11.40 |
| P4 | pAdHM4-SFFV-OCT4Cherry-cutted | 22.24 |
| P5 | pAdHM4-SFFV-M2K | 36.40 |
| P6 | pAAV-CAG-mNeonGreen-Wpre-polyA | 5.74 |
| P7 | pAAVD-COL1A1(HA)-P2A-SOX9-T2A-mNeonGreen-wpre(sgDocut) | 7.02 |
| P8 | pAAVD-Grin1(800)-CAG-Crimson-PolyA(rbG)-Grin1(950) | 7.19 |
| P9 | pRSC-COL1A1-raSOX9-T2A-Puro-Wpre | 9.03 |
| P10 | pRSC-EF1-raSOX9-T2A-Puro-wpre | 9.51 |
| P11 | pU6-sgKLRC1a-gN21(Sa-v2) | 2.60 |
| P12 | pU6-sgKLRC1c-gN21(Sa-v2) | 2.60 |
| P13 | pIVT-T7-Nco-U3.Hba-A10(Gaaaac) | 2.40 |
| P14 | pIVT-T7-Nco-U3.Hba-A20(Gaaaac) | 2.40 |
| P15 | pIVT-T7-Nco-U3.Hba-A30(Gaaaac) | 2.40 |
| P16 | pIVT-T7-Nco-U3.Hba-A60(Gaaaac) | 2.40 |
| P17 | pIVT-T7-Nco-U3.Hba-A90(Gaaaac) | 2.40 |
| P18 | pIVT-T7-Nco-U3.Hba-A120(Gaaaac) | 2.40 |
| P19 | pAdHM4-CMV-M2K | 37.62 |
| P20 | pAdHM4-SFFV-Oct4Cherry | 35.33 |

### Table S2. Sequences of the 24 backbone-targeting sgRNAs.

| Number | sgRNA ID | Spacer sequence (without PAM) | Length (nt) |
|---|---|---|---|
| 1 | sgPlasBB-CS119-F | TGCTTCAATAATATTGAAAA | 20 |
| 2 | sgPlasBB-CS210-F | TTTTGCTCACCCAGAAACGC | 20 |
| 3 | sgPlasBB-CS280-F | ATCGAACTGGATCTCAACAG | 20 |
| 4 | sgPlasBB-CS357-F | TAAAGTTCTGCTATGTGGCG | 20 |
| 5 | sgPlasBB-CS466-F | ACAGAAAAGCATCTTACGGA | 20 |
| 6 | sgPlasBB-CS558-F | GACAACGATCGGAGGACCGA | 20 |
| 7 | sgPlasBB-CS629-R | GGTATGGCTTCATTCAGCTC | 20 |
| 8 | sgPlasBB-CS678-F | CACCACGATGCCTGTAGCAA | 20 |
| 9 | sgPlasBB-CS801-R | AATAAACCAGCCAGCCGGAA | 20 |
| 10 | sgPlasBB-CS869-F | GCGGTATCATTGCAGCACTG | 20 |
| 11 | sgPlasBB-CS914-F | TCGTAGTTATCTACACGACG | 20 |
| 12 | sgPlasBB-CS983-F | GTGCCTCACTGATTAAGCAT | 20 |
| 13 | sgPlasBB-CS1106-R | TGGAACGAAAACTCACGTTA | 20 |
| 14 | sgPlasBB-CS1224-F | AAAACCACCGCTACCAGCGG | 20 |
| 15 | sgPlasBB-CS1339-R | CTACAGAGTTCTTGAAGTGG | 20 |
| 16 | sgPlasBB-CS1405-F | TTACCAGTGGCTGCTGCCAG | 20 |
| 17 | sgPlasBB-CS1483-F | CAGCGGTCGGGCTGAACGGG | 20 |
| 18 | sgPlasBB-CS1587-F | GCTTCCCGAAGGGAGAAAGG | 20 |
| 19 | sgPlasBB-CS1647-F | GCACGAGGGAGCTTCCAG | 18 |
| 20 | sgPlasBB-CS1733-F | TGTGATGCTCGTCAGGGGGG | 20 |
| 21 | sgPlasBB-CS1860-R | GGTATCAGCTCACTCAAAGG | 20 |
| 22 | sgPlasBB-CS1923-F | CGAGTCAGTGAGCGAGGAAG | 20 |
| 23 | sgPlasBB-CS1985-F | GGCCGATTCATTAATGCAGC | 20 |
| 24 | sgPlasBB-CS2096-R | ATTCCACACAACATACGAGC | 20 |

Spacer lengths of 18-20 nt were permitted by design; the variation is intentional.

**Table S3. Primers used for colony genotyping of the P9 93-bp deletion.**

| Primer name | Direction | Sequence (5′-3′) | Expected amplicon |
|---|---|---|---|
| Q1295-raSOX9-D90seq-F | Forward | GACCTTCGACGTCAACGAGT | 548 bp intact / 455 bp deletion |
| Q1339-raSOX9-D90seq-R | Reverse | AGGTGGAGTAGAGGCTGGAG | 548 bp intact / 455 bp deletion |

**Table S4. Per-sample read statistics for pooled runs.**

Read counts and percentages of passing reads classified by terminal-signature matching.

| Read category | Run-N584 | Run-N585 | Run-N586 | Combined |
|---|---|---|---|---|
| Total reads | 422,794 | 474,664 | 327,534 | 1,224,992 |
| Passing reads | 422,790 | 474,658 | 327,532 | 1,224,980 |
| Assigned | 102,683 (24.29%) | 65,327 (13.76%) | 135,651 (41.42%) | 303,661 (24.79%) |
| Single-end | 233,338 (55.19%) | 263,415 (55.50%) | 148,329 (45.29%) | 645,082 (52.66%) |
| Conflicting | 14,479 (3.42%) | 13,678 (2.88%) | 5,875 (1.79%) | 34,032 (2.78%) |
| Single-end + conflicting | 247,817 (58.61%) | 277,093 (58.38%) | 154,204 (47.08%) | 679,114 (55.44%) |
| Unmatched | 72,290 (17.10%) | 132,238 (27.86%) | 37,677 (11.50%) | 242,205 (19.77%) |

### Table S5. Physical DNA mixture design and per-sample results.

| Sample | Pool | Guide | Input P6 (mol%) | Reads assigned | P6 | P7 | Ambiguous | Observed P6 (%) |
|---|---|---|---|---|---|---|---|---|
| V01 | Pool1 | Syn264 | 0 | 1,711 | 0 | 1,350 | 361 | 0.0000 |
| V02 | Pool1 | Syn288 | 0 | 5,423 | 0 | 4,321 | 1,102 | 0.0000 |
| V03 | Pool1 | Syn289 | 0 | 9,613 | 0 | 7,663 | 1,950 | 0.0000 |
| V04 | Pool1 | Syn266 | 0.5 | 8,273 | 20 | 6,545 | 1,708 | 0.3046 |
| V05 | Pool1 | Syn293 | 0.5 | 4,017 | 29 | 3,224 | 764 | 0.8915 |
| V06 | Pool1 | Syn290 | 0.5 | 34,051 | 57 | 26,796 | 7,198 | 0.2123 |
| V07 | Pool1 | Syn291 | 1 | 4,938 | 11 | 3,961 | 966 | 0.2769 |
| V08 | Pool1 | Syn263 | 1 | 1,921 | 3 | 1,488 | 430 | 0.2012 |
| V09 | Pool1 | Syn292 | 1 | 32,736 | 101 | 25,524 | 7,111 | 0.3941 |
| V01 | Pool2 | Syn264 | 0 | 703 | 0 | 546 | 157 | 0.0000 |
| V02 | Pool2 | Syn288 | 0 | 1,575 | 0 | 1,280 | 295 | 0.0000 |
| V03 | Pool2 | Syn289 | 0 | 3,001 | 0 | 2,438 | 563 | 0.0000 |
| V04 | Pool2 | Syn266 | 0.5 | 4,886 | 39 | 3,825 | 1,022 | 1.0093 |
| V05 | Pool2 | Syn293 | 0.5 | 1,432 | 18 | 1,140 | 274 | 1.5544 |
| V06 | Pool2 | Syn290 | 0.5 | 25,912 | 211 | 20,239 | 5,462 | 1.0318 |
| V07 | Pool2 | Syn291 | 1 | 2,447 | 24 | 1,926 | 497 | 1.2308 |
| V08 | Pool2 | Syn263 | 1 | 1,495 | 18 | 1,155 | 322 | 1.5345 |
| V09 | Pool2 | Syn292 | 1 | 23,876 | 328 | 18,336 | 5,212 | 1.7574 |
| V10 | Pool3 | Syn240 | 5 | 10,766 | 706 | 7,864 | 2,196 | 8.2380 |
| V11 | Pool3 | Syn266 | 5 | 10,790 | 771 | 7,751 | 2,268 | 9.0472 |
| V12 | Pool3 | Syn293 | 5 | 6,398 | 361 | 4,825 | 1,212 | 6.9610 |
| V13 | Pool3 | Syn294 | 50 | 18,948 | 11,086 | 4,331 | 3,531 | 71.9076 |
| V14 | Pool3 | Syn241 | 50 | 22,385 | 12,471 | 5,679 | 4,235 | 68.7107 |
| V15 | Pool3 | Syn242 | 50 | 13,108 | 7,237 | 3,574 | 2,297 | 66.9411 |
| V16 | Pool3 | Syn262 | 100 | 35,197 | 28,383 | 0 | 6,814 | 100.0000 |
| V17 | Pool3 | Syn265 | 100 | 18,059 | 14,637 | 2 | 3,420 | 99.9863 |

Each final P6/P7 mixture was prepared separately in its own tube; no mixture was split into aliquots. Tubes within a pool and input level shared an upstream P6 working solution; the two pools used separate working solutions. Observed P6 (%) = 100 x P6 / (P6 + P7); ambiguous reads are excluded from this denominator and reported separately. Group means: 0.834% at 0.5 mol%, 0.899% at 1 mol%, 8.082% at 5 mol% and 69.186% at 50 mol%. Pure-plasmid controls: 0 P6 calls among 17,598 classified reads in the P7-only samples; 2 P7 calls among 43,022 classified reads in the P6-only samples.

### Table S6. Per-colony genotyping results for the P9 deletion.

| Colony ID | Double peaks | Genotyping classification | Deletion sequence detected |
|---|---|---|---|
| P2207A | No | 93-bp deletion | Yes |
| P2207B | Yes | Double-peak profile | Yes |
| P2207C | Yes | Double-peak profile | Yes |
| P2207D | Yes | Double-peak profile | Yes |
| P2207E | No | Intact | No |
| P2207F | Yes | Double-peak profile | Yes |
| P2207G | No | 93-bp deletion | Yes |
| P2207H | No | 93-bp deletion | Yes |
| P2207I | No | Intact | No |
| P2207J | Yes | Double-peak profile | Yes |
| P2207K | No | 93-bp deletion | Yes |
| P2207L | No | 93-bp deletion | Yes |
| P2207M | No | 93-bp deletion | Yes |
| P2207N | Yes | Double-peak profile | Yes |
| P2207O | Yes | Double-peak profile | Yes |
| P2207P | No | Intact | No |
| P2207Q | No | 93-bp deletion | Yes |
| P2207R | No | 93-bp deletion | Yes |
| P2207S | Yes | Double-peak profile | Yes |
| P2207T | Yes | Double-peak profile | Yes |
| P2207U | No | 93-bp deletion | Yes |
| P2207V | Yes | Double-peak profile | Yes |
| P2207W | No | 93-bp deletion | Yes |
| P2207X | Yes | Double-peak profile | Yes |

Totals: 93-bp deletion, 10/24 (41.7%); intact, 3/24 (12.5%); double-peak, 11/24 (45.8%). Scored as carrying detectable deletion sequence, 21/24 (87.5%). Counts verified against Fig. 3E.

### Table S7. Per-subsample results of the computational mixture benchmark.

| Contaminant fraction (%) | Subsamples | Detected (>=3 reads) | Precision | Contaminant reads: geometric mean | min | max | Mean recall | Mean ambiguous |
|---|---|---|---|---|---|---|---|---|
| 0.1 | 20 | 13/20 | 1.000 | 3.0 | 1 | 5 | 0.650 | 0.33 |
| 0.2 | 20 | 20/20 | 1.000 | 6.0 | 3 | 8 | 0.620 | 0.33 |
| 0.5 | 20 | 20/20 | 1.000 | 16.0 | 11 | 20 | 0.648 | 0.33 |
| 1.0 | 20 | 20/20 | 1.000 | 33.7 | 26 | 39 | 0.677 | 0.33 |
| 5.0 | 20 | 20/20 | 1.000 | 160 | 144 | 175 | 0.640 | 0.33 |

All 100 subsamples used 5,000 passing reads. Precision was 1.000 in every subsample: no read was called as contaminant in error among 493,200 declared reads in total, giving a one-sided 95% upper bound on the per-read false-call rate of 6.1 x 10-6. Recall is bounded by the ambiguous-read convention, under which a mean of 33% of reads in each subsample carried insufficient class-specific evidence. The full per-subsample table (level, subsample, counts, TP/FP/FN/TN, precision, recall, F1, classifier version) is provided as a supplementary data file.
